# EphA2 and Phosphoantigen-Mediated Selective Killing of Medulloblastoma by γδT Cells Preserves Neuronal and Stem Cell Integrity

**DOI:** 10.1101/2024.10.12.617193

**Authors:** Lola Boutin, Mingzhi Liu, Julie Déchanet Merville, Oscar Bedoya-Reina, Margareta T Wilhelm

## Abstract

Medulloblastoma (MB) is a pediatric brain tumor that develops in the cerebellum, representing one of the most common malignant brain cancers in children. Standard treatment includes surgery, chemotherapy, and radiation, but despite a 5-year survival rate of approximately 70%, these therapies often lead to significant neurological damage in the developing brain. This underscores the urgent need for less toxic, more effective therapeutic alternatives. Recent advancements in cancer immunotherapy, including immune checkpoint inhibitors and CAR-T cell therapy, have revolutionized cancer treatment. One promising avenue is the use of Gamma Delta (γδ)T cells, a unique T cell population with potential advantages such as non-alloreactivity, potent tumor cell lysis, and broad antigen recognition. However, their capacity to recognize and target MB cells remains underexplored. To investigate the therapeutic potential of γδT cells against MB, we analyzed the proportion and status of MB-infiltrated γδT cells within patient datasets. We next investigated the expression of γδT cell ligands on MB cells and identified EphA2 receptor and the phosphoantigen/Butyrophilin complex as key ligands, activating Vγ9Vδ1 and Vγ9Vδ2 T cells, respectively, leading to significant MB cell lysis in both monolayer and spheroid models. Importantly, preliminary safety data showed that γδT cells did not target differentiated neurons or neuroepithelial stem cells derived from induced pluripotent stem cells, underscoring the selectivity and safety of this approach. In conclusion, γδT cells trigger an efficient and specific killing of MB, and would offer a promising novel therapeutic strategy.

**Key messages:** Medulloblastoma patients often experience significant long-term side effects from current standard treatments. Immunotherapy has emerged as a promising alternative to conventional therapeutic approaches. In our study, we demonstrated that γδT cells can efficiently and specifically target medulloblastoma cells without causing harm to healthy neuronal tissue. These findings suggest that γδ T cell therapy may provide therapeutic benefits while potentially reducing treatment-related toxicity in medulloblastoma patients.

## Introduction

Medulloblastoma (MB), the most common malignant pediatric tumor of the central nervous system (CNS), accounting for 20% of all pediatric CNS tumors, represents a heterogeneous group of brain tumors usually found in cerebellum. Current treatment protocols have improved the 5-year-overall survival (5y-OS) rate to 80% globally^1^. Four main subgroups of MB have been identified by molecular classification: Wingless (WNT), Sonic Hedgehog (SHH), Group (Grp)3, and Group (Grp)4^2^. Prognosis is strongly linked to the MB subgroup, with WNT-activated MB having a more favorable outcome (90% 5y-OS) and Grp3, the less favorable (50% 5y-OS) with a high metastatic rate^3^. MB patients are treated according to a standard of care, which includes surgical resection, adjuvant chemotherapy and, in some cases, craniospinal irradiation. However, the treatment line often leads to systematic and irreversible neurological deficits, resulting in a decline in intellectual and cognitive function, due to damage of the healthy brain tissue^4^. Highlighting the importance of identifying less toxic and more effective therapeutic alternatives for MB.

Recently, immunotherapy has paved the way to propose more specific and safer alternatives to conventional chemotherapies. MB is known as a cold tumor with an immune evasive microenvironment, mostly limited to M2-like microglia and macrophage infiltration^5^. In addition, MB is characterized by a low mutation burden and the absence of PD-L1 expression, which correlates with low T cell infiltration^6^. While immune checkpoint-blockade (ICB) therapy (e.g. PD1/PD-L1, CTLA-4) has led to a major breakthrough for many solid tumors, MB does not seem to benefit from it^7^. Adoptive cell therapies such as CAR-T or NK cell therapy are currently being evaluated in various clinical trials for pediatric brain tumors including MB patients^8,9^, with CAR-T cells targeting B7-H3, GD2, IL-13α, EphA2 or HER2 showing some promise^8^. However, tumor cells can escape CAR-T cells by downregulating the expression of the target molecules, as has been demonstrated in 30-70% of patients undergoing therapy^10^.

γδT cells are a MHC-peptide unrestricted T cell population which recognizes conserved cellular stress patterns upregulated in infected and transformed cells^11^. They possess both adaptive and innate receptors such as a functional T cell receptor (TCR) associated with a CD3 molecule, as well as NKG2D and Toll-like receptors (TLR), which enable them to recognize a broad spectrum of ligands^12^. γδT cells express a TCR composed of Vγ and Vδ chains, the former defining the γδT subpopulation (Vδ1, Vδ2 and Vδ3), and the preferred location. Invariant Vγ9Vδ2 T cells are the main population in the peripheral blood system (>90% of total γδT) sensing phosphoantigen (pAgs) level dysregulation in transformed or infected cells, whereas Vδ1 and Vδ3 subpopulations are mainly present in tissues where they are involved in immune surveillance but also in the maintenance of tissue homeostasis^13^.

Allogeneic therapies based on γδT cells have been shown to be clinically safe and effective in various cancers, demonstrating the potential of using an allogeneic γδT cell bank for tumor immunotherapy^14– 16^. However, the ability of γδT cells to target and eliminate MB cells is poorly documented. Here, we show that both Vγ9Vδ1T and Vγ9Vδ2T cells can specifically eliminate MB cells without affecting healthy neural stem cells or neurons, highlighting the potential of using γδT cells for immunotherapy of medulloblastoma and other pediatric CNS tumors.

## Materials & Methods

### Deconvolution of bulk MB tumor samples and TIL abundance

Bulk RNA sequencing datasets were downloaded from GEO portal: Normal cerebellum (GSE44971) and MB patients (GSE37418 and GSE85217). Assessments of leucocyte fractions from the specified transcriptomes were performed by applying CIBERSORT (https://cibersort.stanford.edu/) with the matrix LM-7 as previously described^17,18^. Abundances were calculated from the CIBERSORT results and Sample Enrichment Score (SES). SES were computed by applying the open-source software AutoCompare-SES (https://sites.google.com/site/fredsoftwares/products/autocompare_ses) with normalized settings. Then, the open-source software DeepTIL (https://sites.google.com/site/fredsoftwares/products/deeptil) was used to automatically compute the abundance of the seven leucocyte subsets.

### TCR repertoire analysis

To conduct the TCR repertoire analysis, we used the MB immune cell landscape dataset obtained by Riemondy et al. using single-cell sequencing^6^. Briefly, 29 samples were single-cell sequenced using 10X to a deep of 50,000 reads per cells, and processed using CellRanger and Seurat. The resulting dataset contains normalized expression for 4,669 MN tumor-infiltrating immune cells including clusters of microglia, myeloid, neutrophil, NK, T, and B cells. This data was downloaded from the pediatric Neuro-oncology Cell Atlas (pneuroonccellatlas.org) available at https://github.com/rnabioco/medulloblast on 08/09/2023. The available metadata from Supplementary Table 1 of Riemondy et al. was included in the analysis. To study the γδ repertoires in T cells of MB, BAM files from single-cell alignments of the 29 samples analyzed were obtained from the GEO SuperSeries accession number GSE156053 on 09/12/2023. These files were further processed with TRUST4, to determine the repertoires for each cell independently, following their barcodes^19^.

### T cell differentiation state analysis

To determine the differentiation state of T cells, gene signatures for 1) naïve, 2) resident, 3) pre-exhausted, 4) exhausted, and 5) effector-memory cells were generated **(Table S1**). For each of these gene lists a signature score was computed as the average expression of each reference gene sets, minus the average of a (n) randomly selected set of genes (where n = max [# reference genes, 50]). The signature score provides an estimation of the transcriptional resemblance of each cell to each reference cluster of interest. Additionally, the normalized expression for each gene within these signatures was obtained.

### Cell culture

Neuroepithelial stem (NES) cells were obtained as previously described^20^. NES cells were cultured in flasks coated with 20 µg/ml poly-L-ornithine (Sigma, P3655) and 1 µg/ml laminin (Sigma, L2020) in complete neural stem cell medium (DMEM/F12+Glutamax (ThermoFisher, 31331093) supplemented with 1% N2 supplement (ThermoFisher, 17502001), 0.1% B27 supplement (ThermoFisher, 17504044), 10 ng/ml FGF2 (Qkine, Qk053), 10 ng/ml EGF (PeproTech, AF100-15) and 1% penicillin-streptomycin (Sigma, P4333)). For neuron differentiation, NES cells were seeded in flat bottom 96 well-plates coated with 20 µg/ml poly-L-ornithine and 2 µg/ml laminin at a concentration of 15×10^3^ cells/well in complete NES media. After 24 hours, NES media was removed and replaced with neuron differentiation media (DMEM/F12+Glutamax with 1% N2 supplement, 0,1% B27 supplement and 1 µg/ml laminin). NES cells were maintained in neuron differentiation media for a minimum of 3 weeks to ensure neuronal differentiation.

CHLA-01-MED and CHLA-01R-MED cells were cultured in DMEM/F12+Glutamax with 20 ng/ml, 20 ng/ml EGF, 2% B27. DAOY and ONS-76 cells were maintained in DMEM (ThermoFisher, 41966052) supplemented with 10% FBS (Hyclone, SV30160.03HI). Additionally, medium for DAOY cells contains 1% MEM non-essential amino acids, 1% HEPES, and 1% Glutamax (ThermoFisher, cat. 11140050, SH30237.01, and 35050061, respectively). UW228-3 cells were cultured in RPMI-1640+Glutamax (ThermoFisher, 61870044) supplemented with 10% FBS. D425 and D458 cells were cultured in DMEM/F12+Glutamax supplemented with 10% FBS. All cell lines tested negative for mycoplasma and were authenticated by STR analysis at Eurofins (DAOY, D425, D458 CHLA-01-MED, CHLA-01R-MED), or Multiplexion (ONS-76, UW228-3). Grp3-MB Patient-Derived Xenograft (PDX) MB-LU-181 was established and cultured as described^21^. Jurkat JRT3 expressing Vγ9Vδ1-MAU TCR were obtained as described and cultured in RPMI-1640+Glutamax supplemented with 10% FBS^22^.

### Sorting and *ex-vivo* expansion of γδT cells

Human peripheral blood mononuclear cells (PBMCs) were isolated from blood of healthy donors obtained from Karolinska University Hospital (Stockholm, Sweden). PBMCs were collected by Ficoll gradient centrifugation (Ficoll-Paque® PREMIUM, Cytiva, 17-5442-02) and resuspended in RPMI-1640 medium supplemented with 10% heat-inactivated FBS. γδT cells were magnetically sorted using EasySep™ Human Gamma/Delta T Cell Isolation Kit (Stemcell Technologies, 19255) according to the manufacturer’s protocol. Sorted fraction was expanded by activation with tetramer CD2/CD3/CD28 (Immunocult®, Stemcell Technologies, 10970) in ImmunoCult-XF T Cell Exp Medium (Stemcell Technologies, 10981) supplemented with 300IU/ml recombinant human IL-2 (Stemcell Technologies, 78036) for 14 days. Expanded γδT cell purity was assessed by flow cytometry by staining with anti-pan γδ TCR mAb and the expanded populations were kept for further experiments if γδT > 85%. After expansion, γδT cells were maintained in ImmunoCult-XF T Cell Exp Medium supplemented with 300IU/ml recombinant human IL-2 for up to three weeks.

### Flow cytometry based-T cell activation

For CD107a surface mobilization assays, target cells were treated for 18 hours with zoledronic acid (Sigma, SML0223) in their maintenance media. Target cells were co-cultured for 4 hours with amplified γδT cells (E/T ratio 1:1) in RPMI-1640 supplemented with 10% FBS containing Golgi Stop (BD Biosciences, 554724) and anti-CD107a mAb for 4 hours. Cells were harvested and stained with anti-pan γδ TCR, anti-Vδ2 and anti-Vδ1 mAbs. For CD69 expression, target cells were co-cultured with JRT3-MAU cells (E/T ratio 1:1) in RPMI-1640 supplemented with 10% FBS for 4 hours. Cells were harvested and stained with anti-Vδ1 and anti-CD69 mAbs. Flow cytometry data were acquired using BDCanto II cytometer (BD Biosciences) and analyzed using FlowJo v.10 software (Treestar).

### Flow cytometry staining

All antibodies used for flow cytometry assays are described in supplemental materials.

### LDH-release cytotoxicity assay

For monolayer killing assay, targets cells were seeded in flat-bottom 96-well plates in their maintenance media one day prior to co-culture. For spheroid killing assay, target cells were seeded in U-bottom 96-well low adherence plates (BRANDplates®, inertGrade™) 4 days prior co-culture in NES media. Target cells were treated for 18 hours with zoledronic acid, followed by co-culture with amplified γδT cells (E/T ratio 10:1) in RPMI-1640 supplemented with 5% Human serum (Sigma, H5667) for 8 hours. Supernatants were collected and used for LDH (lactate dehydrogenase) measurement using CytoTox 96® kit (Promega, G1780) according to the manufacturer’s protocol, and quantified by FLUOstar Omega plate reader (Absorbance 490nm, BMG labtech). The percentage of target cell lysis was calculated as follows: ((experimental release - spontaneous release)/ (maximum release – spontaneous release)) × 100. Spontaneous and maximum release values were determined by adding either medium or lysis buffer (provided in the kit) to target cells without T cells. For NKG2D blocking assay, 10 μg/mL of anti-NKG2D mAb was added to the γδ T cells 20 minutes prior co-coculture and kept throughout the experiment.

### Statistical analysis

Samples collected from anonymized healthy human donors have been used in this study. Data were analyzed with GraphPad Prism software v.10.1.2 and are presented as mean ± SD; or survival curve. Statistical analysis data was performed by unpaired nonparametric t-student test; log-rank test; or by two- and one-way ANOVA followed by Tukey, Sidák, or two-tailed Dunnett’s tests to correct for multiplicity. *P*-values below 0.05 were considered statistically significant.

## Results

### γδT cells are predicted to be present in healthy cerebellum and in MB tumors

The presence of γδT cells in healthy CNS and in CNS tumors has been poorly reported. Moreover, the difficulty of transcriptionally separating γδT from CD8^+^ αβT and NK cells makes it difficult to predict their proportion from tissue or tumor mRNA samples^23^. Recently, an updated version of an immunosignature (LM-7) CIBERSORT deconvolution matrix optimized for γδT cells was developed^17^. Using the LM-7 matrix, we estimated the immune infiltration, comparing healthy cerebellum from five fetal and four adult tissues (GSE44971) with 76 MB patients (GSE37418). As previously described, we found that the immune landscape of MB is not different from that of non-tumoral tissue (**fig.1A, Table S2**)24. In particular, we found no significant differences in the proportion of different immune cell populations between fetal/adult cerebellum and MB (two-side Kruskal-Wallis, *p*>0.05) for the predictions of CIBERSORT and SES+CIBERSORT, with the solely exception of CD4^+^ T cells that present a marginal significant difference between fetal and malignant tissues (two-side Mann-Whitney U, *p*=0.013) for the CIBERSORT prediction (**Fig. S1A**). The abundance value was similar to previous reports in which MB had the lowest immune cell abundance of all tumors, including other brain tumors such as ependymoma (7.43), glioblastoma (15.75) or pilocytic astrocytoma (22.29), making MB one of the most immune-evasive tumors^17^.

**Figure 1:**
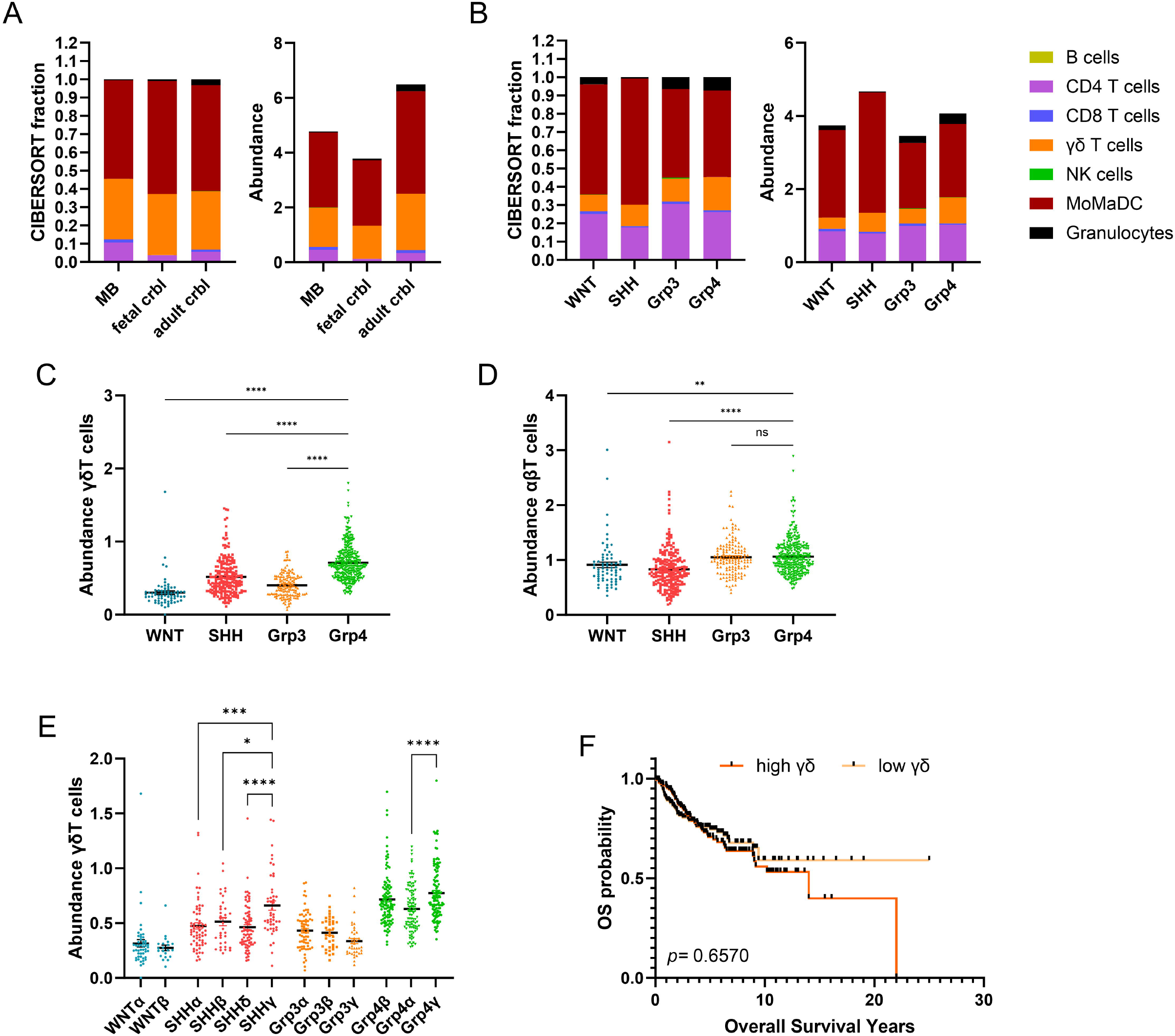
γδT cells abundance in MB tumors. (**A**) Immune cell infiltration and abundance in MB (n=76) versus normal fetal (n=5) and adult (n=4) cerebellum samples. (**B**) Immune cell infiltration and abundance in MB subgroups (WNT-MB n=70; SHH-MB n=222; Grp3-MB n=143; Grp4-MB n=325). γδT and (**D**) αβT cells abundance across MB subgroups. Statistical analysis was performed using one-way ANOVA followed by Dunnett tests to correct for WNT-MB, SHH-MB, and Grp3-MB vs Grp4-MB (**P = 0.0037; ****P < 0.0001). (**E**) γδT abundance across MB subtypes. (**F**) Overall survival of patients grouped according to high or low abundance of γδT cells in MB all subgroups (1^st^ and last quartile, n=306 per group). Significance was determined by log-rank test – nonsignificant differences are not displayed in the figure.

To further analyze the differences in immune cell infiltration, and specifically γδT cell infiltration between the different MB subgroups, we used a larger MB patient cohort (GSE85217, 763 patients)^3^. We observed some differences in immune cell infiltration between the subgroups, with SHH-MB having an increased fraction of myeloid cells and a lower proportion of granulocytes compared to other subgroups, consistent with previous findings (**fig.1B, Table.S3**)25. The ratio of myeloid cells to T cells is also significantly higher in SHH-MB (**Fig. S1B)**. CD4^+^ T cells are the major lymphoid subsets in MB, followed by γδT cells. In contrast, CD8^+^T cells and NK cells, two cytotoxic subsets, are found in very low abundance in the tumor.

Interestingly, the frequency of γδT cells in MB differs between the subgroups, with the highest score in Grp4-MB (**fig.1C**), while αβT cells (CD8^+^ and CD4^+^) are more homogeneously distributed in the subgroups (**fig.1D**). In addition, the four MB subgroups can be further divided into 12 subtypes according to age, histology, methylation profile and driver mutations^3^, and the abundance score of γδT cells varies between subtypes in the same subgroup, except for WNT-MB (**fig.1E)**. We observed an increase of γδT cell score in SHHγ and Grp4γ and a slight decrease in Grp3γ subtypes. γδT cell infiltration in many tumor types have been shown to be associated with a favorable outcome^26,27^. We investigated whether the γδT cell scores correlate with prognosis in MB. Using the same cohort, patients were divided into γδT cell high vs γδT cell low abundance score (1^st^ vs 4^th^ quartile), but we found no significant correlation between γδT cell frequency and survival (p= 0.6570, log-rank test, **fig.1F)**. In conclusion, we show here that γδT cell infiltration is predicted for normal cerebellum and MB, with some differences between the different MB subgroups and subtypes, but γδT cell frequency did not correlate with prognosis, possibly due to its global low frequency.

### Tumor infiltrated γδT cells show a tissue resident phenotype

To further characterize the phenotype and activation state of γδT cells in MB, we extracted the BCR/TCR repertoire using the Trust4 algorithm from a single-cell RNA dataset of 28 MB patients (GSE155446, 1 WNT, 9 SHH, 7 Grp3 and 11 Grp4)^19^. Unfortunately, the number of B and T cells recovered by the algorithm was low compared to the clustering shown in the pediatric neuro-oncologic cell atlas. In total, 21 B cells, 56 αβT cells and 52 γδT cells were identified by the algorithm with a complete or partial BCR/TCR identification (**Table.S4**). Interestingly the γ- and the β-VJC segments were recovered better than their δ- and α-pairs. Therefore, we focused our analysis of γδT-infiltrated cells on the γ-chain. Analysis of the variable (V)γ-segment repertoire revealed a high diversity with a dominance for recombination of *TRGV2, TRGV9* and *TRGV10* genes in all MB subgroups (**fig.2A**). *TRGV10* is characterized as a pseudogene (Vγ type III) and result in a non-functional γδTCR^28^. In addition, *TRGV2, TRGV4* and *TRGV8*, already identified as brain-specific γδTCR signatures^29^, were present in MB. Interestingly, the public *TRGV9* clonotype (CALWEVQELGKKIKVF), which is present in the blood from fetal to adult life, was detected in one MB patient^30^. We next compared the diversity of the TRGV and TRGJ repertoires in the different MB subgroups (**fig.2B**). Grp3-infiltrated γδT cells showed a significant reduction in TRGV and TRGJ diversity compared to the SHH and Grp4 MB and had the highest proportion of non-functional TRGV10^+^ cells. However, the CDR3 length distribution of Grp3-infiltrated γδT cells is greater than in the other subgroups (Δ9 vs Δ6 or Δ7 amino acids for SHH- and Grp4-MB, respectively) (**fig.2C**). Overall, analysis of the γδTCR repertoire revealed no specific clonal expansion, suggesting a lack of tumor-specific reactivity.

**Figure 2:**
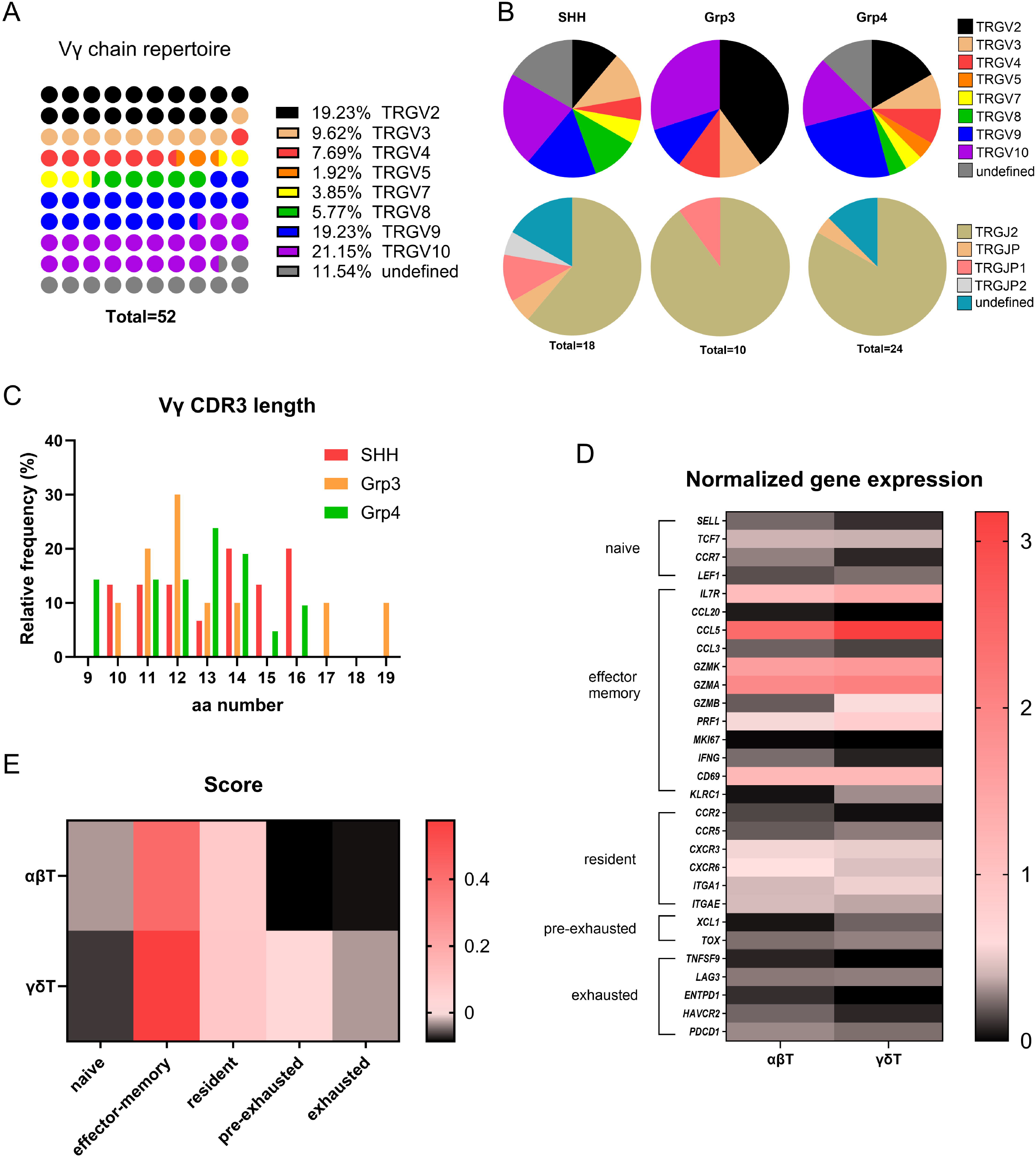
Infiltrated γδTCR repertoire and differentiation state. **(A)** TCR *TRGV* gene repertoire repartition (n=52 γδT cells, in 24 patients, GSE156053). (**B**) *TRGV* and *TRGJ* gene expression repartition across MB subgroups. (**C**) Relative frequency of the Vγ chain CDR3 across MB subgroups. Normalization of 30 gene expression of the identified αβT and γδT cells representative of five T cell differentiation state (n = n=56 αβT cells and n=52 γδT cells). (**E**) Signature score of each T cell differentiation state cluster.

To confirm this hypothesis, we examined the differentiation status of γδT and αβT cells identified by the Trust4 algorithm, independent of MB subgroup. We compared gene expression profile specific to naïve, effector memory, resident memory, pre-exhausted and exhausted T cells. There were few differences between infiltrated γδT and αβT cells, which preferentially expressed *ILR7, CCL5, GZMK, GZMA* and *CD69* (**fig.2D**). Evaluation of T cell differentiation state gene signatures of the T cell differentiation state revealed a phenotype enriched with resident memory cells that neither proliferate (*KI67*^low^), show proinflammatory activity (*IFNG*^low^), nor exhibit chronic activation-induced exhaustion (**fig.2E)**. In summary, γδ T-infiltrated cells in MB do not express markers that would suggest active involvement in anti-tumor immunity in MB.

### MB cells express γδT activating receptor ligands

Our analysis suggests that γδT cells infiltrate MB tumors but are insufficient in number and not functionally active. However, one of the main advantages of γδT cells is that they allow for allogeneic therapy due to their HLA-unrestricted activation. Therefore, we investigated which known ligands of human γδTCRs or co-receptors are expressed by MB cells to identify a γδT cell subset that could target MB cells. We examined a panel of seven MB cell lines covering three of the four MB subgroups (WNT excluded) with two paired primary (P) and recurrent (R) tumors for Grp3 and Grp4-MB, and that are considered relevant *in vitro* models^31^. We stained for γδTCR ligands: CD1d, CD1c, Ephrin Type-A receptor 2 (EphA2) and Annexin A2 (ANXA2); NKG2D ligands: UL16 binding protein (ULBP)2/5/6; MHC class I chain related protein (MIC)A/B; and DNAM-1 (CD226) ligands: CD112 and CD155 (**fig.3, fig.S2**). Flow cytometry results showed that all MB cell lines were negative for CD1c and that only DAOY cells presented weak expression of CD1d, as previously reported^32^. Overall, lipid presentation by CD1 molecules does not appear to be a promising target candidate for MB.

However, all MB cell lines express EphA2 and ANXA2, specific ligands for Vγ9Vδ1 and Vγ8Vδ3T cells, respectively^22,33^. CD112 and CD155, ligands of DNAM-1 an adhesion molecule, are also expressed by all MB cell lines tested. Binding of DNAM-1 promotes activation of T/NK cells and cytolytic degranulation^34^. However, CD112 and CD155 are also ligands for the checkpoint inhibitor TIGIT, which could promote immunosuppression. Interestingly, only the SHH-MB cell lines (DAOY, UW228-3 and ONS-76) were found expressing NKG2D ligands, suggesting that the immunogenicity of Grp3 and Grp4-MB for the activation of NK and γδT cells may be reduced. In summary, we identified a γδT cell ligand signature in MB cells that is subgroup-dependent, guiding to which γδT cell subset that can target MB cells.

**Figure 3:**
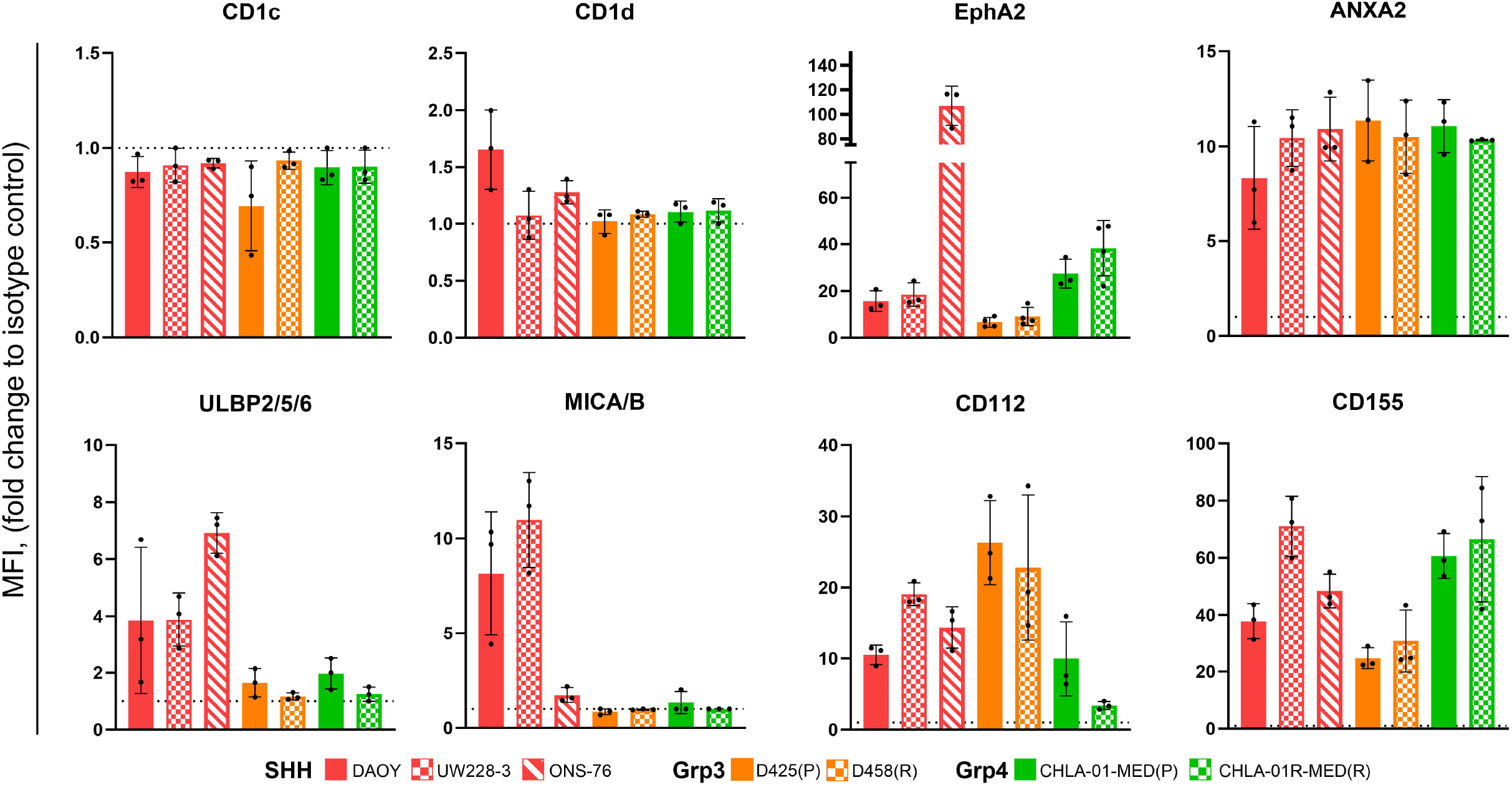
γδTCR, and γδT co-receptor ligand expression by MB cell lines. Fold change of MFI (Median of fluorescence) of expression of γδTCR, and γδT co-receptor ligands on seven MB cell lines compared to corresponding isotype controls, acquired by flow cytometry (dashed line =1, threshold of expression).

### EphA2-expressing MB cells trigger Vγ9Vδ1 T cell activation

EphA2 is a member of the Ephrin receptors, the largest receptor tyrosine kinase receptor family, and is involved in several biological processes such as neuronal development, cytoskeleton dynamic, migration, cell proliferation and angiogenesis. It is found to be overexpressed in tumors compared to normal tissues, making it a good target for cancer therapy^35^, which is supported by the development of CAR-T cells targeting EphA2 for Grp3-MB and ependymoma^36^. As proof-of-principal for EphA2 as a target ligand for Vγ9Vδ1T cells in MB, we used the Jurkat lymphoblastic cell line JRT3 (β-TCR chain defective), engineered to express the Vγ9Vδ1-MAU TCR that was previously shown to recognize EphA2^22^. The JRT3 model lacks cytotoxic capabilities but can be used to assess Vγ9Vδ1T cell activation. We co-cultured JRT3-MAU with MB cell lines or with healthy iPSC-derived neuroepithelial stem (NES) cells and measured the expression of the activation marker CD69 on JRT3-MAU by flow cytometry. Co-culture with MB cell lines induced expression of CD69, regardless of MB subgroup, resulting in strong activation of JRT3-MAU cells, except for the relapse cell line CHLA-01R-MED, which showed a lower response (**fig.4A**). However, co-culture with non-cancerous healthy NES cells resulted in significantly less activation of JRT3-MAU (**fig.4A**). Interestingly, the expression level of EphA2 does not seem to correlate with the activation potential. To confirm this observation, we measured EphA2 expression in NES cells, which was in the same range as in the D425 cell line but activated JRT3-MAU to a much lower extent (**fig.3, fig.4B**). These results suggest the presence of accessory signals on tumor cells that potentiate JRT3-MAU activation in addition to EphA2.

**Figure 4:**
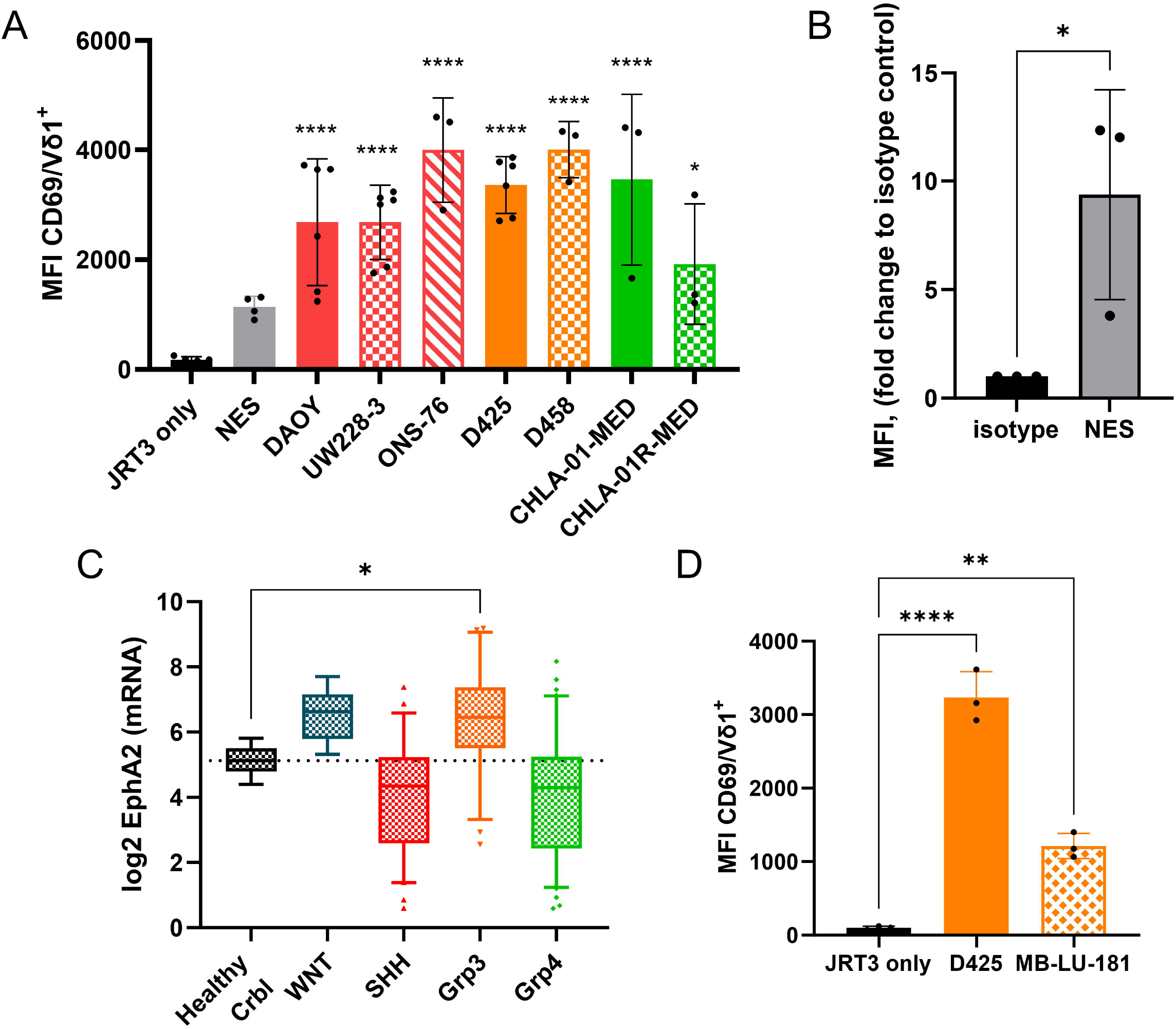
Vγ9Vδ1 TCR recognition of EphA2-positive MB cells. Analysis by flow cytometry of CD69 surface expression on Jurkat JRT3 MAU (Vγ9Vδ1 TCR positive) after a 4 h-co-culture (E/T ratio 1:1) with (**A**) Neuroepithelial Stem cells (NES) and MB cell lines, or (**D**) Grp3 PDX cells. n= 3 independent experiments. Statistical analysis was performed using one-way ANOVA followed by Dunnett tests to correct for multiplicity (\**P <* 0.05; \*\*\**P =* 0.0001; \*\*\*\**P <* 0.0001). (**B**) Fold change of MFI (Median of fluorescence) of EphA2 expression on NES. n= 3 independent experiments. Statistical analysis was performed using Unpaired t-test (\**P <* 0.05). (**C**) *EPHA2* mRNA expression in human MB subgroups versus normal human cerebellum (crbl). Statistical analysis was performed using one-way ANOVA followed by Dunnett tests to correct for multiplicity (\**P <* 0.05) – nonsignificant differences are not displayed in the figure.

Next, we examined EphA2 RNA expression in both healthy cerebellum and MB patients using a publicly available dataset^37^ and found that the *EPHA2* gene expression is upregulated in WNT- and Grp3-MB compared to normal cerebellum (**fig.4C**). To demonstrate the potential of Vγ9Vδ1T cells targeting Grp3-MB, we co-cultured JRT3-MAU with the Grp3-MB PDX MB-LU-181, using D425 cells as a reference. Results showed that JRT3-MAU upregulate CD69 expression in the presence of Grp3-MB PDX, yet to a lesser extent than with the D425 cell line (**fig.4D**). Overall, these results confirmed the potential of targeting EphA2-positive MB cells with Vγ9Vδ1T cells in a tumor-specific manner.

### Amino Bisphosphonates sensitize MB cells, but not healthy neuronal cells to Vγ9Vδ2T lysis

Cancer cells have a dysregulated metabolism with increased protein and lipid synthesis. Cholesterol, an essential component of the cell membrane, is produced via the mevalonate synthesis pathways. One of the intermediates of this pathway, isopentenyl pyrophosphate (IPP), can bind to the intracellular domain of a member of the butyrophilin family, BTN3A1, and trigger the activation of Vγ9Vδ2T cells^38,39^. Vγ9Vδ2T cells are a peripheral population and are thought to naturally recognize and lyse leukemia cells^40^. However, solid tumors often remain undetectable to Vγ9Vδ2T cell targeting^41^. Fortunately, amino-bisphosphonate (NBP) molecules such as Zoledronate (zol) can increase intracellular IPP levels by inhibiting the farnesyl diphosphate synthase in target cells, which then enhance Vγ9Vδ2T cell recognition and activation (**fig.5A**).

**Figure 5:**
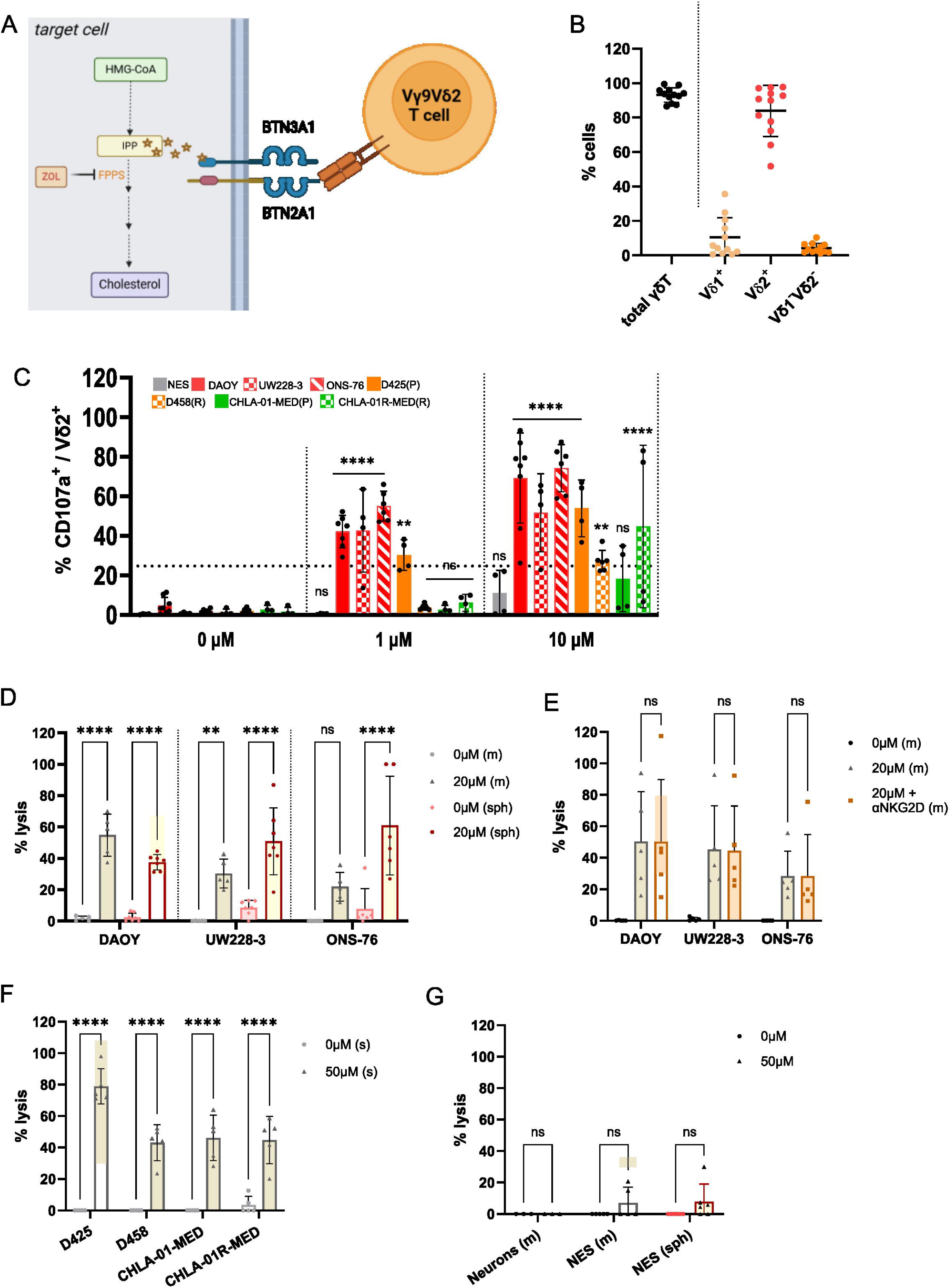
Vγ9Vδ2 TCR recognition of zoledronate-treated MB cells. **(A)** Schematic of the action of zoledronate in the cholesterol synthesis pathway leading to the activation of the Vγ9Vδ2 TCR via the BTN3A1/2A1 complex. (B) Percentage of total γδT cells in the ex vivo expanded population and percentage of the different γδT cell subsets assessed by flow cytometry. n=12 donors (C) Analysis by flow cytometry of CD107a expression on ex vivo-expanded human Vδ2T cells after 4 h of co-culture Neuroepithelial Stem cells (NES) and MB cell lines treated with 0 or 10 µM of zol (E/T ratio 1:1). n = 4-6 donors. (**D**)-(**G**) Analysis of cytotoxicity by LDH release in the supernatant, after 8 h-co-culture with ex vivo expanded γδT cells, by (**D**) SHH-MB cell lines grown in monolayer (m) or spheroid (sph), treated with 0 or 20 µM of zol; (**E**) by SHH-MB cell lines grown in monolayer (m), treated with 0 or 20 µM of zol with or without pre-conditioning with NKG2D blocking antibody (10µg/ml); (**F**) by Grp3- and Grp4-MB grown in suspension (s), treated with 0 or 50 µM of zol; (**G**) by differentiated neurons and NES grown monolayer (m) or spheroid (sph), treated with 0 or 50 µM of zol. n = 3-6 donors; statistical analysis was performed using two-way ANOVA followed by (C) Dunnett, (D) Tukey (E)-(G) Sidak tests to correct for multiplicity (\**P <* 0.05; \*\**P <* 0.005; \*\*\**P =* 0.0001; \*\*\*\**P <* 0.0001) – nonsignificant differences are not displayed in the figure.

BTN3A1 is ubiquitously expressed in all human cells, including healthy brain and MB cells. We tested whether Vγ9Vδ2T cells can naturally recognize MB cells or be sensitized by zol treatment. NES and MB cells were treated overnight with 0, 1 or 10µM of zol followed by co-culture with ex-vivo expanded peripheral γδT cells isolated from healthy donors, the majority of which consisted of Vδ2^+^ γδT cells (**fig.5B**). The expression of the degranulation marker CD107a was analyzed on Vδ2^+^ γδT cells by flow cytometry. As expected, untreated NES and MB cells did not trigger Vγ9Vδ2T cell degranulation (**fig.5C**). However, MB cells treated with 10µM zol, but not NES cells, were able to significantly activate Vγ9Vδ2T cells. The suboptimal dose of 1µM showed a difference in the sensitivity of each cell line to zol-treatment, with SHH cell lines and D425 cells inducing CD107a expression on Vγ9Vδ2T cells, but not D458 and Grp4-MB cells. Furthermore, we detected differences between donors in terms of zol-mediated activation against MB.

Next, we investigated whether zol treatment was sufficient to induce killing of MB cells by Vγ9Vδ2T cells. We cultured MB cells, either as adherent monolayer (m) or as spheroids (sph), together with ex vivo expanded γδT cells and quantified LDH release as a measure of cell death. LDH release results showed that SHH-MB treated with zol induced killing by γδT cells in both monolayer and spheroid conditions (**fig.5D)**. The cytotoxic potential of γδT cells against zol-treated SHH-MB cells is donor- and target-dependent. Interestingly, the UW228-3 and ONS-76 cell lines are eliminated more efficiently in spheroid form. We hypothesized that the percentage of killing in monolayer could be underestimated by the readout method, as brightfield images taken after co-culture showed no remaining adherent MB cells (**fig.S3**). Considering that we found that SHH-MB cells express NKG2D ligands, we further hypothesized that NKG2D might act in synergy with TCR activation against SHH-MB. However, NKG2D blocking on γδT cell did not affect the killing potential of zol-treated SHH-MB (**fig.5E, fig.S3**). The CD107a results indicated that Grp3- and Grp4-MB were less sensitive to zol treatment, so the zol concentration was increased to 50µM. Since Grp3- and Grp4-MB are naturally suspension cell types, the treatment and co-culture were performed accordingly. Zol-treated D425 reached an average of 80% of lysis, while the other three cell lines are closer to 45% (**fig.5F**). The Grp4-MB cell lines CHLA-01 and CHLA-01R grow as large neurospheres and have a lower proliferation rate than the Grp3 cells, which could explain the difference. However, the difference between the D425 and D458 cell lines could be the result of resistance acquired at relapse.

Finally, we examined the off-target effect of zol treatment on both proliferating NES cells and post-mitotic neurons. We treated the neuronal cells both as monolayer and as spheroids with high doses of zol (50 µM). The co-culture of zol-treated NES cells and neurons did not induce significant cytotoxicity-mediated by Vγ9Vδ2T cells, demonstrating that zol treatment trigger specific targeting and killing of MB cells by Vγ9Vδ2T cells while sparing healthy stem and differentiated neuronal cells (**fig.5G)**.

## Discussion

CNS tumors remain among the most challenging to cure due to the complexity of the surrounding environment. Brain cells are very sensitive to conventional cancer therapy such as chemotherapies and radiotherapies^42^, which has a significant impact on treatment outcomes and the long-term quality of life of survivors. Given that MB develops in the cerebellum, which controls physical movements, balance, and coordination^43^, it is crucial to develop less toxic therapies to preserve these important bodily functions.

A healthy immune system can distinguish between normal cells and infected or transformed cells. In cancer patients, however, the immune system is often impaired. Therefore, allogeneic transfer of immune cells from healthy individuals is a promising therapeutic strategy for cancer patients^44,45^. Non-alloreactive and highly cytotoxic, γδT cells are a promising approach for allogeneic cell therapy in cancer patients^46^. Here, we demonstrate that MB patients may benefit from allogeneic γδT cell transfer. We confirm previous studies that the MB microenvironment lacks sufficient functional cytotoxic T cells, including both αβT and γδT cells, to elicit a sufficient anti-tumor response from autologous T cells. The small numbers of αβT and γδT cells identified by the TCR repertoire algorithm Trust4 showed no significant clonal expansion or expression of activation/exhaustion markers, suggesting that conventional ICB therapy may show limited efficacy in MB patients^47^. However, the low TCR recovery could be due to insufficient sequencing quality or depth. Future sequencing experiments with fresh patient samples and protocols optimized for TCR chain repertoire analysis could provide better insights. B7-H3 is the only immune checkpoint protein reported to be significantly expressed in MB and other brain tumors^48^. Strategies to use CAR-T cells targeting B7-H3 or antibody-drug conjugate are currently being investigated in various brain tumors and show some therapeutic efficacy and promising safety profile^49,50^. B7-H3 has been reported to confer resistance to killing by Vγ9Vδ2 T cells in a colon cancer model in a STAT3/ULBP2 axis-dependent manner^51^. However, the direct or indirect effects of B7-H3 expressed by MB on the cytotoxicity of γδT cells remains to be investigated.

We showed that MB cells express several ligands recognized by different γδTCR receptors and co-receptors (NKG2D and DNAM1), indicating the potential to target them with the corresponding γδT cell subset. We tested the activation potential of different γδT cell subsets against MB cell lines based on the TCR ligands they expressed. First, we demonstrated that seven MB cell lines and one PDX expressed EphA2 and activated Vγ9Vδ1-MAU TCR Jurkat cells. Similarly, CAR-T cells targeting EphA2 have shown strong anti-tumor potential in an *in vivo* Grp3-PDX model^36^. However, these CAR-T cells lack the multipotency of multiple activating receptors expressed by γδT cells, which is an advantage in heterogeneous solid tumors. On the other hand, maintaining and expanding T cell clones in culture is challenging, so using the natural MAU clone may not be a viable therapeutic option. An alternative is TCR-engineered T cell therapy, where the Vγ9Vδ1 TCR may be integrated into adoptive cell therapy (ACT) candidates, such as γδT or NKT cells. The latter are particularly interesting as they retain their endogenous TCR and can present dual-TCR recognition^52^. Additionally, zol treatment can sensitize MB cells to killing by *ex vivo* expanded Vγ9Vδ2T cells. Zol, commercially available as Zometa®, is prescribed for bone diseases, including bone malignancies and osteoporosis. We demonstrated the direct anti-tumor potential of zol against MB cells, however, it has also been reported that zol can inhibit pro-tumoral macrophages and microglia in a breast cancer brain metastasis model^53^. Importantly, both Vγ9Vδ1 and Vγ9Vδ2 T cell activation and killing were restricted to tumor cells, as normal NES and differentiated neurons did not triggering γδT cell activation, demonstrating the specificity of tumor cell targeting.

T cell-based therapies, including CAR-T cells and T cell engagers (TCE), face safety concerns, particularly regarding immune effector cell-associated neurotoxicity syndrome (ICANS) and cytokine storm syndrome (CSS)^54^. However, the first clinical trial of allogeneic intravenous Vγ9Vδ2 T cell transfer for solid tumors did not report severe side effects associated with the ACT^14^. Nevertheless, there is currently no clinical data on the safety of intraventricular infusion of allogeneic γδT cells in patients, making it essential to identify potential side effects associated with this therapeutic strategy. A phase II clinical study using autologous Vγ9Vδ2 T cells genetically modified to resist Temozolomide, a conventional chemotherapy used for glioblastoma, is ongoing and could provide future indication regarding the safety of intracranial injection of γδT cells (NCT05664243).

In conclusion, our study is the first to support the therapeutic potential of allogeneic γδT cells for MB patients and their potential as a safer alternative to conventional treatment.

## Author contributions

Conception and design of the study: L.B and M.W.; Analysis and interpretation of the data: L.B. Generation and acquisition of data: L.B, M.L, O.B-R. Contribution of new reagents: J.D.M. Statistical analysis: L.B and O.B-R. Writing original draft of manuscript: L.B, O.B-R. J.D.M. and M.W, All authors have reviewed, read, and approved the submitted version of the manuscript.

## Data Availability Statement

All data relevant to the study are included in the article or uploaded as supplementary information

## Acknowledgements

The authors thank the Biomedicum Flow Cytometry facility (BFC) for technical assistance. Part of the computations and data handling were enabled by resources provided by the National Academic Infrastructure for Supercomputing in Sweden (NAISS), partially funded by the Swedish Research Council through grant agreement no. 2022-06725. We thank everyone in the Wilhelm lab for technical assistance and helpful discussions.

