## Supplemental Figures for "EphA2 and Phosphoantigen-Mediated Selective Killing of Medulloblastoma by γδT Cells Preserves Neuronal and Stem Cell Integrity"

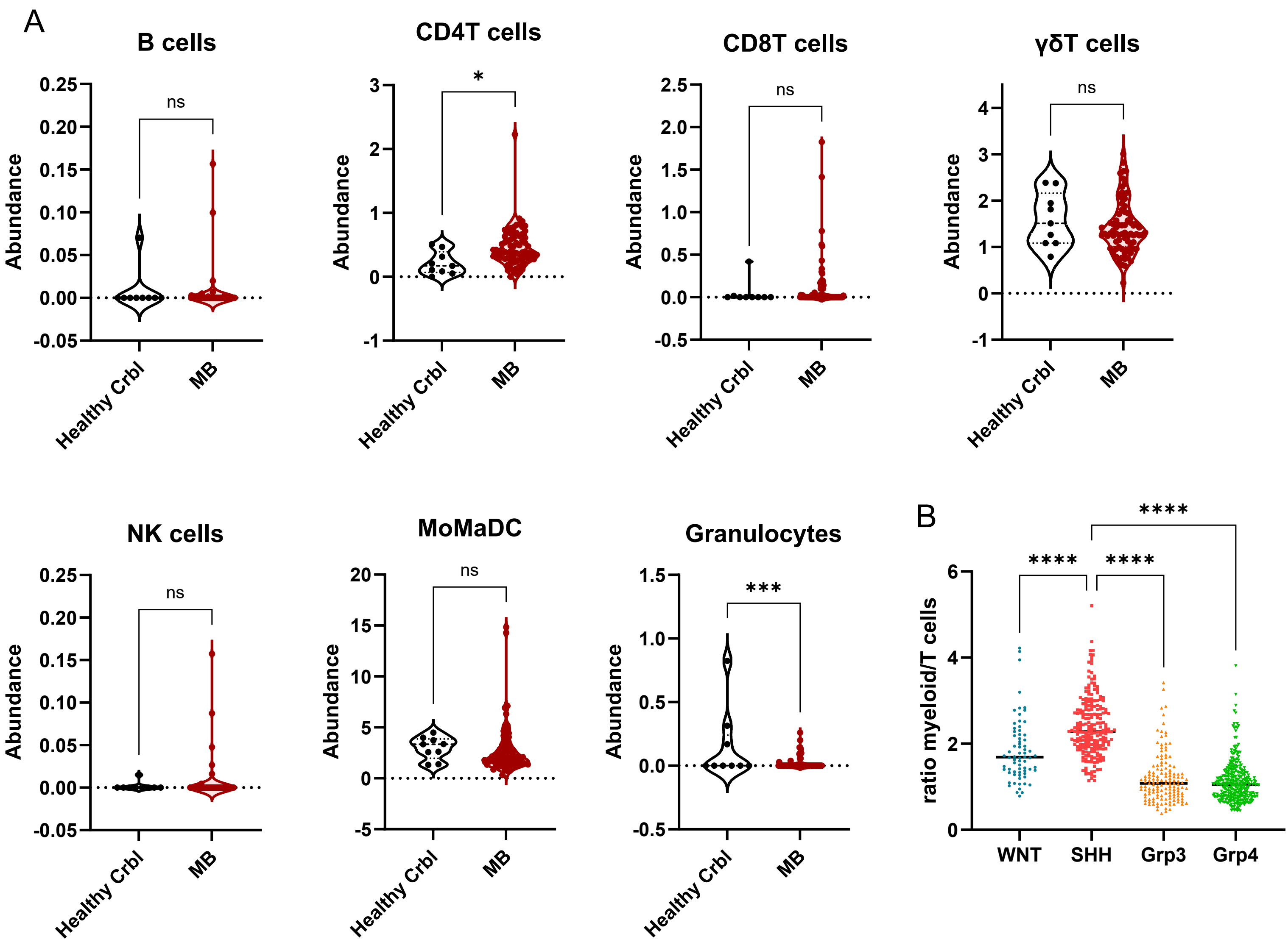


**Figure S1: Immune cell abundance in healthy cerebellum and Medulloblastoma** . (**A**) Abundance calculated after LM -7 CIBERSORT deconvolution of normal cerebellum (n=9) versus MB patients (n=76). Statistical analysis was performed using Unpaired t-test (*$P$ <0.05) – nonsignificant differences are not displayed in the figure. ( **B**) ratio of myeloid cells/ T cells (CD4+CD8+ γδT cells) in SHH-MB versus other subgroups. Statistical analysis was performed using two -way ANOVA followed by Tukey test to correct for multiplicity (****$P$ < 0.0001).

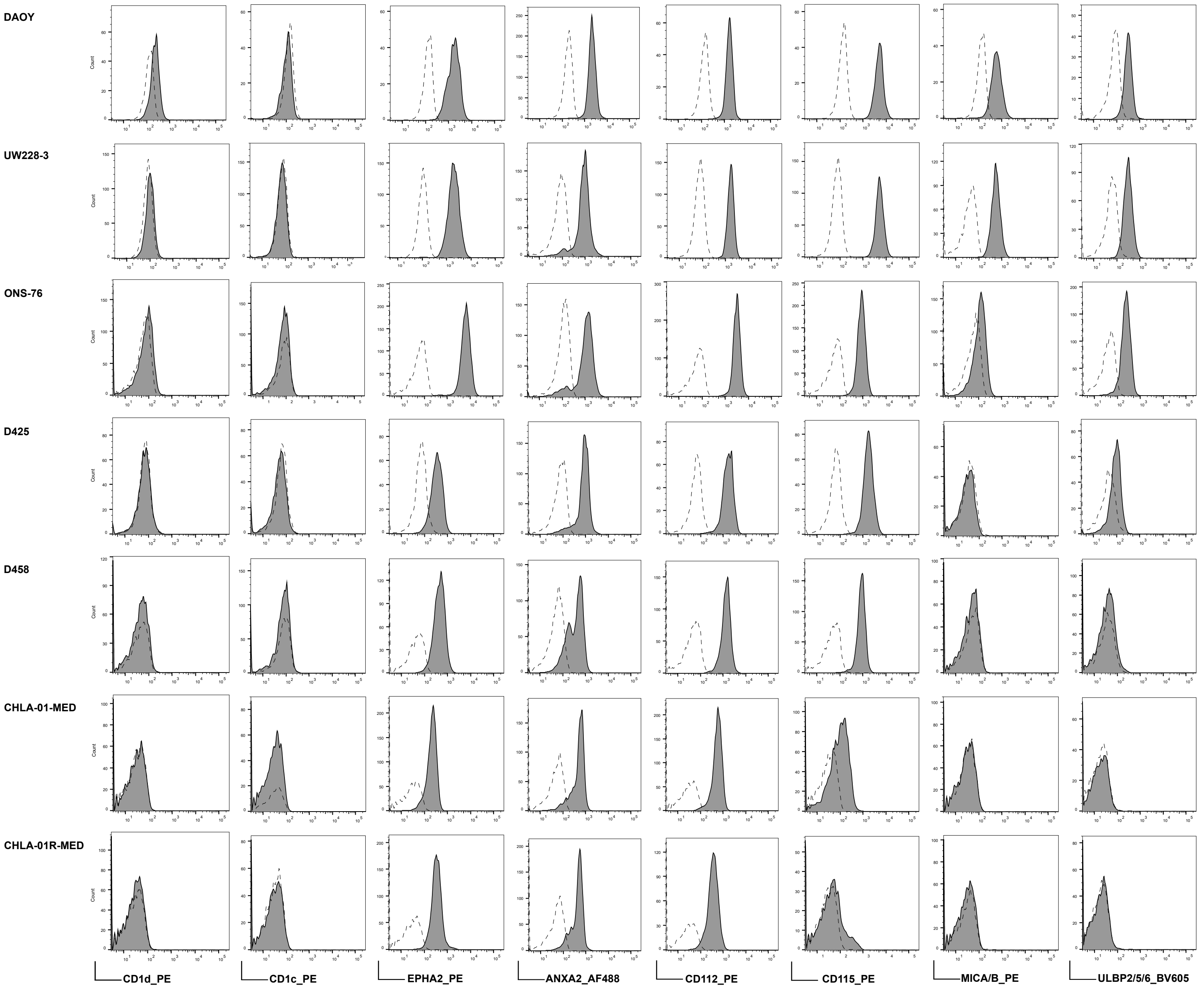


**Figure S2: Representative histograms of the expression of CD1d, CD1c, EphrinA2 (EphA2), annexin A2, CD112, CD155, MICA/B and ULBP2,5,6 on 7 MB cells lines.** Isotype control (dashed line), antibody (full line).

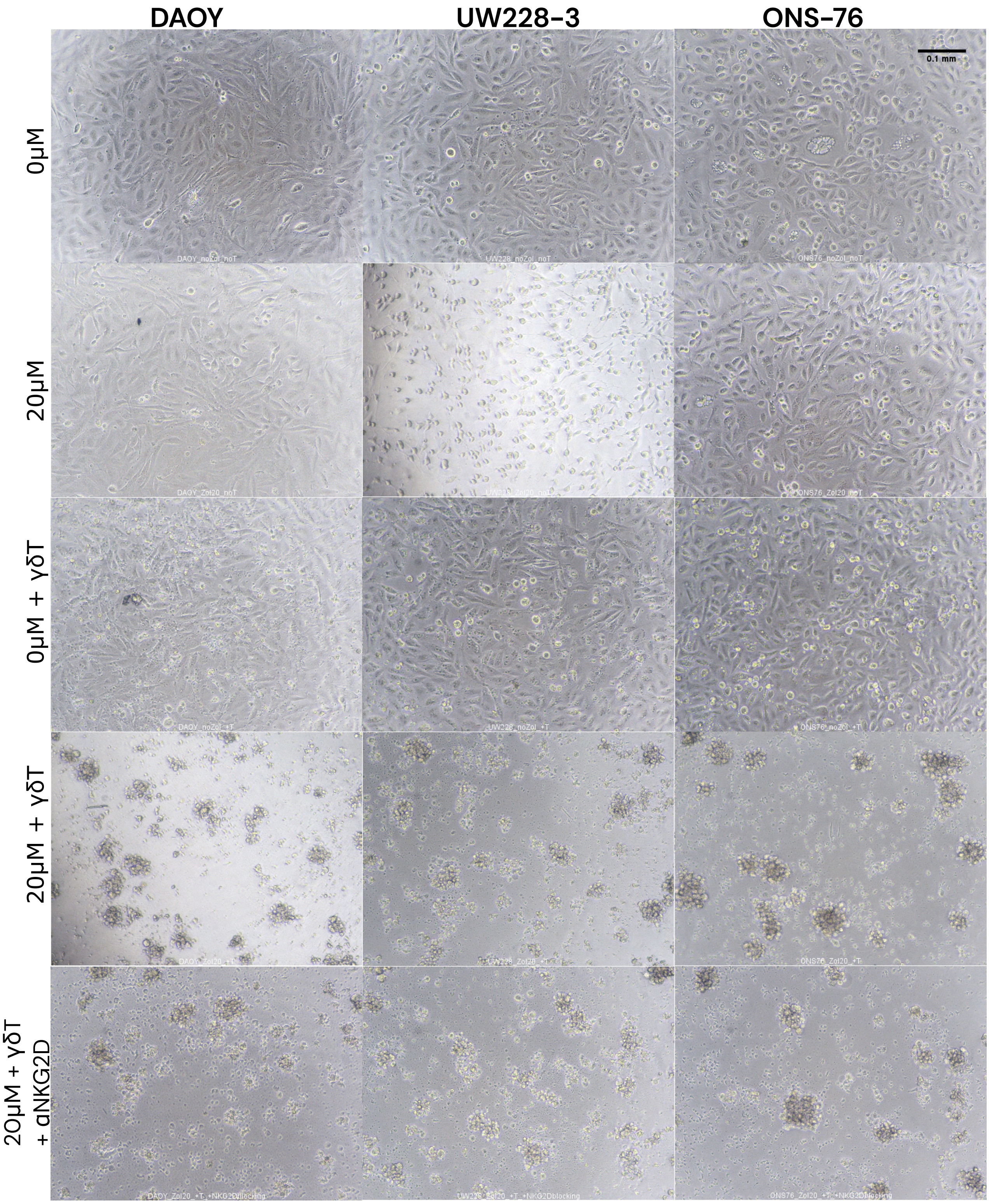


**Figure S3: Brightfield images of DAOY, UW228 -3 and ONS -76 after 8 hours co-culture.** MB treated with or without Zol treatment, anti-NKG2D antibody, and/or ex vivo expanded γδT cells. Scale bar = 0,1 mm [Nikon Eclipse Ts2, objective 10x, N.A 0.25].
