## Supplemental Table 1 for "EphA2 and Phosphoantigen-Mediated Selective Killing of Medulloblastoma by γδT Cells Preserves Neuronal and Stem Cell Integrity"

Table S1

Boutin et al.

**Ephrin-A2 and Phosphoantigen-Mediated Selective Killing of Medulloblastoma by γδT Cells Preserves Neuronal and Stem Cell Integrity**

Gene Signatures

| Naive | Effector Memory | Resident | Pre-exhausted | Exhausted |
|---|---|---|---|---|
| sell | ccl20 | itga1 | xcl1 | pdcd1 |
| tcf7 | il7r | itgae | tox | havcr2 |
| ccr7 | cd69 | cxcr6 | | prf1 |
| lef1 | gzma | ccr2 | | tigit |
| | gzmk | cxcr3 | | lag3 |
| | ccl5 | ccr5 | | gzmb |
| | klrc1 | lfa1 | | entpd1 |
| | ifng | | | ifng |
| | mki67 | | | |
| | gzmb | | | |
| | tnfsf9 | | | |
| | ccl3 | | | |
