## Supplemental Table 2 for "EphA2 and Phosphoantigen-Mediated Selective Killing of Medulloblastoma by γδT Cells Preserves Neuronal and Stem Cell Integrity"

**Table S2**

Boutin et al.

**Ephrin-A2 and Phosphoantigen-Mediated Selective Killing of Medulloblastoma by γδT Cells Preserves Neuronal and Stem Cell Integrity**

**CIBERSORTx_GSE44971**

| Sample | Bcells | TCD4 | TCD8 | Tgd | NK | MoMaDC | granulocytes | P-value | Correlation | RMSE | HG-U133_Plus_2 | age group |
|---|---|---|---|---|---|---|---|---|---|---|---|---|
| GSM1094863 | 0 | 0 | 0 | 0,296381 | 0 | 0,652177 | 0,05144213 | 0 | 0,709788221 | 0,703862 | GSE44971-CBM-07 | fetal |
| GSM1094864 | 0 | 0,042458 | 0 | 0,311428 | 0 | 0,646114 | 0 | 0 | 0,552661778 | 0,843202 | GSE44971-CBM-08 | fetal |
| GSM1094865 | 0 | 0,02244 | 0,007153 | 0,353972 | 0 | 0,616435 | 0 | 0 | 0,50876315 | 0,876936 | GSE44971-CBM-09 | fetal |
| GSM1094866 | 0 | 0,079161 | 0 | 0,273212 | 0 | 0,647627 | 0 | 0 | 0,53941579 | 0,851896 | GSE44971-CBM-10 | fetal |
| GSM1094867 | 0 | 0,043656 | 0 | 0,432214 | 0 | 0,524131 | 0 | 0 | 0,413776482 | 0,940446 | GSE44971-CBM-11 | fetal |
| GSM1094868 | 0 | 0,036438 | 0 | 0,341741 | 0 | 0,592212 | 0,02960967 | 0 | 0,440667092 | 0,924395 | GSE44971-CBM-12 | adult |
| GSM1094869 | 0 | 0,08187 | 0 | 0,381771 | 0 | 0,536359 | 0 | 0 | 0,461304163 | 0,9031 | GSE44971-CBM-13 | adult |
| GSM1094870 | 0 | 0,010311 | 0,051036 | 0,289743 | 0 | 0,548367 | 0,10054307 | 0 | 0,50784149 | 0,871125 | GSE44971-CBM-14 | adult |
| GSM1094871 | 0,012108 | 0,080714 | 0 | 0,259827 | 0,002582 | 0,644769 | 0 | 0 | 0,520897658 | 0,865392 | GSE44971-CBM-15 | adult |
| **average** | **0,001345** | **0,044116** | **0,006465** | **0,326699** | **0,000287** | **0,60091** | **0,02017721** | | | | | |

**SES_CIBERSORTx_GSE44971**

| Sample | Bcells | TCD4 | TCD8 | Tgd | NK | MoMaDC | granulocytes | P-value | Correlation | RMSE |
|---|---|---|---|---|---|---|---|---|---|---|
| GSM1094863 | 0 | 0 | 0 | 1,812349 | 0 | 3,988011 | 0,31456475 | 0 | 0,709788221 | 0,703862 |
| GSM1094864 | 0 | 0,1723 | 0 | 1,263826 | 0 | 2,622038 | 0 | 0 | 0,552661778 | 0,843202 |
| GSM1094865 | 0 | 0,050178 | 0,015994 | 0,791509 | 0 | 1,378398 | 0 | 0 | 0,50876315 | 0,876936 |
| GSM1094866 | 0 | 0,314148 | 0 | 1,084229 | 0 | 2,570074 | 0 | 0 | 0,53941579 | 0,851896 |
| GSM1094867 | 0 | 0,109619 | 0 | 1,085285 | 0 | 1,316087 | 0 | 0 | 0,413776482 | 0,940446 |
| **fetal** | **0** | **0,129249** | **0,003199** | **1,20744** | **0** | **2,374922** | **0,06291295** | | | |
| GSM1094868 | 0 | 0,207553 | 0 | 1,94659 | 0 | 3,373301 | 0,16865976 | 0 | 0,440667092 | 0,924395 |
| GSM1094869 | 0 | 0,511568 | 0 | 2,385505 | 0 | 3,351458 | 0 | 0 | 0,461304163 | 0,9031 |
| GSM1094870 | 0 | 0,084573 | 0,418622 | 2,376607 | 0 | 4,497952 | 0,82469985 | 0 | 0,50784149 | 0,871125 |
| GSM1094871 | 0,070364 | 0,469053 | 0 | 1,509937 | 0,015002 | 3,746958 | 0 | 0 | 0,520897658 | 0,865392 |
| **adult** | **0,017591** | **0,318187** | **0,104655** | **2,05466** | **0,00375** | **3,742417** | **0,2483399** | | | |

**CIBERSORTx_GSE37418**

| Sample | Bcells | TCD4 | TCD8 | Tgd | NK | MoMaDC | granulocytes | P-value | Correlation | RMSE | Group |
|---|---|---|---|---|---|---|---|---|---|---|---|
| GSM918586 | 0,017527 | 0,101786 | 0 | 0,276659 | 0,009757 | 0,594271 | 0 | 0 | 0,745226239 | 0,674223 | G3 |
| GSM918588 | 0 | 0,1209 | 0 | 0,446285 | 0 | 0,377789 | 0,05502652 | 0 | 0,644504791 | 0,764481 | G3 |
| GSM918589 | 0 | 0,154907 | 0 | 0,370216 | 0 | 0,474877 | 0 | 0 | 0,543345164 | 0,839754 | G3 |
| GSM918594 | 0 | 0,152356 | 0 | 0,262159 | 0 | 0,585485 | 0 | 0 | 0,738920529 | 0,681943 | G3 |
| GSM918600 | 0,003214 | 0,09692 | 0 | 0,362661 | 0,012582 | 0,524623 | 0 | 0 | 0,726498127 | 0,693487 | G3 |
| GSM918601 | 0 | 0,063409 | 0,050397 | 0,368759 | 0 | 0,517435 | 0 | 0 | 0,534048051 | 0,84921 | G3 |
| GSM918611 | 0 | 0,191128 | 0 | 0,411861 | 0 | 0,39701 | 0 | 0 | 0,697986611 | 0,723199 | G3 |
| GSM918612 | 0 | 0,052031 | 0,020939 | 0,373635 | 0 | 0,553395 | 0 | 0 | 0,749222808 | 0,669547 | G3 |
| GSM918614 | 0 | 0,079907 | 0 | 0,381801 | 0 | 0,538292 | 0 | 0 | 0,592644438 | 0,805591 | G3 |
| GSM918624 | 0 | 0,038389 | 0,004163 | 0,31672 | 0 | 0,640728 | 0 | 0 | 0,717660113 | 0,696393 | G3 |
| GSM918629 | 0 | 0,12376 | 0 | 0,386114 | 0,001182 | 0,488943 | 0 | 0 | 0,625663426 | 0,778895 | G3 |
| GSM918636 | 0,000899 | 0,140854 | 0 | 0,376218 | 0 | 0,482029 | 0 | 0 | 0,582007313 | 0,812044 | G3 |
| GSM918637 | 0 | 0,13248 | 0,061592 | 0,318227 | 0 | 0,400966 | 0,08673552 | 0 | 0,653961179 | 0,760945 | G3 |
| GSM918639 | 0 | 0,126074 | 0 | 0,33397 | 0 | 0,510225 | 0,02973084 | 0 | 0,593287676 | 0,803682 | G3 |
| GSM918640 | 0 | 0,101895 | 0,102006 | 0,290704 | 0 | 0,505395 | 0 | 0 | 0,527351528 | 0,850685 | G3 |
| GSM918578 | 0 | 0,049312 | 0 | 0,421318 | 0 | 0,529371 | 0 | 0 | 0,6527244 | 0,756816 | G4 |
| GSM918579 | 0 | 0,095753 | 0 | 0,428402 | 0 | 0,465948 | 0,00989776 | 0 | 0,521313151 | 0,858146 | G4 |
| GSM918581 | 0 | 0,129461 | 0 | 0,339247 | 0 | 0,531292 | 0 | 0 | 0,53895604 | 0,844336 | G4 |
| GSM918583 | 0 | 0,18862 | 0 | 0,378516 | 0 | 0,432863 | 0 | 0 | 0,499878255 | 0,868487 | G4 |
| GSM918584 | 0 | 0,15149 | 0 | 0,37631 | 0 | 0,4722 | 0 | 0 | 0,530507863 | 0,848942 | G4 |
| GSM918585 | 0 | 0,134164 | 0 | 0,397818 | 0 | 0,468018 | 0 | 0 | 0,615534399 | 0,786797 | G4 |
| GSM918590 | 0 | 0,017283 | 0,1045 | 0,282472 | 0 | 0,595745 | 0 | 0 | 0,712224729 | 0,703973 | G4 |
| GSM918591 | 0 | 0,093873 | 0 | 0,452079 | 0 | 0,454048 | 0 | 0 | 0,471107816 | 0,89494 | G4 |
| GSM918592 | 0 | 0,028188 | 0 | 0,380902 | 0 | 0,555987 | 0,03492239 | 0 | 0,636919967 | 0,770444 | G4 |
| GSM918595 | 0 | 0,082028 | 0 | 0,244743 | 0 | 0,673229 | 0 | 0 | 0,775950041 | 0,635449 | G4 |
| GSM918598 | 0 | 0,138214 | 0 | 0,403366 | 0 | 0,45842 | 0 | 0 | 0,558704282 | 0,829278 | G4 |
| GSM918599 | 0 | 0,139318 | 0 | 0,379309 | 0 | 0,481373 | 0 | 0 | 0,618088138 | 0,784836 | G4 |
| GSM918602 | 0 | 0,146907 | 0 | 0,271704 | 0 | 0,581389 | 0 | 0 | 0,753500263 | 0,668529 | G4 |
| GSM918605 | 0 | 0,114419 | 0,003569 | 0,352445 | 0 | 0,529568 | 0 | 0 | 0,66108825 | 0,750159 | G4 |
| GSM918610 | 0 | 0,116674 | 0 | 0,381158 | 0 | 0,502168 | 0 | 0 | 0,4638473 | 0,897534 | G4 |
| GSM918613 | 0 | 0,056877 | 0 | 0,392485 | 0 | 0,533965 | 0,01667371 | 0 | 0,612409808 | 0,790184 | G4 |
| GSM918615 | 0,000167 | 0,088609 | 0 | 0,312618 | 0 | 0,57636 | 0,02224662 | 0 | 0,734676686 | 0,683719 | G4 |
| GSM918616 | 0 | 0,165408 | 0 | 0,369675 | 0 | 0,453177 | 0,01173978 | 0 | 0,548080267 | 0,835866 | G4 |
| GSM918622 | 0,0067 | 0,09779 | 0 | 0,423474 | 0 | 0,442807 | 0,02922917 | 0 | 0,566625916 | 0,823888 | G4 |
| GSM918623 | 0 | 0,147114 | 0 | 0,341039 | 0 | 0,511847 | 0 | 0 | 0,498484887 | 0,871822 | G4 |
| GSM918626 | 0 | 0,102766 | 0,08425 | 0,121454 | 0,007254 | 0,684276 | 0 | 0 | 0,825977119 | 0,580267 | G4 |
| GSM918627 | 0 | 0,161139 | 0 | 0,409963 | 0 | 0,398386 | 0,03051227 | 0 | 0,578375764 | 0,814343 | G4 |
| GSM918630 | 0 | 0,167374 | 0 | 0,409457 | 0 | 0,423169 | 0 | 0 | 0,471492241 | 0,888864 | G4 |
| GSM918631 | 0,000629 | 0,116191 | 0,043035 | 0,280929 | 0,003502 | 0,543386 | 0,01232915 | 0 | 0,540565236 | 0,842619 | G4 |
| GSM918632 | 0 | 0,029831 | 0,061692 | 0,413638 | 0 | 0,49484 | 0 | 0 | 0,555617118 | 0,834545 | G4 |
| GSM918633 | 0 | 0,076168 | 0 | 0,36045 | 0 | 0,563382 | 0 | 0 | 0,671703384 | 0,740275 | G4 |
| GSM918634 | 0 | 0,056741 | 0 | 0,377199 | 0 | 0,56606 | 0 | 0 | 0,589257565 | 0,8095 | G4 |
| GSM918635 | 0 | 0,094336 | 0,049433 | 0,28987 | 0 | 0,566362 | 0 | 0 | 0,441561448 | 0,914181 | G4 |
| GSM918645 | 0 | 0,194384 | 0 | 0,376588 | 0 | 0,414603 | 0,01442548 | 0 | 0,509443562 | 0,86147 | G4 |
| GSM918646 | 0 | 0,017692 | 0,070861 | 0,32024 | 0 | 0,591207 | 0 | 0 | 0,633229101 | 0,773426 | G4 |
| GSM918647 | 0 | 0,204733 | 0,003419 | 0,304103 | 0 | 0,487744 | 0 | 0 | 0,534670496 | 0,844582 | G4 |
| GSM918648 | 0 | 0,0979 | 0 | 0,377345 | 0 | 0,524755 | 0 | 0 | 0,569415941 | 0,822842 | G4 |
| GSM918650 | 0 | 0,165699 | 0 | 0,431637 | 0 | 0,345813 | 0,05685123 | 0 | 0,425347739 | 0,917313 | G4 |
| GSM918652 | 0 | 0,165665 | 0 | 0,339137 | 0 | 0,495198 | 0 | 0 | 0,542519369 | 0,840117 | G4 |
| GSM918653 | 0 | 0,070158 | 0 | 0,459559 | 0 | 0,470283 | 0 | 0 | 0,440506923 | 0,918691 | G4 |
| GSM918582 | 0 | 0,130967 | 0 | 0,344869 | 0 | 0,524164 | 0 | 0 | 0,727414629 | 0,694358 | SHH |
| GSM918606 | 0 | 0,129215 | 0 | 0,269355 | 0 | 0,60143 | 0 | 0 | 0,658581927 | 0,751392 | SHH |
| GSM918607 | 0 | 0,068069 | 0,045646 | 0,216363 | 0 | 0,669922 | 0 | 0 | 0,858599791 | 0,540692 | SHH |
| GSM918608 | 0 | 0,05357 | 0 | 0,278668 | 0 | 0,667762 | 0 | 0 | 0,749673853 | 0,66318 | SHH |
| GSM918609 | 0 | 0,095336 | 0 | 0,287636 | 0 | 0,617029 | 0 | 0 | 0,722486335 | 0,693345 | SHH |
| GSM918619 | 0 | 0,088588 | 0 | 0,308486 | 0 | 0,602926 | 0 | 0 | 0,778059103 | 0,63952 | SHH |
| GSM918620 | 0 | 0,049913 | 0,022288 | 0,313055 | 0 | 0,614745 | 0 | 0 | 0,671419956 | 0,740145 | SHH |
| GSM918621 | 0 | 0,065519 | 0 | 0,27318 | 0 | 0,661301 | 0 | 0 | 0,851038968 | 0,548968 | SHH |
| GSM918649 | 0 | 0 | 0,088517 | 0,316578 | 0 | 0,594906 | 0 | 0 | 0,816036975 | 0,598646 | SHH |
| GSM918651 | 0 | 0,167787 | 0,024434 | 0,293743 | 0 | 0,514037 | 0 | 0 | 0,677514809 | 0,739274 | SHH |
| GSM918587 | 0,000384 | 0,094471 | 0,046486 | 0,200965 | 0,007429 | 0,650266 | 0 | 0 | 0,861080381 | 0,54373 | Unknown |
| GSM918596 | 0 | 0,069876 | 0 | 0,32802 | 0 | 0,602105 | 0 | 0 | 0,684991501 | 0,72799 | Unknown |
| GSM918597 | 0 | 0,104133 | 0 | 0,354963 | 0 | 0,540904 | 0 | 0 | 0,550154392 | 0,837322 | Unknown |
| GSM918604 | 0 | 0,065128 | 0 | 0,335515 | 0 | 0,599357 | 0 | 0 | 0,809303792 | 0,605312 | Unknown |
| GSM918617 | 0 | 0,119203 | 0,008339 | 0,347081 | 0 | 0,525376 | 0 | 0 | 0,585912495 | 0,809549 | Unknown |
| GSM918618 | 0,009756 | 0,075906 | 0,011383 | 0,225095 | 0 | 0,67786 | 0 | 0 | 0,81797973 | 0,588089 | Unknown |
| GSM918628 | 0 | 0,016019 | 0,07443 | 0,158443 | 0 | 0,751108 | 0 | 0 | 0,842289815 | 0,546556 | Unknown |
| GSM918644 | 0,001839 | 0,228605 | 0 | 0,249077 | 0 | 0,520479 | 0 | 0 | 0,631841447 | 0,775605 | Unknown |
| GSM918580 | 0 | 0,013484 | 0,083301 | 0,232499 | 0 | 0,670716 | 0 | 0 | 0,78290609 | 0,628202 | WNT |
| GSM918593 | 0,000598 | 0,144525 | 0 | 0,326881 | 0 | 0,527996 | 0 | 0 | 0,618760574 | 0,784224 | WNT |
| GSM918603 | 0 | 0,110514 | 0,005825 | 0,293997 | 0 | 0,589664 | 0 | 0 | 0,639740205 | 0,767274 | WNT |
| GSM918625 | 0 | 0,079669 | 0,035443 | 0,275276 | 0 | 0,609613 | 0 | 0 | 0,70900619 | 0,706534 | WNT |
| GSM918638 | 0 | 0,20525 | 0,005706 | 0,260029 | 0 | 0,529014 | 0 | 0 | 0,568705493 | 0,821243 | WNT |
| GSM918641 | 0 | 0,089878 | 0,013671 | 0,252497 | 0 | 0,643954 | 0 | 0 | 0,755718511 | 0,659483 | WNT |
| GSM918642 | 0 | 0,178385 | 0,055696 | 0,259846 | 0 | 0,506074 | 0 | 0 | 0,538607926 | 0,841881 | WNT |
| GSM918643 | 0 | 0,066477 | 0,042738 | 0,324364 | 0 | 0,566421 | 0 | 0 | 0,612692094 | 0,789519 | WNT |
| average | 0,000549 | 0,106442 | 0,016102 | 0,33162 | 0,000549 | 0,53934 | 0,00539895 | | | | |

**SES_CIBERSORTx_GSE37418**

| Sample | Bcells | TCD4 | TCD8 | Tgd | NK | MoMaDC | granulocytes | P-value | Correlation | RMSE |
|---|---|---|---|---|---|---|---|---|---|---|
| GSM918578 | 0 | 0,183557 | 0 | 1,568301 | 0 | 1,970515 | 0 | 0 | 0,6527244 | 0,756816 |
| GSM918579 | 0 | 0,27968 | 0 | 1,251297 | 0 | 1,360963 | 0,02890987 | 0 | 0,521313151 | 0,858146 |
| GSM918580 | 0 | 0,10025 | 0,619303 | 1,72852 | 0 | 4,986452 | 0 | 0 | 0,78290609 | 0,628202 |
| GSM918581 | 0 | 0,686968 | 0 | 1,800173 | 0 | 2,819239 | 0 | 0 | 0,53895604 | 0,844336 |
| GSM918582 | 0 | 0,824627 | 0 | 2,171454 | 0 | 3,30038 | 0 | 0 | 0,727414629 | 0,694358 |
| GSM918583 | 0 | 0,813135 | 0 | 1,631768 | 0 | 1,866055 | 0 | 0 | 0,499878255 | 0,868487 |
| GSM918584 | 0 | 0,569863 | 0 | 1,415568 | 0 | 1,77628 | 0 | 0 | 0,530507863 | 0,848942 |
| GSM918585 | 0 | 0,371289 | 0 | 1,100937 | 0 | 1,29521 | 0 | 0 | 0,615534399 | 0,786797 |
| GSM918586 | 0,156738 | 0,910249 | 0 | 2,474101 | 0,087255 | 5,314441 | 0 | 0 | 0,745226239 | 0,674223 |
| GSM918587 | 0,002464 | 0,60601 | 0,298194 | 1,289141 | 0,047652 | 4,171295 | 0 | 0 | 0,861080381 | 0,54373 |
| GSM918588 | 0 | 0,239463 | 0 | 0,883945 | 0 | 0,748278 | 0,10898972 | 0 | 0,644504791 | 0,764481 |
| GSM918589 | 0 | 0,281817 | 0 | 0,673522 | 0 | 0,863927 | 0 | 0 | 0,543345164 | 0,839754 |
| GSM918590 | 0 | 0,128648 | 0,777846 | 2,102583 | 0 | 4,434433 | 0 | 0 | 0,712224729 | 0,703973 |
| GSM918591 | 0 | 0,269026 | 0 | 1,295588 | 0 | 1,301229 | 0 | 0 | 0,471107816 | 0,89494 |
| GSM918592 | 0 | 0,116059 | 0 | 1,56827 | 0 | 2,289141 | 0,14378433 | 0 | 0,636919967 | 0,770444 |
| GSM918593 | 0,001596 | 0,385754 | 0 | 0,872483 | 0 | 1,409282 | 0 | 0 | 0,618760574 | 0,784224 |
| GSM918594 | 0 | 0,695899 | 0 | 1,197432 | 0 | 2,674245 | 0 | 0 | 0,738920529 | 0,681943 |
| GSM918595 | 0 | 0,868003 | 0 | 2,589817 | 0 | 7,123974 | 0 | 0 | 0,775950041 | 0,635449 |
| GSM918596 | 0 | 0,364433 | 0 | 1,71077 | 0 | 3,140244 | 0 | 0 | 0,684991501 | 0,72799 |
| GSM918597 | 0 | 0,304978 | 0 | 1,039593 | 0 | 1,584164 | 0 | 0 | 0,550154392 | 0,837322 |
| GSM918598 | 0 | 0,511018 | 0 | 1,491362 | 0 | 1,694913 | 0 | 0 | 0,558704282 | 0,829278 |
| GSM918599 | 0 | 0,479896 | 0 | 1,306567 | 0 | 1,658137 | 0 | 0 | 0,618088138 | 0,784836 |
| GSM918600 | 0,006782 | 0,204529 | 0 | 0,765323 | 0,026552 | 1,107111 | 0 | 0 | 0,726498127 | 0,693487 |
| GSM918601 | 0 | 0,225597 | 0,179304 | 1,311977 | 0 | 1,840941 | 0 | 0 | 0,534048051 | 0,84921 |
| GSM918602 | 0 | 0,682375 | 0 | 1,262056 | 0 | 2,700531 | 0 | 0 | 0,753500263 | 0,668529 |
| GSM918603 | 0 | 0,480095 | 0,025304 | 1,277187 | 0 | 2,561628 | 0 | 0 | 0,639740205 | 0,767274 |
| GSM918604 | 0 | 0,286149 | 0 | 1,474122 | 0 | 2,633344 | 0 | 0 | 0,809303792 | 0,605312 |
| GSM918605 | 0 | 0,494871 | 0,015436 | 1,524354 | 0 | 2,290426 | 0 | 0 | 0,66108825 | 0,750159 |
| GSM918606 | 0 | 0,611231 | 0 | 1,274136 | 0 | 2,844958 | 0 | 0 | 0,658581927 | 0,751392 |
| GSM918607 | 0 | 0,642491 | 0,430844 | 2,042225 | 0 | 6,323302 | 0 | 0 | 0,858599791 | 0,540692 |
| GSM918608 | 0 | 0,273352 | 0 | 1,421968 | 0 | 3,407409 | 0 | 0 | 0,749673853 | 0,66318 |
| GSM918609 | 0 | 0,197351 | 0 | 0,595423 | 0 | 1,277287 | 0 | 0 | 0,722486335 | 0,693345 |
| GSM918610 | 0 | 0,380962 | 0 | 1,244554 | 0 | 1,639675 | 0 | 0 | 0,4638473 | 0,897534 |
| GSM918611 | 0 | 0,345735 | 0 | 0,745022 | 0 | 0,718157 | 0 | 0 | 0,697986611 | 0,723199 |
| GSM918612 | 0 | 0,131526 | 0,052931 | 0,944479 | 0 | 1,398878 | 0 | 0 | 0,749222808 | 0,669547 |
| GSM918613 | 0 | 0,314868 | 0 | 2,172788 | 0 | 2,956021 | 0,09230537 | 0 | 0,612409808 | 0,790184 |
| GSM918614 | 0 | 0,047719 | 0 | 0,228003 | 0 | 0,321457 | 0 | 0 | 0,592644438 | 0,805591 |
| GSM918615 | 0,001502 | 0,799077 | 0 | 2,819199 | 0 | 5,197621 | 0,20062044 | 0 | 0,734676686 | 0,683719 |
| GSM918616 | 0 | 0,555363 | 0 | 1,241193 | 0 | 1,521553 | 0,0394166 | 0 | 0,548080267 | 0,835866 |
| GSM918617 | 0 | 0,386847 | 0,027062 | 1,126375 | 0 | 1,704991 | 0 | 0 | 0,585912495 | 0,809549 |
| GSM918618 | 0,099661 | 0,775415 | 0,116281 | 2,299437 | 0 | 6,92462 | 0 | 0 | 0,81797973 | 0,588089 |
| GSM918619 | 0 | 0,420434 | 0 | 1,464056 | 0 | 2,861454 | 0 | 0 | 0,778059103 | 0,63952 |
| GSM918620 | 0 | 0,376324 | 0,168042 | 2,360325 | 0 | 4,634967 | 0 | 0 | 0,671419956 | 0,740145 |
| GSM918621 | 0 | 0,351575 | 0 | 1,465883 | 0 | 3,548536 | 0 | 0 | 0,851038968 | 0,548968 |
| GSM918622 | 0,0198 | 0,288988 | 0 | 1,251452 | 0 | 1,308584 | 0,08637812 | 0 | 0,566625916 | 0,823888 |
| GSM918623 | 0 | 0,539308 | 0 | 1,25022 | 0 | 1,876388 | 0 | 0 | 0,498484887 | 0,871822 |
| GSM918624 | 0 | 0,21269 | 0,023066 | 1,754768 | 0 | 3,549915 | 0 | 0 | 0,717660113 | 0,696393 |
| GSM918625 | 0 | 0,280741 | 0,124895 | 0,970032 | 0 | 2,148188 | 0 | 0 | 0,70900619 | 0,706534 |
| GSM918626 | 0 | 2,22914 | 1,827485 | 2,634491 | 0,157346 | 14,84286 | 0 | 0 | 0,825977119 | 0,580267 |
| GSM918627 | 0 | 0,760985 | 0 | 1,936066 | 0 | 1,881396 | 0,1440955 | 0 | 0,578375764 | 0,814343 |
| GSM918628 | 0 | 0,304358 | 1,41417 | 3,010416 | 0 | 14,27105 | 0 | 0 | 0,842289815 | 0,546556 |
| GSM918629 | 0 | 0,483824 | 0 | 1,509462 | 0,004622 | 1,911458 | 0 | 0 | 0,625663426 | 0,778895 |
| GSM918630 | 0 | 0,582936 | 0 | 1,426071 | 0 | 1,473827 | 0 | 0 | 0,471492241 | 0,888864 |
| GSM918631 | 0,002877 | 0,53179 | 0,196964 | 1,285777 | 0,016029 | 2,487012 | 0,05642902 | 0 | 0,540565236 | 0,842619 |
| GSM918632 | 0 | 0,068854 | 0,142392 | 0,954731 | 0 | 1,142156 | 0 | 0 | 0,555617118 | 0,834545 |
| GSM918633 | 0 | 0,286017 | 0 | 1,353521 | 0 | 2,11555 | 0 | 0 | 0,671703384 | 0,740275 |
| GSM918634 | 0 | 0,396633 | 0 | 2,63669 | 0 | 3,956866 | 0 | 0 | 0,589257565 | 0,8095 |
| GSM918635 | 0 | 0,373204 | 0,195563 | 1,146761 | 0 | 2,240598 | 0 | 0 | 0,441561448 | 0,914181 |
| GSM918636 | 0,002297 | 0,359813 | 0 | 0,961056 | 0 | 1,231351 | 0 | 0 | 0,582007313 | 0,812044 |
| GSM918637 | 0 | 0,399584 | 0,185773 | 0,959834 | 0 | 1,20939 | 0,26161096 | 0 | 0,653961179 | 0,760945 |
| GSM918638 | 0 | 0,688991 | 0,019154 | 0,872875 | 0 | 1,775811 | 0 | 0 | 0,568705493 | 0,821243 |
| GSM918639 | 0 | 0,420987 | 0 | 1,115197 | 0 | 1,703751 | 0,09927757 | 0 | 0,593287676 | 0,803682 |
| GSM918640 | 0 | 0,334342 | 0,334707 | 0,953868 | 0 | 1,65832 | 0 | 0 | 0,527351528 | 0,850685 |
| GSM918641 | 0 | 0,669193 | 0,101791 | 1,879984 | 0 | 4,79461 | 0 | 0 | 0,755718511 | 0,659483 |
| GSM918642 | 0 | 0,42333 | 0,132173 | 0,616648 | 0 | 1,20098 | 0 | 0 | 0,538607926 | 0,841881 |
| GSM918643 | 0 | 0,157626 | 0,101338 | 0,769115 | 0 | 1,343069 | 0 | 0 | 0,612692094 | 0,789519 |
| GSM918644 | 0,005095 | 0,633269 | 0 | 0,689979 | 0 | 1,441805 | 0 | 0 | 0,631841447 | 0,775605 |
| GSM918645 | 0 | 0,78315 | 0 | 1,517226 | 0 | 1,670384 | 0,0581185 | 0 | 0,509443562 | 0,86147 |
| GSM918646 | 0 | 0,069408 | 0,277996 | 1,256349 | 0 | 2,319391 | 0 | 0 | 0,633229101 | 0,773426 |
| GSM918647 | 0 | 0,528922 | 0,008834 | 0,78564 | 0 | 1,26007 | 0 | 0 | 0,534670496 | 0,844582 |
| GSM918648 | 0 | 0,32173 | 0 | 1,240066 | 0 | 1,724501 | 0 | 0 | 0,569415941 | 0,822842 |
| GSM918649 | 0 | 0 | 0,601399 | 2,150892 | 0 | 4,041908 | 0 | 0 | 0,816036975 | 0,598646 |
| GSM918650 | 0 | 0,383092 | 0 | 0,997935 | 0 | 0,799513 | 0,13143888 | 0 | 0,425347739 | 0,917313 |
| GSM918651 | 0 | 0,733782 | 0,106856 | 1,284625 | 0 | 2,248036 | 0 | 0 | 0,677514809 | 0,739274 |
| GSM918652 | 0 | 0,719068 | 0 | 1,472025 | 0 | 2,149407 | 0 | 0 | 0,542519369 | 0,840117 |
| GSM918653 | 0 | 0,313887 | 0 | 2,056084 | 0 | 2,104063 | 0 | 0 | 0,440506923 | 0,918691 |
| **average** | **0,003932** | **0,450713** | **0,111909** | **1,433199** | **0,004467** | **2,737893** | **0,01909704** | | | |
