## Supplemental Table 3 for "EphA2 and Phosphoantigen-Mediated Selective Killing of Medulloblastoma by γδT Cells Preserves Neuronal and Stem Cell Integrity"

**Table S3**

Boutin et al.

**Ephrin-A2 and Phosphoantigen-Mediated Selective Killing of Medulloblastoma by γδT Cells Preserves Neuronal and Stem Cell Integrity**

**CIBERSORTx_GSE85217_WNT**

| Sample | Bcells | TCD4 | TCD8 | Tgd | NK | MoMaDC | granulocytes | P-value | Correlatior | RMSE | ratio mono/T cells |
|---|---|---|---|---|---|---|---|---|---|---|---|
| GSM2261548 | 0 | 0,202847 | 0,009953 | 0,051976 | 0 | 0,735224 | 0 | 0 | 0,601161 | 0,799082 | 2,776778664 |
| GSM2261551 | 0 | 0,230386 | 0 | 0,063471 | 0 | 0,642135 | 0,06400721 | 0 | 0,495996 | 0,870733 | 2,185192822 |
| GSM2261553 | 0 | 0,1026 | 0 | 0,082732 | 0 | 0,731776 | 0,082891911 | 0 | 0,752767 | 0,667515 | 3,948446011 |
| GSM2261580 | 0 | 0,181016 | 0 | 0,083852 | 0 | 0,735132 | 0 | 0 | 0,646753 | 0,763604 | 2,775469891 |
| GSM2261589 | 0 | 0,222129 | 0 | 0,14597 | 0 | 0,624449 | 0,007452584 | 0 | 0,427527 | 0,909766 | 1,696419392 |
| GSM2261591 | 0 | 0,15856 | 0,048542 | 0,07621 | 0 | 0,716689 | 0 | 0 | 0,603858 | 0,796594 | 2,529690511 |
| GSM2261609 | 0,003235 | 0,281271 | 0 | 0,071934 | 0 | 0,630222 | 0,013338154 | 0 | 0,494501 | 0,870499 | 1,78429972 |
| GSM2261648 | 0 | 0,297956 | 0 | 0,126718 | 0 | 0,505481 | 0,069844985 | 0 | 0,428749 | 0,905268 | 1,190278652 |
| GSM2261687 | 0,00784 | 0,192423 | 0 | 0,066743 | 0 | 0,732994 | 0 | 0 | 0,650231 | 0,761303 | 2,828283218 |
| GSM2261702 | 0 | 0,270884 | 0 | 0,082514 | 0 | 0,646602 | 0 | 0 | 0,484585 | 0,876787 | 1,829671303 |
| GSM2261706 | 0 | 0,122613 | 0,039936 | 0,028074 | 0,004353 | 0,805023 | 0 | 0 | 0,650917 | 0,758661 | 4,223106776 |
| GSM2261707 | 0 | 0,193827 | 0,062548 | 0,031762 | 0 | 0,711863 | 0 | 0 | 0,656084 | 0,758007 | 2,470576124 |
| GSM2261712 | 0 | 0,268773 | 0 | 0,131454 | 0 | 0,531952 | 0,067820943 | 0 | 0,443152 | 0,897929 | 1,329126393 |
| GSM2261713 | 0 | 0,289174 | 0 | 0,155326 | 0 | 0,455228 | 0,100272243 | 0,01 | 0,327465 | 0,956319 | 1,02413381 |
| GSM2261715 | 0 | 0,182856 | 0,007463 | 0,128899 | 0,008961 | 0,619611 | 0,052209699 | 0 | 0,621526 | 0,786082 | 1,941030264 |
| GSM2261717 | 0 | 0,266114 | 0,033853 | 0,117162 | 0 | 0,573514 | 0,009357517 | 0 | 0,444368 | 0,89774 | 1,374907953 |
| GSM2261726 | 0 | 0,280101 | 0 | 0,074539 | 0 | 0,643271 | 0,002088981 | 0 | 0,467365 | 0,887389 | 1,813870584 |
| GSM2261741 | 0 | 0,292879 | 0 | 0,075263 | 0 | 0,631858 | 0 | 0 | 0,499357 | 0,867485 | 1,716343972 |
| GSM2261752 | 0 | 0,21355 | 0,04358 | 0,079559 | 0 | 0,432461 | 0,230851087 | 0 | 0,619838 | 0,792198 | 1,28445366 |
| GSM2261766 | 0 | 0,246785 | 0 | 0,115608 | 0 | 0,576879 | 0,060727775 | 0 | 0,393972 | 0,927413 | 1,591860252 |
| GSM2261771 | 0 | 0,276457 | 0 | 0,077898 | 0 | 0,593149 | 0,052495762 | 0 | 0,472319 | 0,88325 | 1,673880253 |
| GSM2261822 | 0 | 0,231146 | 0,011228 | 0,059906 | 0,010203 | 0,563474 | 0,124042206 | 0 | 0,477189 | 0,880458 | 1,864080211 |
| GSM2261837 | 0 | 0,199828 | 0,114011 | 0,077993 | 0 | 0,541201 | 0,066967071 | 0 | 0,490694 | 0,870835 | 1,381204768 |
| GSM2261849 | 0 | 0,186156 | 0,019278 | 0,111531 | 0 | 0,661065 | 0,021970035 | 0 | 0,477462 | 0,881748 | 2,085606501 |
| GSM2261862 | 0 | 0,285984 | 0 | 0,111258 | 0 | 0,558324 | 0,044434061 | 0 | 0,38411 | 0,931173 | 1,405500176 |
| GSM2261890 | 0 | 0,24424 | 0 | 0,113041 | 0 | 0,596022 | 0,046696534 | 0 | 0,532397 | 0,846199 | 1,66821324 |
| GSM2261905 | 0 | 0,258541 | 0,008895 | 0,096104 | 0,001757 | 0,489725 | 0,144977412 | 0 | 0,623212 | 0,791486 | 1,347096345 |
| GSM2261930 | 0 | 0,158246 | 0,010114 | 0,077596 | 0 | 0,698481 | 0,055563593 | 0 | 0,636453 | 0,77174 | 2,839866138 |
| GSM2261933 | 0 | 0,359245 | 0 | 0,107839 | 0,007014 | 0,443297 | 0,0826042 | 0 | 0,382084 | 0,928264 | 0,94907318 |
| GSM2261958 | 0,007505 | 0,285152 | 0 | 0,110347 | 0 | 0,526307 | 0,070689172 | 0 | 0,521774 | 0,853607 | 1,330742907 |
| GSM2261971 | 0 | 0,243077 | 0,020688 | 0,083016 | 0 | 0,488494 | 0,164724697 | 0 | 0,380234 | 0,933487 | 1,408651294 |
| GSM2261973 | 0 | 0,271411 | 0 | 0,119691 | 0 | 0,590298 | 0,018600506 | 0 | 0,515824 | 0,856524 | 1,509319417 |
| GSM2261975 | 0 | 0,184448 | 0,043836 | 0,039236 | 0 | 0,641548 | 0,090931873 | 0 | 0,55822 | 0,829593 | 2,398127776 |
| GSM2261986 | 0,00124 | 0,249606 | 0 | 0,063378 | 0 | 0,662343 | 0,023433706 | 0 | 0,631936 | 0,778715 | 2,11622088 |
| GSM2261990 | 0 | 0,226879 | 0,036402 | 0,09891 | 0 | 0,579761 | 0,05804753 | 0 | 0,535505 | 0,844257 | 1,600706202 |
| GSM2262002 | 0 | 0,369887 | 0 | 0,153635 | 0 | 0,410726 | 0,065752073 | 0 | 0,33575 | 0,950311 | 0,784542846 |
| GSM2262024 | 0 | 0,272155 | 0 | 0,03233 | 0 | 0,695515 | 0 | 0 | 0,617616 | 0,788453 | 2,284234897 |
| GSM2262040 | 0 | 0,309325 | 0 | 0,126635 | 0 | 0,552313 | 0,011726849 | 0 | 0,418294 | 0,911784 | 1,266888876 |
| GSM2262069 | 0 | 0,265867 | 0,010407 | 0,097151 | 0 | 0,581999 | 0,044575093 | 0 | 0,478341 | 0,878974 | 1,558541492 |
| GSM2262071 | 0 | 0,369748 | 0 | 0,087348 | 0 | 0,491704 | 0,051199264 | 0 | 0,366708 | 0,937655 | 1,075710263 |
| GSM2262088 | 0 | 0,25512 | 0 | 0,076265 | 0 | 0,642922 | 0,025692847 | 0 | 0,48021 | 0,879788 | 1,940107295 |
| GSM2262095 | 0 | 0,184 | 0 | 0,108016 | 0 | 0,688758 | 0,019225697 | 0 | 0,608543 | 0,79352 | 2,358624745 |
| GSM2262113 | 0,001595 | 0,277493 | 0,023408 | 0,065719 | 0 | 0,61509 | 0,016694575 | 0 | 0,537758 | 0,843509 | 1,677730455 |
| GSM2262121 | 0 | 0,312992 | 0 | 0,088829 | 0 | 0,56968 | 0,028499006 | 0 | 0,375882 | 0,936192 | 1,417744909 |
| GSM2262136 | 0 | 0,178014 | 0 | 0,09304 | 0 | 0,726567 | 0,00238027 | 0 | 0,579535 | 0,81457 | 2,680531852 |
| GSM2262147 | 0 | 0,172751 | 0,029465 | 0,082473 | 0 | 0,715311 | 0 | 0 | 0,573235 | 0,818926 | 2,512602228 |
| GSM2262154 | 0 | 0,21083 | 0,029644 | 0,078914 | 0 | 0,654238 | 0,026373333 | 0 | 0,46646 | 0,888574 | 2,048408961 |
| GSM2262160 | 0,000522 | 0,265497 | 0,030302 | 0,075124 | 0 | 0,62837 | 0,000185299 | 0 | 0,630723 | 0,782532 | 1,694069608 |
| GSM2262171 | 0 | 0,348738 | 0 | 0,048763 | 0,006992 | 0,433744 | 0,161763263 | 0 | 0,537696 | 0,846308 | 1,091179185 |
| GSM2262172 | 0 | 0,337613 | 0 | 0,095994 | 0 | 0,566392 | 0 | 0 | 0,528923 | 0,849809 | 1,306231759 |
| GSM2262197 | 0 | 0,234439 | 0 | 0,100045 | 0 | 0,665516 | 0 | 0 | 0,549377 | 0,835332 | 1,9896801 |
| GSM2262210 | 0,002684 | 0,257471 | 0,003313 | 0,083007 | 0 | 0,611948 | 0,041577589 | 0 | 0,513449 | 0,85839 | 1,780000345 |
| GSM2262212 | 0 | 0,286277 | 0 | 0,08499 | 0 | 0,604633 | 0,02410046 | 0 | 0,507229 | 0,862239 | 1,628565269 |
| GSM2262214 | 0 | 0,221815 | 0,006724 | 0,080304 | 0,002879 | 0,680318 | 0,007961011 | 0 | 0,599285 | 0,801004 | 2,202803006 |
| GSM2262220 | 0 | 0,190935 | 0,067602 | 0,043792 | 0 | 0,697671 | 0 | 0 | 0,593125 | 0,805085 | 2,30765542 |
| GSM2262222 | 0 | 0,364278 | 0 | 0,107976 | 0 | 0,504768 | 0,022978307 | 0 | 0,38142 | 0,930139 | 1,068848281 |
| GSM2262223 | 0,001351 | 0,276439 | 0 | 0,133443 | 0 | 0,588767 | 0 | 0 | 0,391807 | 0,927735 | 1,436434327 |
| GSM2262225 | 0,003118 | 0,363386 | 0 | 0,117105 | 0 | 0,501064 | 0,015327174 | 0 | 0,501586 | 0,866181 | 1,042817544 |
| GSM2262230 | 0,016777 | 0,075026 | 0,088128 | 0,026459 | 0,00829 | 0,785319 | 0 | 0 | 0,770196 | 0,650453 | 4,141688821 |
| GSM2262232 | 0 | 0,194163 | 0,036081 | 0,137643 | 0 | 0,632113 | 0 | 0 | 0,586991 | 0,810163 | 1,718226336 |
| GSM2262245 | 0 | 0,303648 | 0,011658 | 0,08275 | 0 | 0,57883 | 0,023113432 | 0 | 0,420081 | 0,912262 | 1,45413856 |
| GSM2262250 | 0,001545 | 0,360328 | 0 | 0,13928 | 0 | 0,434577 | 0,064271135 | 0 | 0,376134 | 0,93091 | 0,869836116 |
| GSM2262252 | 0 | 0,250175 | 0 | 0,060958 | 0 | 0,688867 | 0 | 0 | 0,585842 | 0,810857 | 2,214055884 |
| GSM2262257 | 0 | 0,311845 | 0 | 0,139699 | 0 | 0,47936 | 0,06909674 | 0 | 0,339814 | 0,950503 | 1,061601758 |
| GSM2262267 | 0 | 0,283765 | 0 | 0,117009 | 0 | 0,561461 | 0,03776464 | 0 | 0,457289 | 0,890564 | 1,400942426 |
| GSM2262279 | 0 | 0,349345 | 0 | 0,150509 | 0 | 0,432048 | 0,068097266 | 0 | 0,381033 | 0,92843 | 0,864348683 |
| GSM2262280 | 0 | 0,15559 | 0,080586 | 0 | 0,008489 | 0,755335 | 0 | 0 | 0,745495 | 0,67854 | 3,198188134 |
| GSM2262281 | 0 | 0,295891 | 0 | 0,108852 | 0 | 0,595256 | 0 | 0 | 0,474648 | 0,881185 | 1,470699536 |
| GSM2262292 | 0 | 0,273428 | 0 | 0,067554 | 0 | 0,655728 | 0,003289483 | 0 | 0,575192 | 0,81883 | 1,923053852 |
| GSM2262304 | 0 | 0,307258 | 0,033101 | 0,089618 | 0 | 0,527071 | 0,042952588 | 0 | 0,380803 | 0,931274 | 1,225814456 |
| average | 0,000677 | 0,252038 | 0,013725 | 0,090268 | 0,000842 | 0,603855 | 0,038594841 | | | | |

**SES_CIBERSORTx_GSE85217_WNT**

| Sample | Bcells | TCD4 | TCD8 | Tgd | NK | MoMaDC | granulocytes | P-value | Correlatior | RMSE | | Tab |
|---|---|---|---|---|---|---|---|---|---|---|---|---|
| GSM2261548 | 0 | 1,219359 | 0,059829 | 0,312437 | 0 | 4,41959 | 0 | 0 | 0,601161 | 0,799082 | | 1,279188 |
| GSM2261551 | 0 | 0,650311 | 0 | 0,179161 | 0 | 1,812556 | 0,180673252 | 0 | 0,495996 | 0,870733 | | 0,650311 |
| GSM2261553 | 0 | 0,970223 | 0 | 0,782345 | 0 | 6,919919 | 0,783854127 | 0 | 0,752767 | 0,667515 | | 0,970223 |
| GSM2261580 | 0 | 0,773 | 0 | 0,358076 | 0 | 3,139267 | 0 | 0 | 0,646753 | 0,763604 | | 0,773 |
| GSM2261589 | 0 | 0,600414 | 0 | 0,394556 | 0 | 1,687885 | 0,020144328 | 0 | 0,427527 | 0,909766 | | 0,600414 |
| GSM2261591 | 0 | 0,628135 | 0,192298 | 0,301905 | 0 | 2,839168 | 0 | 0 | 0,603858 | 0,796594 | | 0,820433 |
| GSM2261609 | 0,010798 | 0,938674 | 0 | 0,240061 | 0 | 2,103217 | 0,044512929 | 0 | 0,494501 | 0,870499 | | 0,938674 |
| GSM2261648 | 0 | 0,789512 | 0 | 0,335772 | 0 | 1,339401 | 0,185072278 | 0 | 0,428749 | 0,905268 | | 0,789512 |
| GSM2261687 | 0,051988 | 1,275957 | 0 | 0,442572 | 0 | 4,860486 | 0 | 0 | 0,650231 | 0,761303 | | 1,275957 |
| GSM2261702 | 0 | 0,923946 | 0 | 0,281444 | 0 | 2,205469 | 0 | 0 | 0,484585 | 0,876787 | | 0,923946 |
| GSM2261706 | 0 | 1,107842 | 0,360831 | 0,25366 | 0,039331 | 7,273597 | 0 | 0 | 0,650917 | 0,758661 | | 1,468673 |
| GSM2261707 | 0 | 0,62448 | 0,201519 | 0,102331 | 0 | 2,293509 | 0 | 0 | 0,656084 | 0,758007 | | 0,825999 |
| GSM2261712 | 0 | 0,780267 | 0 | 0,381621 | 0 | 1,544295 | 0,196889086 | 0 | 0,443152 | 0,897929 | | 0,780267 |
| GSM2261713 | 0 | 1,231312 | 0 | 0,661385 | 0 | 1,938375 | 0,426962692 | 0,01 | 0,327465 | 0,956319 | | 1,231312 |
| GSM2261715 | 0 | 2,384368 | 0,09731 | 1,680786 | 0,11685 | 8,07947 | 0,680792414 | 0 | 0,621526 | 0,786082 | | 2,481679 |
| GSM2261717 | 0 | 0,576193 | 0,073299 | 0,253681 | 0 | 1,241779 | 0,020261018 | 0 | 0,444368 | 0,89774 | | 0,649492 |
| GSM2261726 | 0 | 0,843475 | 0 | 0,224461 | 0 | 1,937098 | 0,006290599 | 0 | 0,467365 | 0,887389 | | 0,843475 |
| GSM2261741 | 0 | 0,966768 | 0 | 0,248436 | 0 | 2,085709 | 0 | 0 | 0,499357 | 0,867485 | | 0,966768 |
| GSM2261752 | 0 | 0,607497 | 0,123974 | 0,226325 | 0 | 1,230244 | 0,656714684 | 0 | 0,619838 | 0,792198 | | 0,731471 |
| GSM2261766 | 0 | 0,664672 | 0 | 0,311371 | 0 | 1,553724 | 0,163559788 | 0 | 0,393972 | 0,927413 | | 0,664672 |
| GSM2261771 | 0 | 1,177005 | 0 | 0,331648 | 0 | 2,525304 | 0,223498333 | 0 | 0,472319 | 0,88325 | | 1,177005 |
| GSM2261822 | 0 | 1,045304 | 0,050777 | 0,270909 | 0,046142 | 2,548178 | 0,560951152 | 0 | 0,477189 | 0,880458 | | 1,096081 |
| GSM2261837 | 0 | 0,418372 | 0,238701 | 0,163292 | 0 | 1,133092 | 0,140206462 | 0 | 0,490694 | 0,870835 | | 0,657073 |
| GSM2261849 | 0 | 0,314334 | 0,032551 | 0,188326 | 0 | 1,11624 | 0,037097479 | 0 | 0,477462 | 0,881748 | | 0,346885 |
| GSM2261862 | 0 | 0,499862 | 0 | 0,194464 | 0 | 0,975875 | 0,077664735 | 0 | 0,38411 | 0,931173 | | 0,499862 |
| GSM2261890 | 0 | 0,619944 | 0 | 0,286927 | 0 | 1,512854 | 0,118527598 | 0 | 0,532397 | 0,846199 | | 0,619944 |
| GSM2261905 | 0 | 1,110632 | 0,038213 | 0,412842 | 0,007548 | 2,103742 | 0,622788789 | 0 | 0,623212 | 0,791486 | | 1,148844 |
| GSM2261930 | 0 | 0,985164 | 0,062964 | 0,483073 | 0 | 4,348406 | 0,345912187 | 0 | 0,636453 | 0,77174 | | 1,048128 |
| GSM2261933 | 0 | 0,559936 | 0 | 0,168083 | 0,010933 | 0,690944 | 0,128750817 | 0 | 0,382084 | 0,928264 | | 0,559936 |
| GSM2261958 | 0,011552 | 0,438914 | 0 | 0,169848 | 0 | 0,810105 | 0,108806563 | 0 | 0,521774 | 0,853607 | | 0,438914 |
| GSM2261971 | 0 | 0,586076 | 0,049881 | 0,200159 | 0 | 1,177795 | 0,397163327 | 0 | 0,380234 | 0,933487 | | 0,635957 |
| GSM2261973 | 0 | 0,703212 | 0 | 0,310114 | 0 | 1,529432 | 0,048192981 | 0 | 0,515824 | 0,856524 | | 0,703212 |
| GSM2261975 | 0 | 0,888188 | 0,211086 | 0,188936 | 0 | 3,089292 | 0,437870808 | 0 | 0,55822 | 0,829593 | | 1,099273 |
| GSM2261986 | 0,005129 | 1,032522 | 0 | 0,26217 | 0 | 2,739855 | 0,09693615 | 0 | 0,631936 | 0,778715 | | 1,032522 |
| GSM2261990 | 0 | 0,56255 | 0,090259 | 0,245249 | 0 | 1,437528 | 0,143929784 | 0 | 0,535505 | 0,844257 | | 0,652809 |
| GSM2262002 | 0 | 0,893512 | 0 | 0,371127 | 0 | 0,992164 | 0,158833099 | 0 | 0,33575 | 0,950311 | | 0,893512 |
| GSM2262024 | 0 | 1,63826 | 0 | 0,194614 | 0 | 4,186717 | 0 | 0 | 0,617616 | 0,788453 | | 1,63826 |
| GSM2262040 | 0 | 0,932501 | 0 | 0,381758 | 0 | 1,66502 | 0,035352118 | 0 | 0,418293 | 0,911784 | | 0,932501 |
| GSM2262069 | 0 | 0,680836 | 0,026651 | 0,248785 | 0 | 1,490391 | 0,114148486 | 0 | 0,478341 | 0,878974 | | 0,707488 |
| GSM2262071 | 0 | 0,546359 | 0 | 0,12907 | 0 | 0,726566 | 0,075654574 | 0 | 0,366708 | 0,937655 | | 0,546359 |
| GSM2262088 | 0 | 0,492339 | 0 | 0,147179 | 0 | 1,240733 | 0,049582934 | 0 | 0,48021 | 0,879788 | | 0,492339 |
| GSM2262095 | 0 | 0,472592 | 0 | 0,277432 | 0 | 1,769026 | 0,049379866 | 0 | 0,608543 | 0,79352 | | 0,472592 |
| GSM2262113 | 0,005425 | 0,943827 | 0,079617 | 0,223527 | 0 | 2,09208 | 0,056782546 | 0 | 0,537758 | 0,843509 | | 1,023443 |
| GSM2262121 | 0 | 0,547072 | 0 | 0,155263 | 0 | 0,995732 | 0,04981286 | 0 | 0,375882 | 0,936192 | | 0,547072 |
| GSM2262136 | 0 | 1,307251 | 0 | 0,68324 | 0 | 5,335573 | 0,017479619 | 0 | 0,579535 | 0,81457 | | 1,307251 |
| GSM2262147 | 0 | 0,689361 | 0,117579 | 0,32911 | 0 | 2,854441 | 0 | 0 | 0,573235 | 0,818926 | | 0,80694 |
| GSM2262154 | 0 | 0,432261 | 0,060778 | 0,161797 | 0 | 1,341372 | 0,054072745 | 0 | 0,46646 | 0,888574 | | 0,493039 |
| GSM2262160 | 0,002435 | 1,239096 | 0,14142 | 0,350611 | 0 | 2,932648 | 0,000864805 | 0 | 0,630723 | 0,782532 | | 1,380515 |
| GSM2262171 | 0 | 0,828916 | 0 | 0,115904 | 0,01662 | 1,030968 | 0,384495868 | 0 | 0,537696 | 0,846308 | | 0,828916 |
| GSM2262172 | 0 | 0,893341 | 0 | 0,254006 | 0 | 1,4987 | 0 | 0 | 0,528923 | 0,849809 | | 0,893341 |
| GSM2262197 | 0 | 0,93589 | 0 | 0,399386 | 0 | 2,656772 | 0 | 0 | 0,549377 | 0,835332 | | 0,93589 |
| GSM2262210 | 0,008346 | 0,800676 | 0,010302 | 0,258134 | 0 | 1,903019 | 0,1292969 | 0 | 0,513449 | 0,85839 | | 0,810978 |
| GSM2262212 | 0 | 0,868382 | 0 | 0,257807 | 0 | 1,834073 | 0,073105568 | 0 | 0,507229 | 0,862239 | | 0,868382 |
| GSM2262214 | 0 | 0,831889 | 0,025216 | 0,30117 | 0,010798 | 2,551452 | 0,029856826 | 0 | 0,599285 | 0,801004 | | 0,857105 |
| GSM2262220 | 0 | 0,717963 | 0,254198 | 0,164668 | 0 | 2,623408 | 0 | 0 | 0,593125 | 0,805085 | | 0,972161 |
| GSM2262222 | 0 | 1,098842 | 0 | 0,325708 | 0 | 1,522628 | 0,069313887 | 0 | 0,38142 | 0,930139 | | 1,098842 |
| GSM2262223 | 0,002961 | 0,60565 | 0 | 0,29236 | 0 | 1,289931 | 0 | 0 | 0,391807 | 0,927735 | | 0,60565 |
| GSM2262225 | 0,008029 | 0,935722 | 0 | 0,301546 | 0 | 1,290244 | 0,039467598 | 0 | 0,501586 | 0,866181 | | 0,935722 |
| GSM2262230 | 0,187793 | 0,839785 | 0,986439 | 0,296164 | 0,09279 | 8,79027 | 0 | 0 | 0,770196 | 0,650453 | | 1,826224 |
| GSM2262232 | 0 | 0,52669 | 0,097873 | 0,373372 | 0 | 1,714677 | 0 | 0 | 0,586991 | 0,810163 | | 0,624563 |
| GSM2262245 | 0 | 0,71402 | 0,027414 | 0,194585 | 0 | 1,361102 | 0,054350575 | 0 | 0,420081 | 0,912262 | | 0,741434 |
| GSM2262250 | 0,003238 | 0,755281 | 0 | 0,291944 | 0 | 0,910915 | 0,134718479 | 0 | 0,376134 | 0,93091 | | 0,755281 |
| GSM2262252 | 0 | 0,915934 | 0 | 0,223179 | 0 | 2,522059 | 0 | 0 | 0,585842 | 0,810857 | | 0,915934 |
| GSM2262257 | 0 | 0,547718 | 0 | 0,245363 | 0 | 0,841936 | 0,121359928 | 0 | 0,339814 | 0,950503 | | 0,547718 |
| GSM2262267 | 0 | 0,726239 | 0 | 0,29946 | 0 | 1,436944 | 0,096650788 | 0 | 0,457289 | 0,890564 | | 0,726239 |
| GSM2262279 | 0 | 0,721178 | 0 | 0,310706 | 0 | 0,891908 | 0,140577919 | 0 | 0,381033 | 0,92843 | | 0,721178 |
| GSM2262280 | 0 | 1,981149 | 1,026112 | 0 | 0,108094 | 9,617787 | 0 | 0 | 0,745495 | 0,67854 | | 3,007261 |
| GSM2262281 | 0 | 0,857169 | 0 | 0,315335 | 0 | 1,724401 | 0 | 0 | 0,474648 | 0,881185 | | 0,857169 |
| GSM2262292 | 0 | 1,001759 | 0 | 0,247499 | 0 | 2,402391 | 0,012051686 | 0 | 0,575192 | 0,81883 | | 1,001759 |
| GSM2262304 | 0 | 0,622971 | 0,067112 | 0,181701 | 0 | 1,068645 | 0,087087085 | 0 | 0,380803 | 0,931274 | | 0,690083 |
| **average** | **0,004253** | **0,843441** | **0,068631** | **0,30217** | **0,006416** | **2,390716** | **0,126976016** | | | | | |

**CIBERSORTx_GSE85217_SHH**

| Sample | Bcells | TCD4 | TCD8 | Tgd | NK | MoMaDC | granulocytes | P-value | Correlation | RMSE | ratio mono/T cells |
|---|---|---|---|---|---|---|---|---|---|---|---|
| GSM2261538 | 0 | 0,203724 | 0,000668 | 0,089941 | 0 | 0,705667 | 0 | 0 | 0,6265381 | 0,780385 | 2,39750792 |
| GSM2261543 | 0 | 0,170766 | 0 | 0,092437 | 0 | 0,736797 | 0 | 0 | 0,7034297 | 0,717266 | 2,799353411 |
| GSM2261544 | 0 | 0,20648 | 0 | 0,168306 | 0 | 0,590838 | 0,03437534 | 0 | 0,5703643 | 0,821555 | 1,576466944 |
| GSM2261547 | 0 | 0,149637 | 0 | 0,119766 | 0 | 0,730597 | 0 | 0 | 0,6249725 | 0,780146 | 2,711917793 |
| GSM2261549 | 0 | 0,106643 | 0 | 0,12249 | 0 | 0,770868 | 0 | 0 | 0,7189782 | 0,699516 | 3,36429062 |
| GSM2261550 | 0 | 0,136431 | 0 | 0,125728 | 0 | 0,737842 | 0 | 0 | 0,6567303 | 0,75483 | 2,814488661 |
| GSM2261552 | 0 | 0,194697 | 0 | 0,12945 | 0 | 0,675853 | 0 | 0 | 0,6039521 | 0,797317 | 2,085017629 |
| GSM2261567 | 0 | 0,145612 | 0 | 0,16144 | 0 | 0,678378 | 0,01457062 | 0 | 0,5510153 | 0,833665 | 2,2093295 |
| GSM2261568 | 0 | 0,101089 | 0,025413 | 0,088525 | 0 | 0,784972 | 0 | 0 | 0,6969537 | 0,718482 | 3,650566047 |
| GSM2261569 | 0 | 0,159695 | 0 | 0,149374 | 0 | 0,690931 | 0 | 0 | 0,619177 | 0,785402 | 2,235519279 |
| GSM2261570 | 0 | 0,094101 | 0,003162 | 0,064013 | 0 | 0,838723 | 0 | 0 | 0,7388039 | 0,675543 | 5,200528963 |
| GSM2261571 | 0 | 0,231641 | 0 | 0,11844 | 0 | 0,649919 | 0 | 0 | 0,6126945 | 0,792715 | 1,856479117 |
| GSM2261574 | 0 | 0,137161 | 0 | 0,121307 | 0 | 0,741533 | 0 | 0 | 0,7195002 | 0,702351 | 2,868959263 |
| GSM2261586 | 0 | 0,161956 | 0 | 0,12844 | 0 | 0,709605 | 0 | 0 | 0,5840615 | 0,810819 | 2,443580797 |
| GSM2261597 | 0 | 0,158991 | 0 | 0,098579 | 0 | 0,74243 | 0 | 0 | 0,6484882 | 0,761643 | 2,882444077 |
| GSM2261598 | 0,00042 | 0,208781 | 0 | 0,137174 | 0 | 0,653625 | 0 | 0 | 0,4562617 | 0,894063 | 1,889334584 |
| GSM2261600 | 0 | 0,158242 | 0 | 0,147882 | 0 | 0,693876 | 0 | 0 | 0,54886 | 0,835383 | 2,266646925 |
| GSM2261602 | 0 | 0,171838 | 0 | 0,082983 | 0 | 0,74518 | 0 | 0 | 0,6613598 | 0,751422 | 2,924331403 |
| GSM2261603 | 0 | 0,150681 | 0,018927 | 0,050152 | 0 | 0,78024 | 0 | 0 | 0,7009871 | 0,71586 | 3,550428723 |
| GSM2261614 | 0 | 0,146845 | 0,002556 | 0,181526 | 0 | 0,623766 | 0,04530739 | 0 | 0,503457 | 0,864213 | 1,884907122 |
| GSM2261615 | 0 | 0,140762 | 0,017006 | 0,063603 | 0,00219 | 0,776439 | 0 | 0 | 0,6499408 | 0,759605 | 3,507413289 |
| GSM2261617 | 0 | 0,184059 | 0 | 0,11798 | 0 | 0,697961 | 0 | 0 | 0,6380871 | 0,771673 | 2,310831118 |
| GSM2261619 | 0,000832 | 0,325888 | 0 | 0,114077 | 0 | 0,559203 | 0 | 0 | 0,4724567 | 0,882035 | 1,271018653 |
| GSM2261620 | 0 | 0,148149 | 0 | 0,184496 | 0 | 0,667355 | 0 | 0 | 0,5213626 | 0,85329 | 2,006206867 |
| GSM2261622 | 0 | 0,240823 | 0,060515 | 0,071484 | 0 | 0,627178 | 0 | 0 | 0,7075634 | 0,7283 | 1,682246079 |
| GSM2261629 | 0 | 0,091182 | 0 | 0,112059 | 0,000623 | 0,653883 | 0,142253 | 0 | 0,6940832 | 0,723605 | 3,217276975 |
| GSM2261633 | 0 | 0,144434 | 0 | 0,05522 | 0 | 0,800346 | 0 | 0 | 0,7726248 | 0,646009 | 4,008673665 |
| GSM2261634 | 0 | 0,17637 | 0 | 0,163359 | 0 | 0,66027 | 0 | 0 | 0,6579396 | 0,758588 | 1,943517415 |
| GSM2261641 | 0 | 0,225333 | 0 | 0,129335 | 0 | 0,640993 | 0,00433863 | 0 | 0,6537476 | 0,764259 | 1,807299455 |
| GSM2261644 | 0 | 0,093459 | 0,001163 | 0,14743 | 0,008754 | 0,749193 | 0 | 0 | 0,7377433 | 0,684149 | 3,095162555 |
| GSM2261647 | 0 | 0,112137 | 0,01101 | 0,068804 | 0,00843 | 0,79962 | 0 | 0 | 0,7478752 | 0,670047 | 4,165763842 |
| GSM2261650 | 0 | 0,170686 | 0 | 0,163949 | 0 | 0,665365 | 0 | 0 | 0,6768178 | 0,744034 | 1,988326688 |
| GSM2261655 | 0 | 0,145892 | 0,0495 | 0,1246 | 0 | 0,680008 | 0 | 0 | 0,7089821 | 0,718333 | 2,125073781 |
| GSM2261657 | 0 | 0,163423 | 0 | 0,177986 | 0 | 0,633554 | 0,02503801 | 0 | 0,6324306 | 0,777584 | 1,855708019 |
| GSM2261658 | 0,012305 | 0,101808 | 0,064098 | 0,125576 | 0 | 0,696213 | 0 | 0 | 0,7598601 | 0,674082 | 2,388529262 |
| GSM2261661 | 0 | 0,189945 | 0 | 0,148087 | 0 | 0,661968 | 0 | 0 | 0,5624336 | 0,825973 | 1,958296314 |
| GSM2261673 | 0 | 0,1336 | 0,007549 | 0,13756 | 0 | 0,721292 | 0 | 0 | 0,6910543 | 0,727982 | 2,587980596 |
| GSM2261674 | 0 | 0,10733 | 0,076448 | 0,048979 | 0 | 0,767243 | 0 | 0 | 0,757131 | 0,665799 | 3,296325584 |
| GSM2261680 | 0 | 0,205973 | 0 | 0,112604 | 0 | 0,681423 | 0 | 0 | 0,6564094 | 0,759332 | 2,138960044 |
| GSM2261689 | 0 | 0,119397 | 0 | 0,077659 | 0 | 0,802944 | 0 | 0 | 0,754566 | 0,66326 | 4,074698985 |
| GSM2261691 | 0 | 0,167516 | 0,010399 | 0,093608 | 0 | 0,728477 | 0 | 0 | 0,7434541 | 0,684038 | 2,682929916 |
| GSM2261708 | 0 | 0,188898 | 0,011434 | 0,105754 | 0 | 0,693914 | 0 | 0 | 0,6697864 | 0,748173 | 2,267052974 |
| GSM2261709 | 0 | 0,147189 | 0 | 0,112039 | 0 | 0,740772 | 0 | 0 | 0,7881395 | 0,640714 | 2,85760284 |
| GSM2261710 | 0 | 0,201008 | 0 | 0,181053 | 0 | 0,617938 | 0 | 0 | 0,6561942 | 0,762968 | 1,617378206 |
| GSM2261716 | 0 | 0,225324 | 0 | 0,190022 | 0 | 0,567097 | 0,01755656 | 0 | 0,6699962 | 0,757205 | 1,365360523 |
| GSM2261718 | 0 | 0,188391 | 0 | 0,138204 | 0 | 0,673405 | 0 | 0 | 0,5448257 | 0,837912 | 2,061895126 |
| GSM2261719 | 0 | 0,156635 | 0 | 0,112464 | 0 | 0,730901 | 0 | 0 | 0,6864891 | 0,731474 | 2,716109163 |
| GSM2261725 | 0 | 0,199659 | 0 | 0,168378 | 0 | 0,631963 | 0 | 0 | 0,7281128 | 0,709278 | 1,717116152 |
| GSM2261729 | 0 | 0,116519 | 0 | 0,102284 | 0 | 0,781196 | 0 | 0 | 0,6980232 | 0,717738 | 3,570308047 |
| GSM2261735 | 0 | 0,156026 | 0 | 0,085207 | 0 | 0,758767 | 0 | 0 | 0,6899381 | 0,726738 | 3,145375238 |
| GSM2261738 | 0 | 0,20748 | 0 | 0,146503 | 0 | 0,646018 | 0 | 0 | 0,6182895 | 0,788411 | 1,824999472 |
| GSM2261739 | 0 | 0,122053 | 0,02076 | 0,058868 | 0,005126 | 0,793193 | 0 | 0 | 0,7738673 | 0,64559 | 3,932906164 |
| GSM2261742 | 0 | 0,287882 | 0,046096 | 0,113355 | 0 | 0,538845 | 0,01382284 | 0 | 0,6714076 | 0,76239 | 1,204573687 |
| GSM2261746 | 0 | 0,176738 | 0 | 0,068908 | 0 | 0,754355 | 0 | 0 | 0,7239948 | 0,69799 | 3,070909595 |
| GSM2261747 | 0 | 0,204641 | 0 | 0,121797 | 0,004624 | 0,616644 | 0,05229479 | 0 | 0,7017316 | 0,729321 | 1,889009856 |
| GSM2261750 | 0 | 0,163288 | 0 | 0,086482 | 0,001736 | 0,748493 | 0 | 0 | 0,720737 | 0,701181 | 2,99672266 |
| GSM2261753 | 0 | 0,181179 | 0 | 0,097352 | 0 | 0,719926 | 0,00154348 | 0 | 0,7059125 | 0,716926 | 2,584723697 |
| GSM2261760 | 0 | 0,16529 | 0,012247 | 0,094495 | 0 | 0,727968 | 0 | 0 | 0,6865579 | 0,731971 | 2,676032059 |
| GSM2261762 | 0 | 0,158146 | 0 | 0,112077 | 0 | 0,729777 | 0 | 0 | 0,6734128 | 0,742204 | 2,700647818 |
| GSM2261765 | 0 | 0,140076 | 0 | 0,1169 | 0 | 0,743024 | 0 | 0 | 0,733566 | 0,68999 | 2,891421329 |
| GSM2261775 | 0 | 0,146415 | 0 | 0,130954 | 0 | 0,722632 | 0 | 0 | 0,6435341 | 0,765933 | 2,605314384 |
| GSM2261776 | 0 | 0,172596 | 0 | 0,144557 | 0 | 0,682847 | 0 | 0 | 0,6908382 | 0,7321 | 2,153053946 |
| GSM2261780 | 0 | 0,186705 | 0 | 0,068003 | 0 | 0,745292 | 0 | 0 | 0,7174046 | 0,704861 | 2,926063061 |
| GSM2261782 | 0 | 0,24893 | 0,005447 | 0,13453 | 0 | 0,611093 | 0 | 0 | 0,4662782 | 0,886134 | 1,57130584 |
| GSM2261791 | 0 | 0,114728 | 0,087899 | 0,126257 | 0 | 0,665747 | 0,00536789 | 0 | 0,5710784 | 0,8197 | 2,024256825 |
| GSM2261798 | 0 | 0,188973 | 0 | 0,061509 | 0,002385 | 0,747133 | 0 | 0 | 0,6995686 | 0,719939 | 2,982779981 |
| GSM2261800 | 0 | 0,197009 | 0,037746 | 0,140942 | 0 | 0,468515 | 0,15578794 | 0 | 0,4853692 | 0,873536 | 1,247053937 |
| GSM2261801 | 0 | 0,186019 | 0 | 0,113735 | 0 | 0,700246 | 0 | 0 | 0,6421006 | 0,768612 | 2,336068153 |
| GSM2261802 | 0 | 0,151409 | 0,004436 | 0,116248 | 0 | 0,727907 | 0 | 0 | 0,7054236 | 0,716003 | 2,675216488 |
| GSM2261805 | 0 | 0,198959 | 0 | 0,093941 | 0 | 0,7071 | 0 | 0 | 0,7175217 | 0,70924 | 2,414135333 |
| GSM2261807 | 0 | 0,195158 | 0 | 0,128153 | 0 | 0,669543 | 0,00714621 | 0 | 0,6797794 | 0,742184 | 2,07089579 |
| GSM2261812 | 0 | 0,226868 | 0,053414 | 0,072818 | 0 | 0,6469 | 0 | 0 | 0,7691784 | 0,677958 | 1,832060359 |
| GSM2261815 | 0 | 0,100854 | 0,044507 | 0,10161 | 0 | 0,753028 | 0 | 0 | 0,742107 | 0,680992 | 3,049048096 |
| GSM2261816 | 0 | 0,224956 | 0 | 0,139019 | 0 | 0,624843 | 0,01118232 | 0 | 0,5192649 | 0,854142 | 1,716718827 |

| | | | | | | | | | | | |
|---|---|---|---|---|---|---|---|---|---|---|---|
| GSM2261817 | 0 | 0,193726 | 0 | 0,153745 | 0 | 0,652529 | 0 | 0 | 0,624883 | 0,78322 | 1,877934934 |
| GSM2261821 | 0 | 0,209901 | 0 | 0,104526 | 0 | 0,685573 | 0 | 0 | 0,6649379 | 0,752769 | 2,180387917 |
| GSM2261823 | 0 | 0,108136 | 0 | 0,078405 | 0 | 0,759125 | 0,05433488 | 0 | 0,6912174 | 0,723384 | 4,06949819 |
| GSM2261824 | 0 | 0,185372 | 0 | 0,14195 | 0 | 0,672678 | 0 | 0 | 0,761966 | 0,676645 | 2,05509733 |
| GSM2261826 | 0 | 0,173539 | 0,000649 | 0,130322 | 0 | 0,69549 | 0 | 0 | 0,5774893 | 0,815546 | 2,283966087 |
| GSM2261828 | 0 | 0,19188 | 0 | 0,113514 | 0 | 0,692464 | 0,00214123 | 0 | 0,5859779 | 0,809788 | 2,267443468 |
| GSM2261830 | 0 | 0,228847 | 0 | 0,105976 | 0 | 0,665177 | 0 | 0 | 0,6520148 | 0,764129 | 1,986652337 |
| GSM2261832 | 0 | 0,201448 | 0 | 0,097029 | 0 | 0,701523 | 0 | 0 | 0,710528 | 0,715615 | 2,350341611 |
| GSM2261834 | 0 | 0,16015 | 0 | 0,109364 | 0 | 0,730486 | 0 | 0 | 0,6841324 | 0,733506 | 2,710381869 |
| GSM2261839 | 0 | 0,192769 | 0 | 0,110797 | 0 | 0,655187 | 0,0412472 | 0 | 0,547526 | 0,836293 | 2,15830096 |
| GSM2261842 | 0 | 0,173001 | 0 | 0,126845 | 0 | 0,700154 | 0 | 0 | 0,5938384 | 0,803884 | 2,33504985 |
| GSM2261844 | 0 | 0,228717 | 0 | 0,130643 | 0 | 0,640641 | 0 | 0 | 0,5313439 | 0,846669 | 1,782728933 |
| GSM2261851 | 0 | 0,079254 | 0,08208 | 0,097671 | 0 | 0,740995 | 0 | 0 | 0,6868696 | 0,72988 | 2,860923307 |
| GSM2261853 | 0 | 0,171899 | 0 | 0,065834 | 0 | 0,762267 | 0 | 0 | 0,7232117 | 0,69778 | 3,206407543 |
| GSM2261856 | 0 | 0,185987 | 0 | 0,119909 | 0 | 0,694103 | 0 | 0 | 0,7223113 | 0,706458 | 2,269076817 |
| GSM2261858 | 0 | 0,174577 | 0,006876 | 0,105242 | 0 | 0,713305 | 0 | 0 | 0,7111032 | 0,713328 | 2,488023011 |
| GSM2261859 | 0 | 0,232963 | 0 | 0,063881 | 0 | 0,703156 | 0 | 0 | 0,6777708 | 0,742076 | 2,368770618 |
| GSM2261860 | 0 | 0,202941 | 0 | 0,043731 | 0,005148 | 0,746352 | 0,00182804 | 0 | 0,7528315 | 0,673469 | 3,025686209 |
| GSM2261861 | 0 | 0,181267 | 0 | 0,152629 | 0 | 0,666104 | 0 | 0 | 0,6595063 | 0,757269 | 1,994947748 |
| GSM2261866 | 0 | 0,070747 | 0,027732 | 0,149695 | 0 | 0,751825 | 0 | 0 | 0,6519516 | 0,757564 | 3,029415945 |
| GSM2261867 | 0 | 0,261097 | 0 | 0,126076 | 0 | 0,612827 | 0 | 0 | 0,6957062 | 0,738036 | 1,582825455 |
| GSM2261875 | 0 | 0,129831 | 0,045689 | 0,10187 | 0 | 0,72261 | 0 | 0 | 0,6859647 | 0,732506 | 2,605036404 |
| GSM2261876 | 0 | 0,169683 | 0,000836 | 0,173601 | 0 | 0,627794 | 0,02808699 | 0 | 0,5667229 | 0,822968 | 1,82435113 |
| GSM2261877 | 0 | 0,198972 | 0 | 0,114993 | 0 | 0,686035 | 0 | 0 | 0,6827999 | 0,738862 | 2,185068552 |
| GSM2261888 | 0 | 0,138992 | 0 | 0,054747 | 0 | 0,806262 | 0 | 0 | 0,7706919 | 0,647065 | 4,161601346 |
| GSM2261889 | 0 | 0,143894 | 0,014381 | 0,071638 | 0 | 0,770088 | 0 | 0 | 0,7127898 | 0,705999 | 3,349483183 |
| GSM2261891 | 0 | 0,225589 | 0 | 0,137809 | 0 | 0,581511 | 0,05509086 | 0 | 0,5101323 | 0,859566 | 1,600201767 |
| GSM2261893 | 0 | 0,146965 | 0,00406 | 0,13984 | 0 | 0,709136 | 0 | 0 | 0,6208022 | 0,783645 | 2,438027162 |
| GSM2261894 | 0 | 0,1722 | 0 | 0,119826 | 0 | 0,707974 | 0 | 0 | 0,6070918 | 0,794179 | 2,424357129 |
| GSM2261907 | 0 | 0,12399 | 0,029303 | 0,101709 | 0 | 0,744998 | 0 | 0 | 0,7490567 | 0,676068 | 2,921532165 |
| GSM2261911 | 0 | 0,147075 | 0 | 0,100718 | 0 | 0,752206 | 0 | 0 | 0,7107457 | 0,709194 | 3,035616185 |
| GSM2261914 | 0 | 0,167555 | 0 | 0,123046 | 0 | 0,709399 | 0 | 0 | 0,684359 | 0,734993 | 2,441140515 |
| GSM2261915 | 0 | 0,14301 | 0,032078 | 0,147155 | 0 | 0,677757 | 0 | 0 | 0,5396205 | 0,841234 | 2,103245425 |
| GSM2261937 | 0 | 0,13465 | 0,025118 | 0,111557 | 0 | 0,728674 | 0 | 0 | 0,7127478 | 0,709715 | 2,685609807 |
| GSM2261941 | 0 | 0,155304 | 0,033195 | 0,079938 | 0 | 0,731563 | 0 | 0 | 0,7183237 | 0,705302 | 2,725265416 |
| GSM2261946 | 0 | 0,162993 | 0,00068 | 0,164319 | 0 | 0,672008 | 0 | 0 | 0,6429378 | 0,768668 | 2,048850834 |
| GSM2261949 | 0 | 0,218982 | 0,015686 | 0,090642 | 0 | 0,674689 | 0 | 0 | 0,5016539 | 0,866684 | 2,073981601 |
| GSM2261954 | 0 | 0,132083 | 0 | 0,158735 | 0 | 0,709182 | 0 | 0 | 0,6440198 | 0,765722 | 2,438576752 |
| GSM2261956 | 0 | 0,129075 | 0,127628 | 0,02952 | 0,000375 | 0,713402 | 0 | 0 | 0,718113 | 0,707553 | 2,492469967 |
| GSM2261960 | 0 | 0,08197 | 0,027086 | 0,167397 | 0 | 0,723547 | 0 | 0 | 0,5057173 | 0,86672 | 2,617251709 |
| GSM2261961 | 0 | 0,137591 | 0 | 0,104738 | 0 | 0,728676 | 0,02899597 | 0 | 0,7044439 | 0,715234 | 3,006978469 |
| GSM2261965 | 0 | 0,205791 | 0,039871 | 0,101432 | 0 | 0,652907 | 0 | 0 | 0,6105508 | 0,793961 | 1,881070717 |
| GSM2261968 | 0,004099 | 0,169548 | 0 | 0,124926 | 0 | 0,701427 | 0 | 0 | 0,6835988 | 0,73641 | 2,381968441 |
| GSM2261969 | 0 | 0,147394 | 0,0043 | 0,079341 | 0 | 0,768966 | 0 | 0 | 0,7311787 | 0,689554 | 3,328357909 |
| GSM2261970 | 0 | 0,255984 | 0,005494 | 0,115325 | 0 | 0,556762 | 0,06643399 | 0 | 0,5140558 | 0,857494 | 1,477590123 |
| GSM2261972 | 0 | 0,177467 | 0 | 0,201372 | 0 | 0,621161 | 0 | 0 | 0,5454493 | 0,836913 | 1,639644406 |
| GSM2261974 | 0 | 0,237115 | 0,032166 | 0,068144 | 0 | 0,662574 | 0 | 0 | 0,7185221 | 0,715294 | 1,963617037 |
| GSM2261979 | 0 | 0,14362 | 0,027744 | 0,10573 | 0 | 0,691333 | 0,03157318 | 0 | 0,713385 | 0,711836 | 2,494937238 |
| GSM2261982 | 0 | 0,195999 | 0 | 0,125305 | 0 | 0,678696 | 0 | 0 | 0,7346871 | 0,698653 | 2,112316445 |
| GSM2261983 | 0 | 0,158803 | 0,031672 | 0,085514 | 0 | 0,712669 | 0,01134311 | 0 | 0,6634404 | 0,751204 | 2,582240986 |
| GSM2261984 | 0 | 0,188233 | 0 | 0,177156 | 0 | 0,634612 | 0 | 0 | 0,5344278 | 0,844215 | 1,73681414 |
| GSM2261987 | 0 | 0,119325 | 0,016241 | 0,091283 | 0 | 0,773151 | 0 | 0 | 0,7415587 | 0,679193 | 3,408215243 |
| GSM2261993 | 0,004357 | 0,151375 | 0,046709 | 0,054348 | 0,003449 | 0,703468 | 0,03629455 | 0 | 0,7283684 | 0,698164 | 2,786773489 |
| GSM2261994 | 0 | 0,230897 | 0 | 0,071277 | 0 | 0,697827 | 0 | 0 | 0,7397365 | 0,692878 | 2,309358465 |
| GSM2261995 | 0 | 0,176784 | 0 | 0,081936 | 0 | 0,74128 | 0 | 0 | 0,6949044 | 0,724153 | 2,86518535 |
| GSM2261997 | 0 | 0,260942 | 0 | 0,163896 | 0 | 0,575161 | 0 | 0 | 0,5549027 | 0,832474 | 1,353835091 |
| GSM2261999 | 0 | 0,133805 | 0 | 0,112196 | 0 | 0,753999 | 0 | 0 | 0,7290365 | 0,692671 | 3,065017896 |
| GSM2262000 | 0 | 0,18693 | 0,00024 | 0,053589 | 0 | 0,759241 | 0 | 0 | 0,7489936 | 0,675099 | 3,153527618 |
| GSM2262005 | 0 | 0,213361 | 0 | 0,122805 | 0 | 0,663835 | 0 | 0 | 0,5951274 | 0,804057 | 1,974726282 |
| GSM2262006 | 0 | 0,205565 | 0 | 0,131772 | 0 | 0,658305 | 0,00435918 | 0 | 0,6897073 | 0,736055 | 1,95147897 |
| GSM2262008 | 0 | 0,234173 | 0,001787 | 0,180427 | 0 | 0,580523 | 0,00308858 | 0 | 0,556377 | 0,830966 | 1,394188648 |
| GSM2262009 | 0 | 0,112518 | 0 | 0,207332 | 0 | 0,68015 | 0 | 0 | 0,6835923 | 0,73567 | 2,126461425 |
| GSM2262010 | 0 | 0,230095 | 0,004977 | 0,123413 | 0 | 0,641516 | 0 | 0 | 0,5200665 | 0,853894 | 1,789521353 |
| GSM2262011 | 0,005287 | 0,193694 | 0 | 0,127315 | 0 | 0,673703 | 0 | 0 | 0,6763831 | 0,744832 | 2,098703358 |
| GSM2262012 | 0 | 0,230004 | 0 | 0,09561 | 0 | 0,674387 | 0 | 0 | 0,7257447 | 0,707491 | 2,071129047 |
| GSM2262019 | 0 | 0,12479 | 0,034003 | 0,163072 | 0 | 0,678135 | 0 | 0 | 0,7073556 | 0,718565 | 2,106891078 |
| GSM2262027 | 0 | 0,120715 | 0,011598 | 0,102953 | 0 | 0,764735 | 0 | 0 | 0,6917829 | 0,724305 | 3,250518057 |
| GSM2262029 | 0 | 0,196632 | 0 | 0,101301 | 0,003439 | 0,698628 | 0 | 0 | 0,6572141 | 0,757457 | 2,344912372 |
| GSM2262037 | 0 | 0,163587 | 0 | 0,100261 | 0 | 0,736152 | 0 | 0 | 0,7336602 | 0,691317 | 2,790054175 |
| GSM2262043 | 0 | 0,134659 | 0,043902 | 0,085415 | 0 | 0,736025 | 0 | 0 | 0,7586294 | 0,669232 | 2,788234021 |
| GSM2262045 | 0 | 0,248861 | 0 | 0,119268 | 0 | 0,63187 | 0 | 0 | 0,6768382 | 0,749333 | 1,716435011 |
| GSM2262047 | 0 | 0,159817 | 0 | 0,04577 | 0,001837 | 0,792576 | 0 | 0 | 0,7012707 | 0,71486 | 3,855178674 |
| GSM2262050 | 0 | 0,167239 | 0,056197 | 0,077371 | 0,003004 | 0,696188 | 0 | 0 | 0,6140872 | 0,789565 | 2,314400065 |
| GSM2262052 | 0,00032 | 0,121807 | 0,021757 | 0,090624 | 0 | 0,765492 | 0 | 0 | 0,6558812 | 0,754658 | 3,268705373 |
| GSM2262054 | 0 | 0,230444 | 0 | 0,116031 | 0 | 0,653525 | 0 | 0 | 0,5142247 | 0,857892 | 1,886212648 |
| GSM2262055 | 0 | 0,119133 | 0,027089 | 0,039909 | 0 | 0,813869 | 0 | 0 | 0,7615344 | 0,655312 | 4,372562529 |
| GSM2262057 | 0 | 0,219357 | 0 | 0,122012 | 0 | 0,658631 | 0 | 0 | 0,6577947 | 0,760116 | 1,929381943 |

| | | | | | | | | | | | |
|---|---|---|---|---|---|---|---|---|---|---|---|
| GSM2262062 | 0 | 0,096681 | 0 | 0,153743 | 0 | 0,696527 | 0,05304857 | 0 | 0,6038317 | 0,795828 | 2,781387911 |
| GSM2262066 | 0 | 0,226509 | 0 | 0,131075 | 0,000939 | 0,570549 | 0,07092908 | 0 | 0,5677706 | 0,823883 | 1,595567531 |
| GSM2262073 | 0 | 0,205424 | 0 | 0,120538 | 0 | 0,674039 | 0 | 0 | 0,5487578 | 0,835464 | 2,067847233 |
| GSM2262074 | 0 | 0,193136 | 0 | 0,177353 | 0 | 0,629511 | 0 | 0 | 0,5439876 | 0,838046 | 1,699136341 |
| GSM2262076 | 0 | 0,16022 | 0,019509 | 0,125529 | 0 | 0,694742 | 0 | 0 | 0,6328226 | 0,775458 | 2,27591375 |
| GSM2262081 | 0 | 0,199612 | 0 | 0,10778 | 0 | 0,692607 | 0 | 0 | 0,5765995 | 0,816494 | 2,253167352 |
| GSM2262084 | 0 | 0,145043 | 0 | 0,165105 | 0,010057 | 0,626124 | 0,05367097 | 0 | 0,5611021 | 0,826464 | 2,018789873 |
| GSM2262085 | 0 | 0,240241 | 0 | 0,14558 | 0 | 0,610644 | 0,00353472 | 0 | 0,4387128 | 0,902367 | 1,582714273 |
| GSM2262093 | 0 | 0,201963 | 0 | 0,096555 | 0 | 0,672356 | 0,02912571 | 0 | 0,6138755 | 0,790275 | 2,25231449 |
| GSM2262096 | 0 | 0,205437 | 0,03153 | 0,13794 | 0 | 0,625093 | 0 | 0 | 0,6629221 | 0,758827 | 1,667325055 |
| GSM2262099 | 0 | 0,156556 | 0 | 0,172261 | 0 | 0,671183 | 0 | 0 | 0,7304102 | 0,701129 | 2,041209813 |
| GSM2262100 | 0 | 0,140288 | 0 | 0,092735 | 0 | 0,766977 | 0 | 0 | 0,6930448 | 0,723303 | 3,291429165 |
| GSM2262106 | 0 | 0,166288 | 0 | 0,138704 | 0 | 0,695009 | 0 | 0 | 0,7531293 | 0,680012 | 2,278780782 |
| GSM2262111 | 0 | 0,230902 | 0 | 0,147989 | 0 | 0,621109 | 0 | 0 | 0,5624653 | 0,826703 | 1,639281464 |
| GSM2262115 | 0 | 0,135489 | 0,062649 | 0,126715 | 0 | 0,675147 | 0 | 0 | 0,6774548 | 0,743027 | 2,078318834 |
| GSM2262128 | 0 | 0,138198 | 0,002238 | 0,19484 | 0 | 0,664724 | 0 | 0 | 0,5995547 | 0,799633 | 1,982615287 |
| GSM2262130 | 0 | 0,193522 | 0 | 0,138384 | 0 | 0,668095 | 0 | 0 | 0,5341155 | 0,844985 | 2,012906436 |
| GSM2262132 | 0 | 0,109854 | 0,043329 | 0,097779 | 0 | 0,749039 | 0 | 0 | 0,6513184 | 0,758777 | 2,984676433 |
| GSM2262135 | 0 | 0,170766 | 0 | 0,176546 | 0 | 0,652688 | 0 | 0 | 0,489886 | 0,873154 | 1,879257803 |
| GSM2262139 | 0 | 0,180417 | 0 | 0,089421 | 0 | 0,730162 | 0 | 0 | 0,6417559 | 0,767662 | 2,705930282 |
| GSM2262142 | 0 | 0,264183 | 0 | 0,142861 | 0 | 0,548905 | 0,04405031 | 0 | 0,4999244 | 0,865657 | 1,348514029 |
| GSM2262145 | 0 | 0,219542 | 0 | 0,083194 | 0,004729 | 0,692536 | 0 | 0 | 0,6285738 | 0,779558 | 2,287593303 |
| GSM2262151 | 0 | 0,145468 | 0 | 0,081049 | 0 | 0,773483 | 0 | 0 | 0,6576026 | 0,753243 | 3,414680999 |
| GSM2262156 | 0 | 0,158417 | 0,001864 | 0,09113 | 0 | 0,74859 | 0 | 0 | 0,7392335 | 0,684807 | 2,977562221 |
| GSM2262165 | 0 | 0,158751 | 0,006322 | 0,122307 | 0 | 0,71262 | 0 | 0 | 0,7312037 | 0,696201 | 2,479714204 |
| GSM2262167 | 0 | 0,213761 | 0 | 0,109483 | 0 | 0,676756 | 0 | 0 | 0,6034528 | 0,798001 | 2,093642023 |
| GSM2262169 | 0,00294 | 0,33614 | 0 | 0,09493 | 0,005353 | 0,559139 | 0,00149729 | 0 | 0,5471074 | 0,839169 | 1,297096601 |
| GSM2262170 | 0 | 0,177357 | 0 | 0,140713 | 0 | 0,68193 | 0 | 0 | 0,6070441 | 0,794688 | 2,143958265 |
| GSM2262174 | 0 | 0,177256 | 0 | 0,10991 | 0 | 0,712834 | 0 | 0 | 0,662753 | 0,752008 | 2,482305007 |
| GSM2262178 | 0 | 0,236864 | 0 | 0,172667 | 0 | 0,574406 | 0,01606221 | 0 | 0,638105 | 0,778856 | 1,402593455 |
| GSM2262181 | 0 | 0,189438 | 0 | 0,057883 | 0 | 0,752678 | 0 | 0 | 0,6073107 | 0,794332 | 3,043316486 |
| GSM2262182 | 0 | 0,161459 | 0 | 0,139941 | 0 | 0,698601 | 0 | 0 | 0,621885 | 0,783263 | 2,317857354 |
| GSM2262188 | 0 | 0,251652 | 0 | 0,080716 | 0 | 0,635475 | 0,03215668 | 0 | 0,5537187 | 0,832796 | 1,911960146 |
| GSM2262189 | 0 | 0,223024 | 0 | 0,155236 | 0 | 0,62174 | 0 | 0 | 0,5365041 | 0,843142 | 1,643686363 |
| GSM2262190 | 0 | 0,093263 | 0,062904 | 0,149232 | 0 | 0,637018 | 0,0575827 | 0 | 0,6570659 | 0,758232 | 2,085856675 |
| GSM2262198 | 0 | 0,236814 | 0 | 0,17428 | 0 | 0,566673 | 0,02223289 | 0 | 0,5602972 | 0,828816 | 1,378448707 |
| GSM2262199 | 0 | 0,092289 | 0,001044 | 0,208533 | 0 | 0,698134 | 0 | 0 | 0,6940298 | 0,725549 | 2,312732839 |
| GSM2262202 | 0 | 0,16383 | 0 | 0,058256 | 0 | 0,777913 | 0 | 0 | 0,6848869 | 0,730159 | 3,502749446 |
| GSM2262206 | 0 | 0,189711 | 0 | 0,107178 | 0 | 0,703111 | 0 | 0 | 0,5766977 | 0,816367 | 2,368259263 |
| GSM2262207 | 0 | 0,16107 | 0 | 0,10178 | 0 | 0,73715 | 0 | 0 | 0,6350366 | 0,772413 | 2,804450774 |
| GSM2262217 | 0 | 0,142452 | 0,005568 | 0,115566 | 0 | 0,736413 | 0 | 0 | 0,6238621 | 0,780874 | 2,793818514 |
| GSM2262218 | 0 | 0,167407 | 0 | 0,141928 | 0 | 0,690665 | 0 | 0 | 0,6709808 | 0,746732 | 2,232740921 |
| GSM2262226 | 0 | 0,158998 | 0 | 0,182178 | 0 | 0,658824 | 0 | 0 | 0,5869588 | 0,808821 | 1,931035401 |
| GSM2262233 | 0 | 0,173665 | 0 | 0,061206 | 0 | 0,747852 | 0,01727672 | 0 | 0,7132285 | 0,707159 | 3,184101365 |
| GSM2262234 | 0 | 0,199921 | 0,004383 | 0,061753 | 0 | 0,733943 | 0 | 0 | 0,6488373 | 0,762354 | 2,758594722 |
| GSM2262236 | 0 | 0,204646 | 0 | 0,107403 | 0 | 0,687951 | 0 | 0 | 0,628414 | 0,77961 | 2,204627616 |
| GSM2262240 | 0 | 0,157204 | 0 | 0,092728 | 0 | 0,750068 | 0 | 0 | 0,7149796 | 0,705926 | 3,001096172 |
| GSM2262244 | 0 | 0,240333 | 0 | 0,095283 | 0 | 0,664384 | 0 | 0 | 0,5457269 | 0,837824 | 1,979595782 |
| GSM2262247 | 0 | 0,163253 | 0 | 0,107885 | 0,004423 | 0,72444 | 0 | 0 | 0,6394543 | 0,769293 | 2,671849874 |
| GSM2262248 | 0 | 0,205312 | 0,013374 | 0,09682 | 0 | 0,684494 | 0 | 0 | 0,5976597 | 0,801903 | 2,16950935 |
| GSM2262255 | 0 | 0,238827 | 0 | 0,129486 | 0 | 0,619571 | 0,01211631 | 0 | 0,5023066 | 0,864724 | 1,682186983 |
| GSM2262260 | 0 | 0,146955 | 0 | 0,057137 | 0 | 0,795908 | 0 | 0 | 0,6993803 | 0,71623 | 3,899739185 |
| GSM2262262 | 0 | 0,196145 | 0 | 0,091013 | 0 | 0,712842 | 0 | 0 | 0,612687 | 0,790369 | 2,482403027 |
| GSM2262269 | 0 | 0,175093 | 0 | 0,135361 | 0 | 0,689546 | 0 | 0 | 0,58793 | 0,808169 | 2,221090287 |
| GSM2262271 | 0 | 0,242473 | 0 | 0,132158 | 0 | 0,62537 | 0 | 0 | 0,6198505 | 0,789072 | 1,66929878 |
| GSM2262272 | 0 | 0,271165 | 0 | 0,122806 | 0 | 0,565204 | 0,04082502 | 0 | 0,498744 | 0,866602 | 1,434634379 |
| GSM2262273 | 0 | 0,199922 | 0 | 0,098559 | 0 | 0,701519 | 0 | 0 | 0,6009677 | 0,799138 | 2,350299204 |
| GSM2262277 | 0 | 0,100499 | 0,002396 | 0,15802 | 0 | 0,697412 | 0,04167353 | 0 | 0,7229883 | 0,700754 | 2,672951926 |
| GSM2262278 | 0 | 0,196644 | 0 | 0,140868 | 0 | 0,662488 | 0 | 0 | 0,5990817 | 0,80103 | 1,962859294 |
| GSM2262283 | 0 | 0,192559 | 0 | 0,11304 | 0 | 0,691605 | 0,00279631 | 0 | 0,625889 | 0,781065 | 2,263114326 |
| GSM2262284 | 0 | 0,248529 | 0 | 0,141925 | 0 | 0,609546 | 0 | 0 | 0,4761095 | 0,880243 | 1,561118924 |
| GSM2262285 | 0 | 0,229017 | 0 | 0,132078 | 0 | 0,638096 | 0,00080943 | 0 | 0,7018196 | 0,729872 | 1,767114141 |
| GSM2262286 | 0 | 0,176824 | 0 | 0,095593 | 0 | 0,727583 | 0 | 0 | 0,6686523 | 0,746515 | 2,67083948 |
| GSM2262287 | 0 | 0,261174 | 0 | 0,079744 | 0 | 0,659082 | 0 | 0 | 0,6023634 | 0,80012 | 1,933258106 |
| GSM2262288 | 0 | 0,14217 | 0 | 0,071158 | 0 | 0,786672 | 0 | 0 | 0,6996908 | 0,716361 | 3,687620179 |
| GSM2262289 | 0 | 0,143467 | 0 | 0,116048 | 0 | 0,728509 | 0,01197653 | 0 | 0,6900201 | 0,728091 | 2,807194963 |
| GSM2262295 | 0 | 0,242387 | 0 | 0,18499 | 0 | 0,48563 | 0,08699322 | 0 | 0,4281832 | 0,905146 | 1,136303197 |
| GSM2262298 | 0 | 0,231609 | 0 | 0,193496 | 0 | 0,553862 | 0,02103301 | 0 | 0,4298007 | 0,905622 | 1,302884559 |
| GSM2262302 | 0 | 0,290283 | 0 | 0,142628 | 0 | 0,497108 | 0,06998037 | 0 | 0,5142929 | 0,857867 | 1,148289879 |
| GSM2262306 | 0 | 0,194828 | 0 | 0,094363 | 0 | 0,710808 | 0 | 0 | 0,6251911 | 0,781085 | 2,457915122 |
| GSM2262309 | 0 | 0,180377 | 0 | 0,123331 | 0 | 0,690914 | 0,00537819 | 0 | 0,5864123 | 0,809329 | 2,274925785 |
| GSM2262310 | 0 | 0,179031 | 0 | 0,125664 | 0 | 0,695305 | 0 | 0 | 0,630366 | 0,777411 | 2,28197322 |
| **average** | **0,000137** | **0,177508** | **0,008109** | **0,116807** | **0,000344** | **0,689965** | **0,00713073** | | | | |

**SES_CIBERSORTx_GSE85217_SHH**

| Sample | Bcells | TCD4 | TCD8 | Tgd | NK | MoMaDC | granulocytes | P-value | Correlation | RMSE | | Tab |
|---|---|---|---|---|---|---|---|---|---|---|---|---|
| GSM2261538 | 0 | 0,861694 | 0,002827 | 0,380422 | 0 | 2,9847599 | 0 | 0 | 0,6265381 | 0,780385 | | 0,864521 |
| GSM2261543 | 0 | 1,425353 | 0 | 0,771555 | 0 | 6,1499223 | 0 | 0 | 0,7034297 | 0,717266 | | 1,425353 |
| GSM2261544 | 0 | 0,581533 | 0 | 0,47402 | 0 | 1,6640443 | 0,0968151 | 0 | 0,5703643 | 0,821555 | | 0,581533 |
| GSM2261547 | 0 | 1,007144 | 0 | 0,806094 | 0 | 4,9173509 | 0 | 0 | 0,6249725 | 0,780146 | | 1,007144 |
| GSM2261549 | 0 | 0,814156 | 0 | 0,935139 | 0 | 5,8851358 | 0 | 0 | 0,7189782 | 0,699516 | | 0,814156 |
| GSM2261550 | 0 | 1,312535 | 0 | 1,209569 | 0 | 7,098433 | 0 | 0 | 0,6567303 | 0,75483 | | 1,312535 |
| GSM2261552 | 0 | 0,741504 | 0 | 0,493011 | 0 | 2,5739848 | 0 | 0 | 0,6039521 | 0,797317 | | 0,741504 |
| GSM2261567 | 0 | 0,529225 | 0 | 0,58675 | 0 | 2,4655549 | 0,0529567 | 0 | 0,5510153 | 0,833665 | | 0,529225 |
| GSM2261568 | 0 | 0,625436 | 0,157232 | 0,547701 | 0 | 4,8566012 | 0 | 0 | 0,6969537 | 0,718482 | | 0,782668 |
| GSM2261569 | 0 | 0,487019 | 0 | 0,455545 | 0 | 2,1071181 | 0 | 0 | 0,619177 | 0,785402 | | 0,487019 |
| GSM2261570 | 0 | 1,365022 | 0,045875 | 0,928564 | 0 | 12,166434 | 0 | 0 | 0,7388039 | 0,675543 | | 1,410897 |
| GSM2261571 | 0 | 1,474163 | 0 | 0,753755 | 0 | 4,1360829 | 0 | 0 | 0,6126945 | 0,792715 | | 1,474163 |
| GSM2261574 | 0 | 0,931295 | 0 | 0,823649 | 0 | 5,0348643 | 0 | 0 | 0,7195002 | 0,702351 | | 0,931295 |
| GSM2261586 | 0 | 0,489249 | 0 | 0,388002 | 0 | 2,1436349 | 0 | 0 | 0,5840615 | 0,810819 | | 0,489249 |
| GSM2261597 | 0 | 1,123155 | 0 | 0,696387 | 0 | 5,2447279 | 0 | 0 | 0,6484882 | 0,761643 | | 1,123155 |
| GSM2261598 | 0,002197 | 1,09247 | 0 | 0,717777 | 0 | 3,4201632 | 0 | 0 | 0,4562617 | 0,894063 | | 1,09247 |
| GSM2261600 | 0 | 0,877487 | 0 | 0,820041 | 0 | 3,8476965 | 0 | 0 | 0,54886 | 0,835383 | | 0,877487 |
| GSM2261602 | 0 | 0,857456 | 0 | 0,414077 | 0 | 3,7183848 | 0 | 0 | 0,6613598 | 0,751422 | | 0,857456 |
| GSM2261603 | 0 | 1,098223 | 0,137947 | 0,365528 | 0 | 5,686713 | 0 | 0 | 0,7009871 | 0,71586 | | 1,23617 |
| GSM2261614 | 0 | 0,185541 | 0,003229 | 0,229361 | 0 | 0,7881388 | 0,0572467 | 0 | 0,503457 | 0,864213 | | 0,18877 |
| GSM2261615 | 0 | 0,744322 | 0,089925 | 0,336323 | 0,011578 | 4,105673 | 0 | 0 | 0,6499408 | 0,759605 | | 0,834247 |
| GSM2261617 | 0 | 0,620196 | 0 | 0,39754 | 0 | 2,3518152 | 0 | 0 | 0,6380871 | 0,771673 | | 0,620196 |
| GSM2261619 | 0,003152 | 1,233824 | 0 | 0,4319 | 0 | 2,1171661 | 0 | 0 | 0,4724567 | 0,882035 | | 1,233824 |
| GSM2261620 | 0 | 0,620888 | 0 | 0,773222 | 0 | 2,7968725 | 0 | 0 | 0,5213626 | 0,85329 | | 0,620888 |
| GSM2261622 | 0 | 1,249144 | 0,313891 | 0,370788 | 0 | 3,2531657 | 0 | 0 | 0,7075634 | 0,7283 | | 1,563035 |
| GSM2261629 | 0 | 0,58804 | 0 | 0,722679 | 0,004016 | 4,216946 | 0,9174013 | 0 | 0,6940832 | 0,723605 | | 0,58804 |
| GSM2261633 | 0 | 0,777833 | 0 | 0,29738 | 0 | 4,3101787 | 0 | 0 | 0,7726248 | 0,646009 | | 0,777833 |
| GSM2261634 | 0 | 1,132371 | 0 | 1,048836 | 0 | 4,2392134 | 0 | 0 | 0,6579396 | 0,758588 | | 1,132371 |
| GSM2261641 | 0 | 0,57042 | 0 | 0,327406 | 0 | 1,6226417 | 0,010983 | 0 | 0,6537476 | 0,764259 | | 0,57042 |
| GSM2261644 | 0 | 0,838675 | 0,010438 | 1,322997 | 0,078556 | 6,7230318 | 0 | 0 | 0,7377433 | 0,684149 | | 0,849113 |
| GSM2261647 | 0 | 0,884295 | 0,086824 | 0,542577 | 0,066477 | 6,3057012 | 0 | 0 | 0,7478752 | 0,670047 | | 0,971119 |
| GSM2261650 | 0 | 0,455459 | 0 | 0,437481 | 0 | 1,7754557 | 0 | 0 | 0,6768178 | 0,744034 | | 0,455459 |
| GSM2261655 | 0 | 0,614902 | 0,208633 | 0,525163 | 0 | 2,8660848 | 0 | 0 | 0,7089821 | 0,718333 | | 0,823536 |
| GSM2261657 | 0 | 0,412568 | 0 | 0,449333 | 0 | 1,5994373 | 0,0632097 | 0 | 0,6324306 | 0,777584 | | 0,412568 |
| GSM2261658 | 0,079293 | 0,656027 | 0,413035 | 0,809183 | 0 | 4,4862448 | 0 | 0 | 0,7598601 | 0,674082 | | 1,069062 |
| GSM2261661 | 0 | 0,713285 | 0 | 0,556098 | 0 | 2,4858273 | 0 | 0 | 0,5624336 | 0,825973 | | 0,713285 |
| GSM2261673 | 0 | 0,554765 | 0,031345 | 0,571209 | 0 | 2,9951182 | 0 | 0 | 0,6910543 | 0,727982 | | 0,58611 |
| GSM2261674 | 0 | 0,579495 | 0,412759 | 0,264448 | 0 | 4,1424984 | 0 | 0 | 0,757131 | 0,665799 | | 0,992254 |
| GSM2261680 | 0 | 0,423434 | 0 | 0,231488 | 0 | 1,4008517 | 0 | 0 | 0,6564094 | 0,759332 | | 0,423434 |
| GSM2261689 | 0 | 1,143229 | 0 | 0,743583 | 0 | 7,6881904 | 0 | 0 | 0,754566 | 0,66326 | | 1,143229 |
| GSM2261691 | 0 | 1,026867 | 0,063743 | 0,573812 | 0 | 4,4655269 | 0 | 0 | 0,7434541 | 0,684038 | | 1,09061 |
| GSM2261708 | 0 | 0,91692 | 0,055502 | 0,513336 | 0 | 3,3682917 | 0 | 0 | 0,6697864 | 0,748173 | | 0,972422 |
| GSM2261709 | 0 | 1,909561 | 0 | 1,453543 | 0 | 9,6104178 | 0 | 0 | 0,7881395 | 0,640714 | | 1,909561 |
| GSM2261710 | 0 | 0,779692 | 0 | 0,702288 | 0 | 2,3969223 | 0 | 0 | 0,6561942 | 0,762968 | | 0,779692 |
| GSM2261716 | 0 | 0,843942 | 0 | 0,71172 | 0 | 2,1240388 | 0,0657574 | 0 | 0,6699962 | 0,757205 | | 0,843942 |
| GSM2261718 | 0 | 0,37179 | 0 | 0,272745 | 0 | 1,3289628 | 0 | 0 | 0,5448257 | 0,837912 | | 0,37179 |
| GSM2261719 | 0 | 0,444225 | 0 | 0,318953 | 0 | 2,0728733 | 0 | 0 | 0,6864891 | 0,731474 | | 0,444225 |
| GSM2261725 | 0 | 0,771665 | 0 | 0,650765 | 0 | 2,4424774 | 0 | 0 | 0,7281128 | 0,709278 | | 0,771665 |
| GSM2261729 | 0 | 0,755171 | 0 | 0,662914 | 0 | 5,0630002 | 0 | 0 | 0,6980232 | 0,717738 | | 0,755171 |
| GSM2261735 | 0 | 0,800924 | 0 | 0,437391 | 0 | 3,8949654 | 0 | 0 | 0,6899381 | 0,726738 | | 0,800924 |
| GSM2261738 | 0 | 0,767277 | 0 | 0,541778 | 0 | 2,3890252 | 0 | 0 | 0,6182895 | 0,788411 | | 0,767277 |
| GSM2261739 | 0 | 1,251138 | 0,212812 | 0,603446 | 0,052549 | 8,1308735 | 0 | 0 | 0,7738673 | 0,64559 | | 1,46395 |
| GSM2261742 | 0 | 1,59738 | 0,255772 | 0,628978 | 0 | 2,9899089 | 0,0766993 | 0 | 0,6714076 | 0,76239 | | 1,853152 |
| GSM2261746 | 0 | 0,703699 | 0 | 0,274362 | 0 | 3,0035357 | 0 | 0 | 0,7239948 | 0,69799 | | 0,703699 |
| GSM2261747 | 0 | 0,45312 | 0 | 0,269686 | 0,010238 | 1,3653866 | 0,1157923 | 0 | 0,7017316 | 0,729321 | | 0,45312 |
| GSM2261750 | 0 | 0,814655 | 0 | 0,431466 | 0,008663 | 3,734281 | 0 | 0 | 0,720737 | 0,701181 | | 0,814655 |
| GSM2261753 | 0 | 0,930271 | 0 | 0,499857 | 0 | 3,696485 | 0,007925 | 0 | 0,7059125 | 0,716926 | | 0,930271 |
| GSM2261760 | 0 | 0,721081 | 0,053428 | 0,412237 | 0 | 3,1757708 | 0 | 0 | 0,6865579 | 0,731971 | | 0,774509 |
| GSM2261762 | 0 | 0,616858 | 0 | 0,437163 | 0 | 2,8465392 | 0 | 0 | 0,6734128 | 0,742204 | | 0,616858 |
| GSM2261765 | 0 | 0,824705 | 0 | 0,688253 | 0 | 4,3745997 | 0 | 0 | 0,733566 | 0,68999 | | 0,824705 |
| GSM2261775 | 0 | 0,70249 | 0 | 0,62831 | 0 | 3,467153 | 0 | 0 | 0,6435341 | 0,765933 | | 0,70249 |
| GSM2261776 | 0 | 0,698272 | 0 | 0,584836 | 0 | 2,7626024 | 0 | 0 | 0,6908382 | 0,7321 | | 0,698272 |
| GSM2261780 | 0 | 0,871013 | 0 | 0,317246 | 0 | 3,4769205 | 0 | 0 | 0,7174046 | 0,704861 | | 0,871013 |
| GSM2261782 | 0 | 0,984831 | 0,02155 | 0,532234 | 0 | 2,4176344 | 0 | 0 | 0,4662782 | 0,886134 | | 1,00638 |
| GSM2261791 | 0 | 0,332607 | 0,254828 | 0,366031 | 0 | 1,930062 | 0,015562 | 0 | 0,5710784 | 0,8197 | | 0,587436 |
| GSM2261798 | 0 | 0,747472 | 0 | 0,243295 | 0,009435 | 2,9552396 | 0 | 0 | 0,6995686 | 0,719939 | | 0,747472 |
| GSM2261800 | 0 | 0,223193 | 0,042763 | 0,159675 | 0 | 0,5307846 | 0,1764936 | 0 | 0,4853692 | 0,873536 | | 0,265956 |
| GSM2261801 | 0 | 0,492807 | 0 | 0,301308 | 0 | 1,8551069 | 0 | 0 | 0,6421006 | 0,768612 | | 0,492807 |
| GSM2261802 | 0 | 0,63452 | 0,01859 | 0,487166 | 0 | 3,0504848 | 0 | 0 | 0,7054236 | 0,716003 | | 0,65311 |
| GSM2261805 | 0 | 0,837211 | 0 | 0,395299 | 0 | 2,9754459 | 0 | 0 | 0,7175217 | 0,70924 | | 0,837211 |
| GSM2261807 | 0 | 0,692177 | 0 | 0,454527 | 0 | 2,3747049 | 0,0253458 | 0 | 0,6797794 | 0,742184 | | 0,692177 |
| GSM2261812 | 0 | 2,549381 | 0,600228 | 0,818277 | 0 | 7,2694061 | 0 | 0 | 0,7691784 | 0,677958 | | 3,149609 |
| GSM2261815 | 0 | 0,646208 | 0,285173 | 0,651048 | 0 | 4,8249049 | 0 | 0 | 0,742107 | 0,680992 | | 0,931382 |
| GSM2261816 | 0 | 0,403086 | 0 | 0,2491 | 0 | 1,1196207 | 0,020037 | 0 | 0,5192649 | 0,854142 | | 0,403086 |
| GSM2261817 | 0 | 0,729845 | 0 | 0,57922 | 0 | 2,4583399 | 0 | 0 | 0,624883 | 0,78322 | | 0,729845 |
| GSM2261821 | 0 | 0,677549 | 0 | 0,337405 | 0 | 2,2129939 | 0 | 0 | 0,6649379 | 0,752769 | | 0,677549 |
| GSM2261823 | 0 | 0,956969 | 0 | 0,693857 | 0 | 6,718032 | 0,4808477 | 0 | 0,6912174 | 0,723384 | | 0,956969 |

| | | | | | | | | | | | | |
|---|---|---|---|---|---|---|---|---|---|---|---|---|
| GSM2261824 | 0 | 1,015901 | 0 | 0,77793 | 0 | 3,6864987 | 0 | 0 | 0,761966 | 0,676645 | | 1,015901 |
| GSM2261826 | 0 | 0,566839 | 0,002119 | 0,425676 | 0 | 2,2717088 | 0 | 0 | 0,5774893 | 0,815546 | | 0,568958 |
| GSM2261828 | 0 | 0,364802 | 0 | 0,215812 | 0 | 1,3165103 | 0,0040709 | 0 | 0,5859779 | 0,809788 | | 0,364802 |
| GSM2261830 | 0 | 0,918048 | 0 | 0,425135 | 0 | 2,6684389 | 0 | 0 | 0,6520148 | 0,764129 | | 0,918048 |
| GSM2261832 | 0 | 0,888805 | 0 | 0,4281 | 0 | 3,0951768 | 0 | 0 | 0,710528 | 0,715615 | | 0,888805 |
| GSM2261834 | 0 | 0,718659 | 0 | 0,490758 | 0 | 3,2779842 | 0 | 0 | 0,6841324 | 0,733506 | | 0,718659 |
| GSM2261839 | 0 | 0,558274 | 0 | 0,320875 | 0 | 1,8974688 | 0,1194549 | 0 | 0,547526 | 0,836293 | | 0,558274 |
| GSM2261842 | 0 | 0,536876 | 0 | 0,39364 | 0 | 2,1727993 | 0 | 0 | 0,5938384 | 0,803884 | | 0,536876 |
| GSM2261844 | 0 | 0,744395 | 0 | 0,425197 | 0 | 2,0850657 | 0 | 0 | 0,5313439 | 0,846669 | | 0,744395 |
| GSM2261851 | 0 | 0,434829 | 0,450333 | 0,535872 | 0 | 4,0654677 | 0 | 0 | 0,6868696 | 0,72988 | | 0,885162 |
| GSM2261853 | 0 | 0,662134 | 0 | 0,253584 | 0 | 2,9361678 | 0 | 0 | 0,7232117 | 0,69778 | | 0,662134 |
| GSM2261856 | 0 | 0,533398 | 0 | 0,343892 | 0 | 1,9906392 | 0 | 0 | 0,7223113 | 0,706458 | | 0,533398 |
| GSM2261858 | 0 | 1,091132 | 0,042975 | 0,657777 | 0 | 4,458248 | 0 | 0 | 0,7111032 | 0,713328 | | 1,134106 |
| GSM2261859 | 0 | 1,270456 | 0 | 0,348373 | 0 | 3,8346344 | 0 | 0 | 0,6777708 | 0,742076 | | 1,270456 |
| GSM2261860 | 0 | 1,177498 | 0 | 0,253737 | 0,029868 | 4,3304686 | 0,0106066 | 0 | 0,7528315 | 0,673469 | | 1,177498 |
| GSM2261861 | 0 | 0,695283 | 0 | 0,585434 | 0 | 2,5549647 | 0 | 0 | 0,6595063 | 0,757269 | | 0,695283 |
| GSM2261866 | 0 | 0,149141 | 0,058462 | 0,315571 | 0 | 1,5849132 | 0 | 0 | 0,6519516 | 0,757564 | | 0,207604 |
| GSM2261867 | 0 | 0,75262 | 0 | 0,363417 | 0 | 1,7664929 | 0 | 0 | 0,6957062 | 0,738036 | | 0,75262 |
| GSM2261875 | 0 | 0,58689 | 0,206533 | 0,460493 | 0 | 3,2664953 | 0 | 0 | 0,6859647 | 0,732506 | | 0,793422 |
| GSM2261876 | 0 | 0,281548 | 0,001387 | 0,288049 | 0 | 1,0416751 | 0,0466037 | 0 | 0,5667229 | 0,822968 | | 0,282935 |
| GSM2261877 | 0 | 0,692088 | 0 | 0,39998 | 0 | 2,3862433 | 0 | 0 | 0,6827999 | 0,738862 | | 0,692088 |
| GSM2261888 | 0 | 1,10245 | 0 | 0,434237 | 0 | 6,3950812 | 0 | 0 | 0,7706919 | 0,647065 | | 1,10245 |
| GSM2261889 | 0 | 0,373202 | 0,037298 | 0,1858 | 0 | 1,9972967 | 0 | 0 | 0,7127898 | 0,705999 | | 0,4105 |
| GSM2261891 | 0 | 0,258318 | 0 | 0,157803 | 0 | 0,6658772 | 0,0630835 | 0 | 0,5101323 | 0,859566 | | 0,258318 |
| GSM2261893 | 0 | 0,540082 | 0,014921 | 0,513898 | 0 | 2,6060083 | 0 | 0 | 0,6208022 | 0,783645 | | 0,555003 |
| GSM2261894 | 0 | 0,552193 | 0 | 0,384245 | 0 | 2,2702602 | 0 | 0 | 0,6070918 | 0,794179 | | 0,552193 |
| GSM2261907 | 0 | 0,568067 | 0,134255 | 0,465985 | 0 | 3,4132478 | 0 | 0 | 0,7490567 | 0,676068 | | 0,702322 |
| GSM2261911 | 0 | 1,038452 | 0 | 0,711141 | 0 | 5,3110958 | 0 | 0 | 0,7107457 | 0,709194 | | 1,038452 |
| GSM2261914 | 0 | 0,863827 | 0 | 0,634359 | 0 | 3,6572829 | 0 | 0 | 0,684359 | 0,734993 | | 0,863827 |
| GSM2261915 | 0 | 0,561574 | 0,125964 | 0,577851 | 0 | 2,6614251 | 0 | 0 | 0,5396205 | 0,841234 | | 0,687538 |
| GSM2261937 | 0 | 0,61864 | 0,115402 | 0,512539 | 0 | 3,3478293 | 0 | 0 | 0,7127478 | 0,709715 | | 0,734042 |
| GSM2261941 | 0 | 1,092581 | 0,233534 | 0,562375 | 0 | 5,1466348 | 0 | 0 | 0,7183237 | 0,705302 | | 1,326114 |
| GSM2261946 | 0 | 0,731192 | 0,00305 | 0,737139 | 0 | 3,0146406 | 0 | 0 | 0,6429378 | 0,768668 | | 0,734242 |
| GSM2261949 | 0 | 0,721489 | 0,051683 | 0,298643 | 0 | 2,2229235 | 0 | 0 | 0,5016539 | 0,866684 | | 0,773172 |
| GSM2261954 | 0 | 0,440226 | 0 | 0,529055 | 0 | 2,363667 | 0 | 0 | 0,6440198 | 0,765722 | | 0,440226 |
| GSM2261956 | 0 | 1,101368 | 1,089014 | 0,251884 | 0,003201 | 6,0872746 | 0 | 0 | 0,718113 | 0,707553 | | 2,190382 |
| GSM2261960 | 0 | 0,447404 | 0,14784 | 0,913676 | 0 | 3,9492214 | 0 | 0 | 0,5057173 | 0,86672 | | 0,595244 |
| GSM2261961 | 0 | 0,436697 | 0 | 0,332425 | 0 | 2,312732 | 0,0920298 | 0 | 0,7044439 | 0,715234 | | 0,436697 |
| GSM2261965 | 0 | 0,458091 | 0,088752 | 0,225788 | 0 | 1,4533754 | 0 | 0 | 0,6105508 | 0,793961 | | 0,546844 |
| GSM2261968 | 0,016887 | 0,698541 | 0 | 0,514699 | 0 | 2,8898985 | 0 | 0 | 0,6835988 | 0,73641 | | 0,698541 |
| GSM2261969 | 0 | 0,365316 | 0,010657 | 0,196647 | 0 | 1,905886 | 0 | 0 | 0,7311787 | 0,689554 | | 0,375974 |
| GSM2261970 | 0 | 0,373956 | 0,008026 | 0,168473 | 0 | 0,8133491 | 0,0970505 | 0 | 0,5140558 | 0,857494 | | 0,381983 |
| GSM2261972 | 0 | 0,563148 | 0 | 0,639003 | 0 | 1,9710994 | 0 | 0 | 0,5454493 | 0,836913 | | 0,563148 |
| GSM2261974 | 0 | 1,760579 | 0,238835 | 0,505972 | 0 | 4,91962 | 0 | 0 | 0,7185221 | 0,715294 | | 1,999414 |
| GSM2261979 | 0 | 0,805105 | 0,155528 | 0,592704 | 0 | 3,8754802 | 0,1769933 | 0 | 0,713385 | 0,711836 | | 0,960634 |
| GSM2261982 | 0 | 2,241284 | 0 | 1,432884 | 0 | 7,7610047 | 0 | 0 | 0,7346871 | 0,698653 | | 2,241284 |
| GSM2261983 | 0 | 0,478135 | 0,09536 | 0,257472 | 0 | 2,1457576 | 0,0341527 | 0 | 0,6634404 | 0,751204 | | 0,573495 |
| GSM2261984 | 0 | 0,363784 | 0 | 0,342376 | 0 | 1,2264673 | 0 | 0 | 0,5344278 | 0,844215 | | 0,363784 |
| GSM2261987 | 0 | 1,032397 | 0,14052 | 0,789776 | 0 | 6,6892767 | 0 | 0 | 0,7415587 | 0,679193 | | 1,172916 |
| GSM2261993 | 0,018362 | 0,637958 | 0,196849 | 0,229044 | 0,014535 | 2,9647116 | 0,1529605 | 0 | 0,7283684 | 0,698164 | | 0,834807 |
| GSM2261994 | 0 | 1,451397 | 0 | 0,44804 | 0 | 4,3864813 | 0 | 0 | 0,7397365 | 0,692878 | | 1,451397 |
| GSM2261995 | 0 | 0,638324 | 0 | 0,295852 | 0 | 2,6765864 | 0 | 0 | 0,6949044 | 0,724153 | | 0,638324 |
| GSM2261997 | 0 | 0,635787 | 0 | 0,399334 | 0 | 1,4013838 | 0 | 0 | 0,5549027 | 0,832474 | | 0,635787 |
| GSM2261999 | 0 | 0,661001 | 0 | 0,554254 | 0 | 3,7247782 | 0 | 0 | 0,7290365 | 0,692671 | | 0,661001 |
| GSM2262000 | 0 | 0,986371 | 0,001266 | 0,282772 | 0 | 4,0062675 | 0 | 0 | 0,7489936 | 0,675099 | | 0,987636 |
| GSM2262005 | 0 | 0,395026 | 0 | 0,227366 | 0 | 1,2290545 | 0 | 0 | 0,5951274 | 0,804057 | | 0,395026 |
| GSM2262006 | 0 | 0,561802 | 0 | 0,360128 | 0 | 1,7991255 | 0,0119135 | 0 | 0,6897073 | 0,736055 | | 0,561802 |
| GSM2262008 | 0 | 0,902049 | 0,006886 | 0,695017 | 0 | 2,2362113 | 0,0118974 | 0 | 0,556377 | 0,830966 | | 0,908934 |
| GSM2262009 | 0 | 0,305972 | 0 | 0,563799 | 0 | 1,8495348 | 0 | 0 | 0,6835923 | 0,73567 | | 0,305972 |
| GSM2262010 | 0 | 0,687087 | 0,014861 | 0,368523 | 0 | 1,9156307 | 0 | 0 | 0,5200665 | 0,853894 | | 0,701948 |
| GSM2262011 | 0,017595 | 0,644596 | 0 | 0,423692 | 0 | 2,2420189 | 0 | 0 | 0,6763831 | 0,744832 | | 0,644596 |
| GSM2262012 | 0 | 0,966867 | 0 | 0,401915 | 0 | 2,8349249 | 0 | 0 | 0,7257447 | 0,707491 | | 0,966867 |
| GSM2262019 | 0 | 0,414417 | 0,112922 | 0,54155 | 0 | 2,2520329 | 0 | 0 | 0,7073556 | 0,718565 | | 0,527339 |
| GSM2262027 | 0 | 1,272803 | 0,122283 | 1,085521 | 0 | 8,063259 | 0 | 0 | 0,6917829 | 0,724305 | | 1,395086 |
| GSM2262029 | 0 | 0,610671 | 0 | 0,314607 | 0,01068 | 2,1696956 | 0 | 0 | 0,6572141 | 0,757457 | | 0,610671 |
| GSM2262037 | 0 | 1,074947 | 0 | 0,658823 | 0 | 4,8373129 | 0 | 0 | 0,7336602 | 0,691317 | | 1,074947 |
| GSM2262043 | 0 | 0,644092 | 0,209989 | 0,408551 | 0 | 3,5205146 | 0 | 0 | 0,7586294 | 0,669232 | | 0,854081 |
| GSM2262045 | 0 | 0,716274 | 0 | 0,343279 | 0 | 1,818654 | 0 | 0 | 0,6768382 | 0,749333 | | 0,716274 |
| GSM2262047 | 0 | 0,977394 | 0 | 0,279918 | 0,011235 | 4,8471634 | 0 | 0 | 0,7012707 | 0,71486 | | 0,977394 |
| GSM2262050 | 0 | 0,5627 | 0,189081 | 0,260327 | 0,010109 | 2,3424224 | 0 | 0 | 0,6140872 | 0,789565 | | 0,751781 |
| GSM2262052 | 0,001131 | 0,430524 | 0,076899 | 0,320309 | 0 | 2,7056096 | 0 | 0 | 0,6558812 | 0,754658 | | 0,507423 |
| GSM2262054 | 0 | 0,743462 | 0 | 0,374342 | 0 | 2,1084157 | 0 | 0 | 0,5142247 | 0,857892 | | 0,743462 |
| GSM2262055 | 0 | 0,961622 | 0,218654 | 0,322139 | 0 | 6,5694017 | 0 | 0 | 0,7615344 | 0,655312 | | 1,180276 |
| GSM2262057 | 0 | 0,878675 | 0 | 0,488743 | 0 | 2,6382724 | 0 | 0 | 0,6577947 | 0,760116 | | 0,878675 |
| GSM2262062 | 0 | 0,613506 | 0 | 0,975599 | 0 | 4,4199181 | 0,3366277 | 0 | 0,6038317 | 0,795828 | | 0,613506 |
| GSM2262066 | 0 | 0,334687 | 0 | 0,193675 | 0,001387 | 0,8430376 | 0,1048042 | 0 | 0,5677706 | 0,823883 | | 0,334687 |
| GSM2262073 | 0 | 0,83194 | 0 | 0,488163 | 0 | 2,7297715 | 0 | 0 | 0,5487578 | 0,835464 | | 0,83194 |
| GSM2262074 | 0 | 0,854326 | 0 | 0,784507 | 0 | 2,7846014 | 0 | 0 | 0,5439876 | 0,838046 | | 0,854326 |
| GSM2262076 | 0 | 0,501019 | 0,061007 | 0,392538 | 0 | 2,1725066 | 0 | 0 | 0,6328226 | 0,775458 | | 0,562026 |
| GSM2262081 | 0 | 0,551864 | 0 | 0,297978 | 0 | 1,9148363 | 0 | 0 | 0,5765995 | 0,816494 | | 0,551864 |

| | | | | | | | | | | | | |
|---|---|---|---|---|---|---|---|---|---|---|---|---|
| GSM2262084 | 0 | 0,835655 | 0 | 0,95124 | 0,057942 | 3,6073663 | 0,3092212 | 0 | 0,5611021 | 0,826464 | | 0,835655 |
| GSM2262085 | 0 | 0,77391 | 0 | 0,468972 | 0 | 1,9671266 | 0,0113867 | 0 | 0,4387128 | 0,902367 | | 0,77391 |
| GSM2262093 | 0 | 0,863063 | 0 | 0,412613 | 0 | 2,8732233 | 0,1244648 | 0 | 0,6138755 | 0,790275 | | 0,863063 |
| GSM2262096 | 0 | 0,774213 | 0,118826 | 0,519845 | 0 | 2,3557372 | 0 | 0 | 0,6629221 | 0,758827 | | 0,893039 |
| GSM2262099 | 0 | 1,016506 | 0 | 1,118475 | 0 | 4,3579442 | 0 | 0 | 0,7304102 | 0,701129 | | 1,016506 |
| GSM2262100 | 0 | 0,799583 | 0 | 0,528552 | 0 | 4,3714621 | 0 | 0 | 0,6930448 | 0,723303 | | 0,799583 |
| GSM2262106 | 0 | 1,140013 | 0 | 0,950908 | 0 | 4,7647501 | 0 | 0 | 0,7531293 | 0,680012 | | 1,140013 |
| GSM2262111 | 0 | 0,778743 | 0 | 0,499111 | 0 | 2,0947625 | 0 | 0 | 0,5624653 | 0,826703 | | 0,778743 |
| GSM2262115 | 0 | 0,772713 | 0,357293 | 0,722669 | 0 | 3,8504495 | 0 | 0 | 0,6774548 | 0,743027 | | 1,130006 |
| GSM2262128 | 0 | 0,923555 | 0,014958 | 1,302083 | 0 | 4,442241 | 0 | 0 | 0,5995547 | 0,799633 | | 0,938513 |
| GSM2262130 | 0 | 0,682958 | 0 | 0,488369 | 0 | 2,3577717 | 0 | 0 | 0,5341155 | 0,844985 | | 0,682958 |
| GSM2262132 | 0 | 0,841834 | 0,33204 | 0,749298 | 0 | 5,7400483 | 0 | 0 | 0,6513184 | 0,758777 | | 1,173874 |
| GSM2262135 | 0 | 0,44612 | 0 | 0,461222 | 0 | 1,7051296 | 0 | 0 | 0,489886 | 0,873154 | | 0,44612 |
| GSM2262139 | 0 | 1,275841 | 0 | 0,632351 | 0 | 5,163434 | 0 | 0 | 0,6417559 | 0,767662 | | 1,275841 |
| GSM2262142 | 0 | 0,41815 | 0 | 0,226121 | 0 | 0,8688078 | 0,0697229 | 0 | 0,4999244 | 0,865657 | | 0,41815 |
| GSM2262145 | 0 | 1,137608 | 0 | 0,43109 | 0,024503 | 3,5885424 | 0 | 0 | 0,6285738 | 0,779558 | | 1,137608 |
| GSM2262151 | 0 | 0,617811 | 0 | 0,344219 | 0 | 3,285025 | 0 | 0 | 0,6576026 | 0,753243 | | 0,617811 |
| GSM2262156 | 0 | 1,289078 | 0,015169 | 0,741546 | 0 | 6,091474 | 0 | 0 | 0,7392335 | 0,684807 | | 1,304246 |
| GSM2262165 | 0 | 0,796409 | 0,031716 | 0,613576 | 0 | 3,5750063 | 0 | 0 | 0,7312037 | 0,696201 | | 0,828124 |
| GSM2262167 | 0 | 0,850267 | 0 | 0,435484 | 0 | 2,6919031 | 0 | 0 | 0,6034528 | 0,798001 | | 0,850267 |
| GSM2262169 | 0,018428 | 2,106704 | 0 | 0,594961 | 0,03355 | 3,5043212 | 0,009384 | 0 | 0,5471074 | 0,839169 | | 2,106704 |
| GSM2262170 | 0 | 0,635236 | 0 | 0,503991 | 0 | 2,4424532 | 0 | 0 | 0,6070441 | 0,794688 | | 0,635236 |
| GSM2262174 | 0 | 0,778862 | 0 | 0,482947 | 0 | 3,1321965 | 0 | 0 | 0,662753 | 0,752008 | | 0,778862 |
| GSM2262178 | 0 | 1,010754 | 0 | 0,736809 | 0 | 2,4511211 | 0,0685411 | 0 | 0,638105 | 0,778856 | | 1,010754 |
| GSM2262181 | 0 | 1,265656 | 0 | 0,386725 | 0 | 5,0287183 | 0 | 0 | 0,6073107 | 0,794332 | | 1,265656 |
| GSM2262182 | 0 | 0,592967 | 0 | 0,513942 | 0 | 2,5656573 | 0 | 0 | 0,621885 | 0,783263 | | 0,592967 |
| GSM2262188 | 0 | 0,344664 | 0 | 0,11055 | 0 | 0,8703505 | 0,044042 | 0 | 0,5537187 | 0,832796 | | 0,344664 |
| GSM2262189 | 0 | 1,12128 | 0 | 0,78047 | 0 | 3,1258807 | 0 | 0 | 0,536504 | 0,843142 | | 1,12128 |
| GSM2262190 | 0 | 0,652982 | 0,440423 | 1,044848 | 0 | 4,4600888 | 0,4031657 | 0 | 0,6570659 | 0,758232 | | 1,093405 |
| GSM2262198 | 0 | 0,305534 | 0 | 0,224854 | 0 | 0,7311121 | 0,0286845 | 0 | 0,5602972 | 0,828816 | | 0,305534 |
| GSM2262199 | 0 | 0,638148 | 0,007217 | 1,441937 | 0 | 4,827374 | 0 | 0 | 0,6940298 | 0,725549 | | 0,645366 |
| GSM2262202 | 0 | 1,264416 | 0 | 0,44961 | 0 | 6,0038065 | 0 | 0 | 0,6848869 | 0,730159 | | 1,264416 |
| GSM2262206 | 0 | 0,73903 | 0 | 0,417516 | 0 | 2,7390023 | 0 | 0 | 0,5766977 | 0,816367 | | 0,73903 |
| GSM2262207 | 0 | 0,931751 | 0 | 0,588776 | 0 | 4,2642429 | 0 | 0 | 0,6350366 | 0,772413 | | 0,931751 |
| GSM2262217 | 0 | 0,379231 | 0,014824 | 0,307656 | 0 | 1,9604528 | 0 | 0 | 0,6238621 | 0,780874 | | 0,394055 |
| GSM2262218 | 0 | 0,706652 | 0 | 0,599098 | 0 | 2,9154015 | 0 | 0 | 0,6709808 | 0,746732 | | 0,706652 |
| GSM2262226 | 0 | 0,456091 | 0 | 0,522584 | 0 | 1,8898553 | 0 | 0 | 0,5869588 | 0,808821 | | 0,456091 |
| GSM2262233 | 0 | 0,915975 | 0 | 0,322821 | 0 | 3,9444501 | 0,0911238 | 0 | 0,7132285 | 0,707159 | | 0,915975 |
| GSM2262234 | 0 | 1,00371 | 0,022002 | 0,310032 | 0 | 3,6847774 | 0 | 0 | 0,6488373 | 0,762354 | | 1,025712 |
| GSM2262236 | 0 | 0,990033 | 0 | 0,519595 | 0 | 3,3281676 | 0 | 0 | 0,628414 | 0,77961 | | 0,990033 |
| GSM2262240 | 0 | 0,744176 | 0 | 0,438959 | 0 | 3,5507001 | 0 | 0 | 0,7149796 | 0,705926 | | 0,744176 |
| GSM2262244 | 0 | 0,986687 | 0 | 0,391182 | 0 | 2,7276224 | 0 | 0 | 0,5457269 | 0,837824 | | 0,986687 |
| GSM2262247 | 0 | 0,547707 | 0 | 0,361948 | 0,014838 | 2,4304614 | 0 | 0 | 0,6394543 | 0,769293 | | 0,547707 |
| GSM2262248 | 0 | 0,592637 | 0,038605 | 0,279471 | 0 | 1,975801 | 0 | 0 | 0,5976597 | 0,801903 | | 0,631242 |
| GSM2262255 | 0 | 0,46038 | 0 | 0,249607 | 0 | 1,1943298 | 0,0233563 | 0 | 0,5023066 | 0,864724 | | 0,46038 |
| GSM2262260 | 0 | 0,972855 | 0 | 0,378254 | 0 | 5,268973 | 0 | 0 | 0,6993803 | 0,71623 | | 0,972855 |
| GSM2262262 | 0 | 0,966362 | 0 | 0,448403 | 0 | 3,5120175 | 0 | 0 | 0,612687 | 0,790369 | | 0,966362 |
| GSM2262269 | 0 | 0,509396 | 0 | 0,393804 | 0 | 2,0060889 | 0 | 0 | 0,58793 | 0,808169 | | 0,509396 |
| GSM2262271 | 0 | 1,182078 | 0 | 0,644281 | 0 | 3,0487388 | 0 | 0 | 0,6198505 | 0,789072 | | 1,182078 |
| GSM2262272 | 0 | 0,700369 | 0 | 0,317186 | 0 | 1,45982 | 0,1054436 | 0 | 0,498744 | 0,866602 | | 0,700369 |
| GSM2262273 | 0 | 1,203043 | 0 | 0,593082 | 0 | 4,2214311 | 0 | 0 | 0,6009677 | 0,799138 | | 1,203043 |
| GSM2262277 | 0 | 0,658227 | 0,015692 | 1,03497 | 0 | 4,5677767 | 0,2729454 | 0 | 0,7229883 | 0,700754 | | 0,673919 |
| GSM2262278 | 0 | 1,037532 | 0 | 0,74325 | 0 | 3,495426 | 0 | 0 | 0,5990817 | 0,80103 | | 1,037532 |
| GSM2262283 | 0 | 0,522339 | 0 | 0,306634 | 0 | 1,8760601 | 0,0075853 | 0 | 0,625889 | 0,781065 | | 0,522339 |
| GSM2262284 | 0 | 1,376804 | 0 | 0,786238 | 0 | 3,3767671 | 0 | 0 | 0,4761095 | 0,880243 | | 1,376804 |
| GSM2262285 | 0 | 0,806072 | 0 | 0,464876 | 0 | 2,2459098 | 0,002849 | 0 | 0,7018196 | 0,729872 | | 0,806072 |
| GSM2262286 | 0 | 0,986543 | 0 | 0,533337 | 0 | 4,0593555 | 0 | 0 | 0,6686523 | 0,746515 | | 0,986543 |
| GSM2262287 | 0 | 0,993568 | 0 | 0,303366 | 0 | 2,5073087 | 0 | 0 | 0,6023634 | 0,80012 | | 0,993568 |
| GSM2262288 | 0 | 1,005162 | 0 | 0,503098 | 0 | 5,5618901 | 0 | 0 | 0,6996908 | 0,716361 | | 1,005162 |
| GSM2262289 | 0 | 0,779704 | 0 | 0,630685 | 0 | 3,9592365 | 0,065089 | 0 | 0,6900201 | 0,728091 | | 0,779704 |
| GSM2262295 | 0 | 0,900649 | 0 | 0,687374 | 0 | 1,8044756 | 0,3232444 | 0 | 0,4281832 | 0,905146 | | 0,900649 |
| GSM2262298 | 0 | 0,364962 | 0 | 0,304904 | 0 | 0,8727575 | 0,0331431 | 0 | 0,4298007 | 0,905622 | | 0,364962 |
| GSM2262302 | 0 | 0,334741 | 0 | 0,164472 | 0 | 0,5732412 | 0,080698 | 0 | 0,5142929 | 0,857867 | | 0,334741 |
| GSM2262306 | 0 | 1,509702 | 0 | 0,73121 | 0 | 5,5079701 | 0 | 0 | 0,6251911 | 0,781085 | | 1,509702 |
| GSM2262309 | 0 | 0,473382 | 0 | 0,32367 | 0 | 1,8132344 | 0,0141145 | 0 | 0,5864123 | 0,809329 | | 0,473382 |
| GSM2262310 | 0 | 0,434881 | 0 | 0,30525 | 0 | 1,6889586 | 0 | 0 | 0,630366 | 0,777411 | | 0,434881 |
| **average** | **0,000704** | **0,788224** | **0,044172** | **0,515585** | **0,002033** | **3,2912066** | **0,024814** | | | | | |

**CIBERSORTx_GSE85217_GP3**

| Sample | Bcells | TCD4 | TCD8 | Tgd | NK | MoMaDC | granulocytes | P-value | Correlation | RMSE | ratio mono/T cells |
|---|---|---|---|---|---|---|---|---|---|---|---|
| GSM2261539 | 0 | 0,339832 | 0,054648 | 0,089426 | 0 | 0,44303 | 0,07306385 | 0 | 0,3989844 | 0,919762 | 0,915530295 |
| GSM2261545 | 0 | 0,350243 | 0 | 0,132466 | 0 | 0,457904 | 0,05938724 | 0 | 0,3805862 | 0,929204 | 0,948612796 |
| GSM2261554 | 0 | 0,379944 | 0 | 0,034962 | 0,023633 | 0,515368 | 0,04609337 | 0 | 0,4776851 | 0,879683 | 1,242133122 |
| GSM2261555 | 0 | 0,325486 | 0 | 0,186661 | 0 | 0,389968 | 0,09788521 | 0 | 0,3499741 | 0,943313 | 0,76143693 |
| GSM2261559 | 0 | 0,412242 | 0 | 0,187824 | 0 | 0,225329 | 0,17460546 | 0 | 0,3653978 | 0,935674 | 0,375506286 |
| GSM2261560 | 0 | 0,33191 | 0 | 0,134926 | 0 | 0,471057 | 0,0621067 | 0 | 0,3287346 | 0,955518 | 1,009040812 |
| GSM2261561 | 0 | 0,351578 | 0 | 0,119984 | 0 | 0,450108 | 0,07832994 | 0 | 0,484039 | 0,87573 | 0,954504909 |
| GSM2261565 | 0 | 0,136769 | 0,126071 | 0,068912 | 0 | 0,668248 | 0 | 0 | 0,459582 | 0,892669 | 2,014301316 |
| GSM2261566 | 0 | 0,296221 | 0 | 0,189286 | 0 | 0,424578 | 0,08991439 | 0 | 0,453068 | 0,891142 | 0,874504429 |
| GSM2261577 | 0 | 0,432194 | 0 | 0,143105 | 0 | 0,373086 | 0,05161516 | 0 | 0,4174269 | 0,910485 | 0,6485091 |
| GSM2261581 | 0,008076 | 0,165188 | 0,02601 | 0,107482 | 0 | 0,683598 | 0,00964663 | 0 | 0,6096076 | 0,792979 | 2,288731462 |
| GSM2261590 | 0 | 0,312574 | 0 | 0,149276 | 0,00693 | 0,49282 | 0,03839971 | 0 | 0,5202615 | 0,854703 | 1,067058051 |
| GSM2261592 | 0 | 0,251439 | 0 | 0,137688 | 0,001603 | 0,586064 | 0,02320745 | 0,77 | 0,0092904 | 1,121121 | 1,506101418 |
| GSM2261593 | 0 | 0,297216 | 0,058527 | 0,085489 | 0 | 0,558768 | 0 | 0 | 0,5594566 | 0,831396 | 1,266382738 |
| GSM2261594 | 0 | 0,260203 | 0,032124 | 0,123528 | 0 | 0,512902 | 0,07124301 | 0 | 0,4749077 | 0,879823 | 1,233367235 |
| GSM2261601 | 0 | 0,344624 | 0 | 0,080294 | 0 | 0,538522 | 0,03655952 | 0 | 0,4071596 | 0,918157 | 1,267352308 |
| GSM2261604 | 0 | 0,275713 | 0 | 0,209353 | 0 | 0,398696 | 0,11623767 | 0 | 0,4135276 | 0,911729 | 0,821941678 |
| GSM2261605 | 0 | 0,16271 | 0,010474 | 0,0593 | 0,00817 | 0,759347 | 0 | 0 | 0,6773557 | 0,737439 | 3,266235631 |
| GSM2261612 | 0 | 0,317746 | 0 | 0,107697 | 0 | 0,574557 | 0 | 0 | 0,4538985 | 0,892774 | 1,350490113 |
| GSM2261613 | 0 | 0,242959 | 0 | 0,164368 | 0 | 0,498378 | 0,09429467 | 0 | 0,5261149 | 0,850226 | 1,223533908 |
| GSM2261624 | 0 | 0,152042 | 0,051056 | 0,058198 | 0 | 0,738704 | 0 | 0 | 0,5198403 | 0,85775 | 2,827079359 |
| GSM2261625 | 0 | 0,278217 | 0 | 0,147077 | 0 | 0,557783 | 0,0169222 | 0 | 0,4383712 | 0,900766 | 1,311522556 |
| GSM2261628 | 0 | 0,398793 | 0 | 0,11491 | 0 | 0,339492 | 0,14680537 | 0 | 0,3927966 | 0,922276 | 0,660873675 |
| GSM2261635 | 0 | 0,169048 | 0 | 0,143838 | 0 | 0,687114 | 0 | 0 | 0,5695864 | 0,820977 | 2,19605382 |
| GSM2261636 | 0 | 0,216632 | 0,01801 | 0,108046 | 0 | 0,636475 | 0,02083858 | 0 | 0,5957147 | 0,804431 | 1,857307139 |
| GSM2261642 | 0 | 0,381927 | 0 | 0,093951 | 0 | 0,416883 | 0,10723914 | 0 | 0,3570411 | 0,940656 | 0,876028705 |
| GSM2261643 | 0,001604 | 0,323279 | 0 | 0,152591 | 0,00959 | 0,432005 | 0,08093031 | 0 | 0,4945066 | 0,869424 | 0,907821517 |
| GSM2261645 | 0 | 0,298898 | 0,01425 | 0,14221 | 0,011164 | 0,435433 | 0,09804438 | 0 | 0,4886495 | 0,872298 | 0,956243571 |
| GSM2261646 | 0 | 0,330503 | 0 | 0,178681 | 0,005096 | 0,366721 | 0,11899913 | 0 | 0,4373522 | 0,899283 | 0,720214104 |
| GSM2261649 | 0 | 0,333831 | 0,020026 | 0,075121 | 0,07324 | 0,444648 | 0,05313396 | 0 | 0,4283811 | 0,90383 | 1,036529297 |
| GSM2261667 | 0 | 0,313744 | 0 | 0,108381 | 0 | 0,549039 | 0,02883555 | 0 | 0,5680388 | 0,826357 | 1,300652425 |
| GSM2261668 | 0,001652 | 0,382979 | 0 | 0,10146 | 0,000368 | 0,411167 | 0,10237341 | 0 | 0,4498989 | 0,89412 | 0,848749188 |
| GSM2261675 | 0 | 0,244063 | 0,019386 | 0,134562 | 0 | 0,513049 | 0,08894087 | 0 | 0,5397336 | 0,842362 | 1,289033542 |
| GSM2261676 | 0 | 0,237753 | 0,024423 | 0,06541 | 0,001729 | 0,66391 | 0,0067759 | 0 | 0,6575775 | 0,760446 | 2,026675046 |
| GSM2261678 | 0 | 0,414422 | 0 | 0,134173 | 0 | 0,355414 | 0,09599215 | 0 | 0,3604477 | 0,937853 | 0,647862935 |
| GSM2261700 | 0 | 0,268481 | 0 | 0,153896 | 0,013745 | 0,50312 | 0,06075909 | 0 | 0,4640847 | 0,885468 | 1,191163824 |
| GSM2261701 | 0 | 0,27398 | 0 | 0,11724 | 0 | 0,585268 | 0,02351185 | 0 | 0,4202482 | 0,912164 | 1,49600697 |
| GSM2261711 | 0 | 0,351428 | 0 | 0,121071 | 0,024335 | 0,460253 | 0,04291301 | 0 | 0,4347814 | 0,901108 | 0,974081465 |
| GSM2261720 | 0 | 0,253879 | 0 | 0,24305 | 0 | 0,341421 | 0,16164997 | 0 | 0,4727844 | 0,880114 | 0,687063074 |
| GSM2261722 | 0 | 0,298674 | 0 | 0,168416 | 0 | 0,446724 | 0,08618653 | 0 | 0,4635144 | 0,885779 | 0,956398062 |
| GSM2261732 | 0 | 0,304996 | 0 | 0,095435 | 0 | 0,59957 | 0 | 0 | 0,5470841 | 0,838007 | 1,497312399 |
| GSM2261734 | 0 | 0,241233 | 0 | 0,132746 | 0,065052 | 0,553859 | 0,00711066 | 0 | 0,4908085 | 0,870181 | 1,480989675 |
| GSM2261737 | 0 | 0,306386 | 0 | 0,067542 | 0,030211 | 0,587518 | 0,00834199 | 0 | 0,5475147 | 0,837544 | 1,571204305 |
| GSM2261743 | 0 | 0,216945 | 0,027244 | 0,08606 | 0 | 0,65321 | 0,01653952 | 0 | 0,5594423 | 0,828578 | 1,977926522 |
| GSM2261745 | 0,000712 | 0,333977 | 0,039038 | 0,116972 | 0 | 0,46213 | 0,04717119 | 0 | 0,3947332 | 0,921967 | 0,94314958 |
| GSM2261748 | 0 | 0,294401 | 0,040094 | 0,107592 | 0 | 0,442556 | 0,11535704 | 0 | 0,3710685 | 0,933991 | 1,001062405 |
| GSM2261756 | 0 | 0,158728 | 0,109792 | 0,09474 | 0 | 0,63369 | 0,00304971 | 0 | 0,6147118 | 0,791424 | 1,744454978 |
| GSM2261757 | 0 | 0,252921 | 0,027756 | 0,088126 | 0 | 0,611379 | 0,01981777 | 0 | 0,6266333 | 0,785615 | 1,657734581 |
| GSM2261758 | 0 | 0,249278 | 0 | 0,055956 | 0,010148 | 0,661912 | 0,02270601 | 0 | 0,7571407 | 0,682946 | 2,168535067 |
| GSM2261763 | 0 | 0,318874 | 0 | 0,167761 | 0 | 0,383852 | 0,1295128 | 0 | 0,555284 | 0,836656 | 0,788788628 |
| GSM2261768 | 0 | 0,317641 | 0,014766 | 0,095397 | 0,002385 | 0,533266 | 0,03654618 | 0 | 0,4813294 | 0,876928 | 1,246520401 |
| GSM2261793 | 0,002894 | 0,325332 | 0 | 0,197097 | 0 | 0,319744 | 0,15493243 | 0 | 0,4061 | 0,915005 | 0,612034152 |
| GSM2261794 | 0 | 0,448194 | 0 | 0,178843 | 0 | 0,2948 | 0,07816307 | 0 | 0,4420648 | 0,898033 | 0,470148079 |
| GSM2261797 | 0 | 0,365326 | 0 | 0,16966 | 0 | 0,354266 | 0,11074816 | 0 | 0,3019483 | 0,965555 | 0,662196412 |
| GSM2261818 | 0 | 0,297251 | 0 | 0,175256 | 0,007319 | 0,445581 | 0,07459317 | 0 | 0,5039959 | 0,863666 | 0,943016337 |
| GSM2261820 | 0 | 0,370173 | 0 | 0,065905 | 0,027351 | 0,45592 | 0,08065152 | 0 | 0,3953465 | 0,921862 | 1,045503691 |
| GSM2261840 | 0,009329 | 0,399535 | 0 | 0,192262 | 0 | 0,252195 | 0,14667882 | 0 | 0,4344668 | 0,901239 | 0,426151623 |
| GSM2261841 | 0,009313 | 0,367566 | 0 | 0,165812 | 0 | 0,328645 | 0,12866462 | 0 | 0,3987882 | 0,91862 | 0,616156801 |
| GSM2261843 | 0 | 0,424186 | 0 | 0,089011 | 0 | 0,415792 | 0,07101121 | 0 | 0,4304192 | 0,904558 | 0,810198789 |
| GSM2261845 | 0 | 0,295982 | 0,018299 | 0,184893 | 0,029096 | 0,354921 | 0,11680898 | 0 | 0,4560563 | 0,889052 | 0,711015798 |
| GSM2261850 | 0 | 0,126795 | 0,042133 | 0,189256 | 0 | 0,56041 | 0,08140592 | 0 | 0,6079054 | 0,796139 | 1,564588765 |
| GSM2261855 | 0 | 0,174732 | 0 | 0,097558 | 0 | 0,72771 | 0 | 0 | 0,6826468 | 0,73521 | 2,67255358 |
| GSM2261863 | 0,021679 | 0,180178 | 0,071018 | 0,062452 | 0 | 0,350926 | 0,31374767 | 0 | 0,6454475 | 0,775556 | 1,118852247 |
| GSM2261869 | 0 | 0,226123 | 0,053418 | 0,11884 | 0 | 0,571409 | 0,03021056 | 0 | 0,5136137 | 0,85742 | 1,434328287 |
| GSM2261874 | 0 | 0,218827 | 0,023953 | 0,051881 | 0 | 0,666497 | 0,03884268 | 0 | 0,5433667 | 0,839818 | 2,261912579 |
| GSM2261878 | 0 | 0,346315 | 0 | 0,129847 | 0 | 0,468978 | 0,05485936 | 0 | 0,5493772 | 0,839652 | 0,984911395 |
| GSM2261881 | 0 | 0,296745 | 0,012172 | 0,11196 | 0 | 0,543323 | 0,03579964 | 0 | 0,324478 | 0,961378 | 1,290930105 |
| GSM2261882 | 0 | 0,174231 | 0,055046 | 0,046524 | 0,039693 | 0,682184 | 0,00232164 | 0 | 0,6098018 | 0,792681 | 2,473465046 |
| GSM2261883 | 0 | 0,255699 | 0,007605 | 0,142193 | 0 | 0,594503 | 0 | 0 | 0,4958136 | 0,868219 | 1,466107131 |
| GSM2261885 | 0 | 0,340019 | 0 | 0,098828 | 0 | 0,519576 | 0,04157682 | 0 | 0,3977569 | 0,922261 | 1,18395727 |
| GSM2261896 | 0 | 0,302659 | 0 | 0,180198 | 0 | 0,403748 | 0,11339395 | 0 | 0,3593559 | 0,939034 | 0,836164063 |
| GSM2261900 | 0 | 0,298716 | 0 | 0,092483 | 0 | 0,552782 | 0,05601917 | 0 | 0,4367037 | 0,902406 | 1,413046165 |
| GSM2261919 | 0 | 0,285846 | 0,025684 | 0,160097 | 0 | 0,415305 | 0,1130687 | 0 | 0,5163154 | 0,857292 | 0,880579299 |
| GSM2261926 | 0 | 0,337733 | 0 | 0,114109 | 0 | 0,486049 | 0,06210925 | 0 | 0,5089006 | 0,86188 | 1,075705499 |

| | | | | | | | | | | | |
|---|---|---|---|---|---|---|---|---|---|---|---|
| GSM2261927 | 0 | 0,291025 | 0,002423 | 0,074273 | 0 | 0,618982 | 0,01329702 | 0 | 0,507765 | 0,862172 | 1,683293109 |
| GSM2261943 | 0 | 0,121191 | 0,036173 | 0,068977 | 0 | 0,773659 | 0 | 0 | 0,7627592 | 0,659485 | 3,418113529 |
| GSM2261959 | 0 | 0,319036 | 0,005591 | 0,125156 | 0 | 0,509924 | 0,04029369 | 0 | 0,5181384 | 0,85615 | 1,133712884 |
| GSM2261978 | 0 | 0,242686 | 0,081534 | 0,045494 | 0 | 0,630286 | 0 | 0 | 0,6244936 | 0,786463 | 1,704793391 |
| GSM2261980 | 0 | 0,283319 | 0,058487 | 0,144604 | 0 | 0,46883 | 0,0447594 | 0 | 0,4364772 | 0,900136 | 0,963855221 |
| GSM2261981 | 0 | 0,22393 | 0 | 0,162563 | 0,002612 | 0,513951 | 0,09694326 | 0 | 0,2764317 | 0,98689 | 1,329779681 |
| GSM2262014 | 0 | 0,301363 | 0 | 0,132108 | 0 | 0,535871 | 0,03065814 | 0 | 0,5832039 | 0,817227 | 1,236232899 |
| GSM2262015 | 0,00206 | 0,408096 | 0 | 0,100615 | 0,001825 | 0,45672 | 0,0306831 | 0 | 0,4678447 | 0,8851 | 0,897799501 |
| GSM2262016 | 0 | 0,313584 | 0,008914 | 0,185557 | 0 | 0,388367 | 0,10357712 | 0 | 0,403206 | 0,916742 | 0,764418164 |
| GSM2262018 | 0,000444 | 0,420561 | 0,038728 | 0,114952 | 0,014566 | 0,281795 | 0,12895427 | 0 | 0,331119 | 0,951195 | 0,490724933 |
| GSM2262021 | 0 | 0,254264 | 0 | 0,173866 | 0 | 0,492382 | 0,07948882 | 0 | 0,3490045 | 0,947159 | 1,150075686 |
| GSM2262022 | 0,002085 | 0,308082 | 0 | 0,225655 | 0 | 0,314723 | 0,14945388 | 0 | 0,3748216 | 0,930949 | 0,589659541 |
| GSM2262032 | 0,006476 | 0,257842 | 0 | 0,184268 | 0 | 0,515833 | 0,03558037 | 0 | 0,5030862 | 0,863571 | 1,166754103 |
| GSM2262036 | 0 | 0,248793 | 0 | 0,10466 | 0 | 0,602479 | 0,0440675 | 0 | 0,5508341 | 0,834691 | 1,704550132 |
| GSM2262038 | 0 | 0,319202 | 0,027261 | 0,075092 | 0 | 0,496177 | 0,0822684 | 0 | 0,4909803 | 0,871788 | 1,177016648 |
| GSM2262039 | 0 | 0,303535 | 0 | 0,126904 | 0 | 0,558996 | 0,01056464 | 0 | 0,5524251 | 0,835233 | 1,29866555 |
| GSM2262048 | 0 | 0,341206 | 0 | 0,190593 | 0 | 0,392609 | 0,07559145 | 0 | 0,3142478 | 0,960783 | 0,7382661 |
| GSM2262056 | 0 | 0,394557 | 0 | 0,096161 | 0 | 0,418241 | 0,09104096 | 0 | 0,3890769 | 0,924869 | 0,8523054 |
| GSM2262058 | 0 | 0,294851 | 0,075872 | 0,128354 | 0 | 0,425221 | 0,07570249 | 0 | 0,5727282 | 0,82664 | 0,852015908 |
| GSM2262065 | 0 | 0,211308 | 0,096665 | 0,113645 | 0 | 0,555936 | 0,0224464 | 0 | 0,3792731 | 0,932938 | 1,318578899 |
| GSM2262070 | 0 | 0,361743 | 0 | 0,132984 | 0 | 0,44259 | 0,06268381 | 0 | 0,3669483 | 0,935705 | 0,89461442 |
| GSM2262072 | 0 | 0,295235 | 0 | 0,186857 | 0,016839 | 0,3859 | 0,11516902 | 0 | 0,4361594 | 0,899654 | 0,80046937 |
| GSM2262075 | 0 | 0,370711 | 0,050569 | 0,013285 | 0,053485 | 0,511951 | 0 | 0 | 0,2995212 | 0,971566 | 1,178080499 |
| GSM2262079 | 0 | 0,370298 | 0 | 0,110101 | 0,002999 | 0,447332 | 0,06926935 | 0 | 0,4474734 | 0,895152 | 0,931168506 |
| GSM2262087 | 0 | 0,34713 | 0 | 0,035586 | 0,107308 | 0,360875 | 0,14910101 | 0 | 0,3507288 | 0,942669 | 0,942933197 |
| GSM2262089 | 0 | 0,21549 | 0,040909 | 0,195682 | 0,002151 | 0,35907 | 0,18669875 | 0 | 0,4970119 | 0,866783 | 0,794261212 |
| GSM2262090 | 0,007957 | 0,365869 | 0 | 0,12584 | 0 | 0,38866 | 0,11167517 | 0 | 0,3814708 | 0,927626 | 0,790427218 |
| GSM2262092 | 0,007288 | 0,39548 | 0 | 0,132874 | 0 | 0,31997 | 0,14438751 | 0 | 0,412341 | 0,912428 | 0,605596523 |
| GSM2262097 | 0 | 0,187064 | 0,20312 | 0,176769 | 0 | 0,339795 | 0,09325253 | 0 | 0,3691071 | 0,935101 | 0,59933537 |
| GSM2262101 | 0 | 0,386197 | 0 | 0,134248 | 0 | 0,309754 | 0,1697999 | 0 | 0,3513438 | 0,941912 | 0,595170968 |
| GSM2262108 | 0 | 0,302165 | 0,02645 | 0,135147 | 0 | 0,498657 | 0,03758046 | 0 | 0,3579253 | 0,941664 | 1,075242196 |
| GSM2262109 | 0 | 0,261409 | 0,045728 | 0,11189 | 0,007801 | 0,474574 | 0,09859808 | 0 | 0,4493838 | 0,893473 | 1,13256047 |
| GSM2262110 | 0 | 0,194605 | 0,05218 | 0,104626 | 0 | 0,491097 | 0,15749101 | 0 | 0,5386984 | 0,842347 | 1,397499353 |
| GSM2262112 | 0 | 0,223619 | 0,039978 | 0,178698 | 0,011466 | 0,515123 | 0,03111542 | 0 | 0,5390655 | 0,842145 | 1,164661145 |
| GSM2262114 | 0 | 0,31189 | 0,00525 | 0,108101 | 0,011056 | 0,487132 | 0,07657149 | 0 | 0,4537366 | 0,891517 | 1,145544286 |
| GSM2262118 | 0 | 0,372124 | 0 | 0,061374 | 0,005701 | 0,530414 | 0,03038659 | 0 | 0,4528919 | 0,893443 | 1,22356516 |
| GSM2262125 | 0 | 0,345088 | 0,051711 | 0,122033 | 0 | 0,315863 | 0,16530499 | 0 | 0,3519087 | 0,941429 | 0,608796701 |
| GSM2262129 | 0 | 0,349432 | 0 | 0,107024 | 0 | 0,523894 | 0,01965026 | 0 | 0,3557817 | 0,943992 | 1,147742445 |
| GSM2262133 | 0 | 0,342154 | 0 | 0,080933 | 0 | 0,5607 | 0,01621314 | 0 | 0,4178467 | 0,912948 | 1,325258795 |
| GSM2262134 | 0 | 0,377987 | 0 | 0,069504 | 0,005375 | 0,547133 | 0 | 0 | 0,4743297 | 0,881691 | 1,222667437 |
| GSM2262137 | 0 | 0,444456 | 0 | 0,127217 | 0,005447 | 0,340655 | 0,08222439 | 0 | 0,3782491 | 0,929465 | 0,595889787 |
| GSM2262140 | 0 | 0,285376 | 0,054433 | 0,126742 | 0,014192 | 0,467211 | 0,05204571 | 0 | 0,3892269 | 0,92452 | 1,001415039 |
| GSM2262141 | 0 | 0,415341 | 0 | 0,135811 | 0 | 0,392871 | 0,05597752 | 0 | 0,3514334 | 0,942707 | 0,71281827 |
| GSM2262143 | 0 | 0,374745 | 0 | 0,112771 | 0 | 0,492074 | 0,02041012 | 0 | 0,3752814 | 0,932895 | 1,009348791 |
| GSM2262150 | 0 | 0,265947 | 0 | 0,06734 | 0 | 0,666713 | 0 | 0 | 0,5932932 | 0,806249 | 2,000413985 |
| GSM2262153 | 0 | 0,421456 | 0 | 0,103494 | 0,002502 | 0,380366 | 0,09218122 | 0 | 0,4644542 | 0,887104 | 0,724576592 |
| GSM2262155 | 0 | 0,191178 | 0 | 0,067405 | 0 | 0,741417 | 0 | 0 | 0,7035265 | 0,717157 | 2,86723196 |
| GSM2262157 | 0 | 0,272903 | 0 | 0,07352 | 0,016944 | 0,636633 | 0 | 0 | 0,6416858 | 0,77414 | 1,837736354 |
| GSM2262158 | 0 | 0,477319 | 0 | 0,043997 | 0,025263 | 0,369346 | 0,08407583 | 0 | 0,390585 | 0,924293 | 0,708488652 |
| GSM2262184 | 0 | 0,296921 | 0,001791 | 0,113478 | 0,017404 | 0,456528 | 0,11387697 | 0 | 0,3519385 | 0,944067 | 1,10756694 |
| GSM2262192 | 0 | 0,361969 | 0 | 0,19305 | 0 | 0,304487 | 0,14049282 | 0 | 0,3355429 | 0,949626 | 0,548606322 |
| GSM2262195 | 0,019089 | 0,129576 | 0,062056 | 0,087279 | 0,005585 | 0,647026 | 0,04938896 | 0 | 0,6457936 | 0,767701 | 2,319831505 |
| GSM2262204 | 0 | 0,348078 | 0 | 0,149978 | 0 | 0,413122 | 0,0888231 | 0 | 0,2990947 | 0,967973 | 0,829469366 |
| GSM2262205 | 0 | 0,35045 | 0 | 0,088199 | 0 | 0,498654 | 0,06269725 | 0 | 0,3460754 | 0,948508 | 1,136793714 |
| GSM2262208 | 0 | 0,249936 | 0 | 0,088946 | 0 | 0,645986 | 0,01513198 | 0 | 0,5711593 | 0,821183 | 1,906229804 |
| GSM2262209 | 0 | 0,324151 | 0 | 0,124856 | 0 | 0,491984 | 0,0590093 | 0 | 0,3876344 | 0,926527 | 1,095718664 |
| GSM2262229 | 0,002918 | 0,304692 | 0 | 0,09296 | 0,019064 | 0,552132 | 0,02823328 | 0 | 0,5707688 | 0,824349 | 1,38847927 |
| GSM2262249 | 0 | 0,374452 | 0 | 0,248976 | 0 | 0,290524 | 0,08604792 | 0 | 0,4141227 | 0,911917 | 0,466011432 |
| GSM2262254 | 0 | 0,374033 | 0 | 0,10677 | 0 | 0,519197 | 0 | 0 | 0,5235446 | 0,85393 | 1,07985276 |
| GSM2262259 | 0 | 0,267561 | 0 | 0,18804 | 0 | 0,462259 | 0,08214073 | 0 | 0,4305627 | 0,903364 | 1,014613702 |
| GSM2262261 | 0 | 0,349498 | 0 | 0,235288 | 0 | 0,341104 | 0,07410948 | 0 | 0,3396767 | 0,949187 | 0,583296447 |
| GSM2262263 | 0 | 0,39281 | 0 | 0,146168 | 0 | 0,319012 | 0,14200997 | 0 | 0,3354668 | 0,949338 | 0,591882732 |
| GSM2262275 | 0 | 0,327559 | 0 | 0,134081 | 0,00644 | 0,314628 | 0,21729216 | 0 | 0,4602372 | 0,887854 | 0,681546513 |
| GSM2262291 | 0 | 0,249084 | 0 | 0,093154 | 0 | 0,657762 | 0 | 0 | 0,5870328 | 0,810373 | 1,921946445 |
| GSM2262294 | 0 | 0,419357 | 0 | 0,100454 | 0,003606 | 0,38901 | 0,08757335 | 0 | 0,4021668 | 0,918126 | 0,74836771 |
| GSM2262296 | 0 | 0,20249 | 0,069457 | 0,190965 | 0,023198 | 0,488637 | 0,02525359 | 0 | 0,3708448 | 0,935945 | 1,055571606 |
| GSM2262299 | 0 | 0,338097 | 0,001892 | 0,116058 | 0 | 0,507387 | 0,03656609 | 0 | 0,5090247 | 0,861627 | 1,112576985 |
| GSM2262300 | 0 | 0,404535 | 0 | 0,163417 | 0,001292 | 0,40466 | 0,02609555 | 0 | 0,3689906 | 0,934529 | 0,712490666 |
| GSM2262301 | 0 | 0,369919 | 0 | 0,209146 | 0 | 0,376301 | 0,04463358 | 0 | 0,3857081 | 0,926084 | 0,649842595 |
| GSM2262308 | 0 | 0,32734 | 0 | 0,036505 | 0,006875 | 0,629281 | 0 | 0 | 0,59108 | 0,809812 | 1,729529757 |
| average | 0,000719 | 0,304918 | 0,015001 | 0,124033 | 0,00543 | 0,4844 | 0,06549869 | | | | |

**SES_CIBERSORTx_GSE85217_GP3**

| Sample | Bcells | TCD4 | TCD8 | Tgd | NK | MoMaDC | granulocytes | P-value | Correlation | RMSE | | Tab |
|---|---|---|---|---|---|---|---|---|---|---|---|---|
| GSM2261539 | 0 | 1,08901 | 0,175122 | 0,28657 | 0 | 1,419715 | 0,23413709 | 0 | 0,39898443 | 0,919762 | | 1,264133 |
| GSM2261545 | 0 | 1,987295 | 0 | 0,751621 | 0 | 2,598171 | 0,3369664 | 0 | 0,3805862 | 0,929204 | | 1,987295 |
| GSM2261554 | 0 | 1,289376 | 0 | 0,118647 | 0,0802 | 1,748952 | 0,15642235 | 0 | 0,47768508 | 0,879683 | | 1,289376 |
| GSM2261555 | 0 | 1,063764 | 0 | 0,610053 | 0 | 1,274506 | 0,3199118 | 0 | 0,34997414 | 0,943313 | | 1,063764 |
| GSM2261559 | 0 | 0,661578 | 0 | 0,301425 | 0 | 0,361614 | 0,28021182 | 0 | 0,36539782 | 0,935674 | | 0,661578 |
| GSM2261560 | 0 | 1,056116 | 0 | 0,429327 | 0 | 1,498872 | 0,19761941 | 0 | 0,32873465 | 0,955518 | | 1,056116 |
| GSM2261561 | 0 | 1,004661 | 0 | 0,342865 | 0 | 1,28622 | 0,22383406 | 0 | 0,48403897 | 0,87573 | | 1,004661 |
| GSM2261565 | 0 | 0,571013 | 0,526348 | 0,287708 | 0 | 2,789945 | 0 | 0 | 0,45958202 | 0,892669 | | 1,097361 |
| GSM2261566 | 0 | 0,959864 | 0 | 0,613357 | 0 | 1,375789 | 0,29135549 | 0 | 0,45306802 | 0,891142 | | 0,959864 |
| GSM2261577 | 0 | 1,374496 | 0 | 0,455113 | 0 | 1,186518 | 0,16415052 | 0 | 0,41742689 | 0,910485 | | 1,374496 |
| GSM2261581 | 0,031488 | 0,644102 | 0,101419 | 0,419093 | 0 | 2,665488 | 0,03761417 | 0 | 0,60960762 | 0,792979 | | 0,745521 |
| GSM2261590 | 0 | 0,975453 | 0 | 0,465847 | 0,021628 | 1,53795 | 0,1198344 | 0 | 0,52026151 | 0,854703 | | 0,975453 |
| GSM2261592 | 0 | 1,588517 | 0 | 0,869871 | 0,010124 | 3,702582 | 0,14661797 | 0,77 | 0,00929035 | 1,121121 | | 1,588517 |
| GSM2261593 | 0 | 1,253429 | 0,246822 | 0,360527 | 0 | 2,356457 | 0 | 0 | 0,55945655 | 0,831396 | | 1,500251 |
| GSM2261594 | 0 | 1,064822 | 0,13146 | 0,505512 | 0 | 2,098937 | 0,29154614 | 0 | 0,47490774 | 0,879823 | | 1,196282 |
| GSM2261601 | 0 | 1,19055 | 0 | 0,277387 | 0 | 1,860393 | 0,12629958 | 0 | 0,40715959 | 0,918157 | | 1,19055 |
| GSM2261604 | 0 | 0,679746 | 0 | 0,516141 | 0 | 0,982949 | 0,28657345 | 0 | 0,41352761 | 0,911729 | | 0,679746 |
| GSM2261605 | 0 | 1,203471 | 0,077466 | 0,438606 | 0,060427 | 5,616432 | 0 | 0 | 0,67735575 | 0,737439 | | 1,280937 |
| GSM2261612 | 0 | 0,744987 | 0 | 0,252507 | 0 | 1,347106 | 0 | 0 | 0,45389853 | 0,892774 | | 0,744987 |
| GSM2261613 | 0 | 0,762687 | 0 | 0,51598 | 0 | 1,564492 | 0,29600658 | 0 | 0,52611488 | 0,850226 | | 0,762687 |
| GSM2261624 | 0 | 1,685513 | 0,565996 | 0,645169 | 0 | 8,189139 | 0 | 0 | 0,51984027 | 0,85775 | | 2,251509 |
| GSM2261625 | 0 | 1,630131 | 0 | 0,861756 | 0 | 3,268167 | 0,09915063 | 0 | 0,43837124 | 0,900766 | | 1,630131 |
| GSM2261628 | 0 | 0,890965 | 0 | 0,256726 | 0 | 0,758479 | 0,32798614 | 0 | 0,39279664 | 0,922276 | | 0,890965 |
| GSM2261635 | 0 | 0,681343 | 0 | 0,579736 | 0 | 2,769398 | 0 | 0 | 0,56958643 | 0,820977 | | 0,681343 |
| GSM2261636 | 0 | 0,924306 | 0,076842 | 0,461 | 0 | 2,715659 | 0,08891236 | 0 | 0,59571474 | 0,804431 | | 1,001148 |
| GSM2261642 | 0 | 1,072519 | 0 | 0,263832 | 0 | 1,170682 | 0,30114678 | 0 | 0,35704113 | 0,940656 | | 1,072519 |
| GSM2261643 | 0,005118 | 1,031362 | 0 | 0,486815 | 0,030595 | 1,378234 | 0,25819333 | 0 | 0,49450656 | 0,869424 | | 1,031362 |
| GSM2261645 | 0 | 1,08489 | 0,051722 | 0,516169 | 0,040522 | 1,580461 | 0,35586468 | 0 | 0,48864946 | 0,872298 | | 1,136612 |
| GSM2261646 | 0 | 1,070578 | 0 | 0,578789 | 0,016507 | 1,187898 | 0,38546658 | 0 | 0,43735217 | 0,899283 | | 1,070578 |
| GSM2261649 | 0 | 0,941072 | 0,056453 | 0,211766 | 0,206465 | 1,253466 | 0,14978505 | 0 | 0,42838108 | 0,90383 | | 0,997525 |
| GSM2261667 | 0 | 1,584417 | 0 | 0,547328 | 0 | 2,772659 | 0,14562021 | 0 | 0,56803881 | 0,826357 | | 1,584417 |
| GSM2261668 | 0,004562 | 1,058016 | 0 | 0,280293 | 0,001017 | 1,135889 | 0,2828162 | 0 | 0,44989886 | 0,89412 | | 1,058016 |
| GSM2261675 | 0 | 1,082979 | 0,086021 | 0,59709 | 0 | 2,276549 | 0,39465691 | 0 | 0,53973364 | 0,842362 | | 1,169 |
| GSM2261676 | 0 | 0,983755 | 0,101056 | 0,270646 | 0,007154 | 2,747072 | 0,02803677 | 0 | 0,65757748 | 0,760446 | | 1,084811 |
| GSM2261678 | 0 | 1,213492 | 0 | 0,392879 | 0 | 1,040708 | 0,28108029 | 0 | 0,36044769 | 0,937853 | | 1,213492 |
| GSM2261700 | 0 | 0,97871 | 0 | 0,561005 | 0,050105 | 1,834053 | 0,22148891 | 0 | 0,46408468 | 0,885468 | | 0,97871 |
| GSM2261701 | 0 | 1,315834 | 0 | 0,563065 | 0 | 2,810845 | 0,11291951 | 0 | 0,42024819 | 0,912164 | | 1,315834 |
| GSM2261711 | 0 | 0,647819 | 0 | 0,223181 | 0,044859 | 0,848426 | 0,07910544 | 0 | 0,43478144 | 0,901108 | | 0,647819 |
| GSM2261720 | 0 | 0,533392 | 0 | 0,510641 | 0 | 0,717317 | 0,33962198 | 0 | 0,47278437 | 0,880114 | | 0,533392 |
| GSM2261722 | 0 | 1,022313 | 0 | 0,576461 | 0 | 1,529064 | 0,29500282 | 0 | 0,46351439 | 0,885779 | | 1,022313 |
| GSM2261732 | 0 | 1,07615 | 0 | 0,336733 | 0 | 2,115528 | 0 | 0 | 0,54708414 | 0,838007 | | 1,07615 |
| GSM2261734 | 0 | 0,859221 | 0 | 0,472812 | 0,2317 | 1,972727 | 0,02532666 | 0 | 0,49080854 | 0,870181 | | 0,859221 |
| GSM2261737 | 0 | 0,949353 | 0 | 0,209283 | 0,093612 | 1,820454 | 0,02584809 | 0 | 0,54751469 | 0,837544 | | 0,949353 |
| GSM2261743 | 0 | 1,247029 | 0,156604 | 0,494687 | 0 | 3,754736 | 0,09507129 | 0 | 0,55944228 | 0,828578 | | 1,403633 |
| GSM2261745 | 0,001625 | 0,762184 | 0,08909 | 0,266947 | 0 | 1,05465 | 0,10765169 | 0 | 0,39473323 | 0,921967 | | 0,851275 |
| GSM2261748 | 0 | 1,173503 | 0,159816 | 0,428869 | 0 | 1,764061 | 0,45982134 | 0 | 0,37106853 | 0,933991 | | 1,333319 |
| GSM2261756 | 0 | 0,573824 | 0,396911 | 0,342498 | 0 | 2,290875 | 0,01102512 | 0 | 0,61471179 | 0,791424 | | 0,970735 |
| GSM2261757 | 0 | 1,420234 | 0,155861 | 0,494859 | 0 | 3,433091 | 0,11128327 | 0 | 0,62663327 | 0,785615 | | 1,576094 |
| GSM2261758 | 0 | 1,078112 | 0 | 0,242006 | 0,04389 | 2,862724 | 0,09820199 | 0 | 0,75714073 | 0,682946 | | 1,078112 |
| GSM2261763 | 0 | 0,972298 | 0 | 0,511528 | 0 | 1,170425 | 0,39490473 | 0 | 0,55528403 | 0,836656 | | 0,972298 |
| GSM2261768 | 0 | 0,908477 | 0,042231 | 0,272843 | 0,00682 | 1,525182 | 0,10452495 | 0 | 0,48132936 | 0,876928 | | 0,950708 |
| GSM2261793 | 0,007459 | 0,838563 | 0 | 0,508031 | 0 | 0,824162 | 0,39934808 | 0 | 0,40610003 | 0,915005 | | 0,838563 |
| GSM2261794 | 0 | 0,933903 | 0 | 0,372655 | 0 | 0,614276 | 0,16286859 | 0 | 0,4420648 | 0,898033 | | 0,933903 |
| GSM2261797 | 0 | 1,189009 | 0 | 0,552186 | 0 | 1,153013 | 0,36044717 | 0 | 0,30194831 | 0,965555 | | 1,189009 |
| GSM2261818 | 0 | 0,541606 | 0 | 0,319324 | 0,013335 | 0,811872 | 0,13591247 | 0 | 0,50399595 | 0,863666 | | 0,541606 |
| GSM2261820 | 0 | 1,022859 | 0 | 0,182108 | 0,075575 | 1,259797 | 0,2228559 | 0 | 0,39534652 | 0,921862 | | 1,022859 |
| GSM2261840 | 0,016133 | 0,690932 | 0 | 0,332487 | 0 | 0,436132 | 0,25365774 | 0 | 0,43446676 | 0,901239 | | 0,690932 |
| GSM2261841 | 0,029782 | 1,17546 | 0 | 0,53026 | 0 | 1,050991 | 0,41146406 | 0 | 0,39878816 | 0,91862 | | 1,17546 |
| GSM2261843 | 0 | 1,034827 | 0 | 0,217149 | 0 | 1,014349 | 0,1732362 | 0 | 0,43041917 | 0,904558 | | 1,034827 |
| GSM2261845 | 0 | 0,434177 | 0,026843 | 0,27122 | 0,042681 | 0,520635 | 0,17134757 | 0 | 0,45605627 | 0,889052 | | 0,461021 |
| GSM2261850 | 0 | 0,301806 | 0,100287 | 0,450479 | 0 | 1,333924 | 0,19376749 | 0 | 0,60790536 | 0,796139 | | 0,402093 |
| GSM2261855 | 0 | 0,872206 | 0 | 0,486981 | 0 | 3,6325 | 0 | 0 | 0,68264677 | 0,73521 | | 0,872206 |
| GSM2261863 | 0,13489 | 1,1211 | 0,44189 | 0,388586 | 0 | 2,183525 | 1,95219719 | 0 | 0,64544751 | 0,775556 | | 1,56299 |
| GSM2261869 | 0 | 1,018077 | 0,240504 | 0,535053 | 0 | 2,572661 | 0,13601738 | 0 | 0,51361373 | 0,85742 | | 1,258581 |
| GSM2261874 | 0 | 1,440613 | 0,15769 | 0,341553 | 0 | 4,387783 | 0,25571506 | 0 | 0,54336673 | 0,839818 | | 1,598302 |
| GSM2261878 | 0 | 0,742923 | 0 | 0,278552 | 0 | 1,006062 | 0,11768554 | 0 | 0,5493772 | 0,839652 | | 0,742923 |
| GSM2261881 | 0 | 0,922192 | 0,037825 | 0,347938 | 0 | 1,688479 | 0,11125412 | 0 | 0,32447802 | 0,961378 | | 0,960018 |
| GSM2261882 | 0 | 0,89421 | 0,282511 | 0,238776 | 0,203717 | 3,501181 | 0,01191539 | 0 | 0,60980183 | 0,792681 | | 1,176721 |
| GSM2261883 | 0 | 0,932731 | 0,027742 | 0,518687 | 0 | 2,168607 | 0 | 0 | 0,49581358 | 0,868219 | | 0,960473 |
| GSM2261885 | 0 | 0,81755 | 0 | 0,237624 | 0 | 1,249281 | 0,09996827 | 0 | 0,39775686 | 0,922261 | | 0,81755 |
| GSM2261896 | 0 | 0,629938 | 0 | 0,375054 | 0 | 0,840338 | 0,23601154 | 0 | 0,35935592 | 0,939034 | | 0,629938 |
| GSM2261900 | 0 | 0,928102 | 0 | 0,287342 | 0 | 1,717478 | 0,17405001 | 0 | 0,43670367 | 0,902406 | | 0,928102 |
| GSM2261919 | 0 | 0,672945 | 0,060465 | 0,376903 | 0 | 0,977718 | 0,26618851 | 0 | 0,51631535 | 0,857292 | | 0,73341 |
| GSM2261926 | 0 | 1,0689 | 0 | 0,361148 | 0 | 1,53831 | 0,19657141 | 0 | 0,50890055 | 0,86188 | | 1,0689 |
| GSM2261927 | 0 | 0,867963 | 0,007226 | 0,221513 | 0 | 1,846071 | 0,03965744 | 0 | 0,50776502 | 0,862172 | | 0,875189 |
| GSM2261943 | 0 | 0,890417 | 0,265769 | 0,506788 | 0 | 5,684234 | 0 | 0 | 0,76275925 | 0,659485 | | 1,156186 |
| GSM2261959 | 0 | 0,679737 | 0,011911 | 0,266656 | 0 | 1,086442 | 0,08584954 | 0 | 0,51813839 | 0,85615 | | 0,691648 |

| | | | | | | | | | | | | |
|---|---|---|---|---|---|---|---|---|---|---|---|---|
| GSM2261978 | 0 | 0,860232 | 0,289009 | 0,16126 | 0 | 2,234134 | 0 | 0 | 0,62449365 | 0,786463 | | 1,149241 |
| GSM2261980 | 0 | 0,937626 | 0,19356 | 0,478557 | 0 | 1,551559 | 0,14812811 | 0 | 0,43647718 | 0,900136 | | 1,131186 |
| GSM2261981 | 0 | 1,1304 | 0 | 0,820619 | 0,013184 | 2,594426 | 0,48936948 | 0 | 0,27643171 | 0,98689 | | 1,1304 |
| GSM2262014 | 0 | 0,712834 | 0 | 0,312483 | 0 | 1,267532 | 0,07251776 | 0 | 0,58320387 | 0,817227 | | 0,712834 |
| GSM2262015 | 0,009966 | 1,974077 | 0 | 0,486702 | 0,00883 | 2,209286 | 0,14842285 | 0 | 0,46784468 | 0,8851 | | 1,974077 |
| GSM2262016 | 0 | 0,646572 | 0,018381 | 0,382597 | 0 | 0,800766 | 0,21356344 | 0 | 0,40320599 | 0,916742 | | 0,664953 |
| GSM2262018 | 0,000958 | 0,907828 | 0,0836 | 0,248138 | 0,031443 | 0,608286 | 0,27836256 | 0 | 0,33111901 | 0,951195 | | 0,991428 |
| GSM2262021 | 0 | 0,759483 | 0 | 0,519337 | 0 | 1,470739 | 0,23743243 | 0 | 0,34900451 | 0,947159 | | 0,759483 |
| GSM2262022 | 0,004588 | 0,677827 | 0 | 0,496476 | 0 | 0,692439 | 0,32882115 | 0 | 0,37482161 | 0,930949 | | 0,677827 |
| GSM2262032 | 0,026354 | 1,049207 | 0 | 0,749823 | 0 | 2,099025 | 0,14478334 | 0 | 0,50308616 | 0,863571 | | 1,049207 |
| GSM2262036 | 0 | 1,250748 | 0 | 0,526154 | 0 | 3,028818 | 0,22153871 | 0 | 0,55083412 | 0,834691 | | 1,250748 |
| GSM2262038 | 0 | 1,349483 | 0,115249 | 0,317465 | 0 | 2,097675 | 0,34780416 | 0 | 0,49098025 | 0,871788 | | 1,464732 |
| GSM2262039 | 0 | 0,995612 | 0 | 0,416254 | 0 | 1,833543 | 0,03465268 | 0 | 0,55242507 | 0,835233 | | 0,995612 |
| GSM2262048 | 0 | 0,958005 | 0 | 0,535129 | 0 | 1,102331 | 0,21223838 | 0 | 0,31424776 | 0,960783 | | 0,958005 |
| GSM2262056 | 0 | 1,146684 | 0 | 0,279468 | 0 | 1,215517 | 0,26458846 | 0 | 0,38907686 | 0,924869 | | 1,146684 |
| GSM2262058 | 0 | 0,531186 | 0,136686 | 0,231234 | 0 | 0,766053 | 0,13638119 | 0 | 0,57272816 | 0,82664 | | 0,667872 |
| GSM2262065 | 0 | 0,650296 | 0,297486 | 0,34974 | 0 | 1,710885 | 0,06907849 | 0 | 0,37927308 | 0,932938 | | 0,947782 |
| GSM2262070 | 0 | 1,212715 | 0 | 0,445818 | 0 | 1,483748 | 0,21014265 | 0 | 0,36694828 | 0,935705 | | 1,212715 |
| GSM2262072 | 0 | 0,820588 | 0 | 0,519357 | 0,046802 | 1,072585 | 0,32010493 | 0 | 0,43615936 | 0,899654 | | 0,820588 |
| GSM2262075 | 0 | 1,912695 | 0,260912 | 0,068542 | 0,275956 | 2,641432 | 0 | 0 | 0,29952117 | 0,971566 | | 2,173607 |
| GSM2262079 | 0 | 0,638508 | 0 | 0,189849 | 0,005172 | 0,77134 | 0,11944191 | 0 | 0,44747345 | 0,895152 | | 0,638508 |
| GSM2262087 | 0 | 1,005696 | 0 | 0,103098 | 0,31089 | 1,045519 | 0,43197169 | 0 | 0,35072877 | 0,942669 | | 1,005696 |
| GSM2262089 | 0 | 0,661705 | 0,125619 | 0,600882 | 0,006604 | 1,102598 | 0,57329652 | 0 | 0,49701194 | 0,866783 | | 0,787323 |
| GSM2262090 | 0,017908 | 0,823457 | 0 | 0,283226 | 0 | 0,874753 | 0,25134622 | 0 | 0,38147077 | 0,927626 | | 0,823457 |
| GSM2262092 | 0,013775 | 0,747488 | 0 | 0,251143 | 0 | 0,604767 | 0,27290348 | 0 | 0,41234104 | 0,912428 | | 0,747488 |
| GSM2262097 | 0 | 0,34942 | 0,379412 | 0,33019 | 0 | 0,63471 | 0,17418837 | 0 | 0,36910711 | 0,935101 | | 0,728833 |
| GSM2262101 | 0 | 0,982754 | 0 | 0,341621 | 0 | 0,78823 | 0,43208873 | 0 | 0,35134382 | 0,941912 | | 0,982754 |
| GSM2262108 | 0 | 1,178256 | 0,10314 | 0,526989 | 0 | 1,944452 | 0,14654041 | 0 | 0,35792528 | 0,941664 | | 1,281397 |
| GSM2262109 | 0 | 0,760781 | 0,133084 | 0,325636 | 0,022702 | 1,381159 | 0,28695128 | 0 | 0,44938375 | 0,893473 | | 0,893865 |
| GSM2262110 | 0 | 0,74768 | 0,200479 | 0,401975 | 0 | 1,886812 | 0,60508543 | 0 | 0,53869839 | 0,842347 | | 0,948159 |
| GSM2262112 | 0 | 0,732562 | 0,130965 | 0,585406 | 0,037563 | 1,687517 | 0,10193246 | 0 | 0,53906546 | 0,842145 | | 0,863528 |
| GSM2262114 | 0 | 1,190118 | 0,020033 | 0,412493 | 0,042187 | 1,858811 | 0,29218337 | 0 | 0,45373657 | 0,891517 | | 1,210151 |
| GSM2262118 | 0 | 0,919582 | 0 | 0,151667 | 0,014088 | 1,310743 | 0,07509043 | 0 | 0,45289192 | 0,893443 | | 0,919582 |
| GSM2262125 | 0 | 1,085165 | 0,162611 | 0,383747 | 0 | 0,993265 | 0,51981927 | 0 | 0,3519087 | 0,941429 | | 1,247775 |
| GSM2262129 | 0 | 0,850209 | 0 | 0,260403 | 0 | 1,274697 | 0,04781145 | 0 | 0,35578173 | 0,943992 | | 0,850209 |
| GSM2262133 | 0 | 0,955244 | 0 | 0,225952 | 0 | 1,565391 | 0,04526468 | 0 | 0,41784672 | 0,912948 | | 0,955244 |
| GSM2262134 | 0 | 1,235243 | 0 | 0,227137 | 0,017565 | 1,788005 | 0 | 0 | 0,47432971 | 0,881691 | | 1,235243 |
| GSM2262137 | 0 | 0,707932 | 0 | 0,202633 | 0,008676 | 0,542596 | 0,13096743 | 0 | 0,37824909 | 0,929465 | | 0,707932 |
| GSM2262140 | 0 | 0,845804 | 0,16133 | 0,375642 | 0,042063 | 1,384733 | 0,15425443 | 0 | 0,38922693 | 0,92452 | | 1,007134 |
| GSM2262141 | 0 | 0,852739 | 0 | 0,278834 | 0 | 0,806605 | 0,11492774 | 0 | 0,35143344 | 0,942707 | | 0,852739 |
| GSM2262143 | 0 | 1,403347 | 0 | 0,422307 | 0 | 1,842722 | 0,076432 | 0 | 0,37528144 | 0,932895 | | 1,403347 |
| GSM2262150 | 0 | 1,114516 | 0 | 0,282204 | 0 | 2,79402 | 0 | 0 | 0,59329324 | 0,806249 | | 1,114516 |
| GSM2262153 | 0 | 1,070543 | 0 | 0,262887 | 0,006356 | 0,966172 | 0,23415029 | 0 | 0,46445417 | 0,887104 | | 1,070543 |
| GSM2262155 | 0 | 1,244164 | 0 | 0,438664 | 0 | 4,82506 | 0 | 0 | 0,70352646 | 0,717157 | | 1,244164 |
| GSM2262157 | 0 | 1,363099 | 0 | 0,367217 | 0,084631 | 3,179864 | 0 | 0 | 0,64168576 | 0,77414 | | 1,363099 |
| GSM2262158 | 0 | 1,554363 | 0 | 0,143272 | 0,082266 | 1,202755 | 0,27378838 | 0 | 0,39058504 | 0,924293 | | 1,554363 |
| GSM2262184 | 0 | 0,87477 | 0,005276 | 0,334323 | 0,051276 | 1,344994 | 0,33549699 | 0 | 0,35193851 | 0,944067 | | 0,880046 |
| GSM2262192 | 0 | 1,16285 | 0 | 0,620187 | 0 | 0,978185 | 0,45134215 | 0 | 0,3355429 | 0,949626 | | 1,16285 |
| GSM2262195 | 0,172784 | 1,172879 | 0,561706 | 0,790017 | 0,050557 | 5,856651 | 0,44705126 | 0 | 0,64579358 | 0,767701 | | 1,734584 |
| GSM2262204 | 0 | 0,526835 | 0 | 0,226999 | 0 | 0,625282 | 0,13443858 | 0 | 0,29909467 | 0,967972 | | 0,526835 |
| GSM2262205 | 0 | 1,03365 | 0 | 0,260142 | 0 | 1,470775 | 0,18492508 | 0 | 0,34607545 | 0,948508 | | 1,03365 |
| GSM2262208 | 0 | 1,256061 | 0 | 0,447 | 0 | 3,246426 | 0,07604626 | 0 | 0,57115929 | 0,821183 | | 1,256061 |
| GSM2262209 | 0 | 0,757239 | 0 | 0,291672 | 0 | 1,14931 | 0,13784987 | 0 | 0,38763438 | 0,926527 | | 0,757239 |
| GSM2262229 | 0,008501 | 0,887673 | 0 | 0,270825 | 0,055541 | 1,608551 | 0,08225327 | 0 | 0,57076881 | 0,824349 | | 0,887673 |
| GSM2262249 | 0 | 0,86659 | 0 | 0,576202 | 0 | 0,672358 | 0,1991398 | 0 | 0,41412268 | 0,911917 | | 0,86659 |
| GSM2262254 | 0 | 1,115841 | 0 | 0,318524 | 0 | 1,548903 | 0 | 0 | 0,52354464 | 0,85393 | | 1,115841 |
| GSM2262259 | 0 | 0,512267 | 0 | 0,360018 | 0 | 0,885032 | 0,15726518 | 0 | 0,43056275 | 0,903364 | | 0,512267 |
| GSM2262261 | 0 | 1,006114 | 0 | 0,677334 | 0 | 0,98195 | 0,21334194 | 0 | 0,33967666 | 0,949187 | | 1,006114 |
| GSM2262263 | 0 | 0,754754 | 0 | 0,28085 | 0 | 0,612956 | 0,27286095 | 0 | 0,3354668 | 0,949338 | | 0,754754 |
| GSM2262275 | 0 | 1,189971 | 0 | 0,487097 | 0,023396 | 1,142997 | 0,78939011 | 0 | 0,46023722 | 0,887854 | | 1,189971 |
| GSM2262291 | 0 | 1,66199 | 0 | 0,621559 | 0 | 4,38886 | 0 | 0 | 0,58703278 | 0,810373 | | 1,66199 |
| GSM2262294 | 0 | 1,215057 | 0 | 0,291059 | 0,01045 | 1,127128 | 0,25373771 | 0 | 0,40216684 | 0,918126 | | 1,215057 |
| GSM2262296 | 0 | 0,793401 | 0,272149 | 0,748245 | 0,090896 | 1,914591 | 0,09894943 | 0 | 0,37084481 | 0,935945 | | 1,065551 |
| GSM2262299 | 0 | 0,618097 | 0,003459 | 0,212173 | 0 | 0,927588 | 0,06684887 | 0 | 0,50902471 | 0,861627 | | 0,621555 |
| GSM2262300 | 0 | 1,012165 | 0 | 0,408876 | 0,003233 | 1,012478 | 0,06529223 | 0 | 0,36899064 | 0,934529 | | 1,012165 |
| GSM2262301 | 0 | 1,335194 | 0 | 0,754898 | 0 | 1,358231 | 0,16110154 | 0 | 0,38570807 | 0,926084 | | 1,335194 |
| GSM2262308 | 0 | 1,429598 | 0 | 0,159428 | 0,030024 | 2,748268 | 0 | 0 | 0,59107999 | 0,809812 | | 1,429598 |
| average | 0,003374 | 0,99284 | 0,059111 | 0,400266 | 0,018426 | 1,786536 | 0,19877291 | | | | | |

**CIBERSORTx_GSE85217_GP4**

| Sample | Bcells | TCD4 | TCD8 | Tgd | NK | MoMaDC | granulocytes | P-value | Correlation | RMSE | ratio mono/T cells |
|---|---|---|---|---|---|---|---|---|---|---|---|
| GSM2261540 | 0 | 0,27726 | 0 | 0,263684 | 0 | 0,38037 | 0,07868532 | 0 | 0,3598455 | 0,940333 | 0,70315936 |
| GSM2261541 | 0 | 0,179393 | 0 | 0,217109 | 0 | 0,425363 | 0,1781353 | 0 | 0,4104143 | 0,915444 | 1,072792035 |
| GSM2261542 | 0 | 0,168176 | 0,002903 | 0,186812 | 0 | 0,422746 | 0,21936327 | 0 | 0,4269089 | 0,907196 | 1,181215112 |
| GSM2261546 | 0 | 0,235968 | 0 | 0,255653 | 0 | 0,420279 | 0,08810009 | 0 | 0,3801833 | 0,930376 | 0,854885698 |
| GSM2261556 | 0 | 0,264958 | 0 | 0,249282 | 0 | 0,349748 | 0,13601197 | 0 | 0,3766431 | 0,930994 | 0,680125843 |
| GSM2261557 | 0 | 0,407516 | 0 | 0,198181 | 0 | 0,281825 | 0,11247922 | 0 | 0,3893759 | 0,923897 | 0,465290149 |
| GSM2261558 | 0 | 0,304291 | 0 | 0,141177 | 0 | 0,554532 | 0 | 0 | 0,6588625 | 0,769132 | 1,244830504 |
| GSM2261562 | 0 | 0,266985 | 0,019836 | 0,165446 | 0 | 0,547733 | 0 | 0 | 0,5350412 | 0,844935 | 1,211083682 |
| GSM2261563 | 0 | 0,280191 | 0 | 0,184658 | 0 | 0,483421 | 0,05173 | 0 | 0,4372366 | 0,899988 | 1,039953419 |
| GSM2261564 | 0 | 0,212477 | 0,005913 | 0,141141 | 0 | 0,640468 | 0 | 0 | 0,6266741 | 0,783042 | 1,781395926 |
| GSM2261572 | 0 | 0,350923 | 0 | 0,184079 | 0 | 0,418305 | 0,04669271 | 0 | 0,3896014 | 0,924075 | 0,781875264 |
| GSM2261573 | 0 | 0,173914 | 0 | 0,12285 | 0 | 0,703236 | 0 | 0 | 0,6157798 | 0,787883 | 2,369682741 |
| GSM2261575 | 0 | 0,290185 | 0 | 0,176239 | 0 | 0,527445 | 0,00613139 | 0 | 0,4345432 | 0,902119 | 1,13082824 |
| GSM2261576 | 0 | 0,259546 | 0 | 0,213155 | 0 | 0,508087 | 0,01921137 | 0 | 0,5068694 | 0,861159 | 1,074857504 |
| GSM2261578 | 0 | 0,304768 | 0 | 0,225437 | 0 | 0,462326 | 0,00746912 | 0 | 0,4076931 | 0,915829 | 0,87197593 |
| GSM2261579 | 0 | 0,105618 | 0,032041 | 0,094561 | 0,006141 | 0,573216 | 0,1884236 | 0 | 0,6514869 | 0,761268 | 2,46841736 |
| GSM2261582 | 0 | 0,261627 | 0 | 0,182131 | 0 | 0,538031 | 0,01821106 | 0 | 0,5469887 | 0,837689 | 1,212442892 |
| GSM2261583 | 0 | 0,310098 | 0 | 0,120121 | 0 | 0,541916 | 0,02786477 | 0 | 0,4859709 | 0,874145 | 1,259630033 |
| GSM2261584 | 0 | 0,272433 | 0 | 0,229689 | 0 | 0,430672 | 0,06720567 | 0 | 0,3796134 | 0,929945 | 0,857703357 |
| GSM2261585 | 0 | 0,296199 | 0 | 0,162922 | 0 | 0,535365 | 0,00551363 | 0 | 0,3937326 | 0,924369 | 1,166064948 |
| GSM2261587 | 0 | 0,299795 | 0 | 0,256906 | 0 | 0,295501 | 0,1477975 | 0 | 0,388295 | 0,924851 | 0,530808104 |
| GSM2261588 | 0 | 0,390243 | 0 | 0,127737 | 0 | 0,48202 | 0 | 0 | 0,4135607 | 0,913127 | 0,930575318 |
| GSM2261595 | 0 | 0,335047 | 0 | 0,255337 | 0 | 0,280324 | 0,12929144 | 0 | 0,4410479 | 0,897451 | 0,4748163 |
| GSM2261596 | 0 | 0,278191 | 0 | 0,227698 | 0 | 0,438977 | 0,0551342 | 0 | 0,4368451 | 0,899841 | 0,867733927 |
| GSM2261599 | 0 | 0,340853 | 0 | 0,186719 | 0 | 0,436411 | 0,0360167 | 0 | 0,4039992 | 0,91698 | 0,827207495 |
| GSM2261606 | 0 | 0,188391 | 0 | 0,128415 | 0 | 0,584332 | 0,09886272 | 0 | 0,4950276 | 0,86927 | 1,844450236 |
| GSM2261607 | 0 | 0,360358 | 0 | 0,228561 | 0 | 0,344775 | 0,06630607 | 0 | 0,3578219 | 0,940093 | 0,585437814 |
| GSM2261608 | 0 | 0,313662 | 0 | 0,210036 | 0 | 0,423632 | 0,0526694 | 0 | 0,3775117 | 0,930466 | 0,808924901 |
| GSM2261610 | 0 | 0,276236 | 0 | 0,210787 | 0 | 0,447147 | 0,06582966 | 0 | 0,3560214 | 0,942051 | 0,91812317 |
| GSM2261611 | 0 | 0,32252 | 0 | 0,230605 | 0 | 0,414327 | 0,03254849 | 0 | 0,3679554 | 0,935781 | 0,749066645 |
| GSM2261616 | 0 | 0,30366 | 0 | 0,179855 | 0 | 0,510172 | 0,00631338 | 0 | 0,4352075 | 0,901486 | 1,055130455 |
| GSM2261618 | 0 | 0,241079 | 0 | 0,254207 | 0 | 0,372835 | 0,13187843 | 0 | 0,3724885 | 0,933728 | 0,752767069 |
| GSM2261621 | 0 | 0,31865 | 0 | 0,207004 | 0,00312 | 0,364613 | 0,10661438 | 0 | 0,3856225 | 0,925553 | 0,693636727 |
| GSM2261623 | 0 | 0,198845 | 0 | 0,236529 | 0 | 0,52124 | 0,0433861 | 0 | 0,492003 | 0,869538 | 1,1972247 |
| GSM2261626 | 0 | 0,337175 | 0 | 0,138798 | 0 | 0,461916 | 0,06211128 | 0 | 0,3354146 | 0,951844 | 0,970465924 |
| GSM2261627 | 0 | 0,250447 | 0 | 0,255934 | 0 | 0,326559 | 0,16706088 | 0 | 0,449478 | 0,892751 | 0,644887453 |
| GSM2261630 | 0 | 0,255471 | 0 | 0,152551 | 0 | 0,584466 | 0,00751131 | 0 | 0,4841993 | 0,874958 | 1,432436912 |
| GSM2261631 | 0 | 0,219391 | 0 | 0,231496 | 0 | 0,526874 | 0,02223955 | 0 | 0,5002503 | 0,86473 | 1,168527671 |
| GSM2261632 | 0 | 0,145286 | 6,19E-06 | 0,260498 | 0 | 0,504279 | 0,08993165 | 0 | 0,4272002 | 0,907951 | 1,242710235 |
| GSM2261637 | 0 | 0,23433 | 0,018645 | 0,19879 | 0 | 0,473521 | 0,07471324 | 0 | 0,511416 | 0,858591 | 1,048157931 |
| GSM2261638 | 0 | 0,294337 | 0 | 0,186767 | 0 | 0,438067 | 0,08082848 | 0 | 0,3925381 | 0,922823 | 0,910543979 |
| GSM2261639 | 0 | 0,184705 | 0,085486 | 0,157643 | 0 | 0,571479 | 0,00068637 | 0 | 0,6538752 | 0,7682 | 1,335747659 |
| GSM2261640 | 0 | 0,265551 | 0 | 0,1522 | 0 | 0,51917 | 0,06307851 | 0 | 0,5580011 | 0,831716 | 1,242772861 |
| GSM2261651 | 0 | 0,324325 | 0,012009 | 0,158934 | 0 | 0,378054 | 0,12667767 | 0 | 0,3968833 | 0,919872 | 0,763332727 |
| GSM2261652 | 0 | 0,212673 | 0 | 0,105322 | 0 | 0,682005 | 0 | 0 | 0,7052547 | 0,722208 | 2,144707568 |
| GSM2261653 | 0 | 0,278143 | 0,020418 | 0,18224 | 0 | 0,445936 | 0,07326277 | 0 | 0,5057664 | 0,862532 | 0,927485768 |
| GSM2261654 | 0 | 0,180159 | 0,038839 | 0,163172 | 0 | 0,617831 | 0 | 0 | 0,6413231 | 0,77349 | 1,61664079 |
| GSM2261656 | 0 | 0,236711 | 0 | 0,182708 | 0 | 0,523536 | 0,05704427 | 0 | 0,5801119 | 0,817396 | 1,248241035 |
| GSM2261659 | 0 | 0,353784 | 0 | 0,220095 | 0 | 0,389762 | 0,03635946 | 0 | 0,42779 | 0,904638 | 0,679170132 |
| GSM2261660 | 0 | 0,322675 | 0 | 0,198358 | 0 | 0,413332 | 0,06563485 | 0 | 0,4048685 | 0,916221 | 0,793292863 |
| GSM2261662 | 0 | 0,214831 | 0 | 0,181189 | 0,001667 | 0,551565 | 0,05074703 | 0 | 0,7135303 | 0,727436 | 1,392767132 |
| GSM2261663 | 0 | 0,233546 | 0 | 0,214968 | 0 | 0,45646 | 0,09502633 | 0 | 0,4987043 | 0,865737 | 1,017715021 |
| GSM2261664 | 0 | 0,171605 | 0 | 0,145255 | 0 | 0,61701 | 0,06612958 | 0 | 0,6738141 | 0,748159 | 1,947264912 |
| GSM2261665 | 0 | 0,2403 | 0,000606 | 0,21308 | 0 | 0,492264 | 0,0537501 | 0 | 0,4691196 | 0,882519 | 1,084316011 |
| GSM2261666 | 0 | 0,329028 | 0 | 0,212916 | 0 | 0,335798 | 0,12225843 | 0 | 0,3595558 | 0,938431 | 0,619618108 |
| GSM2261669 | 0 | 0,290617 | 0 | 0,103056 | 0 | 0,597596 | 0,00873081 | 0 | 0,5511342 | 0,835262 | 1,51799931 |
| GSM2261670 | 0 | 0,30349 | 0 | 0,180355 | 0 | 0,378002 | 0,13815255 | 0 | 0,4537057 | 0,890812 | 0,781245236 |
| GSM2261671 | 0 | 0,169339 | 0,019772 | 0,213375 | 0 | 0,571845 | 0,02566903 | 0 | 0,5526488 | 0,832444 | 1,420781891 |
| GSM2261672 | 0 | 0,311506 | 0 | 0,183534 | 0 | 0,382938 | 0,12202214 | 0 | 0,4269737 | 0,904653 | 0,773548514 |
| GSM2261677 | 0 | 0,227688 | 0,04337 | 0,157617 | 0 | 0,539787 | 0,03153786 | 0 | 0,5355867 | 0,844321 | 1,259200376 |
| GSM2261679 | 0 | 0,253968 | 0 | 0,101367 | 0,004124 | 0,639635 | 0,00090556 | 0 | 0,6049073 | 0,798832 | 1,800088836 |
| GSM2261681 | 0 | 0,281557 | 0 | 0,230853 | 0 | 0,419371 | 0,06821942 | 0 | 0,3932821 | 0,922649 | 0,818429844 |
| GSM2261682 | 0 | 0,256667 | 0,028766 | 0,126203 | 0 | 0,573788 | 0,01457583 | 0 | 0,5997248 | 0,804975 | 1,393918919 |
| GSM2261683 | 0 | 0,166866 | 0 | 0,129997 | 0 | 0,703138 | 0 | 0 | 0,6962773 | 0,725957 | 2,368564555 |
| GSM2261684 | 0 | 0,202468 | 0,043942 | 0,17553 | 0 | 0,575525 | 0,00253472 | 0 | 0,6445191 | 0,774085 | 1,363997169 |
| GSM2261685 | 0 | 0,229151 | 0 | 0,183043 | 0 | 0,547167 | 0,0406391 | 0 | 0,5241072 | 0,850864 | 1,327450558 |
| GSM2261686 | 0 | 0,263734 | 0 | 0,200528 | 0 | 0,489125 | 0,04661276 | 0 | 0,5058077 | 0,861965 | 1,053553221 |
| GSM2261688 | 0 | 0,121389 | 0,069377 | 0,154607 | 0 | 0,601324 | 0,05330155 | 0 | 0,5893368 | 0,807998 | 1,741081032 |
| GSM2261690 | 0 | 0,252455 | 0,001617 | 0,137122 | 0 | 0,608806 | 0 | 0 | 0,5850128 | 0,812855 | 1,556273932 |
| GSM2261692 | 0 | 0,186729 | 0,002348 | 0,105929 | 0 | 0,704994 | 0 | 0 | 0,6991991 | 0,724041 | 2,389757818 |
| GSM2261693 | 0 | 0,141646 | 0,10301 | 0,211787 | 0 | 0,504001 | 0,03955683 | 0 | 0,5339336 | 0,844375 | 1,104193117 |
| GSM2261694 | 0 | 0,301802 | 0 | 0,210356 | 0 | 0,443718 | 0,04412456 | 0 | 0,4341332 | 0,901299 | 0,866370585 |
| GSM2261695 | 0 | 0,243059 | 0,007392 | 0,186705 | 0 | 0,512339 | 0,05050431 | 0 | 0,4891955 | 0,871338 | 1,17198261 |
| GSM2261696 | 0 | 0,275437 | 0,027148 | 0,158914 | 0,000535 | 0,422254 | 0,11571123 | 0 | 0,4059575 | 0,915687 | 0,914962002 |
| GSM2261697 | 0 | 0,316049 | 0 | 0,187818 | 0 | 0,418748 | 0,07738469 | 0 | 0,4349284 | 0,900711 | 0,8310696 |
| GSM2261698 | 0 | 0,313377 | 0 | 0,195287 | 0 | 0,462193 | 0,02914217 | 0 | 0,4241211 | 0,906782 | 0,908639926 |
| GSM2261699 | 0 | 0,283347 | 0 | 0,193629 | 0,003782 | 0,459269 | 0,05997383 | 0 | 0,5430439 | 0,841313 | 0,962876793 |
| GSM2261703 | 0 | 0,303872 | 0 | 0,237496 | 0 | 0,350315 | 0,10831568 | 0 | 0,4079494 | 0,914498 | 0,647091841 |
| GSM2261704 | 0 | 0,295944 | 0,024253 | 0,199411 | 0 | 0,43796 | 0,04243248 | 0 | 0,4794055 | 0,876952 | 0,84286858 |
| GSM2261705 | 0 | 0,230286 | 0,015264 | 0,149645 | 0 | 0,511258 | 0,09354692 | 0 | 0,6072588 | 0,800972 | 1,293684511 |

| | | | | | | | | | | | |
|---|---|---|---|---|---|---|---|---|---|---|---|
| GSM2261714 | 0 | 0,301672 | 0,035222 | 0,140783 | 0 | 0,510911 | 0,01141261 | 0 | 0,5729923 | 0,824176 | 1,069574549 |
| GSM2261721 | 0 | 0,267316 | 0,016096 | 0,158674 | 0 | 0,514586 | 0,04332842 | 0 | 0,5950066 | 0,809842 | 1,163995882 |
| GSM2261723 | 0 | 0,174025 | 0,005028 | 0,083833 | 0 | 0,714783 | 0,02233068 | 0 | 0,6421687 | 0,767491 | 2,718979647 |
| GSM2261724 | 0 | 0,356058 | 0 | 0,177024 | 0 | 0,387076 | 0,07984197 | 0 | 0,5032948 | 0,865261 | 0,726109867 |
| GSM2261727 | 0 | 0,313414 | 5,67E-05 | 0,266491 | 0 | 0,293115 | 0,12692296 | 0 | 0,4267358 | 0,905036 | 0,505403421 |
| GSM2261728 | 0 | 0,243769 | 0 | 0,170537 | 0 | 0,542603 | 0,04309063 | 0 | 0,5378819 | 0,842901 | 1,30966513 |
| GSM2261730 | 0 | 0,321327 | 0 | 0,175868 | 0 | 0,376829 | 0,12597548 | 0 | 0,4678298 | 0,883597 | 0,757910169 |
| GSM2261731 | 0 | 0,190721 | 0 | 0,161581 | 0 | 0,362547 | 0,28515187 | 0 | 0,5493692 | 0,835455 | 1,029082115 |
| GSM2261733 | 0 | 0,186502 | 0,003747 | 0,194172 | 0 | 0,430415 | 0,18516346 | 0 | 0,5461101 | 0,837619 | 1,119641187 |
| GSM2261736 | 0 | 0,258267 | 0 | 0,204536 | 0 | 0,48599 | 0,05120726 | 0 | 0,4419507 | 0,897488 | 1,050101639 |
| GSM2261740 | 0 | 0,24113 | 0 | 0,197405 | 0,004721 | 0,471093 | 0,08565056 | 0 | 0,4830352 | 0,874637 | 1,074243673 |
| GSM2261744 | 0 | 0,220601 | 0,009968 | 0,22128 | 0 | 0,432399 | 0,11575188 | 0 | 0,4934046 | 0,868633 | 0,95695415 |
| GSM2261749 | 0 | 0,181875 | 0 | 0,256141 | 0 | 0,505798 | 0,05618667 | 0 | 0,5577714 | 0,829592 | 1,154747376 |
| GSM2261751 | 0 | 0,214145 | 0 | 0,158395 | 0 | 0,560753 | 0,06670757 | 0 | 0,5897116 | 0,809881 | 1,505218452 |
| GSM2261754 | 0 | 0,304529 | 0 | 0,198658 | 0 | 0,359009 | 0,13780381 | 0 | 0,5313333 | 0,849214 | 0,713469132 |
| GSM2261755 | 0 | 0,286287 | 0 | 0,176444 | 0 | 0,458152 | 0,07911689 | 0 | 0,4600442 | 0,887565 | 0,990104403 |
| GSM2261759 | 0 | 0,295741 | 0 | 0,085172 | 0 | 0,619087 | 0 | 0 | 0,5628665 | 0,827744 | 1,62526994 |
| GSM2261761 | 0 | 0,288383 | 0,006525 | 0,220263 | 0 | 0,330609 | 0,15422046 | 0 | 0,3553483 | 0,940864 | 0,641745608 |
| GSM2261764 | 0 | 0,269221 | 0 | 0,246833 | 0 | 0,349841 | 0,13410458 | 0 | 0,483694 | 0,874215 | 0,677914722 |
| GSM2261767 | 0 | 0,229803 | 0 | 0,161292 | 0 | 0,51961 | 0,08929414 | 0 | 0,6105078 | 0,798304 | 1,328601483 |
| GSM2261769 | 0 | 0,218222 | 0,002203 | 0,259933 | 0 | 0,377447 | 0,14219555 | 0 | 0,5389635 | 0,842323 | 0,785762483 |
| GSM2261770 | 0 | 0,254398 | 0 | 0,244557 | 0 | 0,385704 | 0,11534072 | 0 | 0,5190406 | 0,854416 | 0,773023213 |
| GSM2261772 | 0 | 0,306367 | 0 | 0,175055 | 0 | 0,384804 | 0,13377317 | 0 | 0,4421181 | 0,896892 | 0,799307388 |
| GSM2261773 | 0 | 0,263644 | 0 | 0,16509 | 0 | 0,48847 | 0,08279553 | 0 | 0,5601046 | 0,830906 | 1,139332165 |
| GSM2261774 | 0 | 0,291024 | 0 | 0,192691 | 0 | 0,407418 | 0,10886738 | 0 | 0,5008167 | 0,865473 | 0,842270435 |
| GSM2261777 | 0 | 0,209682 | 0,009464 | 0,104947 | 0,003305 | 0,631221 | 0,04138087 | 0 | 0,6777107 | 0,74644 | 1,947656999 |
| GSM2261778 | 0 | 0,305229 | 0 | 0,233224 | 0 | 0,331838 | 0,12970919 | 0 | 0,4872521 | 0,872683 | 0,616279585 |
| GSM2261779 | 0 | 0,295451 | 0 | 0,160696 | 0 | 0,494141 | 0,04971245 | 0 | 0,4701671 | 0,882343 | 1,083295331 |
| GSM2261781 | 0 | 0,330998 | 0,006205 | 0,200551 | 0 | 0,390872 | 0,07137498 | 0 | 0,396338 | 0,920384 | 0,726860243 |
| GSM2261783 | 0 | 0,15623 | 0 | 0,147128 | 0 | 0,696642 | 0 | 0 | 0,6726576 | 0,744748 | 2,296440418 |
| GSM2261784 | 0 | 0,370838 | 0 | 0,199636 | 0 | 0,262103 | 0,1674232 | 0 | 0,3690813 | 0,933556 | 0,459447576 |
| GSM2261785 | 0 | 0,264357 | 0,081046 | 0,14034 | 0 | 0,491408 | 0,02284897 | 0 | 0,5882136 | 0,815089 | 1,011662562 |
| GSM2261786 | 0 | 0,169214 | 0 | 0,150485 | 0 | 0,502611 | 0,17768964 | 0 | 0,591802 | 0,807488 | 1,572138507 |
| GSM2261787 | 0 | 0,145037 | 0,119322 | 0,128177 | 0 | 0,583389 | 0,02407509 | 0 | 0,6046154 | 0,799569 | 1,48620415 |
| GSM2261788 | 0 | 0,155889 | 0,061459 | 0,17901 | 0 | 0,433198 | 0,17044515 | 0 | 0,5807227 | 0,816847 | 1,092948517 |
| GSM2261789 | 0 | 0,201085 | 0,058524 | 0,175028 | 0 | 0,433205 | 0,13215792 | 0 | 0,4291186 | 0,903895 | 0,996703531 |
| GSM2261790 | 0 | 0,240024 | 0,045655 | 0,169905 | 0 | 0,434351 | 0,11006519 | 0 | 0,499551 | 0,865706 | 0,953393321 |
| GSM2261792 | 0 | 0,309278 | 0 | 0,137388 | 0,002117 | 0,45601 | 0,09520729 | 0 | 0,3692487 | 0,935104 | 1,02091888 |
| GSM2261795 | 0 | 0,162379 | 0 | 0,205601 | 0 | 0,487675 | 0,14434486 | 0 | 0,5952146 | 0,805929 | 1,325274365 |
| GSM2261796 | 0 | 0,311401 | 0 | 0,199956 | 0 | 0,441604 | 0,04703951 | 0 | 0,4505615 | 0,892581 | 0,863592987 |
| GSM2261799 | 0 | 0,223163 | 0 | 0,219461 | 0 | 0,478049 | 0,07932793 | 0 | 0,5656993 | 0,826299 | 1,080034207 |
| GSM2261803 | 0 | 0,229349 | 0 | 0,183324 | 0 | 0,565446 | 0,02188076 | 0 | 0,5282532 | 0,848289 | 1,370205108 |
| GSM2261804 | 0 | 0,155076 | 0,015687 | 0,288979 | 0 | 0,407816 | 0,13244202 | 0 | 0,6182628 | 0,791454 | 0,887055855 |
| GSM2261806 | 0 | 0,264113 | 0,001273 | 0,123974 | 0 | 0,567318 | 0,04332206 | 0 | 0,5520775 | 0,834553 | 1,457053429 |
| GSM2261808 | 0 | 0,34549 | 0 | 0,266431 | 0 | 0,328716 | 0,05936347 | 0 | 0,3963517 | 0,921545 | 0,537186914 |
| GSM2261809 | 0 | 0,2078 | 0 | 0,083477 | 0,004284 | 0,704439 | 0 | 0 | 0,6142862 | 0,789484 | 2,418452008 |
| GSM2261810 | 0 | 0,247246 | 0,045584 | 0,182168 | 0 | 0,37313 | 0,15187143 | 0 | 0,4766003 | 0,878267 | 0,785540731 |
| GSM2261811 | 0 | 0,196617 | 0,06281 | 0,166699 | 0 | 0,478272 | 0,09560119 | 0 | 0,5296509 | 0,847947 | 1,122369889 |
| GSM2261813 | 0 | 0,24954 | 0 | 0,124452 | 0 | 0,435271 | 0,19073633 | 0 | 0,3513063 | 0,946729 | 1,163851025 |
| GSM2261814 | 0 | 0,177622 | 0,012343 | 0,196836 | 0 | 0,404957 | 0,20824197 | 0 | 0,5663551 | 0,825365 | 1,046939207 |
| GSM2261819 | 0 | 0,341625 | 0 | 0,173066 | 0 | 0,410951 | 0,07435829 | 0 | 0,3695659 | 0,933831 | 0,798441306 |
| GSM2261825 | 0 | 0,300007 | 0 | 0,179363 | 0,004848 | 0,492146 | 0,0236361 | 0 | 0,504717 | 0,863006 | 1,026651901 |
| GSM2261827 | 0 | 0,101841 | 0,055497 | 0,106906 | 0 | 0,418874 | 0,31688233 | 0 | 0,615351 | 0,78935 | 1,585182994 |
| GSM2261829 | 0 | 0,24916 | 0,000275 | 0,13954 | 0 | 0,609617 | 0,00140933 | 0 | 0,5873989 | 0,811199 | 1,567242438 |
| GSM2261831 | 0 | 0,28386 | 0 | 0,107198 | 0 | 0,549298 | 0,05964384 | 0 | 0,5001605 | 0,865971 | 1,404646123 |
| GSM2261833 | 0 | 0,198227 | 0 | 0,18097 | 0 | 0,60554 | 0,01526232 | 0 | 0,6095571 | 0,795336 | 1,596896741 |
| GSM2261835 | 0 | 0,361367 | 0 | 0,204179 | 0 | 0,304053 | 0,13040102 | 0 | 0,4044414 | 0,916024 | 0,537627715 |
| GSM2261836 | 0 | 0,302002 | 0,009204 | 0,124192 | 0 | 0,564602 | 0 | 0 | 0,5172621 | 0,855992 | 1,296750384 |
| GSM2261838 | 0 | 0,216889 | 0 | 0,311885 | 0 | 0,344815 | 0,12640999 | 0 | 0,4601186 | 0,887492 | 0,652103097 |
| GSM2261846 | 0 | 0,311447 | 0,015532 | 0,232138 | 0 | 0,40332 | 0,03756301 | 0 | 0,5405976 | 0,842796 | 0,72135113 |
| GSM2261847 | 0 | 0,22528 | 0,015992 | 0,214877 | 0 | 0,542674 | 0,00117695 | 0,02 | 0,2070456 | 1,023101 | 1,189686162 |
| GSM2261848 | 0 | 0,20118 | 0 | 0,179209 | 0 | 0,583538 | 0,03607348 | 0 | 0,5425085 | 0,839174 | 1,534056041 |
| GSM2261852 | 0 | 0,184791 | 0,018215 | 0,183617 | 0 | 0,51183 | 0,1015469 | 0 | 0,3172743 | 0,966598 | 1,323846309 |
| GSM2261854 | 0 | 0,278225 | 0 | 0,165609 | 0 | 0,556166 | 0 | 0 | 0,5745233 | 0,821133 | 1,25309692 |
| GSM2261857 | 0 | 0,370935 | 0 | 0,122477 | 0 | 0,46787 | 0,03871827 | 0 | 0,4356592 | 0,901352 | 0,948233129 |
| GSM2261864 | 0 | 0,219817 | 0 | 0,204711 | 0 | 0,441606 | 0,13386521 | 0 | 0,4304391 | 0,90366 | 1,0402269 |
| GSM2261865 | 0 | 0,238282 | 0,013444 | 0,138114 | 0 | 0,551444 | 0,05871661 | 0 | 0,5624251 | 0,828012 | 1,414538552 |
| GSM2261868 | 0 | 0,210119 | 0 | 0,130907 | 0 | 0,658975 | 0 | 0 | 0,6524915 | 0,763674 | 1,932333252 |
| GSM2261870 | 0 | 0,274846 | 0 | 0,132628 | 0 | 0,498253 | 0,09427245 | 0 | 0,4699791 | 0,882658 | 1,222785557 |
| GSM2261871 | 0 | 0,216601 | 0 | 0,234591 | 0 | 0,481257 | 0,06755103 | 0 | 0,4814587 | 0,875448 | 1,066632723 |
| GSM2261872 | 0 | 0,234539 | 0,022461 | 0,179731 | 0 | 0,465967 | 0,09730122 | 0 | 0,5136798 | 0,857482 | 1,066939692 |
| GSM2261873 | 0 | 0,277012 | 0 | 0,114579 | 0 | 0,601974 | 0,00643431 | 0 | 0,48286 | 0,87638 | 1,537251508 |
| GSM2261879 | 0 | 0,335902 | 0 | 0,124842 | 0 | 0,401092 | 0,13816499 | 0 | 0,4673105 | 0,884427 | 0,870532407 |
| GSM2261880 | 0,000733 | 0,308809 | 0 | 0,224267 | 0 | 0,436333 | 0,02985769 | 0 | 0,330597 | 0,954901 | 0,818518766 |
| GSM2261884 | 0 | 0,265331 | 0 | 0,081563 | 0 | 0,653105 | 0 | 0 | 0,530529 | 0,847949 | 1,882719257 |
| GSM2261886 | 0 | 0,229826 | 0,034736 | 0,164441 | 0 | 0,4921 | 0,0788975 | 0 | 0,5152799 | 0,856498 | 1,147077115 |
| GSM2261887 | 0 | 0,293228 | 0 | 0,144099 | 0 | 0,500783 | 0,06188979 | 0 | 0,4315346 | 0,903496 | 1,145100208 |
| GSM2261892 | 0 | 0,18704 | 0 | 0,216383 | 0 | 0,573135 | 0,02344193 | 0 | 0,5826326 | 0,813359 | 1,420678511 |
| GSM2261895 | 0 | 0,218929 | 0 | 0,271 | 0 | 0,407529 | 0,10254157 | 0 | 0,5276288 | 0,848669 | 0,831811835 |
| GSM2261897 | 0 | 0,379586 | 0 | 0,166597 | 0 | 0,395377 | 0,0584395 | 0 | 0,4217677 | 0,907771 | 0,72389091 |
| GSM2261898 | 0 | 0,253478 | 0 | 0,164965 | 0 | 0,512028 | 0,06952964 | 0 | 0,5271453 | 0,849692 | 1,223651575 |
| GSM2261899 | 0,001232 | 0,344772 | 0 | 0,219666 | 0 | 0,314694 | 0,11963561 | 0 | 0,4587644 | 0,8882 | 0,557535101 |
| GSM2261901 | 0 | 0,339249 | 0,04465 | 0,045198 | 0 | 0,570903 | 0 | 0 | 0,7196436 | 0,728777 | 1,330475407 |

| | | | | | | | | | | | |
|---|---|---|---|---|---|---|---|---|---|---|---|
| GSM2261902 | 0 | 0,206713 | 0,066053 | 0,167296 | 0 | 0,559937 | 0 | 0 | 0,6840242 | 0,74898 | 1,272402482 |
| GSM2261903 | 0 | 0,173962 | 0,000761 | 0,202546 | 0 | 0,611681 | 0,01104989 | 0 | 0,5790735 | 0,814911 | 1,621337003 |
| GSM2261904 | 0 | 0,240623 | 0,043109 | 0,151338 | 0 | 0,489191 | 0,07573998 | 0 | 0,4815911 | 0,87569 | 1,124397674 |
| GSM2261906 | 0 | 0,313132 | 0 | 0,181918 | 0 | 0,422677 | 0,0822737 | 0 | 0,4696933 | 0,882469 | 0,853806049 |
| GSM2261908 | 0 | 0,276194 | 0,027918 | 0,200354 | 0 | 0,344048 | 0,15148603 | 0 | 0,441654 | 0,896779 | 0,68200369 |
| GSM2261909 | 0 | 0,304838 | 0 | 0,205111 | 0 | 0,408679 | 0,08137222 | 0 | 0,4270284 | 0,904727 | 0,801412038 |
| GSM2261910 | 0 | 0,151974 | 0,053279 | 0,198002 | 0 | 0,42219 | 0,17455519 | 0 | 0,5534735 | 0,833148 | 1,046955353 |
| GSM2261912 | 0 | 0,252507 | 0,007014 | 0,209101 | 0 | 0,46616 | 0,06521757 | 0 | 0,4619535 | 0,886329 | 0,994746731 |
| GSM2261913 | 0 | 0,293797 | 0 | 0,190114 | 0 | 0,376704 | 0,13938619 | 0 | 0,4884523 | 0,872292 | 0,778457578 |
| GSM2261916 | 0 | 0,181285 | 0,000375 | 0,202817 | 0 | 0,557345 | 0,05817896 | 0 | 0,5911319 | 0,808104 | 1,449619036 |
| GSM2261917 | 0 | 0,282196 | 0 | 0,203846 | 0 | 0,435833 | 0,07812497 | 0 | 0,4566155 | 0,889185 | 0,896698387 |
| GSM2261918 | 0 | 0,202335 | 0 | 0,21476 | 0 | 0,541301 | 0,04160388 | 0 | 0,625186 | 0,786979 | 1,297789677 |
| GSM2261920 | 0 | 0,281554 | 0 | 0,120428 | 0 | 0,553112 | 0,04490547 | 0 | 0,5645381 | 0,827573 | 1,37596095 |
| GSM2261921 | 0 | 0,2255 | 0 | 0,228958 | 0 | 0,46707 | 0,0784718 | 0 | 0,4701074 | 0,88181 | 1,02775157 |
| GSM2261922 | 0 | 0,247315 | 0 | 0,257671 | 0 | 0,435013 | 0,06000159 | 0,27 | 0,0982178 | 1,069497 | 0,861437479 |
| GSM2261923 | 0 | 0,265132 | 0 | 0,174537 | 0 | 0,511478 | 0,04885189 | 0 | 0,5005883 | 0,865071 | 1,163323281 |
| GSM2261924 | 0 | 0,183484 | 0,117029 | 0,108659 | 0 | 0,590828 | 0 | 0 | 0,5948551 | 0,806592 | 1,443957262 |
| GSM2261925 | 0 | 0,270512 | 0,003901 | 0,168633 | 0 | 0,472627 | 0,08432709 | 0 | 0,530595 | 0,848364 | 1,066769306 |
| GSM2261928 | 0 | 0,141847 | 0 | 0,21974 | 0 | 0,391981 | 0,24643124 | 0 | 0,6368631 | 0,778209 | 1,084056832 |
| GSM2261935 | 0 | 0,239889 | 0,034884 | 0,121054 | 0 | 0,593027 | 0,01114613 | 0 | 0,5993144 | 0,80421 | 1,498198712 |
| GSM2261939 | 0 | 0,182263 | 0 | 0,142156 | 0 | 0,543284 | 0,13229675 | 0 | 0,6501716 | 0,768159 | 1,674633844 |
| GSM2261945 | 0 | 0,250856 | 0,019905 | 0,15451 | 0 | 0,523185 | 0,05154414 | 0 | 0,5158019 | 0,856299 | 1,230236977 |
| GSM2261951 | 0 | 0,275446 | 0 | 0,154291 | 0 | 0,458224 | 0,1120395 | 0 | 0,4501185 | 0,89308 | 1,066290603 |
| GSM2261953 | 0 | 0,362946 | 0 | 0,168711 | 0 | 0,412508 | 0,05583588 | 0 | 0,4583452 | 0,888939 | 0,775891282 |
| GSM2261955 | 0 | 0,274932 | 0 | 0,189142 | 0 | 0,535926 | 0 | 0 | 0,5246564 | 0,851012 | 1,154830595 |
| GSM2261957 | 0 | 0,315784 | 0 | 0,127422 | 0 | 0,458136 | 0,09865756 | 0 | 0,4695108 | 0,883046 | 1,033684286 |
| GSM2261962 | 0 | 0,18142 | 0,026745 | 0,140901 | 0 | 0,574624 | 0,07631024 | 0 | 0,535473 | 0,843627 | 1,646177731 |
| GSM2261963 | 0 | 0,301021 | 0 | 0,243992 | 0 | 0,353928 | 0,10105815 | 0 | 0,4444857 | 0,895461 | 0,649393856 |
| GSM2261964 | 0 | 0,177865 | 0,047281 | 0,211452 | 0 | 0,545353 | 0,01804893 | 0 | 0,5097905 | 0,858975 | 1,249096497 |
| GSM2261966 | 0 | 0,318703 | 0 | 0,223412 | 0 | 0,324928 | 0,13295658 | 0 | 0,3826069 | 0,927091 | 0,599370686 |
| GSM2261967 | 0 | 0,198045 | 0 | 0,243653 | 0 | 0,48747 | 0,07083205 | 0 | 0,4510051 | 0,892944 | 1,103627595 |
| GSM2261976 | 0 | 0,363491 | 0 | 0,131817 | 0,007089 | 0,416927 | 0,08067701 | 0 | 0,4969954 | 0,868953 | 0,841754136 |
| GSM2261977 | 0 | 0,243446 | 0 | 0,179576 | 0 | 0,53088 | 0,04609814 | 0 | 0,5332475 | 0,845713 | 1,254968028 |
| GSM2261985 | 0 | 0,172276 | 0 | 0,120351 | 0 | 0,707373 | 0 | 0 | 0,725332 | 0,702014 | 2,417324409 |
| GSM2261988 | 0 | 0,262288 | 0 | 0,235319 | 0 | 0,316808 | 0,18558439 | 0 | 0,4155771 | 0,910635 | 0,636663825 |
| GSM2261989 | 0 | 0,300972 | 0 | 0,225205 | 0 | 0,413035 | 0,06078761 | 0 | 0,3942484 | 0,92196 | 0,784973139 |
| GSM2261991 | 0 | 0,278432 | 0 | 0,194242 | 0 | 0,389447 | 0,13787967 | 0 | 0,397588 | 0,919879 | 0,823924019 |
| GSM2261992 | 0 | 0,351217 | 0 | 0,186698 | 0 | 0,37005 | 0,09203569 | 0 | 0,3866439 | 0,924998 | 0,687933952 |
| GSM2261996 | 0 | 0,175411 | 0,108702 | 0,122063 | 0 | 0,415151 | 0,17867282 | 0 | 0,4878526 | 0,871913 | 1,022097181 |
| GSM2261998 | 0 | 0,315002 | 0 | 0,277258 | 0 | 0,31673 | 0,0910109 | 0,32 | 0,0869392 | 1,071926 | 0,534782023 |
| GSM2262001 | 0 | 0,342952 | 0 | 0,177993 | 0 | 0,405033 | 0,07402206 | 0 | 0,4645449 | 0,885459 | 0,777498195 |
| GSM2262003 | 0 | 0,241119 | 0,026029 | 0,143556 | 0 | 0,325399 | 0,26389811 | 0 | 0,5558548 | 0,833846 | 0,792296761 |
| GSM2262004 | 0 | 0,313401 | 0 | 0,203941 | 0 | 0,318823 | 0,16383543 | 0 | 0,421385 | 0,907345 | 0,616271286 |
| GSM2262007 | 0 | 0,223737 | 0,001176 | 0,165976 | 0 | 0,583845 | 0,0252664 | 0 | 0,5794066 | 0,816283 | 1,493637987 |
| GSM2262013 | 0 | 0,257274 | 0,026118 | 0,17963 | 0 | 0,481759 | 0,05521871 | 0 | 0,4700908 | 0,881977 | 1,040466978 |
| GSM2262017 | 0 | 0,229177 | 0,009509 | 0,153575 | 0 | 0,528855 | 0,07888374 | 0 | 0,4450849 | 0,896743 | 1,348221313 |
| GSM2262020 | 0 | 0,267192 | 0 | 0,220336 | 0 | 0,380254 | 0,13221846 | 0 | 0,4629899 | 0,885545 | 0,779963742 |
| GSM2262023 | 0 | 0,320795 | 0 | 0,219457 | 0 | 0,347467 | 0,11228159 | 0 | 0,3747954 | 0,931054 | 0,643158671 |
| GSM2262025 | 0 | 0,240226 | 0 | 0,165749 | 0 | 0,543949 | 0,05007601 | 0 | 0,6199034 | 0,792073 | 1,339859426 |
| GSM2262026 | 0 | 0,27309 | 0 | 0,258475 | 0 | 0,302606 | 0,16582919 | 0 | 0,4048642 | 0,916389 | 0,569273969 |
| GSM2262028 | 0 | 0,306654 | 0 | 0,200368 | 0 | 0,388925 | 0,10405368 | 0 | 0,451529 | 0,891864 | 0,767078671 |
| GSM2262030 | 0 | 0,317877 | 0 | 0,214207 | 0 | 0,409809 | 0,05810746 | 0 | 0,4604197 | 0,887231 | 0,77019639 |
| GSM2262031 | 0 | 0,333899 | 0 | 0,257743 | 0 | 0,358173 | 0,05018493 | 0 | 0,3336406 | 0,953516 | 0,605387091 |
| GSM2262033 | 0 | 0,27024 | 0 | 0,203311 | 0 | 0,505621 | 0,0208284 | 0 | 0,4415626 | 0,897947 | 1,067723313 |
| GSM2262034 | 0 | 0,346484 | 0,002526 | 0,201689 | 0 | 0,391391 | 0,05790955 | 0 | 0,3632424 | 0,937107 | 0,710717335 |
| GSM2262035 | 0 | 0,261697 | 0 | 0,232849 | 0 | 0,412468 | 0,09298627 | 0 | 0,47578 | 0,878573 | 0,834032861 |
| GSM2262041 | 0 | 0,238937 | 0 | 0,287957 | 0 | 0,351702 | 0,12140382 | 0 | 0,5056593 | 0,861458 | 0,667499765 |
| GSM2262042 | 0 | 0,193828 | 0,03513 | 0,100321 | 0 | 0,60081 | 0,06991031 | 0 | 0,5178231 | 0,855071 | 1,824617006 |
| GSM2262044 | 0 | 0,323322 | 0,017346 | 0,19887 | 0 | 0,38749 | 0,07297138 | 0 | 0,4761532 | 0,878985 | 0,718189061 |
| GSM2262046 | 0 | 0,336333 | 0 | 0,227409 | 0 | 0,349057 | 0,08720151 | 0 | 0,3509421 | 0,943222 | 0,619178161 |
| GSM2262049 | 0 | 0,151985 | 0,00728 | 0,048479 | 0 | 0,792256 | 0 | 0 | 0,6882746 | 0,726453 | 3,813608302 |
| GSM2262051 | 0 | 0,334595 | 0 | 0,137475 | 0,008598 | 0,372266 | 0,14706646 | 0 | 0,3803882 | 0,928085 | 0,788583381 |
| GSM2262053 | 0 | 0,304028 | 0 | 0,261645 | 0 | 0,25021 | 0,18411723 | 0 | 0,3749055 | 0,932024 | 0,442322194 |
| GSM2262059 | 0 | 0,228082 | 0,002178 | 0,184605 | 0 | 0,570874 | 0,01426058 | 0 | 0,5463267 | 0,837194 | 1,376046229 |
| GSM2262060 | 0 | 0,315082 | 0 | 0,246558 | 0 | 0,372215 | 0,06614419 | 0 | 0,3567773 | 0,941196 | 0,662728261 |
| GSM2262061 | 0 | 0,240772 | 0 | 0,268121 | 0 | 0,404186 | 0,08692069 | 0 | 0,4351719 | 0,900894 | 0,794243416 |
| GSM2262063 | 0 | 0,37526 | 0 | 0,172292 | 0 | 0,427638 | 0,02480942 | 0 | 0,3871657 | 0,925586 | 0,781000493 |
| GSM2262064 | 0 | 0,218544 | 0,00828 | 0,202709 | 0 | 0,517893 | 0,05257467 | 0 | 0,4257767 | 0,90712 | 1,205715363 |
| GSM2262067 | 0 | 0,266352 | 0 | 0,127049 | 0,003055 | 0,547507 | 0,05603643 | 0 | 0,5007263 | 0,865277 | 1,391729274 |
| GSM2262068 | 0 | 0,287788 | 0 | 0,142126 | 0 | 0,485303 | 0,08478383 | 0 | 0,4565132 | 0,889888 | 1,128838043 |
| GSM2262077 | 0 | 0,351772 | 0 | 0,165433 | 0,004254 | 0,354266 | 0,12427471 | 0 | 0,3212869 | 0,95636 | 0,68496321 |
| GSM2262078 | 0 | 0,319029 | 0 | 0,209013 | 0 | 0,328971 | 0,14298711 | 0 | 0,4866052 | 0,87344 | 0,623000936 |
| GSM2262080 | 0 | 0,30995 | 0,00215 | 0,202254 | 0 | 0,396443 | 0,08920341 | 0 | 0,3586326 | 0,939376 | 0,770758496 |
| GSM2262082 | 0 | 0,270924 | 0,01189 | 0,213779 | 0 | 0,458149 | 0,04525803 | 0 | 0,4364645 | 0,900131 | 0,922586343 |
| GSM2262083 | 0 | 0,197475 | 0 | 0,223247 | 0 | 0,478689 | 0,10058817 | 0 | 0,5079103 | 0,860149 | 1,137778929 |
| GSM2262086 | 0 | 0,263008 | 0 | 0,1399 | 0,002874 | 0,538357 | 0,05586115 | 0 | 0,5383725 | 0,843068 | 1,336178039 |
| GSM2262091 | 0 | 0,242911 | 0 | 0,131981 | 0 | 0,625108 | 0 | 0 | 0,4982298 | 0,867255 | 1,667436866 |
| GSM2262094 | 0 | 0,223532 | 0,063702 | 0,141522 | 0 | 0,517812 | 0,05343148 | 0 | 0,5159647 | 0,856076 | 1,207707597 |
| GSM2262098 | 0 | 0,344739 | 0 | 0,121158 | 0 | 0,505943 | 0,02816049 | 0 | 0,4216614 | 0,909028 | 1,085956186 |
| GSM2262102 | 0 | 0,198693 | 0,025661 | 0,125544 | 0 | 0,650102 | 0 | 0 | 0,5750324 | 0,817855 | 1,85797951 |
| GSM2262103 | 0 | 0,254013 | 0 | 0,211219 | 0 | 0,353198 | 0,18156987 | 0 | 0,4592268 | 0,887574 | 0,759187862 |
| GSM2262104 | 0 | 0,179977 | 0,013312 | 0,132121 | 0 | 0,519735 | 0,15485495 | 0 | 0,5562343 | 0,830695 | 1,597169159 |
| GSM2262105 | 0 | 0,280032 | 0 | 0,145006 | 0 | 0,501241 | 0,07372088 | 0 | 0,5068106 | 0,861925 | 1,179284109 |

| | | | | | | | | | | | |
|---|---|---|---|---|---|---|---|---|---|---|---|
| GSM2262107 | 0 | 0,259284 | 0,063639 | 0,191599 | 0 | 0,374069 | 0,11140979 | 0 | 0,5167811 | 0,856551 | 0,727022327 |
| GSM2262116 | 0,014739 | 0,104479 | 0,02167 | 0,105623 | 0,006221 | 0,670325 | 0,07694189 | 0 | 0,6550247 | 0,757527 | 2,892174228 |
| GSM2262117 | 0 | 0,295512 | 0 | 0,177497 | 0 | 0,409496 | 0,11749505 | 0 | 0,47924 | 0,877283 | 0,865725743 |
| GSM2262119 | 0 | 0,262642 | 0,012564 | 0,081865 | 0 | 0,642929 | 0 | 0 | 0,661068 | 0,760206 | 1,800560069 |
| GSM2262120 | 0 | 0,332788 | 0,000884 | 0,22195 | 0 | 0,36527 | 0,07910752 | 0 | 0,470707 | 0,881799 | 0,657407243 |
| GSM2262122 | 0 | 0,392371 | 0 | 0,141939 | 0 | 0,336357 | 0,12933291 | 0 | 0,4097048 | 0,91372 | 0,629515931 |
| GSM2262123 | 0 | 0,270588 | 0,003018 | 0,179856 | 0 | 0,479665 | 0,06687356 | 0 | 0,5400997 | 0,84275 | 1,057786136 |
| GSM2262124 | 0 | 0,341541 | 0 | 0,109762 | 0 | 0,548697 | 0 | 0 | 0,4419429 | 0,898983 | 1,215804641 |
| GSM2262126 | 0 | 0,125585 | 0,115379 | 0,128797 | 0 | 0,63024 | 0 | 0 | 0,6633705 | 0,756865 | 1,704454424 |
| GSM2262127 | 0 | 0,310225 | 0 | 0,194676 | 0 | 0,474822 | 0,02027744 | 0 | 0,4311099 | 0,903232 | 0,94042707 |
| GSM2262131 | 0 | 0,280924 | 0 | 0,211505 | 0 | 0,414477 | 0,0930941 | 0 | 0,4406624 | 0,897578 | 0,841699349 |
| GSM2262138 | 0 | 0,31151 | 0 | 0,160743 | 0 | 0,500581 | 0,02716564 | 0 | 0,3899282 | 0,925325 | 1,059984459 |
| GSM2262144 | 0 | 0,342588 | 0 | 0,16173 | 0 | 0,455865 | 0,0398166 | 0 | 0,4193203 | 0,909286 | 0,903924068 |
| GSM2262146 | 0 | 0,284539 | 0 | 0,11254 | 2,18E-05 | 0,589109 | 0,01379047 | 0 | 0,5212913 | 0,853405 | 1,483605277 |
| GSM2262148 | 0,003911 | 0,268477 | 0 | 0,168873 | 0,01396 | 0,495678 | 0,04910115 | 0 | 0,5410698 | 0,841946 | 1,133366471 |
| GSM2262149 | 0 | 0,175427 | 0,060993 | 0,132475 | 0 | 0,542669 | 0,08843644 | 0 | 0,5743293 | 0,819805 | 1,471065735 |
| GSM2262152 | 0 | 0,309629 | 0 | 0,216658 | 0 | 0,418418 | 0,05529525 | 0 | 0,4589056 | 0,888003 | 0,795036533 |
| GSM2262159 | 0 | 0,281339 | 0 | 0,232556 | 0 | 0,43769 | 0,04841515 | 0 | 0,3755406 | 0,932241 | 0,851709625 |
| GSM2262161 | 0 | 0,240112 | 0 | 0,202253 | 0,001941 | 0,531867 | 0,02382742 | 0 | 0,5607044 | 0,828933 | 1,202328108 |
| GSM2262162 | 0 | 0,198492 | 0,008497 | 0,201865 | 0 | 0,541374 | 0,04977116 | 0 | 0,5826182 | 0,814449 | 1,324123795 |
| GSM2262163 | 0 | 0,278046 | 0 | 0,228185 | 0 | 0,475502 | 0,01826578 | 0 | 0,6166922 | 0,796243 | 0,939298028 |
| GSM2262164 | 0 | 0,291692 | 0 | 0,199213 | 0 | 0,503666 | 0,00542933 | 0 | 0,4566785 | 0,889652 | 1,025993571 |
| GSM2262166 | 0 | 0,302299 | 0,027281 | 0,207658 | 0 | 0,399636 | 0,06312583 | 0 | 0,50214 | 0,864665 | 0,743872332 |
| GSM2262168 | 0 | 0,179467 | 0 | 0,150198 | 0 | 0,655617 | 0,01471768 | 0 | 0,5470404 | 0,836218 | 1,988736695 |
| GSM2262173 | 0 | 0,357411 | 0 | 0,197336 | 0,002476 | 0,397275 | 0,04550174 | 0 | 0,3596405 | 0,938995 | 0,716135523 |
| GSM2262175 | 0 | 0,254797 | 0,013124 | 0,089817 | 0 | 0,642262 | 0 | 0 | 0,5932091 | 0,806795 | 1,795344723 |
| GSM2262176 | 0 | 0,237851 | 0 | 0,11601 | 0 | 0,519777 | 0,12636216 | 0 | 0,4821169 | 0,876197 | 1,468870857 |
| GSM2262177 | 0 | 0,251294 | 0 | 0,146468 | 0 | 0,543568 | 0,05867016 | 0 | 0,5015412 | 0,864605 | 1,366566257 |
| GSM2262179 | 0 | 0,254796 | 0 | 0,254001 | 0 | 0,428747 | 0,06245608 | 0 | 0,5280591 | 0,848883 | 0,842666661 |
| GSM2262180 | 0 | 0,27167 | 0,000111 | 0,19885 | 0 | 0,465969 | 0,0634008 | 0 | 0,503316 | 0,863555 | 0,990095338 |
| GSM2262183 | 0 | 0,44675 | 0 | 0,174825 | 0 | 0,292713 | 0,08571219 | 0 | 0,3437489 | 0,946435 | 0,470921899 |
| GSM2262185 | 0 | 0,36632 | 0 | 0,184121 | 0 | 0,417268 | 0,03229062 | 0 | 0,384678 | 0,926699 | 0,758061385 |
| GSM2262186 | 0 | 0,282196 | 0 | 0,157968 | 0 | 0,511206 | 0,04863053 | 0 | 0,4440619 | 0,89675 | 1,161399592 |
| GSM2262187 | 0 | 0,244197 | 0,059318 | 0,165178 | 0 | 0,531307 | 0 | 0 | 0,5882715 | 0,812983 | 1,133594522 |
| GSM2262191 | 0 | 0,328264 | 0 | 0,195279 | 0 | 0,415142 | 0,06131527 | 0 | 0,3040853 | 0,966383 | 0,792946408 |
| GSM2262193 | 0 | 0,219466 | 0,012826 | 0,229056 | 0 | 0,523451 | 0,01520158 | 0 | 0,5098974 | 0,859023 | 1,134611434 |
| GSM2262194 | 0 | 0,280284 | 0 | 0,180221 | 0 | 0,452798 | 0,08669708 | 0 | 0,4476145 | 0,894178 | 0,983265607 |
| GSM2262196 | 0 | 0,284524 | 0,112861 | 0,089407 | 0 | 0,488209 | 0,02499976 | 0 | 0,5938764 | 0,812888 | 1,002912417 |
| GSM2262200 | 0 | 0,243691 | 0 | 0,248068 | 0 | 0,508241 | 0 | 0 | 0,4214932 | 0,909665 | 1,033516037 |
| GSM2262201 | 0 | 0,259992 | 0 | 0,20125 | 0 | 0,519123 | 0,01963491 | 0 | 0,4604516 | 0,887635 | 1,12548993 |
| GSM2262203 | 0 | 0,24266 | 0,022493 | 0,170567 | 0 | 0,476378 | 0,08790162 | 0 | 0,4199396 | 0,909243 | 1,093311995 |
| GSM2262211 | 0 | 0,290829 | 0 | 0,150412 | 0 | 0,55876 | 0 | 0 | 0,4619132 | 0,887522 | 1,266338691 |
| GSM2262213 | 0 | 0,367623 | 0 | 0,263978 | 0 | 0,279253 | 0,08914544 | 0 | 0,3136758 | 0,963696 | 0,44213394 |
| GSM2262215 | 0 | 0,296621 | 0 | 0,209504 | 0 | 0,482792 | 0,0110836 | 0 | 0,3859745 | 0,927401 | 0,953899535 |
| GSM2262216 | 0 | 0,261102 | 0 | 0,195423 | 0 | 0,510223 | 0,03325247 | 0 | 0,401476 | 0,919817 | 1,117625849 |
| GSM2262219 | 0 | 0,142752 | 0 | 0,145513 | 0 | 0,711734 | 0 | 0 | 0,5214516 | 0,854933 | 2,469023144 |
| GSM2262221 | 0 | 0,163941 | 0,009104 | 0,102634 | 0 | 0,59333 | 0,13099073 | 0 | 0,6937203 | 0,732027 | 2,152251404 |
| GSM2262224 | 0 | 0,207713 | 0,006406 | 0,12171 | 0 | 0,606595 | 0,057577 | 0 | 0,4893891 | 0,872758 | 1,806266082 |
| GSM2262227 | 0 | 0,090243 | 0,003911 | 0,183794 | 0,017903 | 0,407568 | 0,29658096 | 0 | 0,6385374 | 0,77279 | 1,466346356 |
| GSM2262228 | 0 | 0,267776 | 0 | 0,129438 | 0 | 0,546245 | 0,05654149 | 0 | 0,5490932 | 0,836705 | 1,375192035 |
| GSM2262231 | 0 | 0,106445 | 0,010019 | 0,073086 | 0 | 0,596948 | 0,21350129 | 0 | 0,6988897 | 0,720178 | 3,149286783 |
| GSM2262235 | 0 | 0,272759 | 0,001144 | 0,150037 | 0 | 0,5188 | 0,05726129 | 0 | 0,4779209 | 0,878122 | 1,22375947 |
| GSM2262237 | 0 | 0,188073 | 0,015726 | 0,061962 | 0 | 0,734238 | 0 | 0 | 0,6451119 | 0,765157 | 2,762772119 |
| GSM2262238 | 0 | 0,209969 | 0 | 0,260129 | 0 | 0,362156 | 0,16774561 | 0 | 0,40348 | 0,918029 | 0,770385054 |
| GSM2262239 | 0 | 0,271283 | 0 | 0,210187 | 0 | 0,467746 | 0,05078312 | 0 | 0,4315011 | 0,90292 | 0,971494526 |
| GSM2262241 | 0 | 0,173522 | 0 | 0,177886 | 0 | 0,502063 | 0,14652939 | 0 | 0,4644185 | 0,886191 | 1,428716348 |
| GSM2262242 | 0 | 0,363293 | 0 | 0,208263 | 0 | 0,371051 | 0,05739338 | 0 | 0,3867926 | 0,925352 | 0,649193744 |
| GSM2262243 | 0 | 0,298722 | 0 | 0,224848 | 0 | 0,311076 | 0,16535399 | 0 | 0,4067348 | 0,914912 | 0,59414388 |
| GSM2262246 | 0 | 0,281566 | 0 | 0,077437 | 0,007718 | 0,633279 | 0 | 0 | 0,5904901 | 0,809188 | 1,763991084 |
| GSM2262251 | 0 | 0,306048 | 0 | 0,211315 | 0 | 0,356418 | 0,12621937 | 0 | 0,3741888 | 0,931335 | 0,6889122 |
| GSM2262253 | 0 | 0,388774 | 0 | 0,23795 | 0 | 0,286666 | 0,08660929 | 0 | 0,3323123 | 0,953218 | 0,457404397 |
| GSM2262256 | 0 | 0,364277 | 0 | 0,214575 | 0 | 0,356961 | 0,06418744 | 0 | 0,3754936 | 0,931012 | 0,616669918 |
| GSM2262258 | 0 | 0,243027 | 0 | 0,194719 | 0 | 0,387728 | 0,17452612 | 0 | 0,4600763 | 0,88724 | 0,885735536 |
| GSM2262264 | 0 | 0,349256 | 0 | 0,23245 | 0,008692 | 0,39632 | 0,01328201 | 0 | 0,4431298 | 0,896665 | 0,681306814 |
| GSM2262265 | 0 | 0,335482 | 0 | 0,178499 | 0 | 0,352916 | 0,13310365 | 0 | 0,4040753 | 0,916165 | 0,686632906 |
| GSM2262266 | 0 | 0,277653 | 0 | 0,247161 | 0 | 0,365363 | 0,10982275 | 0 | 0,4296568 | 0,903319 | 0,696177223 |
| GSM2262268 | 0 | 0,301624 | 0 | 0,214355 | 0 | 0,351414 | 0,1326064 | 0 | 0,4538456 | 0,890527 | 0,681063186 |
| GSM2262270 | 0 | 0,350599 | 0 | 0,19801 | 0 | 0,384294 | 0,06709688 | 0 | 0,3166383 | 0,959711 | 0,700488638 |
| GSM2262274 | 0 | 0,307468 | 0 | 0,162862 | 0 | 0,52967 | 0 | 0 | 0,4404346 | 0,898985 | 1,1261663 |
| GSM2262276 | 0 | 0,277476 | 0 | 0,182542 | 0 | 0,448489 | 0,09149347 | 0 | 0,4626012 | 0,88609 | 0,974937811 |
| GSM2262282 | 0 | 0,274133 | 0 | 0,20497 | 0 | 0,520898 | 0 | 0 | 0,4254132 | 0,907108 | 1,087236982 |
| GSM2262290 | 0 | 0,220096 | 0 | 0,174806 | 0 | 0,52084 | 0,08425753 | 0 | 0,5509595 | 0,834812 | 1,318910387 |
| GSM2262293 | 0 | 0,224589 | 0 | 0,347374 | 0 | 0,340788 | 0,08724883 | 0 | 0,4097772 | 0,917322 | 0,595821362 |
| GSM2262297 | 0 | 0,283189 | 0 | 0,174355 | 0 | 0,537242 | 0,00521447 | 0 | 0,4349881 | 0,902066 | 1,174189695 |
| GSM2262303 | 0 | 0,184638 | 0,021264 | 0,186729 | 0 | 0,501315 | 0,10605433 | 0 | 0,5506549 | 0,834665 | 1,276808565 |
| GSM2262305 | 0 | 0,28583 | 0 | 0,223337 | 0 | 0,490833 | 0 | 0 | 0,4668311 | 0,883927 | 0,963991126 |
| GSM2262307 | 0 | 0,320683 | 0 | 0,19952 | 0 | 0,450515 | 0,0292826 | 0 | 0,5112922 | 0,859671 | 0,866036351 |
| GSM2262311 | 0 | 0,328465 | 0 | 0,246541 | 0,014404 | 0,358989 | 0,05160085 | 0 | 0,3985158 | 0,919988 | 0,624321341 |
| GSM2262312 | 0 | 0,359627 | 0 | 0,217841 | 0 | 0,361882 | 0,06064947 | 0 | 0,4177022 | 0,909678 | 0,626670209 |
| GSM2262313 | 0 | 0,291208 | 0 | 0,21012 | 0 | 0,412632 | 0,08604071 | 0 | 0,4714277 | 0,8812 | 0,823077375 |
| **average** | **6,32365E-(** | **0,262823** | **0,008781** | **0,182259** | **0,000423** | **0,473163** | **0,07248865** | | | | |

**SES_CIBERSORTx_GSE85217_GP4**

| Sample | Bcells | TCD4 | TCD8 | Tgd | NK | MoMaDC | granulocytes | P-value | Correlation | RMSE | | Tab |
|---|---|---|---|---|---|---|---|---|---|---|---|---|
| GSM2261540 | 0 | 1,048256 | 0 | 0,99693 | 0 | 1,438091 | 0,29749091 | 0 | 0,3598455 | 0,940333 | | 1,048256 |
| GSM2261541 | 0 | 0,68413 | 0 | 0,827963 | 0 | 1,622162 | 0,67933499 | 0 | 0,4104143 | 0,915444 | | 0,68413 |
| GSM2261542 | 0 | 0,755829 | 0,013047 | 0,839584 | 0 | 1,899937 | 0,98587899 | 0 | 0,4269089 | 0,907196 | | 0,768876 |
| GSM2261546 | 0 | 0,817528 | 0 | 0,88573 | 0 | 1,456091 | 0,30522966 | 0 | 0,3801833 | 0,930376 | | 0,817528 |
| GSM2261556 | 0 | 0,629614 | 0 | 0,592365 | 0 | 0,831099 | 0,32320261 | 0 | 0,3766431 | 0,930994 | | 0,629614 |
| GSM2261557 | 0 | 1,220233 | 0 | 0,593416 | 0 | 0,843873 | 0,33679887 | 0 | 0,3893759 | 0,923897 | | 1,220233 |
| GSM2261558 | 0 | 2,890453 | 0 | 1,341038 | 0 | 5,267489 | 0 | 0 | 0,6588625 | 0,769132 | | 2,890453 |
| GSM2261562 | 0 | 1,483726 | 0,110233 | 0,919442 | 0 | 3,043939 | 0 | 0 | 0,5350412 | 0,844935 | | 1,593959 |
| GSM2261563 | 0 | 1,286151 | 0 | 0,847632 | 0 | 2,219035 | 0,23745486 | 0 | 0,4372366 | 0,899988 | | 1,286151 |
| GSM2261564 | 0 | 0,923103 | 0,02569 | 0,613186 | 0 | 2,782502 | 0 | 0 | 0,6266741 | 0,783042 | | 0,948793 |
| GSM2261572 | 0 | 1,272266 | 0 | 0,667374 | 0 | 1,516556 | 0,16928342 | 0 | 0,3896014 | 0,924075 | | 1,272266 |
| GSM2261573 | 0 | 1,221358 | 0 | 0,862742 | 0 | 4,938655 | 0 | 0 | 0,6157798 | 0,787883 | | 1,221358 |
| GSM2261575 | 0 | 1,088281 | 0 | 0,66095 | 0 | 1,97808 | 0,02299456 | 0 | 0,4345432 | 0,902119 | | 1,088281 |
| GSM2261576 | 0 | 1,371158 | 0 | 1,126079 | 0 | 2,684175 | 0,10149182 | 0 | 0,5068694 | 0,861159 | | 1,371158 |
| GSM2261578 | 0 | 1,314017 | 0 | 0,971978 | 0 | 1,993332 | 0,03220333 | 0 | 0,4076931 | 0,915829 | | 1,314017 |
| GSM2261579 | 0 | 0,928242 | 0,281596 | 0,831071 | 0,053972 | 5,037814 | 1,6559968 | 0 | 0,6514869 | 0,761268 | | 1,209837 |
| GSM2261582 | 0 | 0,908717 | 0 | 0,632604 | 0 | 1,868764 | 0,06325316 | 0 | 0,5469887 | 0,837689 | | 0,908717 |
| GSM2261583 | 0 | 1,284178 | 0 | 0,497447 | 0 | 2,244189 | 0,11539378 | 0 | 0,4859709 | 0,874145 | | 1,284178 |
| GSM2261584 | 0 | 0,916461 | 0 | 0,77267 | 0 | 1,448774 | 0,22607879 | 0 | 0,3796134 | 0,929945 | | 0,916461 |
| GSM2261585 | 0 | 1,211025 | 0 | 0,666116 | 0 | 2,188869 | 0,02254274 | 0 | 0,3937326 | 0,924369 | | 1,211025 |
| GSM2261587 | 0 | 0,965575 | 0 | 0,827437 | 0 | 0,951745 | 0,47602321 | 0 | 0,388295 | 0,924851 | | 0,965575 |
| GSM2261588 | 0 | 1,39701 | 0 | 0,457278 | 0 | 1,725554 | 0 | 0 | 0,4135607 | 0,913127 | | 1,39701 |
| GSM2261595 | 0 | 0,888439 | 0 | 0,677073 | 0 | 0,74333 | 0,34283977 | 0 | 0,4410479 | 0,897451 | | 0,888439 |
| GSM2261596 | 0 | 1,2184 | 0 | 0,997252 | 0 | 1,922597 | 0,24147242 | 0 | 0,4368451 | 0,899841 | | 1,2184 |
| GSM2261599 | 0 | 1,577004 | 0 | 0,863878 | 0 | 2,019116 | 0,16663607 | 0 | 0,4039992 | 0,91698 | | 1,577004 |
| GSM2261606 | 0 | 1,050527 | 0 | 0,716082 | 0 | 3,258423 | 0,55129042 | 0 | 0,4950276 | 0,86927 | | 1,050527 |
| GSM2261607 | 0 | 1,023169 | 0 | 0,648957 | 0 | 0,978926 | 0,18826391 | 0 | 0,3578219 | 0,940093 | | 1,023169 |
| GSM2261608 | 0 | 1,11907 | 0 | 0,749357 | 0 | 1,511417 | 0,18791156 | 0 | 0,3775117 | 0,930466 | | 1,11907 |
| GSM2261610 | 0 | 0,825345 | 0 | 0,629794 | 0 | 1,335997 | 0,19668739 | 0 | 0,3560214 | 0,942051 | | 0,825345 |
| GSM2261611 | 0 | 0,943665 | 0 | 0,674728 | 0 | 1,212284 | 0,09523397 | 0 | 0,3679554 | 0,935781 | | 0,943665 |
| GSM2261616 | 0 | 1,068174 | 0 | 0,63267 | 0 | 1,794612 | 0,02220834 | 0 | 0,4352075 | 0,901486 | | 1,068174 |
| GSM2261618 | 0 | 0,733854 | 0 | 0,773816 | 0 | 1,134924 | 0,4014429 | 0 | 0,3724885 | 0,933728 | | 0,733854 |
| GSM2261621 | 0 | 1,080545 | 0 | 0,701953 | 0,010578 | 1,236406 | 0,36153087 | 0 | 0,3856225 | 0,925553 | | 1,080545 |
| GSM2261623 | 0 | 0,924789 | 0 | 1,100049 | 0 | 2,424186 | 0,20178022 | 0 | 0,492003 | 0,869538 | | 0,924789 |
| GSM2261626 | 0 | 1,118992 | 0 | 0,460634 | 0 | 1,532973 | 0,20613058 | 0 | 0,3354146 | 0,951844 | | 1,118992 |
| GSM2261627 | 0 | 1,176643 | 0 | 1,202424 | 0 | 1,53423 | 0,7848818 | 0 | 0,449478 | 0,892751 | | 1,176643 |
| GSM2261630 | 0 | 1,98556 | 0 | 1,185647 | 0 | 4,542554 | 0,05837899 | 0 | 0,4841993 | 0,874958 | | 1,98556 |
| GSM2261631 | 0 | 1,119775 | 0 | 1,181555 | 0 | 2,689168 | 0,11351087 | 0 | 0,5002503 | 0,86473 | | 1,119775 |
| GSM2261632 | 0 | 0,643897 | 2,74E-05 | 1,15451 | 0 | 2,234932 | 0,39857146 | 0 | 0,4272002 | 0,907951 | | 0,643924 |
| GSM2261637 | 0 | 0,65133 | 0,051823 | 0,552545 | 0 | 1,316171 | 0,20766824 | 0 | 0,511416 | 0,858591 | | 0,703153 |
| GSM2261638 | 0 | 1,553379 | 0 | 0,985672 | 0 | 2,311917 | 0,42657581 | 0 | 0,3925381 | 0,922823 | | 1,553379 |
| GSM2261639 | 0 | 0,84341 | 0,390353 | 0,719839 | 0 | 2,60952 | 0,00313412 | 0 | 0,6538752 | 0,7682 | | 1,233763 |
| GSM2261640 | 0 | 0,744217 | 0 | 0,426546 | 0 | 1,454992 | 0,17677976 | 0 | 0,5580011 | 0,831716 | | 0,744217 |
| GSM2261651 | 0 | 0,654346 | 0,02423 | 0,32066 | 0 | 0,762749 | 0,2555805 | 0 | 0,3968833 | 0,919872 | | 0,678576 |
| GSM2261652 | 0 | 1,622227 | 0 | 0,803377 | 0 | 5,202212 | 0 | 0 | 0,7052547 | 0,722208 | | 1,622227 |
| GSM2261653 | 0 | 0,988988 | 0,072601 | 0,647989 | 0 | 1,58561 | 0,26049951 | 0 | 0,5057664 | 0,862532 | | 1,061589 |
| GSM2261654 | 0 | 1,383109 | 0,29817 | 1,252694 | 0 | 4,74318 | 0 | 0 | 0,6413231 | 0,77349 | | 1,681279 |
| GSM2261656 | 0 | 1,404887 | 0 | 1,084376 | 0 | 3,107201 | 0,33855909 | 0 | 0,5801119 | 0,817396 | | 1,404887 |
| GSM2261659 | 0 | 1,436977 | 0 | 0,893967 | 0 | 1,583108 | 0,14768249 | 0 | 0,42779 | 0,904638 | | 1,436977 |
| GSM2261660 | 0 | 0,998833 | 0 | 0,614013 | 0 | 1,279459 | 0,20317114 | 0 | 0,4048685 | 0,916221 | | 0,998833 |
| GSM2261662 | 0 | 1,085295 | 0 | 0,915342 | 0,008422 | 2,786422 | 0,25636623 | 0 | 0,7135303 | 0,727436 | | 1,085295 |
| GSM2261663 | 0 | 0,785171 | 0 | 0,722711 | 0 | 1,534594 | 0,31947367 | 0 | 0,4987043 | 0,865737 | | 0,785171 |
| GSM2261664 | 0 | 1,215712 | 0 | 1,029042 | 0 | 4,37113 | 0,46848641 | 0 | 0,6738141 | 0,748159 | | 1,215712 |
| GSM2261665 | 0 | 0,672991 | 0,001696 | 0,596758 | 0 | 1,378647 | 0,15053387 | 0 | 0,4691196 | 0,882519 | | 0,674687 |
| GSM2261666 | 0 | 1,035429 | 0 | 0,670034 | 0 | 1,056736 | 0,38473979 | 0 | 0,3595558 | 0,938431 | | 1,035429 |
| GSM2261669 | 0 | 1,249035 | 0 | 0,442922 | 0 | 2,568389 | 0,03752389 | 0 | 0,5511342 | 0,835262 | | 1,249035 |
| GSM2261670 | 0 | 0,875152 | 0 | 0,520077 | 0 | 1,090016 | 0,39838009 | 0 | 0,4537057 | 0,890812 | | 0,875152 |
| GSM2261671 | 0 | 0,665064 | 0,077654 | 0,838009 | 0 | 2,245868 | 0,10081272 | 0 | 0,5526488 | 0,832444 | | 0,742718 |
| GSM2261672 | 0 | 1,49065 | 0 | 0,878266 | 0 | 1,832471 | 0,58391245 | 0 | 0,4269737 | 0,904653 | | 1,49065 |
| GSM2261677 | 0 | 0,79135 | 0,150736 | 0,547813 | 0 | 1,876082 | 0,10961277 | 0 | 0,5355867 | 0,844321 | | 0,942087 |
| GSM2261679 | 0 | 1,256445 | 0 | 0,501491 | 0,020403 | 3,164441 | 0,00448003 | 0 | 0,6049073 | 0,798832 | | 1,256445 |
| GSM2261681 | 0 | 0,848171 | 0 | 0,695429 | 0 | 1,263328 | 0,20550657 | 0 | 0,3932821 | 0,922649 | | 0,848171 |
| GSM2261682 | 0 | 0,990756 | 0,11104 | 0,487155 | 0 | 2,214868 | 0,05626392 | 0 | 0,5997248 | 0,804975 | | 1,101795 |
| GSM2261683 | 0 | 1,360553 | 0 | 1,059936 | 0 | 5,733083 | 0 | 0 | 0,6962773 | 0,725957 | | 1,360553 |
| GSM2261684 | 0 | 1,245356 | 0,270279 | 1,079664 | 0 | 3,539981 | 0,01559074 | 0 | 0,6445191 | 0,774085 | | 1,515636 |
| GSM2261685 | 0 | 0,826423 | 0 | 0,660135 | 0 | 1,973333 | 0,14656305 | 0 | 0,5241072 | 0,850864 | | 0,826423 |
| GSM2261686 | 0 | 1,382597 | 0 | 1,051248 | 0 | 2,564184 | 0,24436231 | 0 | 0,5058077 | 0,861965 | | 1,382597 |
| GSM2261688 | 0 | 0,610191 | 0,348741 | 0,777167 | 0 | 3,022689 | 0,26793199 | 0 | 0,5893368 | 0,807998 | | 0,958932 |
| GSM2261690 | 0 | 0,933381 | 0,005978 | 0,506971 | 0 | 2,250884 | 0 | 0 | 0,5850128 | 0,812855 | | 0,939358 |
| GSM2261692 | 0 | 1,462673 | 0,018389 | 0,829756 | 0 | 5,522296 | 0 | 0 | 0,6991991 | 0,724041 | | 1,481062 |
| GSM2261693 | 0 | 0,704258 | 0,512164 | 1,052997 | 0 | 2,505876 | 0,19667536 | 0 | 0,5339336 | 0,844375 | | 1,216421 |
| GSM2261694 | 0 | 0,711163 | 0 | 0,495681 | 0 | 1,045574 | 0,10397477 | 0 | 0,4341332 | 0,901299 | | 0,711163 |
| GSM2261695 | 0 | 0,637272 | 0,01938 | 0,489519 | 0 | 1,343293 | 0,13241631 | 0 | 0,4891955 | 0,871338 | | 0,656652 |
| GSM2261696 | 0 | 0,722042 | 0,071168 | 0,416583 | 0,001403 | 1,106915 | 0,30333034 | 0 | 0,4059575 | 0,915687 | | 0,79321 |
| GSM2261697 | 0 | 0,795732 | 0 | 0,47288 | 0 | 1,054305 | 0,1948355 | 0 | 0,4349284 | 0,900711 | | 0,795732 |
| GSM2261698 | 0 | 0,913594 | 0 | 0,569324 | 0 | 1,347439 | 0,08495863 | 0 | 0,4241211 | 0,906782 | | 0,913594 |
| GSM2261699 | 0 | 0,956357 | 0 | 0,653539 | 0,012764 | 1,550131 | 0,2024246 | 0 | 0,5430439 | 0,841313 | | 0,956357 |
| GSM2261703 | 0 | 0,954428 | 0 | 0,745948 | 0 | 1,100299 | 0,34020677 | 0 | 0,4079494 | 0,914498 | | 0,954428 |
| GSM2261704 | 0 | 1,164233 | 0,09541 | 0,784476 | 0 | 1,722923 | 0,16692809 | 0 | 0,4794055 | 0,876952 | | 1,259643 |
| GSM2261705 | 0 | 0,938567 | 0,062212 | 0,6099 | 0 | 2,08371 | 0,38126488 | 0 | 0,6072588 | 0,800972 | | 1,000779 |

| | | | | | | | | | | | | |
|---|---|---|---|---|---|---|---|---|---|---|---|---|
| GSM2261714 | 0 | 0,956597 | 0,111689 | 0,44642 | 0 | 1,62009 | 0,03618921 | 0 | 0,5729923 | 0,824176 | | 1,068286 |
| GSM2261721 | 0 | 1,188984 | 0,071592 | 0,705763 | 0 | 2,28881 | 0,1927191 | 0 | 0,5950066 | 0,809842 | | 1,260576 |
| GSM2261723 | 0 | 0,822407 | 0,02376 | 0,396178 | 0 | 3,377912 | 0,10553007 | 0 | 0,6421687 | 0,767491 | | 0,846167 |
| GSM2261724 | 0 | 0,754453 | 0 | 0,375095 | 0 | 0,820176 | 0,16917731 | 0 | 0,5032948 | 0,865261 | | 0,754453 |
| GSM2261727 | 0 | 0,86947 | 0,000157 | 0,739297 | 0 | 0,813156 | 0,35210826 | 0 | 0,4267358 | 0,905036 | | 0,869627 |
| GSM2261728 | 0 | 0,886498 | 0 | 0,62018 | 0 | 1,973243 | 0,15670444 | 0 | 0,5378819 | 0,842901 | | 0,886498 |
| GSM2261730 | 0 | 0,975358 | 0 | 0,533829 | 0 | 1,143828 | 0,38238616 | 0 | 0,4678298 | 0,883597 | | 0,975358 |
| GSM2261731 | 0 | 0,710287 | 0 | 0,601762 | 0 | 1,350206 | 1,06196951 | 0 | 0,5493692 | 0,835455 | | 0,710287 |
| GSM2261733 | 0 | 0,539116 | 0,010832 | 0,561286 | 0 | 1,244183 | 0,53524498 | 0 | 0,5461101 | 0,837619 | | 0,549947 |
| GSM2261736 | 0 | 0,782602 | 0 | 0,619788 | 0 | 1,472651 | 0,15516873 | 0 | 0,4419507 | 0,897488 | | 0,782602 |
| GSM2261740 | 0 | 0,703655 | 0 | 0,576057 | 0,013777 | 1,374723 | 0,24994155 | 0 | 0,4830352 | 0,874637 | | 0,703655 |
| GSM2261744 | 0 | 0,823726 | 0,037222 | 0,82626 | 0 | 1,614581 | 0,43221829 | 0 | 0,4934046 | 0,868633 | | 0,860947 |
| GSM2261749 | 0 | 0,716183 | 0 | 1,008625 | 0 | 1,991718 | 0,2212506 | 0 | 0,5577714 | 0,829592 | | 0,716183 |
| GSM2261751 | 0 | 1,036691 | 0 | 0,766802 | 0 | 2,714651 | 0,3229368 | 0 | 0,5897116 | 0,809881 | | 1,036691 |
| GSM2261754 | 0 | 0,929729 | 0 | 0,606503 | 0 | 1,096054 | 0,42071517 | 0 | 0,5313333 | 0,849214 | | 0,929729 |
| GSM2261755 | 0 | 0,877843 | 0 | 0,541032 | 0 | 1,404835 | 0,24259664 | 0 | 0,4600442 | 0,887565 | | 0,877843 |
| GSM2261759 | 0 | 1,799296 | 0 | 0,51819 | 0 | 3,76654 | 0 | 0 | 0,5628665 | 0,827744 | | 1,799296 |
| GSM2261761 | 0 | 1,393163 | 0,031521 | 1,064075 | 0 | 1,59715 | 0,74502943 | 0 | 0,3553483 | 0,940864 | | 1,424684 |
| GSM2261764 | 0 | 1,209939 | 0 | 1,109323 | 0 | 1,572261 | 0,60269523 | 0 | 0,483694 | 0,874215 | | 1,209939 |
| GSM2261767 | 0 | 1,888293 | 0 | 1,325339 | 0 | 4,269636 | 0,73372971 | 0 | 0,6105078 | 0,798304 | | 1,888293 |
| GSM2261769 | 0 | 1,118174 | 0,011288 | 1,331906 | 0 | 1,93405 | 0,72861464 | 0 | 0,5389635 | 0,842323 | | 1,129462 |
| GSM2261770 | 0 | 1,087045 | 0 | 1,044993 | 0 | 1,648115 | 0,49285145 | 0 | 0,5190406 | 0,854416 | | 1,087045 |
| GSM2261772 | 0 | 0,870871 | 0 | 0,497607 | 0 | 1,093835 | 0,38025994 | 0 | 0,4421181 | 0,896892 | | 0,870871 |
| GSM2261773 | 0 | 1,067698 | 0 | 0,668575 | 0 | 1,978191 | 0,33530254 | 0 | 0,5601046 | 0,830906 | | 1,067698 |
| GSM2261774 | 0 | 1,061541 | 0 | 0,70286 | 0 | 1,486103 | 0,39710574 | 0 | 0,5008167 | 0,865473 | | 1,061541 |
| GSM2261777 | 0 | 1,360532 | 0,061408 | 0,680954 | 0,021447 | 4,095716 | 0,26850224 | 0 | 0,6777107 | 0,74644 | | 1,42194 |
| GSM2261778 | 0 | 1,428505 | 0 | 1,091514 | 0 | 1,553037 | 0,60705319 | 0 | 0,4872521 | 0,872683 | | 1,428505 |
| GSM2261779 | 0 | 1,022638 | 0 | 0,556212 | 0 | 1,710361 | 0,17206864 | 0 | 0,4701671 | 0,882343 | | 1,022638 |
| GSM2261781 | 0 | 1,446289 | 0,027112 | 0,876303 | 0 | 1,707906 | 0,31187169 | 0 | 0,396338 | 0,920384 | | 1,473401 |
| GSM2261783 | 0 | 1,232568 | 0 | 1,160756 | 0 | 5,496128 | 0 | 0 | 0,6726576 | 0,744748 | | 1,232568 |
| GSM2261784 | 0 | 1,23352 | 0 | 0,664052 | 0 | 0,871835 | 0,55690105 | 0 | 0,3690813 | 0,933556 | | 1,23352 |
| GSM2261785 | 0 | 1,585562 | 0,486096 | 0,841734 | 0 | 2,94737 | 0,13704368 | 0 | 0,5882136 | 0,815089 | | 2,071658 |
| GSM2261786 | 0 | 0,795009 | 0 | 0,707016 | 0 | 2,361391 | 0,83482962 | 0 | 0,591802 | 0,807488 | | 0,795009 |
| GSM2261787 | 0 | 0,731784 | 0,602036 | 0,646718 | 0 | 2,943485 | 0,12147076 | 0 | 0,6046154 | 0,799569 | | 1,333821 |
| GSM2261788 | 0 | 0,666208 | 0,262651 | 0,765017 | 0 | 1,85132 | 0,72841648 | 0 | 0,5807227 | 0,816847 | | 0,928859 |
| GSM2261789 | 0 | 0,980915 | 0,285488 | 0,853806 | 0 | 2,11322 | 0,64468081 | 0 | 0,4291186 | 0,903895 | | 1,266403 |
| GSM2261790 | 0 | 0,938383 | 0,178491 | 0,66425 | 0 | 1,698112 | 0,43030422 | 0 | 0,499551 | 0,865706 | | 1,116874 |
| GSM2261792 | 0 | 1,013187 | 0 | 0,450079 | 0,006935 | 1,493876 | 0,31189654 | 0 | 0,3692487 | 0,935104 | | 1,013187 |
| GSM2261795 | 0 | 0,624147 | 0 | 0,790281 | 0 | 1,874505 | 0,55482687 | 0 | 0,5952146 | 0,805929 | | 0,624147 |
| GSM2261796 | 0 | 1,159373 | 0 | 0,744452 | 0 | 1,64413 | 0,17513222 | 0 | 0,4505615 | 0,892581 | | 1,159373 |
| GSM2261799 | 0 | 0,802469 | 0 | 0,789158 | 0 | 1,719011 | 0,28525474 | 0 | 0,5656993 | 0,826299 | | 0,802469 |
| GSM2261803 | 0 | 1,084783 | 0 | 0,867092 | 0 | 2,674469 | 0,10349239 | 0 | 0,5282532 | 0,848289 | | 1,084783 |
| GSM2261804 | 0 | 0,68776 | 0,069572 | 1,28162 | 0 | 1,808665 | 0,58738012 | 0 | 0,6182628 | 0,791454 | | 0,757333 |
| GSM2261806 | 0 | 0,993857 | 0,00479 | 0,466515 | 0 | 2,13482 | 0,16302104 | 0 | 0,5520775 | 0,834553 | | 0,998647 |
| GSM2261808 | 0 | 0,885413 | 0 | 0,682802 | 0 | 0,842425 | 0,15213522 | 0 | 0,3963517 | 0,921545 | | 0,885413 |
| GSM2261809 | 0 | 0,901648 | 0 | 0,362207 | 0,01859 | 3,056572 | 0 | 0 | 0,6142862 | 0,789484 | | 0,901648 |
| GSM2261810 | 0 | 0,904454 | 0,166751 | 0,666392 | 0 | 1,364954 | 0,55556328 | 0 | 0,4766003 | 0,878267 | | 1,071206 |
| GSM2261811 | 0 | 0,592747 | 0,189355 | 0,502553 | 0 | 1,441858 | 0,28821123 | 0 | 0,5296509 | 0,847947 | | 0,782102 |
| GSM2261813 | 0 | 1,027599 | 0 | 0,512491 | 0 | 1,792435 | 0,7854468 | 0 | 0,3513063 | 0,946729 | | 1,027599 |
| GSM2261814 | 0 | 0,652696 | 0,045357 | 0,723302 | 0 | 1,488072 | 0,76521457 | 0 | 0,5663551 | 0,825365 | | 0,698053 |
| GSM2261819 | 0 | 1,021012 | 0 | 0,517241 | 0 | 1,228205 | 0,222234 | 0 | 0,3695659 | 0,933831 | | 1,021012 |
| GSM2261825 | 0 | 0,968673 | 0 | 0,579133 | 0,015652 | 1,589058 | 0,07631701 | 0 | 0,504717 | 0,863006 | | 0,968673 |
| GSM2261827 | 0 | 0,474065 | 0,258336 | 0,497641 | 0 | 1,949842 | 1,47507408 | 0 | 0,615351 | 0,78935 | | 0,732401 |
| GSM2261829 | 0 | 1,166357 | 0,001287 | 0,653208 | 0 | 2,853716 | 0,00659731 | 0 | 0,5873989 | 0,811199 | | 1,167644 |
| GSM2261831 | 0 | 1,348745 | 0 | 0,509348 | 0 | 2,609963 | 0,28339474 | 0 | 0,5001605 | 0,865971 | | 1,348745 |
| GSM2261833 | 0 | 0,740838 | 0 | 0,676343 | 0 | 2,263092 | 0,05704009 | 0 | 0,6095571 | 0,795336 | | 0,740838 |
| GSM2261835 | 0 | 0,907218 | 0 | 0,512595 | 0 | 0,763331 | 0,3273741 | 0 | 0,4044414 | 0,916024 | | 0,907218 |
| GSM2261836 | 0 | 1,040414 | 0,031708 | 0,427848 | 0 | 1,945086 | 0 | 0 | 0,5172621 | 0,855992 | | 1,072121 |
| GSM2261838 | 0 | 0,490442 | 0 | 0,705252 | 0 | 0,779716 | 0,28584523 | 0 | 0,4601186 | 0,887492 | | 0,490442 |
| GSM2261846 | 0 | 0,892727 | 0,044522 | 0,665395 | 0 | 1,156069 | 0,10766994 | 0 | 0,5405976 | 0,842796 | | 0,937248 |
| GSM2261847 | 0 | 0,982226 | 0,069725 | 0,936865 | 0 | 2,366067 | 0,00513153 | 0,02 | 0,2070456 | 1,023101 | | 1,051951 |
| GSM2261848 | 0 | 0,479711 | 0 | 0,42732 | 0 | 1,391436 | 0,08601664 | 0 | 0,5425085 | 0,839174 | | 0,479711 |
| GSM2261852 | 0 | 0,922141 | 0,090897 | 0,916282 | 0 | 2,554124 | 0,50673746 | 0 | 0,3172743 | 0,966598 | | 1,013038 |
| GSM2261854 | 0 | 1,425317 | 0 | 0,848398 | 0 | 2,849185 | 0 | 0 | 0,5745233 | 0,821133 | | 1,425317 |
| GSM2261857 | 0 | 1,622493 | 0 | 0,535723 | 0 | 2,046492 | 0,16935621 | 0 | 0,4356592 | 0,901352 | | 1,622493 |
| GSM2261864 | 0 | 0,756955 | 0 | 0,704938 | 0 | 1,520701 | 0,46097389 | 0 | 0,4304391 | 0,90366 | | 0,756955 |
| GSM2261865 | 0 | 0,819474 | 0,046235 | 0,474986 | 0 | 1,896463 | 0,2019316 | 0 | 0,5624251 | 0,828012 | | 0,865708 |
| GSM2261868 | 0 | 0,723974 | 0 | 0,451045 | 0 | 2,270527 | 0 | 0 | 0,6524915 | 0,763674 | | 0,723974 |
| GSM2261870 | 0 | 1,114556 | 0 | 0,537834 | 0 | 2,020518 | 0,38229382 | 0 | 0,4699791 | 0,882658 | | 1,114556 |
| GSM2261871 | 0 | 0,811533 | 0 | 0,878937 | 0 | 1,803111 | 0,25309158 | 0 | 0,4814587 | 0,875448 | | 0,811533 |
| GSM2261872 | 0 | 0,821395 | 0,078663 | 0,629448 | 0 | 1,63189 | 0,34076448 | 0 | 0,5136798 | 0,857482 | | 0,900057 |
| GSM2261873 | 0 | 0,799093 | 0 | 0,330526 | 0 | 1,736508 | 0,01856096 | 0 | 0,48286 | 0,87638 | | 0,799093 |
| GSM2261879 | 0 | 0,779783 | 0 | 0,289815 | 0 | 0,93112 | 0,32074511 | 0 | 0,4673105 | 0,884427 | | 0,779783 |
| GSM2261880 | 0,001842 | 0,775632 | 0 | 0,563289 | 0 | 1,095932 | 0,0749932 | 0 | 0,330597 | 0,954901 | | 0,775632 |
| GSM2261884 | 0 | 1,011861 | 0 | 0,311047 | 0 | 2,490664 | 0 | 0 | 0,530529 | 0,847949 | | 1,011861 |
| GSM2261886 | 0 | 1,419208 | 0,214501 | 1,015447 | 0 | 3,038787 | 0,48720368 | 0 | 0,5152799 | 0,856498 | | 1,63371 |
| GSM2261887 | 0 | 1,130602 | 0 | 0,555602 | 0 | 1,930872 | 0,23862873 | 0 | 0,4315346 | 0,903496 | | 1,130602 |
| GSM2261892 | 0 | 0,905053 | 0 | 1,047041 | 0 | 2,773298 | 0,11343137 | 0 | 0,5826326 | 0,813359 | | 0,905053 |
| GSM2261895 | 0 | 0,655694 | 0 | 0,811647 | 0 | 1,220552 | 0,30711261 | 0 | 0,5276288 | 0,848669 | | 0,655694 |
| GSM2261897 | 0 | 1,276972 | 0 | 0,560453 | 0 | 1,330096 | 0,19659745 | 0 | 0,4217677 | 0,907771 | | 1,276972 |
| GSM2261898 | 0 | 1,021611 | 0 | 0,664871 | 0 | 2,063666 | 0,28023076 | 0 | 0,5271453 | 0,849692 | | 1,021611 |
| GSM2261899 | 0,004456 | 1,247001 | 0 | 0,794507 | 0 | 1,138212 | 0,43270817 | 0 | 0,4587644 | 0,8882 | | 1,247001 |
| GSM2261901 | 0 | 2,316034 | 0,304823 | 0,308564 | 0 | 3,897522 | 0 | 0 | 0,7196436 | 0,728777 | | 2,620857 |

| | | | | | | | | | | | | |
|---|---|---|---|---|---|---|---|---|---|---|---|---|
| GSM2261902 | 0 | 1,084188 | 0,346441 | 0,877449 | 0 | 2,936804 | 0 | 0 | 0,6840242 | 0,74898 | | 1,430629 |
| GSM2261903 | 0 | 1,544813 | 0,006758 | 1,798645 | 0 | 5,431828 | 0,09812484 | 0 | 0,5790735 | 0,814911 | | 1,551571 |
| GSM2261904 | 0 | 0,77029 | 0,138003 | 0,484468 | 0 | 1,566017 | 0,24246181 | 0 | 0,4815911 | 0,87569 | | 0,908293 |
| GSM2261906 | 0 | 1,258689 | 0 | 0,731254 | 0 | 1,699025 | 0,33071415 | 0 | 0,4696933 | 0,882469 | | 1,258689 |
| GSM2261908 | 0 | 0,83413 | 0,084315 | 0,605088 | 0 | 1,039055 | 0,45750117 | 0 | 0,441654 | 0,896779 | | 0,918444 |
| GSM2261909 | 0 | 0,989088 | 0 | 0,665509 | 0 | 1,326014 | 0,26402308 | 0 | 0,4270284 | 0,904727 | | 0,989088 |
| GSM2261910 | 0 | 0,477237 | 0,16731 | 0,621774 | 0 | 1,325781 | 0,54814678 | 0 | 0,5534735 | 0,833148 | | 0,644547 |
| GSM2261912 | 0 | 1,006144 | 0,027949 | 0,83319 | 0 | 1,857474 | 0,25986759 | 0 | 0,4619535 | 0,886329 | | 1,034093 |
| GSM2261913 | 0 | 1,196698 | 0 | 0,774374 | 0 | 1,534395 | 0,56775014 | 0 | 0,4884523 | 0,872292 | | 1,196698 |
| GSM2261916 | 0 | 0,679217 | 0,001404 | 0,759889 | 0 | 2,08819 | 0,21797778 | 0 | 0,5911319 | 0,808104 | | 0,680621 |
| GSM2261917 | 0 | 0,944026 | 0 | 0,681922 | 0 | 1,457985 | 0,26135023 | 0 | 0,4566155 | 0,889185 | | 0,944026 |
| GSM2261918 | 0 | 0,819083 | 0 | 0,86938 | 0 | 2,19127 | 0,16841883 | 0 | 0,625186 | 0,786979 | | 0,819083 |
| GSM2261920 | 0 | 1,181169 | 0 | 0,505217 | 0 | 2,3204 | 0,18838617 | 0 | 0,5645381 | 0,827573 | | 1,181169 |
| GSM2261921 | 0 | 0,642343 | 0 | 0,652193 | 0 | 1,330461 | 0,22352897 | 0 | 0,4701074 | 0,88181 | | 0,642343 |
| GSM2261922 | 0 | 0,888919 | 0 | 0,926142 | 0 | 1,563562 | 0,21566285 | 0,27 | 0,0982178 | 1,069497 | | 0,888919 |
| GSM2261923 | 0 | 0,727003 | 0 | 0,478588 | 0 | 1,402492 | 0,13395364 | 0 | 0,5005883 | 0,865071 | | 0,727003 |
| GSM2261924 | 0 | 0,846174 | 0,539701 | 0,501103 | 0 | 2,724716 | 0 | 0 | 0,5948551 | 0,806592 | | 1,385875 |
| GSM2261925 | 0 | 0,809248 | 0,01167 | 0,504473 | 0 | 1,413886 | 0,25226829 | 0 | 0,530595 | 0,848364 | | 0,820918 |
| GSM2261928 | 0 | 0,960463 | 0 | 1,487887 | 0 | 2,654151 | 1,66861423 | 0 | 0,6368631 | 0,778209 | | 0,960463 |
| GSM2261935 | 0 | 0,965051 | 0,140334 | 0,48699 | 0 | 2,385694 | 0,04483987 | 0 | 0,5993144 | 0,80421 | | 1,105385 |
| GSM2261939 | 0 | 0,822275 | 0 | 0,641335 | 0 | 2,45101 | 0,59685311 | 0 | 0,6501716 | 0,768159 | | 0,822275 |
| GSM2261945 | 0 | 1,132871 | 0,08989 | 0,697772 | 0 | 2,362711 | 0,23277428 | 0 | 0,5158019 | 0,856299 | | 1,222761 |
| GSM2261951 | 0 | 0,759835 | 0 | 0,425621 | 0 | 1,264041 | 0,30906824 | 0 | 0,4501185 | 0,89308 | | 0,759835 |
| GSM2261953 | 0 | 1,476843 | 0 | 0,686491 | 0 | 1,678512 | 0,2271987 | 0 | 0,4583452 | 0,888939 | | 1,476843 |
| GSM2261955 | 0 | 0,897569 | 0 | 0,61749 | 0 | 1,749637 | 0 | 0 | 0,5246564 | 0,851012 | | 0,897569 |
| GSM2261957 | 0 | 0,768126 | 0 | 0,309946 | 0 | 1,114386 | 0,23997832 | 0 | 0,4695108 | 0,883046 | | 0,768126 |
| GSM2261962 | 0 | 0,994896 | 0,146666 | 0,772693 | 0 | 3,151204 | 0,41848077 | 0 | 0,535473 | 0,843627 | | 1,141562 |
| GSM2261963 | 0 | 0,950046 | 0 | 0,770058 | 0 | 1,117025 | 0,31894724 | 0 | 0,4444857 | 0,895461 | | 0,950046 |
| GSM2261964 | 0 | 0,912079 | 0,242453 | 1,084314 | 0 | 2,796534 | 0,09255368 | 0 | 0,5097905 | 0,858975 | | 1,154531 |
| GSM2261966 | 0 | 1,113108 | 0 | 0,780292 | 0 | 1,134848 | 0,46436588 | 0 | 0,3826069 | 0,927091 | | 1,113108 |
| GSM2261967 | 0 | 0,725047 | 0 | 0,892022 | 0 | 1,784642 | 0,25931823 | 0 | 0,4510051 | 0,892944 | | 0,725047 |
| GSM2261976 | 0 | 1,06205 | 0 | 0,385143 | 0,020712 | 1,218181 | 0,23572282 | 0 | 0,4969954 | 0,868953 | | 1,06205 |
| GSM2261977 | 0 | 1,468317 | 0 | 1,08309 | 0 | 3,201934 | 0,27803523 | 0 | 0,5332475 | 0,845713 | | 1,468317 |
| GSM2261985 | 0 | 1,030853 | 0 | 0,720145 | 0 | 4,23273 | 0 | 0 | 0,725332 | 0,702014 | | 1,030853 |
| GSM2261988 | 0 | 1,122696 | 0 | 1,007256 | 0 | 1,356064 | 0,79437342 | 0 | 0,4155771 | 0,910635 | | 1,122696 |
| GSM2261989 | 0 | 0,8714 | 0 | 0,652034 | 0 | 1,195855 | 0,17599759 | 0 | 0,3942484 | 0,92196 | | 0,8714 |
| GSM2261991 | 0 | 0,732335 | 0 | 0,510898 | 0 | 1,024329 | 0,36265317 | 0 | 0,397588 | 0,919879 | | 0,732335 |
| GSM2261992 | 0 | 1,19863 | 0 | 0,637161 | 0 | 1,262903 | 0,31409884 | 0 | 0,3866439 | 0,924998 | | 1,19863 |
| GSM2261996 | 0 | 0,661072 | 0,409666 | 0,460017 | 0 | 1,56458 | 0,67336413 | 0 | 0,4878526 | 0,871913 | | 1,070738 |
| GSM2261998 | 0 | 1,19384 | 0 | 1,050792 | 0 | 1,200389 | 0,34492663 | 0,32 | 0,0869392 | 1,071926 | | 1,19384 |
| GSM2262001 | 0 | 0,988677 | 0 | 0,513125 | 0 | 1,167649 | 0,21339416 | 0 | 0,4645449 | 0,885459 | | 0,988677 |
| GSM2262003 | 0 | 1,154311 | 0,124608 | 0,687246 | 0 | 1,557786 | 1,26336332 | 0 | 0,5558548 | 0,833846 | | 1,278919 |
| GSM2262004 | 0 | 0,854783 | 0 | 0,556237 | 0 | 0,869571 | 0,44685177 | 0 | 0,421385 | 0,907345 | | 0,854783 |
| GSM2262007 | 0 | 0,874289 | 0,004594 | 0,648578 | 0 | 2,281474 | 0,09873271 | 0 | 0,5794066 | 0,816283 | | 0,878884 |
| GSM2262013 | 0 | 0,955026 | 0,096953 | 0,666804 | 0 | 1,788337 | 0,2049772 | 0 | 0,4700908 | 0,881977 | | 1,051979 |
| GSM2262017 | 0 | 0,75196 | 0,031199 | 0,503901 | 0 | 1,735243 | 0,25882795 | 0 | 0,4450849 | 0,896743 | | 0,78316 |
| GSM2262020 | 0 | 0,75915 | 0 | 0,626022 | 0 | 1,080384 | 0,37566128 | 0 | 0,4629899 | 0,885545 | | 0,75915 |
| GSM2262023 | 0 | 1,044303 | 0 | 0,71441 | 0 | 1,131131 | 0,36551717 | 0 | 0,3747954 | 0,931054 | | 1,044303 |
| GSM2262025 | 0 | 0,997389 | 0 | 0,688168 | 0 | 2,25841 | 0,20790942 | 0 | 0,6199034 | 0,792073 | | 0,997389 |
| GSM2262026 | 0 | 0,748002 | 0 | 0,70797 | 0 | 0,828847 | 0,45421099 | 0 | 0,4048642 | 0,916389 | | 0,748002 |
| GSM2262028 | 0 | 1,211892 | 0 | 0,79185 | 0 | 1,537028 | 0,41121897 | 0 | 0,451529 | 0,891864 | | 1,211892 |
| GSM2262030 | 0 | 0,697469 | 0 | 0,470002 | 0 | 0,899182 | 0,12749642 | 0 | 0,4604197 | 0,887231 | | 0,697469 |
| GSM2262031 | 0 | 1,404471 | 0 | 1,084138 | 0 | 1,506572 | 0,21109153 | 0 | 0,3336406 | 0,953516 | | 1,404471 |
| GSM2262033 | 0 | 0,668875 | 0 | 0,503218 | 0 | 1,251471 | 0,05155271 | 0 | 0,4415626 | 0,897947 | | 0,668875 |
| GSM2262034 | 0 | 0,992888 | 0,007238 | 0,577963 | 0 | 1,121575 | 0,16594617 | 0 | 0,3632424 | 0,937107 | | 1,000126 |
| GSM2262035 | 0 | 0,94901 | 0 | 0,844395 | 0 | 1,495759 | 0,33720228 | 0 | 0,47578 | 0,878573 | | 0,94901 |
| GSM2262041 | 0 | 0,733097 | 0 | 0,883498 | 0 | 1,079077 | 0,37248603 | 0 | 0,5056593 | 0,861458 | | 0,733097 |
| GSM2262042 | 0 | 1,115726 | 0,202221 | 0,577477 | 0 | 3,458421 | 0,40242242 | 0 | 0,5178231 | 0,855071 | | 1,317946 |
| GSM2262044 | 0 | 0,96761 | 0,051913 | 0,595162 | 0 | 1,159649 | 0,21838261 | 0 | 0,4761532 | 0,878985 | | 1,019523 |
| GSM2262046 | 0 | 0,78126 | 0 | 0,528244 | 0 | 0,810816 | 0,20255851 | 0 | 0,3509421 | 0,943222 | | 0,78126 |
| GSM2262049 | 0 | 1,997731 | 0,09569 | 0,637218 | 0 | 10,41359 | 0 | 0 | 0,6882746 | 0,726453 | | 2,093421 |
| GSM2262051 | 0 | 0,891794 | 0 | 0,366411 | 0,022916 | 0,992199 | 0,39197547 | 0 | 0,3803882 | 0,928085 | | 0,891794 |
| GSM2262053 | 0 | 0,918094 | 0 | 0,790105 | 0 | 0,755574 | 0,55599042 | 0 | 0,3749055 | 0,932024 | | 0,918094 |
| GSM2262059 | 0 | 1,279507 | 0,012219 | 1,03561 | 0 | 3,202523 | 0,07999983 | 0 | 0,5463267 | 0,837194 | | 1,291727 |
| GSM2262060 | 0 | 0,740619 | 0 | 0,579549 | 0 | 0,874912 | 0,1554756 | 0 | 0,3567773 | 0,941196 | | 0,740619 |
| GSM2262061 | 0 | 0,854742 | 0 | 0,95183 | 0 | 1,434858 | 0,30856822 | 0 | 0,4351719 | 0,900894 | | 0,854742 |
| GSM2262063 | 0 | 1,238529 | 0 | 0,568641 | 0 | 1,411401 | 0,08188231 | 0 | 0,3871657 | 0,925586 | | 1,238529 |
| GSM2262064 | 0 | 0,5772 | 0,021868 | 0,535378 | 0 | 1,367819 | 0,13885603 | 0 | 0,4257767 | 0,90712 | | 0,599068 |
| GSM2262067 | 0 | 0,638993 | 0 | 0,304798 | 0,00733 | 1,313501 | 0,13443456 | 0 | 0,5007263 | 0,865277 | | 0,638993 |
| GSM2262068 | 0 | 0,868868 | 0 | 0,429095 | 0 | 1,46519 | 0,25597316 | 0 | 0,4565132 | 0,889888 | | 0,868868 |
| GSM2262077 | 0 | 0,878206 | 0 | 0,413007 | 0,010621 | 0,884433 | 0,31025454 | 0 | 0,3212869 | 0,95636 | | 0,878206 |
| GSM2262078 | 0 | 0,915356 | 0 | 0,599699 | 0 | 0,943881 | 0,41025768 | 0 | 0,4866052 | 0,87344 | | 0,915356 |
| GSM2262080 | 0 | 1,082445 | 0,007507 | 0,706337 | 0 | 1,384505 | 0,31152702 | 0 | 0,3586326 | 0,939376 | | 1,089952 |
| GSM2262082 | 0 | 0,786648 | 0,034523 | 0,620722 | 0 | 1,330271 | 0,13141001 | 0 | 0,4364645 | 0,900131 | | 0,821171 |
| GSM2262083 | 0 | 0,5536 | 0 | 0,625849 | 0 | 1,341952 | 0,28198768 | 0 | 0,5079103 | 0,860149 | | 0,5536 |
| GSM2262086 | 0 | 1,251756 | 0 | 0,665837 | 0,013681 | 2,562245 | 0,26586461 | 0 | 0,5383725 | 0,843068 | | 1,251756 |
| GSM2262091 | 0 | 0,837325 | 0 | 0,454944 | 0 | 2,154776 | 0 | 0 | 0,4982298 | 0,867255 | | 0,837325 |
| GSM2262094 | 0 | 0,830983 | 0,236813 | 0,526107 | 0 | 1,92497 | 0,19863178 | 0 | 0,5159647 | 0,856076 | | 1,067796 |
| GSM2262098 | 0 | 0,917855 | 0 | 0,322578 | 0 | 1,347055 | 0,07497628 | 0 | 0,4216614 | 0,909028 | | 0,917855 |
| GSM2262102 | 0 | 1,892167 | 0,244372 | 1,195565 | 0 | 6,190982 | 0 | 0 | 0,5750324 | 0,817855 | | 2,136539 |
| GSM2262103 | 0 | 0,556983 | 0 | 0,463146 | 0 | 0,77447 | 0,39813433 | 0 | 0,4592268 | 0,887574 | | 0,556983 |
| GSM2262104 | 0 | 1,003895 | 0,074255 | 0,736957 | 0 | 2,899033 | 0,86376635 | 0 | 0,5562343 | 0,830695 | | 1,07815 |
| GSM2262105 | 0 | 1,1104 | 0 | 0,574988 | 0 | 1,98755 | 0,29232245 | 0 | 0,5068106 | 0,861925 | | 1,1104 |

| | | | | | | | | | | | | |
|---|---|---|---|---|---|---|---|---|---|---|---|---|
| GSM2262107 | 0 | 0,768525 | 0,188627 | 0,567904 | 0 | 1,10875 | 0,33022187 | 0 | 0,5167811 | 0,856551 | | 0,957153 |
| GSM2262116 | 0,071006 | 0,503331 | 0,104396 | 0,508838 | 0,029971 | 3,229301 | 0,37066845 | 0 | 0,6550247 | 0,757527 | | 0,607727 |
| GSM2262117 | 0 | 1,347192 | 0 | 0,809178 | 0 | 1,866825 | 0,53564069 | 0 | 0,47924 | 0,877283 | | 1,347192 |
| GSM2262119 | 0 | 1,482336 | 0,070913 | 0,462039 | 0 | 3,628647 | 0 | 0 | 0,661068 | 0,760206 | | 1,553249 |
| GSM2262120 | 0 | 1,014328 | 0,002695 | 0,676499 | 0 | 1,113333 | 0,24111748 | 0 | 0,470707 | 0,881799 | | 1,017022 |
| GSM2262122 | 0 | 1,205933 | 0 | 0,436243 | 0 | 1,033776 | 0,39749843 | 0 | 0,4097048 | 0,91372 | | 1,205933 |
| GSM2262123 | 0 | 1,214183 | 0,013542 | 0,80705 | 0 | 2,152357 | 0,30007557 | 0 | 0,5400997 | 0,84275 | | 1,227725 |
| GSM2262124 | 0 | 1,49866 | 0 | 0,481628 | 0 | 2,407643 | 0 | 0 | 0,4419429 | 0,898983 | | 1,49866 |
| GSM2262126 | 0 | 0,666995 | 0,612787 | 0,684053 | 0 | 3,347268 | 0 | 0 | 0,6633705 | 0,756865 | | 1,279782 |
| GSM2262127 | 0 | 1,044737 | 0 | 0,655605 | 0 | 1,599047 | 0,06828787 | 0 | 0,4311099 | 0,903232 | | 1,044737 |
| GSM2262131 | 0 | 1,010467 | 0 | 0,760769 | 0 | 1,490848 | 0,33485372 | 0 | 0,4406624 | 0,897578 | | 1,010467 |
| GSM2262138 | 0 | 0,906831 | 0 | 0,467936 | 0 | 1,457231 | 0,07908132 | 0 | 0,3899282 | 0,925325 | | 0,906831 |
| GSM2262144 | 0 | 1,531685 | 0 | 0,723081 | 0 | 2,038137 | 0,17801687 | 0 | 0,4193203 | 0,909286 | | 1,531685 |
| GSM2262146 | 0 | 1,192792 | 0 | 0,471769 | 9,15E-05 | 2,469551 | 0,05780982 | 0 | 0,5212913 | 0,853405 | | 1,192792 |
| GSM2262148 | 0,0125 | 0,858128 | 0 | 0,539764 | 0,044621 | 1,584324 | 0,15694091 | 0 | 0,5410698 | 0,841946 | | 0,858128 |
| GSM2262149 | 0 | 0,632833 | 0,220026 | 0,477891 | 0 | 1,95762 | 0,31902512 | 0 | 0,5743293 | 0,819805 | | 0,852858 |
| GSM2262152 | 0 | 1,392357 | 0 | 0,97428 | 0 | 1,881563 | 0,24865476 | 0 | 0,4589056 | 0,888003 | | 1,392357 |
| GSM2262159 | 0 | 0,939856 | 0 | 0,776889 | 0 | 1,462169 | 0,1617382 | 0 | 0,3755406 | 0,932241 | | 0,939856 |
| GSM2262161 | 0 | 1,22301 | 0 | 1,030175 | 0,009888 | 2,709068 | 0,1213651 | 0 | 0,5607044 | 0,828933 | | 1,22301 |
| GSM2262162 | 0 | 0,733395 | 0,031396 | 0,745858 | 0 | 2,000286 | 0,18389606 | 0 | 0,5826182 | 0,814449 | | 0,764791 |
| GSM2262163 | 0 | 0,967567 | 0 | 0,794056 | 0 | 1,654689 | 0,06356263 | 0 | 0,6166922 | 0,796243 | | 0,967567 |
| GSM2262164 | 0 | 1,071235 | 0 | 0,731609 | 0 | 1,849707 | 0,01993916 | 0 | 0,4566785 | 0,889652 | | 1,071235 |
| GSM2262166 | 0 | 1,187301 | 0,10715 | 0,815592 | 0 | 1,569603 | 0,24793154 | 0 | 0,50214 | 0,864665 | | 1,294451 |
| GSM2262168 | 0 | 0,646912 | 0 | 0,541411 | 0 | 2,363261 | 0,05305185 | 0 | 0,5470404 | 0,836218 | | 0,646912 |
| GSM2262173 | 0 | 0,941787 | 0 | 0,519986 | 0,006524 | 1,046827 | 0,11989811 | 0 | 0,3596405 | 0,938995 | | 0,941787 |
| GSM2262175 | 0 | 1,60675 | 0,082762 | 0,566384 | 0 | 4,05011 | 0 | 0 | 0,5932091 | 0,806795 | | 1,689511 |
| GSM2262176 | 0 | 0,57794 | 0 | 0,281887 | 0 | 1,262975 | 0,30704015 | 0 | 0,4821169 | 0,876197 | | 0,57794 |
| GSM2262177 | 0 | 0,667487 | 0 | 0,389047 | 0 | 1,443823 | 0,15583942 | 0 | 0,5015412 | 0,864605 | | 0,667487 |
| GSM2262179 | 0 | 1,318548 | 0 | 1,314434 | 0 | 2,218726 | 0,3232047 | 0 | 0,5280591 | 0,848883 | | 1,318548 |
| GSM2262180 | 0 | 1,01288 | 0,000414 | 0,741382 | 0 | 1,737297 | 0,23638062 | 0 | 0,503316 | 0,863555 | | 1,013294 |
| GSM2262183 | 0 | 1,076304 | 0 | 0,421185 | 0 | 0,7052 | 0,20649657 | 0 | 0,3437489 | 0,946435 | | 1,076304 |
| GSM2262185 | 0 | 1,265058 | 0 | 0,635848 | 0 | 1,441003 | 0,11151311 | 0 | 0,384678 | 0,926699 | | 1,265058 |
| GSM2262186 | 0 | 1,134208 | 0 | 0,634909 | 0 | 2,054652 | 0,1954571 | 0 | 0,4440619 | 0,89675 | | 1,134208 |
| GSM2262187 | 0 | 1,668276 | 0,405238 | 1,12844 | 0 | 3,629716 | 0 | 0 | 0,5882715 | 0,812983 | | 2,073513 |
| GSM2262191 | 0 | 1,329915 | 0 | 0,791147 | 0 | 1,681889 | 0,24841034 | 0 | 0,3040853 | 0,966383 | | 1,329915 |
| GSM2262193 | 0 | 0,876506 | 0,051225 | 0,914807 | 0 | 2,090565 | 0,06071233 | 0 | 0,5098974 | 0,859023 | | 0,927731 |
| GSM2262194 | 0 | 0,840302 | 0 | 0,540308 | 0 | 1,357506 | 0,25992104 | 0 | 0,4476145 | 0,894178 | | 0,840302 |
| GSM2262196 | 0 | 1,336333 | 0,530078 | 0,419921 | 0 | 2,29299 | 0,11741735 | 0 | 0,5938764 | 0,812888 | | 1,866411 |
| GSM2262200 | 0 | 1,293634 | 0 | 1,316874 | 0 | 2,698003 | 0 | 0 | 0,4214932 | 0,909665 | | 1,293634 |
| GSM2262201 | 0 | 0,942295 | 0 | 0,729397 | 0 | 1,881473 | 0,07116336 | 0 | 0,4604516 | 0,887635 | | 0,942295 |
| GSM2262203 | 0 | 1,199868 | 0,111221 | 0,843393 | 0 | 2,35552 | 0,43464218 | 0 | 0,4199396 | 0,909243 | | 1,311088 |
| GSM2262211 | 0 | 1,111797 | 0 | 0,575002 | 0 | 2,136058 | 0 | 0 | 0,4619132 | 0,887522 | | 1,111797 |
| GSM2262213 | 0 | 1,164335 | 0 | 0,836071 | 0 | 0,884448 | 0,28234096 | 0 | 0,3136758 | 0,963696 | | 1,164335 |
| GSM2262215 | 0 | 0,992825 | 0 | 0,701233 | 0 | 1,615962 | 0,03709813 | 0 | 0,3859745 | 0,927401 | | 0,992825 |
| GSM2262216 | 0 | 0,696042 | 0 | 0,520955 | 0 | 1,360147 | 0,08864406 | 0 | 0,401476 | 0,919817 | | 0,696042 |
| GSM2262219 | 0 | 0,929801 | 0 | 0,947784 | 0 | 4,6358 | 0 | 0 | 0,5214516 | 0,854933 | | 0,929801 |
| GSM2262221 | 0 | 1,37351 | 0,076271 | 0,859874 | 0 | 4,970959 | 1,09744861 | 0 | 0,6937203 | 0,732027 | | 1,449781 |
| GSM2262224 | 0 | 0,717845 | 0,022138 | 0,420624 | 0 | 2,096364 | 0,19898349 | 0 | 0,4893891 | 0,872758 | | 0,739983 |
| GSM2262227 | 0 | 0,750154 | 0,032509 | 1,527802 | 0,148816 | 3,387942 | 2,46535159 | 0 | 0,6385374 | 0,77279 | | 0,782663 |
| GSM2262228 | 0 | 1,045973 | 0 | 0,505603 | 0 | 2,133714 | 0,22085948 | 0 | 0,5490932 | 0,836705 | | 1,045973 |
| GSM2262231 | 0 | 1,168408 | 0,109973 | 0,802239 | 0 | 6,55247 | 2,34352053 | 0 | 0,6988897 | 0,720178 | | 1,278381 |
| GSM2262235 | 0 | 1,212078 | 0,005082 | 0,66673 | 0 | 2,305428 | 0,25445623 | 0 | 0,4779209 | 0,878122 | | 1,21716 |
| GSM2262237 | 0 | 1,738872 | 0,145395 | 0,572887 | 0 | 6,78856 | 0 | 0 | 0,6451119 | 0,765157 | | 1,884268 |
| GSM2262238 | 0 | 0,821748 | 0 | 1,018054 | 0 | 1,417356 | 0,65649893 | 0 | 0,40348 | 0,918029 | | 0,821748 |
| GSM2262239 | 0 | 0,936137 | 0 | 0,725308 | 0 | 1,614085 | 0,17524089 | 0 | 0,4315011 | 0,90292 | | 0,936137 |
| GSM2262241 | 0 | 0,999033 | 0 | 1,02416 | 0 | 2,890568 | 0,84362642 | 0 | 0,4644185 | 0,886191 | | 0,999033 |
| GSM2262242 | 0 | 1,19374 | 0 | 0,684329 | 0 | 1,219231 | 0,18858824 | 0 | 0,3867926 | 0,925352 | | 1,19374 |
| GSM2262243 | 0 | 1,264235 | 0 | 0,951588 | 0 | 1,316518 | 0,69980172 | 0 | 0,4067348 | 0,914912 | | 1,264235 |
| GSM2262246 | 0 | 1,279523 | 0 | 0,351899 | 0,035073 | 2,877814 | 0 | 0 | 0,5904901 | 0,809188 | | 1,279523 |
| GSM2262251 | 0 | 0,9681 | 0 | 0,668438 | 0 | 1,127431 | 0,39926092 | 0 | 0,3741888 | 0,931335 | | 0,9681 |
| GSM2262253 | 0 | 0,576111 | 0 | 0,35261 | 0 | 0,424801 | 0,12834332 | 0 | 0,3323123 | 0,953218 | | 0,576111 |
| GSM2262256 | 0 | 1,445917 | 0 | 0,851706 | 0 | 1,416875 | 0,25477772 | 0 | 0,3754936 | 0,931012 | | 1,445917 |
| GSM2262258 | 0 | 0,915204 | 0 | 0,733283 | 0 | 1,460124 | 0,65723919 | 0 | 0,4600763 | 0,88724 | | 0,915204 |
| GSM2262264 | 0 | 1,105952 | 0 | 0,736074 | 0,027525 | 1,254984 | 0,04205874 | 0 | 0,4431298 | 0,896665 | | 1,105952 |
| GSM2262265 | 0 | 1,255857 | 0 | 0,668201 | 0 | 1,321122 | 0,4982664 | 0 | 0,4040753 | 0,916165 | | 1,255857 |
| GSM2262266 | 0 | 1,164688 | 0 | 1,03678 | 0 | 1,532612 | 0,46068021 | 0 | 0,4296568 | 0,903319 | | 1,164688 |
| GSM2262268 | 0 | 0,883083 | 0 | 0,627582 | 0 | 1,028859 | 0,38824034 | 0 | 0,4538456 | 0,890527 | | 0,883083 |
| GSM2262270 | 0 | 0,8513 | 0 | 0,480793 | 0 | 0,933116 | 0,16291981 | 0 | 0,3166383 | 0,959711 | | 0,8513 |
| GSM2262274 | 0 | 1,261371 | 0 | 0,668132 | 0 | 2,172941 | 0 | 0 | 0,4404346 | 0,898985 | | 1,261371 |
| GSM2262276 | 0 | 0,785829 | 0 | 0,516969 | 0 | 1,270146 | 0,25911483 | 0 | 0,4626012 | 0,88609 | | 0,785829 |
| GSM2262282 | 0 | 0,899365 | 0 | 0,672457 | 0 | 1,708943 | 0 | 0 | 0,4254132 | 0,907108 | | 0,899365 |
| GSM2262290 | 0 | 0,755577 | 0 | 0,600102 | 0 | 1,788019 | 0,28925185 | 0 | 0,5509595 | 0,834812 | | 0,755577 |
| GSM2262293 | 0 | 1,097832 | 0 | 1,698028 | 0 | 1,665833 | 0,42648811 | 0 | 0,4097772 | 0,917322 | | 1,097832 |
| GSM2262297 | 0 | 1,216532 | 0 | 0,748998 | 0 | 2,307905 | 0,0224005 | 0 | 0,4349881 | 0,902066 | | 1,216532 |
| GSM2262303 | 0 | 0,586272 | 0,067519 | 0,592911 | 0 | 1,591799 | 0,33674886 | 0 | 0,5506549 | 0,834665 | | 0,65379 |
| GSM2262305 | 0 | 1,24077 | 0 | 0,969494 | 0 | 2,130675 | 0 | 0 | 0,4668311 | 0,883927 | | 1,24077 |
| GSM2262307 | 0 | 1,30512 | 0 | 0,812009 | 0 | 1,83351 | 0,11917472 | 0 | 0,5112922 | 0,859671 | | 1,30512 |
| GSM2262311 | 0 | 0,990851 | 0 | 0,743717 | 0,043452 | 1,082928 | 0,1556595 | 0 | 0,3985158 | 0,919988 | | 0,990851 |
| GSM2262312 | 0 | 0,95132 | 0 | 0,576254 | 0 | 0,957285 | 0,16043575 | 0 | 0,4177022 | 0,909678 | | 0,95132 |
| GSM2262313 | 0 | 0,890586 | 0 | 0,6426 | 0 | 1,26193 | 0,26313395 | 0 | 0,4714277 | 0,8812 | | 0,890586 |
| **average** | **0,000275** | **1,020955** | **0,041167** | **0,711592** | **0,001856** | **2,005664** | **0,28087843** | | | | | |
