## Supplemental Table 4 for "EphA2 and Phosphoantigen-Mediated Selective Killing of Medulloblastoma by γδT Cells Preserves Neuronal and Stem Cell Integrity"

**Table S4**

Boutin et al.

**Ephrin-A2 and Phosphoantigen-Mediated Selective Killing of Medulloblastoma by γδT Cells Preserves Neuronal and Stem Cell Integrity**

| CDR3 analysis | | | | | | | | | | |
|---|---|---|---|---|---|---|---|---|---|---|
| **Barcode** | **cell_type** | **V_chain1** | **V_chain2** | **J_chain1** | **J_chain2** | **C_chain1** | **C_chain2** | **CDR3_chain1** | **CDR3_chain2** | **Note** |
| 1066_GTGAGGACACAAGGTG | abT | TRBV7-9 | * | TRBJ2-7 | * | * | * | CASSSKAGGFYEQYF | * | |
| 1195_AGAAATGGTGGTGATG | abT | TRBV6-5 | * | TRBJ2-3 | * | * | * | CASQRAGAADTQYF | * | |
| 1224_CGAAGTTGTAGCACAG | abT | TRBV7-6 | * | TRBJ2-2 | * | * | * | CASGYASGELFF | * | |
| 1224_GGGATCCCACAGAGCA | abT | TRBV7-2 | * | TRBJ2-1 | * | TRBC2 | * | CASAIEAGAPNEQFF | * | |
| 1238_AATGGCTGTCACTCTC | abT | TRBV23-1 | * | TRBJ1-5 | * | TRBC1 | * | CASSPHLQGPYQPQHF | * | |
| 1238_ACCAACAGTATTTCTC | abT | TRBV6-7 | * | TRBJ2-7 | * | * | * | CNIVGTVSYEQYF | * | |
| 1238_AGGAATAGTCACTTCC | abT | TRBV23-1 | * | TRBJ1-5 | * | * | * | CASSPHLQGPYQPQHF | * | |
| 1238_GATCACATCCACCTCA | abT | TRBV20-1 | * | TRBJ2-7 | * | * | * | CSAISPVSVGEQYF | * | |
| 1238_TCCATGCGTAGCTGTT | abT | TRBV2 | * | TRBJ2-5 | * | TRBC2 | * | CASSYGQGAYETQYF | * | |
| 1416_CAACCAAAGGACATCG | abT | TRBV20-1 | * | TRBJ2-3 | * | TRBC2 | * | CSARDPGGLAYTDTQYF | * | |
| 925_CTTGATTGTTCTCCAC | abT | TRBV6-2 | * | TRBJ2-5 | * | * | * | CASSLAGGSWTQYF | * | |
| 925_GCCGTGACAAGTGGTG | abT | TRBV20-1 | * | TRBJ1-1 | * | * | * | CSAWDSTEAFF | * | |
| 925_TATTTCGTCGTGTCAA | abT | TRBV20-1 | * | TRBJ2-7 | * | * | * | CSARVTAGGSSYEQYF | * | |
| 943_AGTAACCCACCGCTGA | abT | TRBV15 | * | TRBJ2-7 | * | * | * | CATSRDGTGDSYEQYF | * | |
| 943_CCTCCAAGTGACTATC | abT | TRBV7-9 | * | TRBJ1-2 | * | * | * | CASSRLLTHYGYTF | * | |
| 898_CATCCGTTCTCATTTG | abT | * | TRAV10 | * | TRAJ7 | * | TRAC | * | CVTVGPGNNRLAF | |
| 996_TACAACGAGCGCTTCG | abT | * | TRAV12-1 | * | TRAJ11 | * | * | * | CVVNGGSGYSTLTF | |
| 1238_GGTAATCCATAATGCC | abT | * | TRAV12-1 | * | TRAJ37 | * | * | * | CVVGSSNTGKLIF | |
| 925_GCCATGGCACAGTGAG | abT | * | TRAV12-1 | * | TRAJ39 | * | * | * | CVVNINNAGNMLTF | |
| 1224_TTTCAGTAGGGACCAT | abT | TRBV28 | TRAV12-1 | TRBJ2-3 | TRAJ9 | * | TRAC | CASSFPDTQYF | CVVPSDTGGFKTIF | |
| 1238_GAGACTTCAACACAGG | abT | * | TRAV12-2 | * | * | * | TRAC | * | CAVNQAGTALIF | |
| 1238_GTAGAGGAGTGAGCCA | abT | * | TRAV12-2 | * | * | * | * | * | CAVNQAGTALIF | |
| 831_TACAACGTCAGCTCTC | abT | * | TRAV12-2 | * | TRAJ24 | * | TRAC | * | CAVNSRTDSWGKLQF | |
| 1238_GCAGCCATCAGACATC | abT | * | TRAV12-2 | * | TRAJ3 | * | TRAC | * | CAVIPRGYSSASKIIF | |
| 934_GATGGAGGTAACGGTG | abT | * | TRAV12-3 | * | TRAJ17 | * | TRAC | * | CAMLKAAGNKLTF | |
| 898_TACGCTCCAATAGGGC | abT | * | TRAV12-3 | * | TRAJ8 | * | * | * | CAMSRTGFQKLVF | |
| 1130_AACCCAATCACTTTGT | abT | * | TRAV13-1 | * | TRAJ13 | * | * | * | CAATLSGGYQKVTF | |
| 925_GCCCGAAGTTGTCCCT | abT | * | TRAV13-1 | * | TRAJ31 | * | TRAC | * | CAASNGNNARLMF | |
| 1238_GTGAGCCTCGGTAACT | abT | * | TRAV13-1 | * | TRAJ48 | * | * | * | CAALYEKLTF | |
| 1224_GTCTGTCTCGTAGGAG | abT | * | TRAV13-2 | * | TRAJ13 | * | * | * | CAETGGYQKVTF | |
| 1195_AAGGTAACACACGGTC | abT | * | TRAV16 | * | TRAJ22 | * | * | * | CALTAGSARQLTF | |
| 1224_CCTCAGTAGGTCGACA | abT | * | TRAV17 | * | TRAJ30 | * | * | * | CATDIALNRDDKIIF | |
| 1066_CTGCATCCAATAGAGT | abT | * | TRAV17 | * | TRAJ42 | * | * | * | CAPYGGSQGNLIF | |
| 1224_GCAGGCTCATGAGATA | abT | * | TRAV19 | * | TRAJ17 | * | * | * | CALSEALGKAAGNKLTF | |
| 1235_GTACAACAGGAGAATG | abT | * | TRAV19 | * | TRAJ39 | * | * | * | CALSEVPNNAGNMLTF | |
| 1416_TTTATGCGTGCCCTTT | abT | * | TRAV19 | * | TRAJ50 | * | TRAC | * | CALKTSYDKVIF | |
| 898_CATCCACCAGACTCTA | abT | * | TRAV19 | * | TRAJ7 | * | * | * | CALSEPYGNNRLAF | |
| 801_GAAACCTTCGTGAGAG | abT | * | TRAV20 | * | TRAJ42 | * | * | * | CAVEGGSQGNLIF | |
| 1028_CTAGACAAGGTAGTAT | abT | * | TRAV22 | * | TRAJ57 | * | * | * | CAAEGGSEKLVF | |
| 1195_CTGATCCCAAGGTCTT | abT | * | TRAV23DV6 | * | TRAJ31 | * | * | * | CAASGNARLMF | |
| 1238_CAGGCCACAAAGACTA | abT | * | TRAV26-2 | * | TRAJ53 | * | TRAC | * | CILLSGSNYKLTF | |
| 996_TGAGTCAAGGTCGTAG | abT | * | TRAV29DV5 | * | TRAJ33 | * | * | * | CAADSNYQLIW | |
| 1355_ACCCACTAGTATGACA | abT | * | TRAV29DV5 | * | TRAJ47 | * | * | * | CAPKREYGNKLVF | |
| 925_AAACCCAGTGTTCCAA | abT | * | TRAV3 | * | TRAJ22 | * | * | * | CAVRVPSGSARQLTF | |
| 925_CCTAAGATCGAGATGG | abT | * | TRAV34 | * | TRAJ47 | * | * | * | CGASLEEYGNKLVF | |
| 801_AAACGCTTCGGCCCAA | abT | * | TRAV36DV7 | * | TRAJ54 | * | * | * | CAVEAF | |
| 996_GTGATGTAGGTGATAT | abT | * | TRAV38-2DV8 | * | TRAJ22 | * | TRAC | * | CAYRRTFSGSARQLTF | |
| 1224_CTAACTTGTCACTGAT | abT | * | TRAV38-2DV8 | * | TRAJ47 | * | * | * | CAYIWEYGNKLVF | |
| 1416_GTGTGATCACACGCCA | abT | * | TRAV38-2DV8 | * | TRAJ49 | * | * | * | CAYSGNQFYF | |
| 925_GGCTTGGGTTTCACTT | abT | * | TRAV38-2DV8 | * | TRAJ57 | * | * | * | CAYRSDPQGGSEKLVF | |
| 1130_ATTCTACCACCAGCTG | abT | * | TRAV39 | * | TRAJ30 | * | * | * | CAVGMNRDDKIIF | |
| 925_GTCATCCCAGCACCCA | abT | * | TRAV39 | * | TRAJ54 | * | * | * | CAVEKGGAQKLVF | |
| 943_CCCTTAGGTTCTGACA | abT | * | TRAV41 | * | TRAJ42 | * | * | * | CALNYGGSQGNLIF | |
| 966_AACAAAGAGCTTTCTT | abT | * | TRAV5 | * | TRAJ8 | * | * | * | CAEIPDTGFQKLVF | |
| 1238_CCCTCAATCGGCATAT | abT | * | TRAV9-2 | * | TRAJ37 | * | * | * | CALSLSSSNTGKLIF | |
| 1238_CTGTATTGTCCGGACT | abT | * | TRAV9-2 | * | TRAJ42 | * | * | * | CALSPFGGSQGNLIF | |
| 1195_CCTCACAAGGATATGT | gdT | TRDV2 | * | TRDJ3 | * | TRDC | * | CACDTLGDTSWDTRQMFF | * | |
| 1238_GGGCTACAGTAGTGCG | gdT | TRDV2 | * | TRDJ1 | * | TRDC | * | CACDNLGGSTDKLIF | * | |
| 1397_TGCGATACAGAACGCA | gdT | TRDV1 | * | TRDJ1 | * | TRDC | * | CALGDRQAFLRSSKPPYWGPHTDKLIF | * | |
| 1416_CGACAGCAGATGGGCT | gdT | TRDV2 | * | TRDJ1 | * | TRDC | * | CACDTVVKDTTDKLIF | * | |
| 1416_GTCTTTAGTTTCAGAC | gdT | TRDV2 | * | TRDJ1 | * | * | * | CACDRLPSGGYTDKLIF | * | |
| 943_GTTATGGCAGCCTATA | gdT | TRDV2 | * | TRDJ3 | * | TRDC | * | CDTVLGDLSSWDTRQMFF | * | |
| 1125_GGTAGAGGTTTCCCAC | gdT | TRDV2 | * | TRDJ3 | TRDC | * | * | CACDGLGDIPPRDSWDTRQMFF | * | |
| 1066_GGTAGAGCACGACTAT | gdT | * | TRGV10 | * | TRGJ2 | * | TRGC1 | * | CAAWVCWNYYKKLF | |
| 1167_AAGTTCGGTCCTTTGC | gdT | * | TRGV10 | * | TRGJ2 | * | * | * | CAASGNYKKLF | |
| 1195_TAGAGTCAGCGGTAAC | gdT | * | TRGV10 | * | TRGJ2 | * | TRGC2 | * | CAAFYYYKKLF | |
| 1224_AGTCATGCAATTGTGC | gdT | * | TRGV10 | * | TRGJ2 | * | * | * | CAAWDYWPKLF | |
| 1224_CATACAGGTGGACCAA | gdT | * | TRGV10 | * | TRGJ2 | * | TRGC2 | * | CAAWDYNYKKLF | |
| 1238_AATCACGTCCATAGGT | gdT | * | TRGV10 | * | TRGJ2 | * | * | * | CAAWDANYYKKLF | |
| 1238_GACGCTGTCCGCTTAC | gdT | * | TRGV10 | * | TRGJ2 | * | TRGC2 | * | CAAWEHYKKLF | |
| 1355_TGGCCAGTCTTCGGTC | gdT | * | TRGV10 | * | TRGJ2 | * | * | * | CAASGIEKLF | |
| 1130_CCTCCAATCCTACCGT | gdT | * | TRGV10 | * | TRGJP1 | * | * | * | CAAWEYYTRRDTTGWFKIF | |
| 1416_AATGCCAGTACAGCGA | gdT | TRDV2 | TRGV10 | TRDJ3 | TRGJP1 | TRDC | TRGC1 | CACDKMLGDNSWDTRQMFF | CATFRLRATTGWFKIF | |
| 1224_GTCTCACTCACACCCT | gdT | * | TRGV2 | * | TRGJ2 | * | * | * | CATWDGPEGDYKKLF | |
| 1224_GTGAGCCGTAGTAAGT | gdT | * | TRGV2 | * | TRGJ2 | * | * | * | CATWDDFSYYKKLF | |
| 1238_ACGATGTTCACGAACT | gdT | * | TRGV2 | * | TRGJ2 | * | TRGC2 | * | CATWDGLYYKKLF | |
| 1238_TGTTACTGTTCCTAAG | gdT | * | TRGV2 | * | TRGJ2 | * | TRGC1 | * | CATWEGGYKKLF | |
| 1238_TGTTACTTCTGAACGT | gdT | * | TRGV2 | * | TRGJ2 | * | TRGC2 | * | CATWDGQEKLF | |
| 925_AGGCCACGTACTGACT | gdT | * | TRGV2 | * | TRGJ2 | * | * | * | CATWDVPPWKKLF | |
| 925_TAGGTACAGACATATG | gdT | * | TRGV2 | * | TRGJ2 | * | * | * | CATWDIYYKKLF | |
| 925_TGCTTCGAGATCGCCC | gdT | * | TRGV2 | * | TRGJ2 | * | * | * | CATWDELGKKLF | |
| 945_GAGCTGCGTTATGGTC | gdT | * | TRGV2 | * | TRGJ2 | * | TRGC2 | * | CATWDGPTQESYYMKLF | |
| 1167_ATTTACCCAAATGATG | gdT | * | TRGV3 | * | TRGJ2 | * | TRGC2 | * | CATWDSLYKKLF | |
| 1235_ATGAAAGCAGAGGTAC | gdT | * | TRGV3 | * | TRGJ2 | * | * | * | CATLHYYKKLF | |
| 1238_AGAGCCCTCACAATGC | gdT | * | TRGV3 | * | TRGJ2 | * | * | * | CHLGQGPLIFYYKKLF | |
| 966-2_CCGTGAGAGAGGGTAA | gdT | * | TRGV3 | * | TRGJ2 | * | TRGC1 | * | CATWDRPRYYKKLF | |
| 1235_TCAAGCAGTAGACTGG | gdT | * | TRGV3 | * | TRGJP2 | * | * | * | CATWDRLPDWIKTF | |
| 1125_CTAAGTGTCTACGCAA | gdT | * | TRGV4 | * | TRGJ2 | * | * | * | CATPNNKLF | |
| 1224_ACGTCCTTCATTATCC | gdT | * | TRGV4 | * | TRGJ2 | * | TRGC2 | * | CATWDPSRNYYKKLF | |
| 925_GTCCTCATCCATGATG | gdT | * | TRGV4 | * | TRGJ2 | * | * | * | CATRYYYKKLF | |
| 934_GTGACGCCAATGAACA | gdT | * | TRGV4 | * | TRGJ2 | * | * | * | CAPPVKKLF | |
| 966_TCAGTTTTCGCCAGAC | gdT | * | TRGV5 | * | TRGJ2 | * | * | * | CATWDRRYYKKLF | |
| 1224_AGCATCAGTGGCTTAT | gdT | * | TRGV7 | * | TRGJ2 | * | * | * | CATWDKAKLF | |
| 1224_CATGGTAAGCTGAAAT | gdT | * | TRGV8 | * | TRGJ2 | * | TRGC2 | * | CATWDRWYKKLF | |
| 1235_TCAGTGATCCTTATAC | gdT | * | TRGV8 | * | TRGJ2 | * | * | * | CATWAMHPNFYYKKLF | |
| 1238_CGGGCATAGCACCGTC | gdT | * | TRGV8 | * | TRGJ2 | * | * | * | CATWDMRGYKKLF | |
| 1066_GCCGTGACAATCGTCA | gdT | * | TRGV9 | * | TRGJ2 | * | TRGC2 | * | CALWGRWDKKLF | |
| 1195_ACATTTCAGGGAGGAC | gdT | * | TRGV9 | * | TRGJ2 | * | * | * | CALWEHQNYYKKLF | |
| 1235_TATACCTTCTCACCCA | gdT | * | TRGV9 | * | TRGJ2 | * | * | * | CALWTMNYYKKLF | |
| 1238_AGTTAGCTCTTAGCTT | gdT | * | TRGV9 | * | TRGJ2 | * | * | * | CALWEDYYKKLF | |
| 801_GCTGGGTCAAAGACTA | gdT | * | TRGV9 | * | TRGJ2 | * | * | * | CALWEFSNYYKKLF | |
| 945_GAAACCTTCACACCCT | gdT | TRDV1 | TRGV9 | TRDJ1 | TRGJ2 | TRDC | TRGC2 | CALGGGSVGGYGDKLIF | CAFQPNTSYYKKLF | |
| 996_CTCATGCGTGGACAGT | gdT | * | TRGV9 | * | TRGJ2 | * | TRGC2 | * | CALSISASYYKKLF | |
| 898_GCAACATCAAACCACT | gdT | TRDV2 | TRGV9 | TRDJ3 | TRGJP | TRDC | TRGC1 | CACDTLLGDTWDTRQMFF | CALWEVQELGKKIKVF | blood clonotype |
| 996_TATTCCAAGGACGCTA | gdT | * | TRGV9 | * | TRGJP | * | TRGC1 | * | CALWGNKSWAKKIKVF | |

| Normalized gene expression | | | | | | | | | | | | | | | | | | | | | | | | | | | | | | | |
|---|---|---|---|---|---|---|---|---|---|---|---|---|---|---|---|---|---|---|---|---|---|---|---|---|---|---|---|---|---|---|---|
| **Barcode** | **cell_type** | **XCL1** | **SELL** | **CCL20** | **PDCD1** | **CXCR6** | **CCR2** | **CCR5** | **TIGIT** | **LEF1** | **IL7R** | **ITGA1** | **GZMK** | **GZMA** | **TCF7** | **HAVCR2** | **CXCR3** | **TOX** | **PRF1** | **ENTPD1** | **MKI67** | **LAG3** | **CD69** | **KLRC1** | **IFNG** | **GZMB** | **ITGAE** | **CCL5** | **CCL3** | **CCR7** | **TNFSF9** |
| 1066_GTGAGGACACAAGGTG | abT | 0 | 0 | 0,546758 | 0,898243 | 1,157802 | 0,546758 | 0 | 2,197677 | 0 | 0 | 1,363684 | 0 | 5,989014 | 0 | 1,157802 | 1,363684 | 0,546758 | 2,113412 | 1,920027 | 0 | 3,677947 | 1,680056 | 0 | 1,807223 | 0,546758 | 0,546758 | 3,404715 | 4,376832 | 0 | 0,546758 |
| 1195_AGAAATGGTGGTGATG | abT | 0 | 0 | 1,312228 | 0,857483 | 0 | 0 | 0 | 1,860801 | 0 | 1,860801 | 0 | 0 | 0 | 0 | 0 | 0 | 0 | 0 | 0,857483 | 0 | 0 | 0 | 0 | 0 | 0 | 0,857483 | 0 | 0 | 0 | 0 |
| 1224_CGAAGTTGTAGCACAG | abT | 0 | 0 | 0 | 1,970125 | 1,632056 | 0 | 1,117496 | 1,117496 | 0 | 0 | 0 | 3,162478 | 3,834238 | 0 | 0 | 1,117496 | 0 | 0 | 0 | 0 | 0 | 1,632056 | 0 | 0 | 1,632056 | 0 | 3,959474 | 0 | 0 | 0 |
| 1224_GGGATCCCACAGAGCA | abT | 0 | 1,142606 | 0 | 0 | 1,142606 | 0 | 0 | 0 | 0 | 0 | 0 | 2,002126 | 3,006392 | 0 | 0 | 0 | 0 | 1,142606 | 0 | 0 | 0 | 2,255463 | 0 | 0 | 0 | 0 | 1,662003 | 0 | 0 | 0 |
| 1238_AATGGCTGTCACTCTC | abT | 0 | 0 | 0 | 1,400946 | 0 | 0 | 0 | 0 | 0 | 1,400946 | 0 | 2,962911 | 2,582952 | 1,400946 | 0 | 0 | 1,400946 | 0 | 0 | 0 | 0 | 1,400946 | 0 | 0 | 0 | 0 | 3,237592 | 0 | 1,400946 | 0 |
| 1238_ACCAACAGTATTTCTC | abT | 0 | 1,366017 | 0 | 1,366017 | 0 | 0 | 0 | 0 | 0 | 4,033788 | 1,366017 | 0 | 1,922702 | 0 | 0 | 0 | 0 | 0 | 0 | 0 | 0 | 2,278203 | 0 | 0 | 0 | 0 | 1,366017 | 0 | 0 | 0 |
| 1238_AGGAATAGTCACTTCC | abT | 0 | 0 | 0 | 0 | 0 | 1,48534 | 0 | 0 | 0 | 0 | 0 | 3,560042 | 2,420315 | 0 | 0 | 0 | 0 | 1,48534 | 0 | 0 | 0 | 1,48534 | 0 | 2,058337 | 0 | 0 | 2,894936 | 0 | 1,48534 | 1,48534 |
| 1238_GATCACATCCACCTCA | abT | 1,093307 | 0 | 0 | 0 | 1,093307 | 0 | 0 | 1,093307 | 0 | 0 | 1,603068 | 2,700615 | 3,603166 | 0 | 0 | 1,093307 | 1,093307 | 1,603068 | 0 | 1,093307 | 1,603068 | 1,093307 | 0 | 0 | 0 | 1,093307 | 4,664936 | 0 | 0 | 0 |
| 1238_TCCATGCGTAGCTGTT | abT | 0 | 0 | 0 | 1,655461 | 0 | 0 | 0 | 0 | 0 | 0 | 0 | 2,887143 | 3,273855 | 0 | 0 | 2,248233 | 0 | 0 | 0 | 0 | 0 | 0 | 0 | 0 | 0 | 0 | 3,862632 | 0 | 0 | 0 |
| 1416_CAACCAAAGGACATCG | abT | 0 | 0 | 0 | 0 | 2,937568 | 1,024962 | 0 | 0 | 0 | 0 | 0 | 0 | 0 | 1,024962 | 0 | 0 | 0 | 1,024962 | 0 | 0 | 0 | 1,850181 | 0 | 0 | 0 | 0 | 0 | 0 | 0 | 0 |
| 925_CTTGATTGTTCTCCAC | abT | 0 | 0 | 0 | 0 | 0 | 0 | 1,874918 | 1,324457 | 1,324457 | 0 | 0 | 2,865673 | 2,488282 | 1,324457 | 0 | 1,874918 | 1,324457 | 0 | 0 | 0 | 0 | 2,694676 | 0 | 0 | 0 | 1,324457 | 3,529934 | 1,324457 | 0 | 0 |
| 925_GCCGTGACAAGTGGTG | abT | 0 | 0 | 0 | 0 | 0 | 0 | 0 | 0 | 1,134671 | 1,652555 | 0 | 0 | 0 | 0 | 0 | 0 | 0 | 0 | 0 | 0 | 0 | 0 | 0 | 0 | 0 | 0 | 0 | 0 | 0 | 0 |
| 925_TATTTCGTCGTGTCAA | abT | 0 | 0 | 0 | 0 | 1,381886 | 0 | 0 | 0 | 0 | 3,085428 | 0 | 1,94087 | 0 | 0 | 0 | 1,381886 | 0 | 0 | 0 | 0 | 0 | 1,381886 | 0 | 0 | 0 | 1,381886 | 3,755455 | 0 | 0 | 0 |
| 943_AGTAACCCACCGCTGA | abT | 0 | 0 | 0 | 0 | 1,311435 | 0 | 0 | 0 | 0 | 0 | 0 | 2,471989 | 1,311435 | 0 | 0 | 1,311435 | 0 | 0 | 0 | 0 | 0 | 1,859884 | 0 | 1,311435 | 0 | 0 | 0 | 0 | 0 | 0 |
| 943_CCTCCAAGTGACTATC | abT | 0 | 0 | 0 | 0 | 0 | 0 | 0 | 0 | 0 | 0 | 0 | 1,49627 | 1,49627 | 0 | 0 | 0 | 0 | 1,49627 | 0 | 0 | 0 | 3,081452 | 0 | 0 | 0 | 0 | 2,070654 | 0 | 0 | 0 |
| 898_CATCCGTTCTCATTTG | abT | 0 | 0 | 0 | 0 | 0 | 0 | 0 | 0 | 0 | 0 | 0 | 2,9218 | 3,737851 | 0 | 1,925449 | 1,925449 | 1,368414 | 2,28109 | 0 | 0 | 0 | 2,750188 | 2,28109 | 0 | 2,28109 | 0 | 4,4537 | 0 | 0 | 0 |
| 996_TACAACGAGCGCTTCG | abT | 0 | 1,484284 | 0 | 0 | 0 | 0 | 0 | 0 | 2,419071 | 3,342665 | 0 | 0 | 0 | 0 | 0 | 0 | 0 | 0 | 0 | 0 | 0 | 1,484284 | 0 | 0 | 0 | 0 | 0 | 0 | 2,419071 | 0 |
| 1238_GGTAATCCATAATGCC | abT | 0 | 0 | 0 | 0 | 0 | 0 | 0 | 0 | 0 | 0 | 0 | 3,082246 | 3,752198 | 0 | 1,64122 | 0 | 0 | 1,64122 | 0 | 0 | 0 | 0 | 0 | 1,64122 | 0 | 1,64122 | 4,081943 | 0 | 0 | 0 |
| 925_GCCATGGCACAGTGAG | abT | 0 | 0 | 0 | 0 | 0 | 0 | 0 | 0 | 0 | 4,221184 | 0 | 2,500262 | 0 | 0 | 0 | 0 | 0 | 0 | 0 | 0 | 2,500262 | 2,500262 | 0 | 0 | 0 | 0 | 0 | 0 | 0 | 0 |
| 1224_TTTCAGTAGGGACCAT | abT | 0 | 0 | 0 | 1,272575 | 0 | 0 | 0 | 0 | 0 | 0 | 0 | 0 | 2,943925 | 0 | 0 | 1,272575 | 0 | 1,814836 | 0 | 0 | 1,272575 | 0 | 0 | 0 | 2,798512 | 1,272575 | 3,958919 | 1,272575 | 0 | 0 |
| 1238_GAGACTTCAACACAGG | abT | 0 | 0 | 0 | 1,563098 | 2,989107 | 0 | 0 | 0 | 0 | 0 | 2,1456 | 3,311094 | 2,1456 | 0 | 0 | 1,563098 | 1,563098 | 0 | 0 | 0 | 0 | 0 | 0 | 1,563098 | 0 | 0 | 3,913154 | 1,563098 | 0 | 0 |
| 1238_GTAGAGGAGTGAGCCA | abT | 0 | 0 | 0 | 0 | 0 | 0 | 0 | 0 | 0 | 0 | 1,836234 | 3,095192 | 3,485454 | 0 | 0 | 0 | 0 | 0 | 0 | 0 | 1,291004 | 1,291004 | 0 | 0 | 0 | 1,291004 | 4,535555 | 1,291004 | 1,291004 | 0 |
| 831_TACAACGTCAGCTCTC | abT | 0 | 0 | 0 | 0 | 1,124465 | 1,124465 | 0 | 0 | 1,124465 | 0 | 2,74406 | 2,231549 | 1,979028 | 1,124465 | 0 | 1,124465 | 0 | 1,640382 | 0 | 0 | 1,640382 | 0 | 0 | 0 | 1,640382 | 1,124465 | 3,5928 | 0 | 0 | 0 |
| 1238_GCAGCCATCAGACATC | abT | 0 | 1,318611 | 0 | 0 | 2,480974 | 0 | 1,318611 | 0 | 0 | 0 | 0 | 2,858156 | 3,244311 | 0 | 0 | 0 | 1,318611 | 0 | 0 | 0 | 1,318611 | 1,318611 | 0 | 0 | 0 | 1,318611 | 3,739415 | 0 | 1,868173 | 0 |
| 934_GATGGAGGTAACGGTG | abT | 0 | 0 | 0 | 0 | 0 | 0 | 0 | 0 | 0 | 3,636502 | 0 | 2,188267 | 2,823727 | 1,837343 | 1,29196 | 1,29196 | 0 | 0 | 0 | 0 | 0 | 1,29196 | 0 | 0 | 0 | 0 | 2,653212 | 0 | 0 | 0 |
| 898_TACGCTCCAATAGGGC | abT | 0 | 0 | 0 | 0 | 0 | 0 | 0 | 0 | 0 | 3,213516 | 0 | 0 | 1,841126 | 0 | 1,295225 | 0 | 0 | 0 | 0 | 0 | 0 | 0 | 0 | 0 | 0 | 0 | 1,295225 | 0 | 0 | 0 |
| 1130_AACCCAATCACTTTGT | abT | 0 | 0 | 0 | 0 | 0 | 0 | 1,528817 | 0 | 0 | 0 | 2,107231 | 2,107231 | 2,737665 | 0 | 0 | 1,528817 | 0 | 0 | 0 | 0 | 0 | 0 | 0 | 0 | 0 | 0 | 4,388006 | 0 | 0 | 0 |
| 925_GCCCGAAGTTGTCCCT | abT | 0 | 0 | 0 | 0 | 0 | 0 | 0 | 0 | 0 | 3,954978 | 0 | 0 | 0 | 2,254699 | 0 | 0 | 0 | 0 | 0 | 0 | 0 | 2,254699 | 0 | 0 | 0 | 0 | 0 | 0 | 0 | 0 |
| 1238_GTGAGCCTCGGTAACT | abT | 0 | 0 | 0 | 0 | 0 | 0 | 0 | 0 | 0 | 0 | 2,236201 | 3,849454 | 3,538953 | 0 | 0 | 0 | 0 | 0 | 0 | 0 | 0 | 0 | 0 | 0 | 0 | 0 | 4,333616 | 0 | 0 | 0 |
| 1224_GTCTGTCTCGTAGGAG | abT | 0 | 1,400025 | 0 | 0,92729 | 0 | 0 | 0 | 0 | 1,719718 | 1,961587 | 0 | 0 | 0 | 1,400025 | 0 | 0 | 1,400025 | 0 | 0 | 0 | 0 | 1,961587 | 0 | 0 | 0 | 0 | 0 | 0 | 0,92729 | 0 |
| 1195_AAGGTAACACACGGTC | abT | 0 | 0 | 0 | 0 | 0 | 0 | 0 | 0 | 0 | 0 | 0 | 0 | 1,994592 | 0 | 0 | 0 | 0 | 0 | 0 | 0 | 0 | 0 | 0 | 0 | 0 | 1,429019 | 1,429019 | 0 | 0 | 0 |
| 1224_CCTCAGTAGGTCGACA | abT | 0 | 0 | 0 | 0 | 0 | 0 | 0 | 0 | 0 | 0 | 2,938584 | 0 | 0 | 2,297024 | 0 | 0 | 0 | 0 | 0 | 0 | 0 | 0 | 0 | 0 | 0 | 0 | 2,297024 | 0 | 0 | 0 |
| 1066_CTGCATCCAATAGAGT | abT | 0 | 1,443925 | 0 | 0 | 0 | 0 | 0 | 0 | 0 | 2,63542 | 0 | 1,443925 | 2,512078 | 1,767457 | 0 | 0,962658 | 0 | 0 | 0 | 0 | 0 | 0,962658 | 0 | 0 | 0 | 0 | 0 | 0 | 0 | 0 |
| 1224_GCAGGCTCATGAGATA | abT | 0 | 0 | 0 | 0 | 1,827496 | 0 | 0 | 0 | 1,007751 | 2,812704 | 0 | 2,271834 | 2,912042 | 0 | 0 | 1,827496 | 0 | 0 | 0 | 0 | 0 | 2,578399 | 0 | 0 | 0 | 0 | 3,085262 | 0 | 0 | 0 |
| 1235_GTACAACAGGAGAATG | abT | 0 | 0 | 0 | 0 | 0 | 0 | 0 | 0 | 0 | 0 | 0 | 3,459691 | 0 | 0 | 0 | 0 | 0 | 1,96871 | 0 | 0 | 0 | 0 | 0 | 0 | 0 | 0 | 4,31665 | 0 | 0 | 0 |
| 1416_TTTATGCGTGCCCTTT | abT | 0 | 0 | 0 | 0 | 2,206283 | 0 | 1,306701 | 0 | 0 | 0 | 1,85441 | 1,306701 | 3,330047 | 0 | 0 | 0 | 0 | 0 | 0 | 0 | 0 | 1,306701 | 0 | 0 | 0 | 0 | 2,466052 | 0 | 0 | 0 |
| 898_CATCCACCAGACTCTA | abT | 0 | 0,970123 | 0 | 0 | 0 | 1,453144 | 0,970123 | 0 | 0 | 2,021955 | 0 | 1,453144 | 3,028163 | 0,970123 | 0 | 0 | 0,970123 | 0 | 0 | 0 | 0 | 0 | 0 | 0 | 0 | 0 | 3,028163 | 0 | 0 | 0,970123 |
| 801_GAAACCTTCGTGAGAG | abT | 0 | 0,660161 | 1,556125 | 0 | 0 | 0,660161 | 0 | 0 | 0 | 2,709819 | 0 | 0 | 2,020981 | 0 | 0,660161 | 0 | 0 | 0 | 0 | 0 | 0 | 2,503176 | 0 | 0 | 0 | 0,660161 | 0 | 0 | 0,660161 | 0 |
| 1028_CTAGACAAGGTAGTAT | abT | 0 | 0 | 0 | 0 | 0 | 0 | 0 | 0 | 0 | 3,848309 | 0 | 0 | 3,848309 | 0 | 0 | 0 | 0 | 0 | 0 | 0 | 0 | 2,320694 | 0 | 0 | 0 | 0 | 2,963492 | 0 | 0 | 0 |
| 1195_CTGATCCCAAGGTCTT | abT | 0 | 0 | 0 | 0 | 1,774471 | 0 | 0 | 0 | 0 | 2,583462 | 0 | 0 | 2,753118 | 1,237962 | 0 | 0 | 0 | 1,237962 | 0 | 0 | 0 | 1,774471 | 0 | 0 | 0 | 0 | 2,583462 | 0 | 0 | 0 |
| 1238_CAGGCCACAAAGACTA | abT | 0 | 0 | 0 | 0 | 0 | 0 | 0 | 1,22355 | 0 | 1,757593 | 0 | 0 | 0 | 0 | 0 | 0 | 0 | 0 | 0 | 0 | 0 | 0 | 0 | 0 | 0 | 0 | 0 | 0 | 0 | 0 |
| 996_TGAGTCAAGGTCGTAG | abT | 0 | 0 | 0 | 0,99158 | 0 | 0 | 0 | 1,806068 | 0 | 0 | 0 | 2,413495 | 0,99158 | 0 | 1,479552 | 0 | 0 | 0,99158 | 0 | 0 | 0 | 1,479552 | 0 | 0 | 0,99158 | 0 | 3,33676 | 0 | 0 | 0 |
| 1355_ACCCACTAGTATGACA | abT | 0 | 0 | 0 | 0 | 0 | 0 | 0 | 0 | 0 | 0 | 0 | 0 | 0 | 0 | 0 | 0 | 0 | 0 | 0 | 0 | 0 | 0 | 0 | 0 | 0 | 0 | 4,03386 | 0 | 0 | 0 |
| 925_AAACCCAGTGTTCCAA | abT | 0 | 0 | 0 | 0 | 1,415411 | 0 | 0 | 0 | 0 | 2,600667 | 1,415411 | 2,981079 | 1,979118 | 0 | 1,979118 | 0 | 0 | 0 | 0 | 0 | 0 | 0 | 0 | 0 | 0 | 0 | 3,563882 | 0 | 0 | 0 |
| 925_CCTAAGATCGAGATGG | abT | 0 | 0 | 0 | 0 | 0 | 0 | 0 | 0 | 0 | 0 | 0 | 3,643784 | 3,399914 | 0 | 0 | 0 | 0 | 2,227293 | 0 | 0 | 1,636539 | 2,227293 | 0 | 0 | 0 | 1,636539 | 4,076232 | 0 | 0 | 0 |
| 801_AAACGCTTCGGCCCAA | abT | 0 | 0 | 0 | 1,347603 | 0 | 0 | 0 | 0 | 0 | 0 | 1,347603 | 2,517113 | 3,475999 | 0 | 0 | 0 | 1,347603 | 0 | 0 | 0 | 0 | 0 | 0 | 1,901567 | 0 | 1,347603 | 4,355441 | 0 | 1,347603 | 0 |
| 996_GTGATGTAGGTGATAT | abT | 0 | 1,053167 | 0 | 0 | 0 | 0 | 0 | 1,053167 | 1,053167 | 1,554651 | 0 | 0 | 0 | 1,053167 | 0 | 0 | 0 | 1,053167 | 1,053167 | 0 | 0 | 0 | 0 | 0 | 0 | 1,053167 | 0 | 0 | 0 | 0 |
| 1224_CTAACTTGTCACTGAT | abT | 0 | 2,051513 | 0 | 0 | 0 | 0 | 0 | 0 | 0 | 2,788354 | 0 | 0 | 0 | 1,805785 | 0 | 0 | 0 | 0 | 0 | 0 | 0 | 1,805785 | 0 | 0 | 0 | 0,991367 | 0 | 0 | 2,554464 | 0 |
| 1416_GTGTGATCACACGCCA | abT | 0 | 0 | 0 | 0 | 1,546603 | 1,046528 | 1,046528 | 0 | 0 | 0 | 0 | 1,546603 | 3,219703 | 0 | 0 | 1,046528 | 0 | 0 | 0 | 0 | 0 | 1,046528 | 0 | 0 | 0 | 1,046528 | 3,142986 | 0 | 0 | 0 |
| 925_GGCTTGGGTTTCACTT | abT | 0 | 0 | 0 | 0 | 0 | 0 | 0 | 0 | 0 | 0 | 0 | 0 | 0 | 0 | 0 | 0 | 0 | 0 | 0 | 0 | 0 | 3,105571 | 0 | 3,105571 | 0 | 0 | 0 | 0 | 0 | 0 |
| 1130_ATTCTACCACCAGCTG | abT | 1,588336 | 0 | 0 | 1,588336 | 0 | 0 | 0 | 2,173751 | 0 | 0 | 0 | 3,341772 | 1,588336 | 0 | 0 | 0 | 0 | 1,588336 | 0 | 0 | 0 | 2,540562 | 0 | 0 | 0 | 0 | 3,585195 | 2,173751 | 0 | 1,588336 |
| 925_GTCATCCCAGCACCCA | abT | 0 | 0 | 0 | 1,173145 | 0 | 0 | 0 | 0 | 0 | 2,498211 | 0 | 0 | 2,040756 | 1,698233 | 1,173145 | 2,040756 | 0 | 0 | 0 | 0 | 0 | 2,810908 | 0 | 0 | 0 | 1,173145 | 2,498211 | 0 | 0 | 0 |
| 943_CCCTTAGGTTCTGACA | abT | 0 | 0 | 0 | 0 | 0 | 0 | 0 | 0 | 0 | 2,433413 | 0 | 0 | 0 | 1,280775 | 0 | 0 | 0 | 0 | 0 | 0 | 0 | 2,174557 | 0 | 0 | 0 | 0 | 0 | 0 | 0 | 0 |
| 966_AACAAAGAGCTTTCTT | abT | 0 | 0 | 0 | 0 | 0 | 0 | 0 | 0 | 0 | 0 | 0 | 3,01824 | 3,904759 | 0 | 0 | 0 | 0 | 1,445163 | 0 | 0 | 0 | 0 | 0 | 0 | 1,445163 | 2,012912 | 4,235459 | 0 | 0 | 0 |
| 1238_CCCTCAATCGGCATAT | abT | 0 | 0 | 0 | 0 | 1,985324 | 0 | 1,985324 | 0 | 0 | 0 | 0 | 1,985324 | 0 | 0 | 0 | 0 | 0 | 0 | 1,985324 | 0 | 0 | 0 | 0 | 0 | 0 | 0 | 1,985324 | 0 | 1,985324 | 0 |
| 1238_CTGTATTGTCCGGACT | abT | 0 | 0 | 0 | 0 | 0 | 1,455408 | 0 | 0 | 0 | 0 | 0 | 3,777828 | 1,455408 | 0 | 0 | 1,455408 | 1,455408 | 0 | 0 | 0 | 0 | 0 | 0 | 0 | 0 | 0 | 3,777828 | 0 | 0 | 0 |
| 1195_CCTCACAAGGATATGT | gdT | 0 | 0 | 0 | 0 | 0,967587 | 0 | 0 | 0 | 0 | 3,843411 | 0 | 2,214621 | 2,214621 | 1,774064 | 0 | 0 | 0 | 1,774064 | 0 | 0 | 0 | 2,378576 | 0 | 0 | 0 | 0 | 3,171458 | 0 | 0 | 0 |
| 1238_GGGCTACAGTAGTGCG | gdT | 0 | 0 | 0 | 0 | 1,431438 | 0 | 1,431438 | 0 | 0 | 2,356518 | 0 | 3,148153 | 1,431438 | 0 | 0 | 1,431438 | 0 | 0 | 0 | 0 | 0 | 2,356518 | 1,431438 | 0 | 0 | 2,356518 | 3,148153 | 0 | 0 | 0 |
| 1397_TGCGATACAGAACGCA | gdT | 0 | 0 | 0 | 0,831026 | 0 | 0 | 0 | 1,278527 | 0 | 0 | 0 | 2,430572 | 3,339645 | 0 | 0 | 0,831026 | 0,831026 | 1,278527 | 0 | 0 | 0,831026 | 0 | 0 | 0 | 0,831026 | 0,831026 | 3,685628 | 0 | 0 | 0 |
| 1416_CGACAGCAGATGGGCT | gdT | 1,250624 | 0 | 0 | 0 | 2,914937 | 0 | 1,250624 | 0 | 0 | 1,789264 | 0 | 0 | 3,255216 | 0 | 0 | 1,789264 | 1,250624 | 0 | 0 | 0 | 0 | 0 | 2,137424 | 0 | 1,250624 | 0 | 3,431088 | 0 | 0 | 0 |
| 1416_GTCTTTAGTTTCAGAC | gdT | 0 | 0 | 0 | 0 | 1,997523 | 0 | 1,138984 | 0 | 0 | 1,138984 | 0 | 0 | 2,764125 | 0 | 0 | 0 | 0 | 1,657692 | 0 | 0 | 0 | 1,138984 | 0 | 0 | 0 | 1,138984 | 2,250699 | 0 | 0 | 0 |
| 943_GTTATGGCAGCCTATA | gdT | 0 | 0 | 0 | 0 | 0 | 0 | 0 | 0 | 0 | 1,920131 | 0 | 0 | 2,370532 | 0 | 0 | 0 | 0 | 1,078619 | 0 | 0 | 1,078619 | 0 | 1,585398 | 0 | 1,078619 | 0 | 3,777201 | 0 | 0 | 0 |
| 1125_GGTAGAGGTTTCCCAC | gdT | 0 | 0 | 0 | 0 | 0 | 0 | 0 | 0 | 0 | 4,119866 | 1,953751 | 3,227654 | 1,393158 | 0 | 0 | 1,393158 | 0 | 2,310816 | 0 | 0 | 0 | 2,310816 | 0 | 0 | 1,393158 | 1,393158 | 2,781162 | 0 | 0 | 0 |
| 1066_GGTAGAGCACGACTAT | gdT | 0 | 0 | 0 | 0 | 0 | 0 | 0 | 0 | 0 | 2,323805 | 1,195018 | 1,724057 | 2,966763 | 0 | 0 | 0 | 0 | 2,068235 | 0 | 0 | 1,195018 | 2,840558 | 1,195018 | 0 | 0 | 0 | 2,323805 | 0 | 0 | 0 |
| 1167_AAGTTCGGTCCTTTGC | gdT | 0 | 0 | 0 | 0 | 0 | 0 | 1,844929 | 0 | 0 | 0 | 0 | 2,455761 | 3,105062 | 0 | 0 | 0 | 1,844929 | 0 | 0 | 0 | 0 | 0 | 0 | 0 | 0 | 0 | 1,844929 | 0 | 0 | 0 |
| 1195_TAGAGTCAGCGGTAAC | gdT | 0 | 0 | 0 | 0 | 1,428537 | 0 | 0 | 0 | 0 | 0 | 1,428537 | 2,825119 | 2,616692 | 0 | 1,428537 | 1,994044 | 0 | 1,994044 | 0 | 0 | 1,428537 | 1,994044 | 0 | 0 | 2,616692 | 1,428537 | 3,580695 | 0 | 0 | 0 |
| 1224_AGTCATGCAATTGTGC | gdT | 0 | 0 | 0 | 0 | 0 | 0 | 0 | 0 | 0 | 0 | 1,341104 | 3,769283 | 2,88701 | 0 | 0 | 0 | 1,341104 | 0 | 0 | 0 | 1,341104 | 3,375286 | 0 | 0 | 0 | 1,341104 | 4,099142 | 0 | 0 | 0 |
| 1224_CATACAGGTGGACCAA | gdT | 0 | 0 | 0 | 1,199373 | 0 | 0 | 0 | 0 | 0 | 2,846434 | 0 | 2,701912 | 3,438315 | 1,199373 | 0 | 0 | 0 | 1,199373 | 0 | 0 | 0 | 0 | 0 | 0 | 1,729187 | 0 | 3,951181 | 0 | 0 | 0 |
| 1238_AATCACGTCCATAGGT | gdT | 0 | 0 | 0 | 1,823276 | 1,823276 | 0 | 1,823276 | 0 | 0 | 0 | 2,432228 | 2,432228 | 2,432228 | 0 | 0 | 0 | 0 | 1,823276 | 0 | 0 | 3,080461 | 1,823276 | 0 | 0 | 0 | 0 | 4,491619 | 0 | 0 | 0 |
| 1238_GACGCTGTCCGCTTAC | gdT | 0 | 0 | 0 | 0 | 0 | 0 | 0 | 0 | 0 | 0 | 0 | 3,481501 | 2,818651 | 0 | 0 | 0 | 1,597073 | 0 | 0 | 0 | 0 | 2,550669 | 0 | 0 | 0 | 0 | 3,791529 | 1,597073 | 0 | 0 |
| 1355_TGGCCAGTCTTCGGTC | gdT | 0 | 0 | 0 | 0 | 2,085326 | 0 | 0 | 0 | 0 | 0 | 2,085326 | 2,714326 | 3,373787 | 0 | 0 | 0 | 0 | 2,085326 | 0 | 0 | 0 | 2,714326 | 2,085326 | 0 | 0 | 0 | 3,918612 | 0 | 0 | 0 |
| 1130_CCTCCAATCCTACCGT | gdT | 1,336329 | 1,336329 | 0 | 0 | 0 | 0 | 0 | 0 | 1,336329 | 3,885388 | 0 | 1,888599 | 2,50309 | 0 | 1,336329 | 1,336329 | 0 | 1,888599 | 0 | 0 | 0 | 1,336329 | 0 | 0 | 2,50309 | 0 | 3,154461 | 0 | 0 | 0 |
| 1416_AATGCCAGTACAGCGA | gdT | 1,18879 | 0 | 0 | 0 | 2,31574 | 0 | 0 | 0 | 0 | 0 | 0 | 0 | 3,562368 | 0 | 0 | 0 | 0 | 0 | 0 | 0 | 1,18879 | 0 | 2,060426 | 0 | 0 | 0 | 2,958282 | 0 | 0 | 0 |
| 1224_GTCTCACTCACACCCT | gdT | 0 | 0,941269 | 0 | 0 | 0 | 0 | 0 | 0 | 0 | 0 | 0 | 0 | 2,480114 | 0 | 0 | 0,941269 | 0 | 0,941269 | 0 | 0 | 0 | 0,941269 | 0 | 0 | 0 | 0 | 4,447517 | 0 | 0 | 0 |
| 1224_GTGAGCCGTAGTAAGT | gdT | 0 | 0 | 0 | 0 | 0 | 0 | 0 | 0 | 0 | 0 | 0 | 0 | 1,381886 | 0 | 0 | 0 | 0 | 1,381886 | 0 | 0 | 0 | 0 | 0 | 0 | 2,55952 | 0 | 3,428295 | 0 | 0 | 0 |
| 1238_ACGATGTTCACGAACT | gdT | 0 | 0 | 0 | 0 | 1,65529 | 0 | 0 | 0 | 0 | 1,136967 | 1,65529 | 2,248044 | 2,248044 | 1,136967 | 0 | 1,136967 | 0 | 0 | 0 | 0 | 0 | 0 | 0 | 0 | 0 | 0 | 3,551821 | 1,65529 | 0 | 0 |
| 1238_TGTTACTGTTCCTAAG | gdT | 0 | 0 | 0 | 0 | 0 | 0 | 0 | 0 | 0 | 0 | 0 | 1,606564 | 3,493023 | 0 | 0 | 0 | 0 | 1,606564 | 0 | 0 | 0 | 1,606564 | 0 | 0 | 0 | 0 | 4,286915 | 0 | 0 | 0 |
| 1238_TGTTACTTCTGAACGT | gdT | 0 | 0 | 0 | 1,524025 | 1,524025 | 0 | 0 | 1,524025 | 0 | 0 | 0 | 2,465723 | 2,941979 | 0 | 0 | 0 | 1,524025 | 2,465723 | 0 | 0 | 0 | 0 | 0 | 1,524025 | 0 | 0 | 3,506037 | 0 | 0 | 0 |
| 925_AGGCCACGTACTGACT | gdT | 0 | 0 | 0 | 1,015435 | 1,015435 | 0 | 0 | 1,015435 | 1,015435 | 1,508761 | 2,084709 | 2,28267 | 1,508761 | 0 | 0 | 1,015435 | 0 | 0 | 0 | 0 | 0 | 1,015435 | 1,837639 | 0 | 0 | 1,015435 | 3,310851 | 0 | 0 | 0 |
| 925_TAGGTACAGACATATG | gdT | 1,227629 | 0 | 0 | 0 | 0 | 0 | 0 | 0 | 0 | 2,884348 | 1,227629 | 2,569986 | 0 | 0 | 0 | 0 | 0 | 0 | 0 | 0 | 1,762374 | 0 | 0 | 0 | 0 | 1,762374 | 2,884348 | 0 | 0 | 0 |
| 925_TGCTTCGAGATCGCCC | gdT | 0 | 0 | 0 | 0 | 0 | 0 | 0 | 1,105729 | 0 | 1,105729 | 1,105729 | 3,14569 | 3,054674 | 0 | 0 | 1,105729 | 0 | 1,617972 | 0 | 0 | 0 | 1,955051 | 1,617972 | 0 | 0 | 0 | 3,565702 | 1,105729 | 0 | 0 |
| 945_GAGCTGCGTTATGGTC | gdT | 0 | 0 | 0 | 0 | 0 | 0 | 0 | 0 | 0 | 3,992089 | 0 | 2,659698 | 1,692061 | 0 | 0 | 0 | 0 | 1,692061 | 0 | 0 | 1,692061 | 1,692061 | 0 | 0 | 0 | 0 | 0 | 0 | 0 | 0 |
| 1167_ATTTACCCAAATGATG | gdT | 0 | 0 | 0 | 0 | 0 | 0 | 1,566387 | 0 | 0 | 0 | 0 | 2,149274 | 2,515108 | 0 | 0 | 0 | 0 | 1,566387 | 0 | 0 | 0 | 2,149274 | 0 | 0 | 0 | 0 | 2,515108 | 0 | 0 | 0 |
| 1235_ATGAAAGCAGAGGTAC | gdT | 0 | 0 | 0 | 1,673868 | 0 | 0 | 0 | 0 | 0 | 0 | 0 | 3,295708 | 2,268564 | 0 | 0 | 0 | 0 | 0 | 0 | 0 | 0 | 0 | 0 | 0 | 0 | 0 | 4,189523 | 0 | 0 | 0 |
| 1238_AGAGCCCTCACAATGC | gdT | 0 | 0 | 0 | 0 | 0 | 0 | 0 | 0 | 0 | 2,897845 | 0 | 2,061024 | 0 | 0 | 0 | 0 | 0 | 0 | 0 | 0 | 0 | 0 | 0 | 0 | 0 | 1,487723 | 3,460819 | 1,487723 | 0 | 0 |
| 966-2_CCGTGAGAGAGGGTAA | gdT | 0 | 1,604655 | 0 | 0 | 0 | 0 | 0 | 0 | 2,191909 | 3,605098 | 0 | 0 | 0 | 2,191909 | 0 | 0 | 0 | 0 | 0 | 0 | 0 | 0 | 0 | 0 | 0 | 0 | 2,191909 | 0 | 0 | 0 |
| 1235_TCAAGCAGTAGACTGG | gdT | 0 | 0 | 0 | 0 | 0 | 0 | 0 | 1,009078 | 1,009078 | 2,438725 | 0 | 0 | 0 | 1,82925 | 0 | 1,009078 | 0 | 0 | 0 | 0 | 0 | 1,500993 | 0 | 0 | 0 | 1,500993 | 3,163783 | 0 | 1,500993 | 0 |
| 1125_CTAAGTGTCTACGCAA | gdT | 0 | 0 | 0 | 0 | 0 | 0 | 0 | 0 | 0 | 3,348498 | 0 | 0 | 1,71859 | 1,71859 | 0 | 0 | 0 | 2,317801 | 0 | 0 | 0 | 1,71859 | 0 | 0 | 1,71859 | 0 | 2,689885 | 0 | 0 | 0 |
| 1224_ACGTCCTTCATTATCC | gdT | 1,049655 | 0 | 0 | 1,049655 | 0 | 0 | 0 | 1,550395 | 1,049655 | 0 | 0 | 3,224327 | 2,973827 | 0 | 0 | 1,049655 | 0 | 0 | 0 | 0 | 0 | 1,049655 | 0 | 0 | 0 | 0 | 4,217151 | 0 | 0 | 0 |
| 925_GTCCTCATCCATGATG | gdT | 0 | 0 | 0 | 0 | 0 | 0 | 0 | 0 | 0 | 2,564211 | 0 | 0 | 0 | 0 | 0 | 0 | 0 | 0 | 0 | 0 | 0 | 2,564211 | 0 | 0 | 0 | 2,564211 | 2,564211 | 0 | 0 | 0 |
| 934_GTGACGCCAATGAACA | gdT | 0 | 1,328524 | 0 | 0 | 0 | 0 | 1,328524 | 1,328524 | 0 | 1,328524 | 0 | 1,328524 | 2,493359 | 0 | 0 | 0 | 1,328524 | 0 | 0 | 0 | 0 | 1,328524 | 1,328524 | 0 | 2,493359 | 1,328524 | 3,984043 | 0 | 0 | 0 |
| 966_TCAGTTTTCGCCAGAC | gdT | 2,843817 | 0 | 0 | 0 | 0 | 0 | 0 | 0 | 0 | 3,229691 | 0 | 0 | 0 | 2,207242 | 0 | 0 | 0 | 0 | 0 | 0 | 0 | 0 | 0 | 0 | 0 | 0 | 0 | 0 | 0 | 0 |
| 1224_AGCATCAGTGGCTTAT | gdT | 0 | 0 | 0 | 0 | 0 | 0 | 0 | 0 | 0 | 0 | 0 | 0 | 2,089348 | 0 | 1,512888 | 0 | 0 | 2,718614 | 0 | 0 | 0 | 0 | 0 | 0 | 3,492208 | 0 | 3,59452 | 0 | 0 | 0 |
| 1224_CATGGTAAGCTGAAAT | gdT | 0 | 0 | 0 | 0 | 0 | 0 | 0 | 0 | 0 | 0 | 0 | 0 | 2,604621 | 0 | 0 | 0 | 0 | 1,9828 | 0 | 0 | 0 | 1,127421 | 0 | 0 | 2,235455 | 0 | 2,873647 | 0 | 0 | 0 |
| 1235_TCAGTGATCCTTATAC | gdT | 0 | 0 | 0 | 0 | 0 | 0 | 0 | 0 | 2,007035 | 3,96138 | 0 | 0 | 0 | 1,43998 | 0 | 1,43998 | 0 | 0 | 0 | 0 | 0 | 1,43998 | 0 | 0 | 0 | 0 | 2,630625 | 0 | 0 | 0 |
| 1238_CGGGCATAGCACCGTC | gdT | 0 | 0 | 0 | 0 | 0 | 0 | 0 | 0 | 0 | 0 | 0 | 3,023436 | 2,177557 | 1,591755 | 0 | 0 | 1,591755 | 1,591755 | 0 | 0 | 0 | 2,544518 | 0 | 0 | 0 | 0 | 3,197619 | 0 | 0 | 0 |
| 1066_GCCGTGACAATCGTCA | gdT | 0 | 0 | 0 | 0 | 0 | 0 | 0 | 0 | 0 | 3,517249 | 0 | 0 | 0 | 1,533075 | 0 | 1,533075 | 0 | 0 | 0 | 0 | 0 | 1,533075 | 0 | 0 | 0 | 0 | 2,952929 | 0 | 1,533075 | 0 |
| 1195_ACATTTCAGGGAGGAC | gdT | 0 | 0 | 0 | 0 | 0 | 0 | 1,469679 | 2,401844 | 0 | 0 | 0 | 2,875777 | 2,875777 | 0 | 0 | 0 | 0 | 0 | 0 | 0 | 0 | 2,040659 | 0 | 0 | 0 | 0 | 4,058947 | 1,469679 | 0 | 0 |
| 1235_TATACCTTCTCACCCA | gdT | 0 | 0 | 0 | 0 | 0 | 0 | 0 | 0 | 1,726712 | 3,435403 | 0 | 2,8436 | 0 | 1,726712 | 0 | 1,726712 | 0 | 0 | 0 | 0 | 0 | 1,197271 | 0 | 0 | 0 | 0 | 2,699105 | 0 | 1,197271 | 0 |
| 1238_AGTTAGCTCTTAGCTT | gdT | 0 | 0 | 0 | 0 | 1,629578 | 0 | 0 | 0 | 0 | 0 | 1,629578 | 3,391553 | 3,242996 | 0 | 0 | 0 | 1,629578 | 0 | 0 | 0 | 0 | 0 | 0 | 2,219578 | 0 | 0 | 3,916427 | 0 | 0 | 0 |
| 801_GCTGGGTCAAAGACTA | gdT | 0 | 0 | 0 | 0 | 0 | 0 | 0 | 0 | 1,617162 | 1,617162 | 0 | 0 | 0 | 0 | 0 | 0 | 0 | 0 | 0 | 0 | 0 | 0 | 0 | 0 | 0 | 0 | 0 | 0 | 0 | 0 |
| 945_GAAACCTTCACACCCT | gdT | 0 | 0 | 0 | 2,024613 | 0 | 1,683083 | 0 | 0 | 0 | 0 | 2,024613 | 2,649457 | 3,383841 | 0 | 0 | 0 | 1,160353 | 0 | 0 | 0 | 0 | 1,160353 | 0 | 0 | 0 | 0 | 3,522409 | 0 | 0 | 0 |
| 996_CTCATGCGTGGACAGT | gdT | 0 | 0 | 0 | 0 | 0 | 0 | 1,5014 | 0 | 0 | 0 | 2,439203 | 1,5014 | 3,580171 | 0 | 0 | 0 | 0 | 1,5014 | 0 | 0 | 0 | 2,439203 | 0 | 0 | 0 | 0 | 4,518382 | 0 | 0 | 0 |
| 898_GCAACATCAAACCACT | gdT | 0 | 0 | 0 | 0 | 0 | 0 | 0 | 0 | 0 | 0 | 0 | 2,188736 | 3,209706 | 0 | 0 | 1,601802 | 0 | 0 | 0 | 0 | 0 | 2,188736 | 0 | 0 | 0 | 0 | 3,704252 | 0 | 0 | 0 |
| 996_TATTCCAAGGACGCTA | gdT | 1,718715 | 0 | 0 | 0,926552 | 0 | 0 | 0 | 0 | 0 | 1,960537 | 0 | 0,926552 | 2,457921 | 0,926552 | 0 | 0,926552 | 0 | 0 | 0 | 0 | 0 | 0 | 0 | 0 | 0 | 0 | 3,667264 | 0 | 0 | 0 |

| Score | | | | | | |
|---|---|---|---|---|---|---|
| Barcode | cell_type | Exhausted | pre-exhausted | Naive | effector-memory | Resident |
| 1066_GTGAGGACACAAGGTG | abT | 1,334852413 | -0,122349977 | -0,535360187 | 1,087109179 | 0,368755254 |
| 1195_AGAAATGGTGGTGATG | abT | -0,077947844 | -0,455787645 | -0,50754744 | -0,224078057 | -0,312617886 |
| 1224_CGAAGTTGTAGCACAG | abT | 0,241103127 | -0,272733524 | -0,354436711 | 0,835285504 | 0,311628506 |
| 1224_GGGATCCCACAGAGCA | abT | -0,259803944 | -0,257092772 | -0,082070968 | 0,343240804 | -0,17177495 |
| 1238_AATGGCTGTCACTCTC | abT | -0,147811718 | 0,452933281 | 0,423195909 | 0,660618037 | -0,279127702 |
| 1238_ACCAACAGTATTTCTC | abT | -0,212463374 | -0,280450205 | 0,006204369 | 0,449249549 | -0,108392291 |
| 1238_AGGAATAGTCACTTCC | abT | 0,079724165 | -0,223588732 | 0,045513061 | 0,79061959 | -0,04214554 |
| 1238_GATCACATCCACCTCA | abT | 0,040639343 | 0,752838881 | -0,478491569 | 0,613925907 | 0,381181693 |
| 1238_TCCATGCGTAGCTGTT | abT | -0,049071853 | -0,156650597 | -0,234194993 | 0,554547457 | 0,157645823 |
| 1416_CAACCAAAGGACATCG | abT | -0,372028979 | -0,359980675 | -0,213541372 | -0,345183055 | 0,252777885 |
| 925_CTTGATTGTTCTCCAC | abT | -0,186337718 | 0,462869554 | 0,396032974 | 0,766763166 | 0,558313482 |
| 925_GCCGTGACAAGTGGTG | abT | -0,353551432 | -0,215606148 | -0,001528039 | -0,203026562 | -0,261332527 |
| 925_TATTTCGTCGTGTCAA | abT | -0,159366401 | -0,109278881 | -0,124232469 | 0,643039356 | 0,523289582 |
| 943_AGTAACCCACCGCTGA | abT | -0,157705265 | -0,166997318 | -0,25818439 | 0,218129264 | 0,167776862 |
| 943_CCTCCAAGTGACTATC | abT | -0,10261222 | -0,206833621 | -0,229500465 | 0,325565428 | -0,180029558 |
| 898_CATCCGTTCTCATTTG | abT | 0,435410669 | 0,392391161 | -0,36108976 | 1,174434174 | 0,00939746 |
| 996_TACAACGAGCGCTTCG | abT | -0,490557241 | -0,272139263 | 1,215353137 | 0,024837037 | -0,370158479 |
| 1238_GGTAATCCATAATGCC | abT | 0,341791465 | -0,219489341 | -0,259576969 | 0,752049371 | 0,015602975 |
| 925_GCCATGGCACAGTGAG | abT | 0,065795149 | -0,104090255 | -0,139377025 | 0,576374089 | -0,12166068 |
| 1224_TTTCAGTAGGCACCAT | abT | 0,564584583 | -0,218726746 | -0,316635827 | 0,586461861 | 0,134560474 |
| 1238_GAGACTTCAACACAGG | abT | 0,024595688 | 0,578015431 | -0,272975256 | 0,697392744 | 0,889047706 |
| 1238_GTAGAGGAGTGAGCCA | abT | -0,245328693 | -0,228046622 | -0,046071253 | 0,758432245 | 0,182055757 |
| 831_TACAACGTCAGCTCTC | abT | -0,072801485 | -0,548543872 | -0,141742385 | 0,147767569 | 0,571742968 |
| 1238_GCAGCCATCAGACATC | abT | -0,232270705 | 0,445626621 | 0,493717148 | 0,549330341 | 0,531326216 |
| 934_GATGGAGGTAACGGTG | abT | -0,071672131 | -0,135789917 | 0,273321542 | 0,780172513 | 0,049387116 |
| 898_TACGCTCCAATAGGGC | abT | -0,064637169 | -0,158331262 | -0,210018861 | 0,252633033 | -0,193685441 |
| 1130_AACCCAATCACTTTGT | abT | -0,343826277 | -0,36021691 | -0,399630373 | 0,387247751 | 0,541689174 |
| 925_GCCCGAAGTTGTCCCT | abT | -0,179879794 | -0,121155528 | 0,407269009 | 0,406936563 | -0,136254915 |
| 1238_GTGAGCCTCGGTAACT | abT | -0,305704338 | -0,129573046 | -0,260266353 | 0,676814988 | 0,184803858 |
| 1224_GTCTGTCTCGTAGGAG | abT | -0,326538858 | 0,381586997 | 0,981222606 | -0,074879492 | -0,346238526 |
| 1195_AAGGTAACACACGGTC | abT | -0,232793302 | -0,19860204 | -0,231954965 | -0,009227872 | 0,046236249 |
| 1224_CCTCAGTAGGTCGACA | abT | -0,467845316 | -0,337466111 | 0,164779392 | -0,165787612 | 0,106837183 |
| 1066_CTGCATCCAATAGAGT | abT | -0,270560037 | -0,158775761 | 0,591550799 | 0,314572148 | -0,034611481 |
| 1224_GCAGGCTCATGAGATA | abT | -0,371272644 | -0,231861782 | -0,054827818 | 0,726687069 | 0,351805938 |
| 1235_GTACAACAGGAGAATG | abT | -0,016604617 | -0,143207291 | -0,20431776 | 0,403547435 | -0,20916989 |
| 1416_TTTATGCGTGCCCTTT | abT | -0,339678596 | -0,171945666 | -0,269414561 | 0,331546499 | 0,64944993 |
| 898_CATCCACCAGACTCTA | abT | -0,507712512 | 0,093203261 | 0,004316436 | 0,381821424 | -0,019950126 |
| 801_GAAACCTTCGTGAGAG | abT | -0,39215973 | -0,324985398 | -0,062752674 | 0,27527618 | -0,124065734 |
| 1028_CTAGACAAGGTAGTAT | abT | -0,127706667 | -0,166810844 | -0,145416951 | 0,909204961 | -0,140402877 |
| 1195_CTGATCCCAAGGTCTT | abT | -0,185052796 | -0,250236614 | -0,014903632 | 0,436470048 | -0,012302691 |
| 1238_CAGGCCACAAAGACTA | abT | -0,254349656 | -0,331220061 | -0,39328093 | -0,257118531 | -0,400794672 |
| 996_TGAGTCAAGGTCGTAG | abT | 0,440836395 | -0,255952401 | -0,361812774 | 0,437901673 | -0,350855104 |
| 1355_ACCCACTAGTATGACA | abT | -0,07355189 | -0,020712597 | -0,057473391 | 0,182946713 | -0,057473391 |
| 925_AAACCCAGTGTTCCAA | abT | 0,064740901 | -0,133239396 | -0,168216719 | 0,671532649 | 0,321008706 |
| 925_CCTAAGATCGAGATGG | abT | 0,219983564 | -0,189989828 | -0,244495587 | 0,851737696 | 0,061087145 |
| 801_AAACGCTTCGGCCCAA | abT | 0,05220043 | 0,457643751 | 0,032921349 | 0,657613912 | 0,171392017 |
| 996_GTGATGTAGGTGATAT | abT | 0,040402139 | -0,279284725 | 0,468529327 | -0,245529831 | -0,128883991 |
| 1224_CTAACTTGTCACTGAT | abT | -0,199477838 | -0,159238742 | 1,409375419 | 0,171394336 | -0,024981004 |
| 1416_GTGTGATCACACGCCA | abT | -0,327690389 | -0,222848683 | -0,322270845 | 0,358399673 | 0,653236017 |
| 925_GGCTTGGGTTTCACTT | abT | 0,267928013 | -0,073687449 | -0,106387608 | 0,386738006 | -0,098816286 |
| 1130_ATTCTACCACCAGCTG | abT | 0,402826584 | 0,568600486 | -0,232252274 | 0,935616086 | -0,204960672 |
| 925_GTCATCCCAGCACCCA | abT | -0,00460847 | -0,206175377 | 0,22145806 | 0,545285746 | 0,350912126 |
| 943_CCCTTAGGTTCTGACA | abT | -0,235994376 | -0,143260874 | 0,122955603 | 0,147270537 | -0,165664091 |
| 966_AACAAAGAGCTTTCTT | abT | 0,183140667 | -0,087560179 | -0,161601133 | 0,810310769 | 0,16807599 |
| 1238_CCCTCAATCGGCATAT | abT | -0,422477933 | -0,392141761 | -0,001594113 | -0,175730703 | 0,222685621 |
| 1238_CTGTATTGTCCGGACT | abT | -0,399754369 | 0,395285055 | -0,353453621 | 0,392245401 | 0,195161154 |
| 1195_CCTCACAAGGATATGT | gdT | -0,100696731 | -0,19890153 | 0,189342914 | 0,812325977 | -0,08186083 |
| 1238_GGGCTACAGTAGTGCG | gdT | -0,289659514 | -0,162087502 | -0,240636916 | 0,903715723 | 0,897917554 |
| 1397_TGCGATACAGAACGCA | gdT | 0,193006795 | 0,128662079 | -0,36646232 | 0,464853109 | -0,060039997 |
| 1416_CGACAGCAGATGGGCT | gdT | -0,358659544 | 0,861341852 | -0,493025058 | 0,517727474 | 0,533904417 |
| 1416_GTCTTTAGTTTCAGAC | gdT | -0,035447665 | -0,161566439 | -0,229408294 | 0,297387793 | 0,50502288 |
| 943_GTTATGGCAGCCTATA | gdT | 0,179858477 | -0,208577282 | -0,187683616 | 0,629620594 | -0,155204106 |
| 1125_GGTAGAGGTTTCCCAC | gdT | 0,214206544 | -0,115582916 | -0,174195097 | 1,002333661 | 0,636594512 |
| 1066_GGTAGAGCACGACTAT | gdT | 0,09696155 | -0,22737036 | -0,241297318 | 0,801448039 | -0,02512646 |
| 1167_AAGTTCGGTCCTTTGC | gdT | -0,232768802 | 0,721187503 | -0,215813364 | 0,355100533 | 0,123663065 |
| 1195_TAGAGTCAGCGGTAAC | gdT | 0,633321265 | -0,192456759 | -0,249438014 | 0,86743684 | 0,812244007 |
| 1224_AGTCATGCAATTGTGC | gdT | -0,119798578 | 0,482586345 | -0,251693549 | 0,864626338 | 0,233015833 |
| 1224_CATACAGGTGGACCAA | gdT | 0,263037947 | -0,224814139 | 0,090482578 | 0,907111219 | -0,175446539 |
| 1238_AATCACGTCCATAGGT | gdT | 0,533344219 | -0,18763242 | -0,266762606 | 0,60288517 | 0,724827886 |
| 1238_GACGCTGTCCGCTTAC | gdT | -0,258787834 | 0,676268675 | -0,24024041 | 0,907209309 | -0,208298941 |
| 1355_TGGCCAGTCTTCGGTC | gdT | 0,189150419 | -0,041706524 | -0,056632447 | 1,098020619 | 0,610671935 |
| 1130_CCTCCAATCCTACCGT | gdT | 0,380319025 | 0,382248198 | 0,381465704 | 0,935985738 | -0,042372747 |
| 1416_AATGCCAGTACAGCGA | gdT | -0,189729729 | 0,321518155 | -0,359838286 | 0,3646926 | 0,010276802 |
| 1224_GTCTCACTCACACCCT | gdT | -0,359447169 | -0,368615832 | -0,209643387 | 0,214447583 | -0,277844291 |
| 1224_GTGAGCCGTAGTAAGT | gdT | 0,242216074 | -0,187563117 | -0,224997131 | 0,312104379 | -0,227003284 |
| 1238_ACGATGTTCACGAACT | gdT | -0,320423357 | -0,252402859 | -0,031041967 | 0,551018023 | 0,477171384 |
| 1238_TGTTACTGTTCCTAAG | gdT | -0,052222765 | -0,221586376 | -0,225737302 | 0,621194583 | -0,184618184 |
| 1238_TGTTACTTCTGAACGT | gdT | 0,6793636 | 0,503985544 | -0,256818713 | 0,609854471 | 0,051838431 |
| 925_AGGCCACGTACTGACT | gdT | -0,03945058 | -0,287008068 | -0,080801397 | 0,633099918 | 0,594019729 |
| 925_TAGGTACAGACATATG | gdT | -0,080169142 | 0,428242811 | -0,193280096 | 0,387621761 | 0,333171344 |
| 925_TGCTTCGAGATCGCCC | gdT | 0,013233551 | -0,212908389 | -0,330906159 | 0,991844852 | 0,089492261 |
| 945_GAGCTGCGTTATGGTC | gdT | 0,212894758 | -0,106865655 | -0,161665327 | 0,612824121 | -0,128779445 |
| 1167_ATTTACCCAAATGATG | gdT | -0,047700719 | -0,152096604 | -0,248174218 | 0,525968378 | 0,035783652 |
| 1235_ATGAAAGCAGAGGTAC | gdT | -0,027403302 | -0,083119734 | -0,155458409 | 0,564939293 | -0,163410691 |
| 1238_AGAGCCCTCACAATGC | gdT | -0,293796504 | -0,263924743 | -0,230165922 | 0,597915843 | 0,056247255 |
| 966-2_CCGTGAGAGAGGGTAA | gdT | -0,168356819 | -0,123160188 | 1,343570712 | 0,266851926 | -0,164245194 |
| 1235_TCAAGCAGTAGACTGG | gdT | -0,221531974 | -0,255563252 | 0,743381022 | 0,264059275 | 0,067926699 |
| 1125_CTAAGTGTCTACGCAA | gdT | 0,115687393 | -0,12680039 | 0,218855344 | 0,632921213 | -0,233706664 |
| 1224_ACGTCCTTCATTATCC | gdT | -0,115763664 | 0,181737699 | -0,208893326 | 0,518824437 | -0,259838894 |
| 925_GTCCTCATCCATGATG | gdT | -0,242720295 | -0,191310207 | -0,246119535 | 0,372667914 | 0,215438468 |
| 934_GTGACGCCAATGAACA | gdT | 0,274841428 | 0,403809988 | 0,05839005 | 0,906107999 | 0,227412644 |
| 966_TCAGTTTTCGCCAGAC | gdT | -0,580757728 | 1,045533888 | 0,153152121 | -0,192723871 | -0,438137739 |
| 1224_AGCATCAGTGGCTTAT | gdT | 0,734452119 | -0,248866605 | -0,29379883 | 0,508953955 | -0,225751127 |
| 1224_CATGGTAAGCTGAAAT | gdT | 0,194859912 | -0,244412016 | -0,305787956 | 0,386110004 | -0,271774275 |
| 1235_TCAGTGATCCTTATAC | gdT | -0,368957054 | -0,205557822 | 0,61773694 | 0,401646605 | -0,014720555 |
| 1238_CGGGCATAGCACCGTC | gdT | -0,009246411 | 0,639778568 | 0,175812455 | 0,658573415 | -0,162830867 |
| 1066_GCCGTGACAATCGTCA | gdT | -0,311108257 | -0,229775201 | 0,515647497 | 0,389880679 | 0,081186914 |
| 1195_ACATTTCAGGGAGGAC | gdT | -0,01782335 | -0,17720558 | -0,231556043 | 0,84417351 | 0,017197049 |
| 1235_TATACCTTCTCACCCA | gdT | -0,387965359 | -0,194549868 | 0,890187699 | 0,529561629 | 0,017419154 |
| 1238_AGTTAGCTCTTAGCTT | gdT | -0,02250084 | 0,649080819 | -0,220007804 | 0,803891212 | 0,337982127 |
| 801_GCTGGGTCAAAGACTA | gdT | -0,432255887 | -0,404092752 | -0,056270479 | -0,228623622 | -0,394165641 |
| 945_GAAACCTTCACACCCT | gdT | -0,167163151 | 0,303487175 | -0,410312269 | 0,470886858 | 0,238227829 |
| 996_CTCATGCGTGGACAGT | gdT | -0,172931637 | -0,289343515 | -0,347628644 | 0,633701643 | 0,323601884 |
| 898_GCAACATCAAACCACT | gdT | -0,196388199 | -0,172196122 | -0,201915488 | 0,690989003 | 0,120126922 |
| 996_TATTCCAAGGACGCTA | gdT | -0,274473897 | 0,607195706 | -0,081942352 | 0,376119211 | -0,145917697 |
| average abT | | -0,07929738 | -0,086101106 | -0,027771141 | 0,422973835 | 0,082723578 |
| average gdT | | -0,027456151 | 0,033000903 | -0,064245409 | 0,575303815 | 0,088393642 |
