## Supplemental Material and Methods for "EphA2 and Phosphoantigen-Mediated Selective Killing of Medulloblastoma by γδT Cells Preserves Neuronal and Stem Cell Integrity"

| TARGET | CONJUGATED | CLONE | SUPPLIER |
| --- | --- | --- | --- |
| CD107a | PerCP-Cy5.5 | #H4A3 | Biolegend |
| pan γδ TCR | FITC | #IMM510 | Beckman Coulter |
| Vδ2 | BV421 | #B6 | Biolegend |
| Vδ1 | APC | #REA173 | Miltenyi Biotec |
| CD69 | PE | #FN50 | Biolegend |
| CD1d | PE | #51.1 | Biolegend |
| CD1c | PE | #L161 | Biolegend |
| EphA2 | PE | #SHM16 | Biolegend |
| CD112 | PE | #TX31 | Biolegend |
| CD155 | PE | #SKII.4 | Biolegend |
| MICA-B | PE | #6D4 | BD Biosciences |
| ULBP2,5,6 | BV605 | #165903 | BD Biosciences |
| Mouse IgG2b isotype | PE | #MPC-11 | Biolegend |
| Mouse IgG1 isotype | PE | #MOPC-21 | Biolegend |
| Mouse IgG2a isotype | BV605 | #G155-178 | BD Biosciences |
| Mouse IgG2a isotype | PE | #G155-178 | BD Biosciences |
| NKG2D | Purified | #1D11 | Nordic BioSite |
| Annexin A2 | Purified | # 1C1E12 | Proteintech |
| Mouse IgG2a isotype | Purified | # 11A1B2 | Proteintech |
| Donkey anti-mouse | AF488 | polyclonal | Life Technologies |

**Supplemental Material & Methods:** list of antibodies used for Flow Cytometry analysis and blocking experiments.
